# Comparative Proteomic Analysis of Environmental and Genetic Models of Parkinson’s Disease Highlights the Role of Purine Metabolism

**DOI:** 10.64898/2026.02.02.703282

**Authors:** Pablo-Reina Gonzalez, Müberra Fatma Cesur, Aiesha Anchan, Abdulla Abu-Salah, Tunahan Çakır, Emir Malovic, Souvarish Sarkar

## Abstract

Parkinson’s Disease (PD) is the second most common neurodegenerative disease, with many cases being attributed to environmental contaminant exposures. Paraquat (PQ), is a pesticide and environmental neurotoxicant that has been strongly associated with increased risk of PD. PQ is known to be a weak inhibitor of complex I of the electron transport chain, and while its acute toxicity is well understood, the underlying mechanism by which PQ exposure contributes to PD pathophysiology remains unclear. Additionally, the mechanism of PQ neurotoxicity has yet to be effectively compared and related to genetic forms of PD. Given that PD is a heterogeneous disease with both genetic and environmental determinants, we sought to systematically compare the proteomic changes that occur in different genetic and environmental models of PD. In this study, we leveraged untargeted omics approaches to differentiate between systemic, peripheral, and CNS-specific changes in the proteome. We did this by performing a comparative proteomic analysis on the heads and bodies of *Drosophila* models of PQ ingestion and neuronal α-synuclein expression in males. Additionally, we validated the findings with metabolomic analysis of male and female brain stems from a murine PQ inhalation model using C57BL/6J mice. Our findings indicate shared dysregulated pathways across all models, highlighting similar mechanisms of action. Specifically, we identified a glia-specific role in purine nucleotide metabolism upstream of inosine catabolism, which may protect against PQ neurotoxicity. This work identifies potential early points for biomarker detection and potential targets for drug intervention.

**Significance Statement:** Neurodegenerative diseases such as Parkinson’s disease (PD) pose a growing public health burden, yet disease-modifying therapies remain limited due to lack of mechanistic understanding and disease heterogeneity. Both genetic and environmental factors contribute to PD, complicating the identification of shared therapeutic targets. Here, we identify a convergent pathway common to genetic and environmental models of Parkinsonism that not only affects the brain but also systemically. Using integrated metabolomics, proteomics, and genome-scale metabolic modeling, we demonstrate that purine metabolism is dysregulated across models. Reverse genetic screening of key enzymes in this pathway mitigates locomotor deficits induced by neurotoxic pesticide exposure in *Drosophila*. These findings reveal a shared metabolic vulnerability in PD and highlight purine metabolism as a potential therapeutic target.

## INTRODUCTION

Parkinson’s Disease (PD) is the fastest-growing neurodegenerative disease in the world, and the second most common disease behind Alzheimer’s disease (AD) ^1,2^. PD has both genetic and environmental contributors, with only around 10% of all cases attributed to Mendelian genetic inheritance ^3,4^. Many have postulated a multi-hit hypothesis of PD in which genetic mutations, which increase the risk for PD, are paired with exposure to environmental toxicants that target similar pathways in a complementary fashion ^5,6^. The summation of these risk factors acting on the same or related pathways can exacerbate the damage to dopaminergic neurons, a key pathophysiological marker of PD, causing them to die off at a faster rate than in normal aging.

Paraquat (PQ), a commonly used non-selective herbicide, is one such environmental neurotoxicant. PQ is known to inhibit complex I of the Electron Transport Chain (ETC) in humans and can cross the blood-brain barrier when inhaled or ingested ^7,8^. It is used to kill weeds and to clear green plant matter for soy and cotton during harvest season, facilitating the picking process. In humans, epidemiological studies have reported positive correlations between pesticide exposure and increased PD risk ^9–15^. While some studies have reported contradictory findings ^16^, recent work in the California Central Valley has identified clear associations between working near agricultural sites utilizing paraquat (PQ) as well as residential proximity to these sites, and increased risk of PD irrespective of sex (odds ratio = 2.08-2.15 and 1.72-1.91, respectively) ^17^.

PD is mostly studied as a disease of the central nervous system, but some of the early symptoms of PD are autonomic. Constipation is one of the most common non-motor symptoms of PD and can arise decades before motor symptoms manifest ^18–20^. Furthermore, recent studies have demonstrated that α-synuclein, a key pathological protein involved in the etiology of PD, can aggregate in the gut and propagate to the brain ^21–23^. Notably, gut microbiome dysbiosis has also been linked to PD ^23,24^, particularly due to the presence of enteric neurons and glial cells that could provide access to the CNS through retrograde transport ^25^. Hence, we hypothesize that PD is a systemic disorder that involves both the autonomic and central nervous system.

While many omics studies have highlighted the cellular signatures of the central nervous system in PD ^26,27^, the mechanistic commonalities between the brain and the body have been overlooked. Furthermore, while omics-based analyses of various genetic models have previously been conducted to identify potential mechanisms, exploring the mechanistic similarities between genetic and environmental factors has been challenging. Here, we sought to identify the mechanistic commonalities between genetic and toxicant-induced models of PD both in the brain and systemically. We achieved this by performing comparative proteomic analyses of a *Drosophila* model of PQ-induced neurotoxicity (as an environmental model) and the pan-neuronal α-synuclein model (nSyb-QF2 driver; as a genetic model) to identify convergent or divergent pathogenic pathways and potential therapeutic targets.

Using comparative proteomics, we have identified purine metabolism as a common dysregulated pathway in genetic and environmental models of PD in *Drosophila*. Furthermore, a cell-specific knockdown genetic screen of purine metabolism pathways in flies identified dysfunction in the inosine-metabolizing enzymes in glia as a potential driving factor in PQ-induced neurodegeneration. Finally, we validated our findings in flies in a rodent model of PQ toxicity by performing metabolomics on brains from a PQ inhalation model in C57BL/6J mice, revealing similar alterations in purine metabolism pathways in both male and female mice.

## METHODS

### Fly Husbandry

All flies were maintained at 25 °C in an incubator for crosses and aging. For our PQ group, adult flies were exposed to 5mM PQ through their food ^28–31^ or control food (Formula 4-24® Instant Drosophila Medium, Carolina Biological Supply Company, item#: 173200) starting 2–3 days following eclosion. The food was changed every three days to ensure it remained moist and fresh. Behavior assays were conducted following 7 days of exposure to measure changes in locomotor activity. Our α-synuclein group was generated using the previously described human *QUAS-wild-type α-synuclein* line, which contains both the *nSyb-QF2* and *nSyb-GAL4* drivers ^32,33^. This line was crossed with the *W-* line to eliminate the QS transcriptional suppressor in half of the resulting offspring. Flies lacking the QS construct were selected based on their darker eye color. Genetic controls were generated by crossing Syb-GAL4 with W-flies. This group was kept on normal, untreated food for 7-days prior to proteomics and metabolomics analyses.

### Drosophila Behavior Assays

#### Climbing Assay

Climbing assays were carried out as described previously ^33^. In brief, flies were transferred to empty vials and acclimated for 3 min. Flies were then tapped to the bottom of the vial, and the percentage that climbed beyond 5 cm up the side of the vial within 10 s was recorded. The assay was repeated three times per group.

#### Locomotor Assay

Locomotor assays were performed according to previously established protocols ^33^. Flies were transferred to empty vials and acclimated for 3 min. Flies were tapped to the bottom of the vial, and placed horizontally for 10 secs. The number of flies moving after the 10 secs was recorded. The assay was repeated three times per group.

### Animal Study

Male and female C57BL/6J mice were obtained from Jackson Laboratories (Strain #:000664) and arrived at postnatal week 12. They were pair-housed in standard mouse caging with ∼3 mm high-performance bedding (BioFresh), supplied with standard rodent chow (LabDiet Autoclavable Diet 5010) and water *ad libitum*, and maintained at 22°C ± 2°C under a 12-hour light-dark cycle (lights on at 06:00) at the University of Rochester Medical Center’s Inhalation Exposure Facility. At postnatal week 14, during early adulthood, male and female animals were randomly assigned to either the PQ or the filtered-air control groups. Within these two treatment groups, animals were further divided into either the behavioral or non-behavioral control groups, as behavioral enrichment has been shown to lessen or mitigate the effects of neurotoxicants ^34^. This allocation yielded sample sizes of n = 7-8 per treatment and per behavior group for males and females. However, only samples from the non-behavior mice were used for the purposes of this publication. These mice were sacrificed the evening of their final exposure day at 2 and 3 months (40 and 60 days of exposure), using CO_2_ and cervical dislocation, followed by cardiac perfusion with phosphate-buffered saline.

The animals were weighed every Friday from the beginning of exposure to the completion of the study to observe signs of systemic toxicity or excessive anxiety and depression. All mice used in this study were treated humanely and with consideration for their well-being and the alleviation of suffering. Mice that presented with consistent weight loss and signs of pain or discomfort were euthanized early to minimize suffering in accordance with university policies. All procedures were approved by the University Committee on Animal Resources (UCAR) at the University of Rochester.

#### Inhalation exposure paradigm

Whole-body inhalation exposures were conducted in the University of Rochester Inhalation Exposure Facility. Exposures were conducted 5 days a week for 12 weeks to mimic a typical work schedule for agricultural workers during the harvest season. Animals were exposed to either filtered air or PQ aerosols for 4 hours per day beginning at 08:00, during which time the mice were singly housed in wire-mesh chambers inside a 30-L steel-reinforced Lexan exposure chamber. The exposure chamber was maintained at 40–45% relative humidity and 22–24°C. PQ aerosols were produced by adding a 0.2 mg/mL solution of PQ in ddH2O (made fresh every day) into an ultrasonic nebulizer (Ultrasonic2000, Nouvag Dental and Medical Equipment). Clean, dry air (approximately 500 mL/min) was passed through the nebulizer at 2.4 MHz to produce a PQ mist, which then passed through a heated drying tube and a cold trap to remove moisture. The subsequent PQ particles were mixed with diluting air and passed into the exposure chamber at a rate of 25–30 L/min. Using a peristaltic pump (Masterflex, Cole Palmer Inc.), the PQ solution was slowly transferred into the nebulizer for the duration of the exposure (approximately 18 mL/h). A condensation particle counter (CPC Model 3022A, TSI Inc.) was used to capture real-time exposure measurements, enabling quantification of continuous particle counts. The particle size distribution of the aerosol was measured daily using an electrostatic classifier (SMPS Model 3071, TSI Inc.). Three filters (Pallflex Membrane Filters, Pall Life Science) were collected hourly, starting after the first 30 minutes of exposure and continuing through the last 30 minutes, to determine the gravimetric exposure concentration. The concentration of PQ on the filters was quantified chemically using a UV-Vis spectrophotometer (HP-Agilent Technologies) at 259 nm. Analytical standard grade PQ (PQ dichloride hydrate PESTANAL®, ≥98.0% purity) was purchased from Millipore Sigma (CAS No: 75365-73-0).

### Proteomics

This protocol was carried out in accordance with our group’s previous publications using proteomics ^35^.

#### Sample Preparation

Drosophila heads and bodies (n = 10 per sample) were homogenized by bead beating (Bead Ruptor Elite, Omni International) for 30 s at 6 m/s in 100 µL (heads) or 200 µL (bodies) of 5% SDS, 100 mM TEAB. Homogenates were centrifuged at 15,000 × g for 5 min to pellet debris, and supernatants were collected. Protein concentrations were determined using a BCA assay (Thermo Scientific). Samples were then diluted to 1 mg/mL in 5% SDS, 50 mM TEAB. For each sample, 25 µg of protein was reduced with 2 mM dithiothreitol at 55 °C for 60 min, followed by alkylation with 10 mM iodoacetamide in the dark at room temperature for 30 min. Next, phosphoric acid was added to 1.2%, and proteins were precipitated by adding six volumes of 90% methanol, 100 mM TEAB. The mixture was loaded onto S-Trap micros (Protifi) and centrifuged at 4,000 × g for 1 min. Bound proteins were washed twice with 90% methanol, 100 mM TEAB. Proteins were digested by adding 1 µg trypsin in 20 µL of 100 mM TEAB to each S-Trap, followed by an additional 20 µL TEAB to prevent drying. Samples were incubated overnight at 37 °C. The following morning, digested peptides were eluted by centrifugation at 4,000 × g for 1 min, followed by sequential elutions with 0.1% TFA in acetonitrile and 0.1% TFA in 50% acetonitrile. Eluates were pooled, frozen, and dried in a SpeedVac (Labconco). Dried peptides were resuspended in 0.1% TFA prior to mass spectrometry analysis.

#### Mass Spectrometry

Peptides were first loaded onto a 75 µm × 2 cm trap column (Thermo Fisher) and then separated on an Aurora Elite 75 µm × 15 cm C18 analytical column (IonOpticks) using a Vanquish Neo UHPLC (Thermo Fisher) coupled to an Orbitrap Astral mass spectrometer (Thermo Fisher). Solvent A consisted of 0.1% formic acid in water, and solvent B was 0.1% formic acid in 80% acetonitrile. Ionization was achieved with an Easy-Spray source operated at 2 kV. The LC gradient began at 1% B, ramped to 5% B in 0.1 min, increased to 30% B over 12.1 min, then to 40% B in 0.7 min, and finally to 99% B in 0.1 min. The column was washed at 99% B for 2 min, giving a total runtime of 15 min. Following each run, the column was re-equilibrated with 1% B prior to the next injection. The Orbitrap Astral was operated in data-independent acquisition (DIA) mode. MS1 scans were acquired in the Orbitrap at 240,000 resolution, with a maximum injection time of 5 ms, across an m/z range of 380–980. DIA MS2 scans were collected in the Astral mass analyzer with a maximum injection time of 3 ms using a variable windowing scheme: 2 Da windows for 380–680 m/z, 4 Da windows for 680–800 m/z, and 8 Da windows for 800–980 m/z. Higher-energy collisional dissociation (HCD) was performed at 25% collision energy, with normalized AGC set to 500%. Fragment ions were measured across 150–2000 m/z, with a total cycle time of 0.6 s.

#### Data Analysis

The data were not blinded prior to analysis. The raw data were processed with DIA-NN version 1.8.1 (https://github.com/vdemichev/DIA-NN) ^36^. For all experiments, data analyses were performed using library-free analysis mode in DIA-NN. To annotate the library, the *Drosophila melanogaster* UniProt ‘one protein sequence per gene’ database (UP000000803_7227, downloaded 4/27/2021) was used with ‘deep learning-based spectra and RT prediction’ enabled. For precursor ion generation, the maximum number of missed cleavages was set to 1, the maximum number of variable modifications to 1 for Ox(M), the peptide length range to 7-30, the precursor charge range to 2-4, the precursor m/z range to 380-980, and the fragment m/z range to 150-2000. The quantification was set to ‘Robust LC (high precision)’ mode with normalization set to RT-dependent, MBR enabled, protein inferences set to ‘Genes’, and ‘Heuristic protein inference’ turned off. MS1 and MS2 mass tolerances, along with the scan window size, were automatically set by the software. Precursors were subsequently filtered at library precursor q-value (1%), library protein group q-value (1%), and posterior error probability (20%). Protein quantification was carried out using the MaxLFQ algorithm as implemented in the DIA-NN R package (https://github.com/vdemichev/diann-rpackage), and the number of peptides quantified in each protein group was counted as implemented in the DiannReportGenerator Package (https://github.com/URMC-MSRL/DiannReportGenerator) ^37^.

### Metabolomics

#### Sample preparation

Frozen tissue was homogenized in a 40:40:20 mixture of acetonitrile, methanol, and water using a Precellys cold tissue homogenizer (Bertin) at a ratio of 15 mg striatum tissue or 20 mg brain stem tissue per 1 mL of solvent. After homogenization, samples were incubated at −20 °C for 30 minutes and then transferred to regular ice for an additional 30 minutes, with vortexing every 10 minutes throughout the incubation. Samples were then centrifuged at 17,000 × g for 10 minutes, after which the supernatants were collected and dried in a vacuum evaporator (Thermo Scientific). The resulting dried extracts were reconstituted in 50% acetonitrile (A955, Fisher Scientific) at a volume equivalent to 10% of the original dried-down volume and transferred to glass vials for LC/MS analysis.

#### LC/MS Analysis

Metabolite extracts were analyzed by high-resolution mass spectrometry using an Orbitrap Exploris 240 (Thermo) coupled to a Vanquish Flex liquid chromatography system (Thermo). A 2 µL injection was loaded onto a Waters XBridge XP BEH Amide column (150 mm length × 2.1 mm inner diameter, 2.5 µm particle size) maintained at 25 °C, with a Waters XBridge XP VanGuard BEH Amide guard column (5 mm × 2.1 mm inner diameter, 2.5 µm particle size) placed upstream. For positive mode acquisition, mobile phase A consisted of 100% LC–MS grade water containing 10 mM ammonium formate and 0.125% formic acid, while mobile phase B consisted of 90% acetonitrile containing 10 mM ammonium formate and 0.125% formic acid. For negative mode acquisition, mobile phase A was prepared as 100% LC–MS grade water containing 10 mM ammonium acetate, 0.1% ammonium hydroxide, and 0.1% medronic acid (Agilent), while mobile phase B contained 90% acetonitrile with the same additives. Chromatographic separation was performed with the following gradient: 0 minutes, 100% B; 2 minutes, 100% B; 3 minutes, 90% B; 5 minutes, 90% B; 6 minutes, 85% B; 7 minutes, 85% B; 8 minutes, 75% B; 9 minutes, 75% B; 10 minutes, 55% B; 12 minutes, 55% B; 13 minutes, 35% B; 20 minutes, 35% B; 20.1 minutes, 35% B; 20.6 minutes, 100% B; and 22.2 minutes, 100% B, all at a flow rate of 150 µL/min. At 22.7 minutes, the flow rate was increased to 300 µL/min and held at 100% B until 27.9 minutes, after which the flow was returned to 150 µL/min at 28 minutes, for a total run length of 28 minutes. The H-ESI source was operated in positive mode at a spray voltage of 3500 V or in negative mode at a spray voltage of 2500 V, with sheath gas set to 35 arbitrary units, auxiliary gas to 7 arbitrary units, and sweep gas to 0. The ion transfer tube temperature was maintained at 320 °C and the vaporizer temperature at 275 °C. Data were collected across a mass range of 70–1000 m/z with full scan MS1 at a resolution of 120,000 FWHM, RF lens set to 70%, and standard automatic gain control (AGC). Data-dependent MS2 fragmentation for compound identification was performed using the AcquireX workflow (Thermo Scientific), consisting of three deep scans with MS1 resolution at 60,000 and MS2 resolution at 15,000. LC–MS data were analyzed using Compound Discover v3.3 (Thermo Scientific) and El-Maven software for peak area determination and compound identification. Compounds were identified based on retention time matching to LC–MS method-specific external standards and MS2 spectral matching to both external standards and the mzCloud database (Thermo Scientific).

### Proteomics-Driven Metabolic Network Analyses

In the metabolic network analyses, the methodology described in Abu-Salah et al. 2024 was followed using the iDrosophila1 genome-scale metabolic model, which includes 8,230 reactions, 6,990 metabolites, and 2,388 genes ^38^. To align with the gene identifiers used in the iDrosophila1 model, FlyBase gene IDs were mapped to the proteomic datasets from each PD group (PQ_head_, PQ_body_, αSyn_head_, and αSyn_body_) and their corresponding control groups. Gene annotation was facilitated using resources from the UniProt ^39,40^ and FlyBase databases ^41,42^. More than 65% of the proteins in each proteomic dataset were successfully annotated with the FlyBase gene IDs. Next, two network-based approaches were employed to identify differential metabolic activities induced by the PQ treatment or α-synuclein expression. Both network-based analyses were conducted in MATLAB R2023a, utilizing Gurobi Optimizer version 11.0.3 (Gurobi Optimization, LLC) as the solver.

#### ΔFBA Approach

The first metabolic network-based approach, ΔFBA ^43^ aims to optimize the agreement between differential protein abundances and predicted changes in metabolic fluxes by maximizing consistencies and minimizing inconsistencies between them. To implement this method, the fold change values of significantly changed proteins relative to the corresponding controls (*p*-value < 0.05) were calculated using the limma-trend function from the limma package in R (version 4.3.0), with the parameter “robust = TRUE” ^44^. For each PD group, the significant fold changes were mapped to the iDrosophila1 reactions through *mapExpressionToReactions* in the COBRA Toolbox ^45^. Nongrowth-associated maintenance (NGAM) reaction flux was held constant by preventing its flux change, while other reaction boundaries remained unconstrained. Then, ΔFBA was run with the cut-off of 0.1 (representing the most upregulated and most downregulated 10% of reactions in terms of mapped fold changes) and three *ε* thresholds (1, 3, and 5; defining the magnitude of minimum flux change required to classify reactions as upregulated or downregulated), respectively. To ensure robustness, only reactions consistently classified as differential across at least two *ε* thresholds were finally retained for each PD group. The metabolites and genes involved in these differential reaction sets were extracted from the iDrosophila1 model and listed for further analysis.

#### Reaction Activity Analysis

The second approach used to simulate proteome-based metabolic shifts in PD was reaction activity analysis. This method involved reconstructing sample-specific metabolic models using the integrative Metabolic Analysis Tool (iMAT) algorithm ^46^, after applying minimal flux constraints to the fundamental reactions (i.e., 10⁻³ h⁻¹ for biomass production, 10⁻⁴ mmol g⁻¹ h⁻¹ for glucose and oxygen uptakes, and a 50% reduction in the maximum NGAM flux; originally set at 8.55 mmol ATP g⁻¹ h⁻¹) to ensure that they remain active. Protein abundance data were subsequently mapped to iDrosophila1 reactions using the *mapExpressionToReactions* function for the reconstruction of sample- and tissue-specific metabolic network models. In total, 28 models were generated: 14 for head tissues and 14 for body tissues, covering the PQ-treated and α-synuclein-expressing fly groups along with their relevant controls. During model reconstruction, the iMAT algorithm utilized the 25^th^ and 75^th^ percentiles of proteomics data to classify reactions as active or inactive by retaining the high-abundance reactions under mass-balance constraints ^47^. Then, binary vectors representing reaction activities (1 = active, 0 = inactive) were created for each reconstructed model. A reaction was defined as differential if its activity status consistently differed between PD and control models within the same experimental group (PQ_head_, PQ_body_, αSyn_head_, or αSyn_body_); specifically, it was active in more than half of the PD models and present at negligible levels (0 or 1 occurrence) in the corresponding control models, or vice versa. The genes and metabolites associated with the differential reactions were finally listed.

### Inferring Pathway-Level Insights from ΔFBA- and iMAT-Based Predictions

To characterize the metabolic effects of PQ treatment and α-synuclein expression in flies, the gene and metabolite lists compiled in the previous section from either ΔFBA or iMAT-based reaction activity analysis were merged for each PD group. This integration aimed to capture complementary information from both network-based approaches, as highlighted in Abu-Salah et al. ^35^.

The resulting gene lists were analyzed using PANGEA (PAthway, Network and Gene-set Enrichment Analysis) to uncover significantly enriched KEGG pathways at a false discovery rate (FDR) threshold of 0.05 ^48^. Pathway enrichment analyses were also conducted on the metabolite lists using MetaboAnalyst 6.0 ^49^. In this step, KEGG compound IDs were submitted to the online tool, and the organism was set to *Homo sapiens* to align the results with PD–related metabolic pathways. Pathways were considered significantly enriched if they met the criteria of *p*-value < 0.05 and pathway impact > 0.10 ^50,51^. The heatmaps illustrating key pathway enrichment results were generated via the *SHeatmap* function in MATLAB ^52,53^.

## RESULTS

### Proteomic analysis reveals differential regulation of purine metabolism pathways in PQ-treated Drosophila heads and bodies

PD is characterized by changes not only in the brain but also systemically. To identify commonalities in mechanistic changes between the brain and the body, we first performed quantitative proteomics on the heads and bodies of flies fed potato-starch feed dosed with 5 mM PQ (or control food) post-eclosion for 7 days *ad libitum*.

The data from the heads and bodies of PQ-exposed flies were neatly segregated, indicating separate protein-expression profiles in the treatment groups (Figure 1A, Figure 2A). Our differential expression analysis between treated and control groups revealed that 755 proteins were significantly downregulated and 485 were significantly upregulated in the heads of PQ-exposed flies (Figure 1B). The bodies of these same conversely had 329 proteins significantly downregulated and 449 significantly upregulated (Figure 2B).

**Figure 1.**
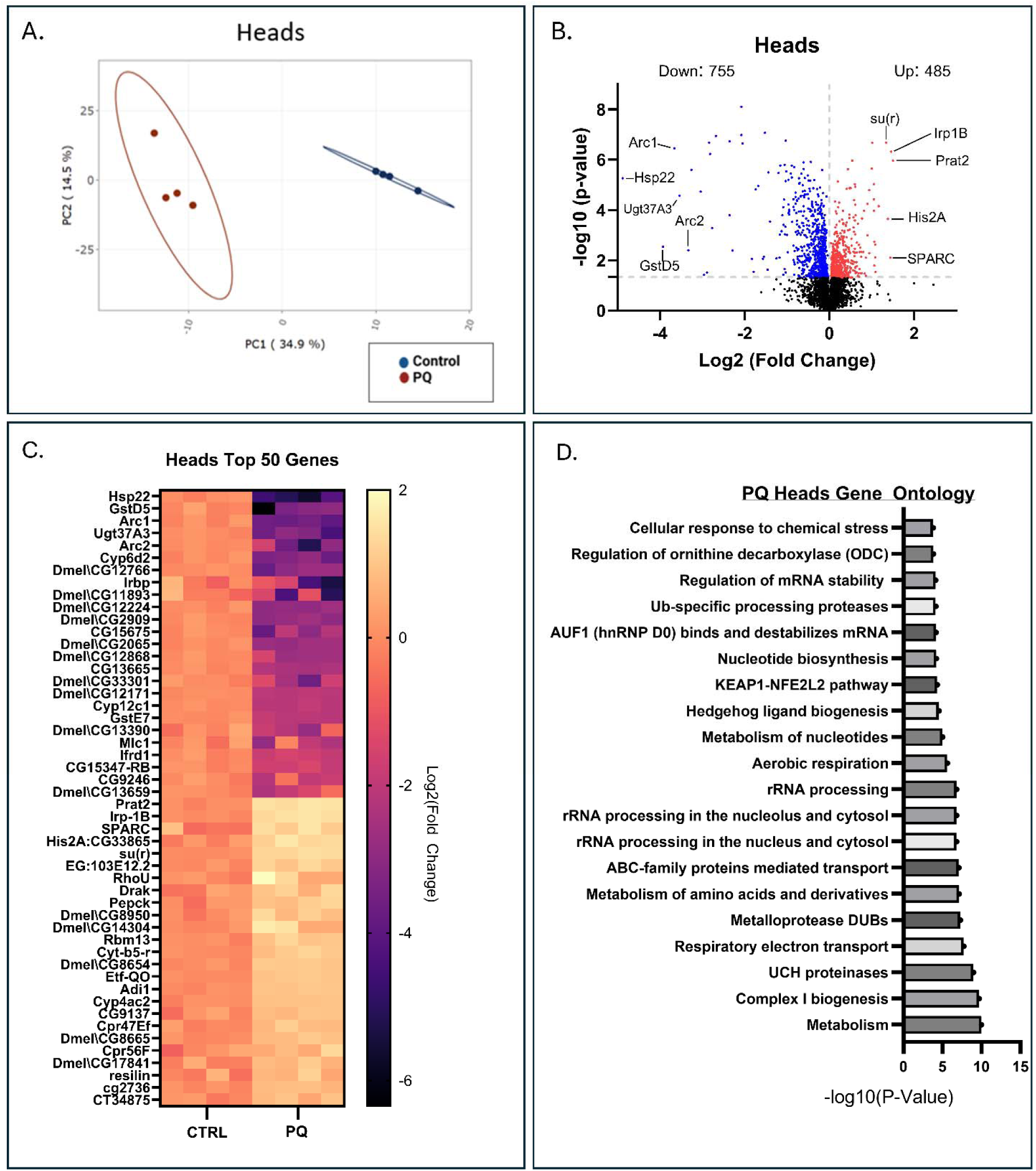
Proteomics analysis of PQ-treated *Drosophila* heads identified differentially regulated pathways. (A) PCA plot of proteomics data from PQ-treated *Drosophila* heads and no treatment controls, demonstrating neat segregation of treatment groups and good separability. (B) Volcano plot showing the most significant up- and down-regulated proteins in the proteomics dataset. (C) Heatmap of the 50 most statistically significant protein expression changes between heads of PQ-treated *Drosophila* heads and control heads, colored based on Log_2_ fold change. The first 25 genes correspond to the most down-regulated proteins, while the next 25 are the most up-regulated. (D) Bar graph of gene ontology analysis of the 20 most statistically significant differentially regulated pathways as identified by the DAVID Functional Annotation Bioinformatics Microarray Analysis.

**Figure 2.**
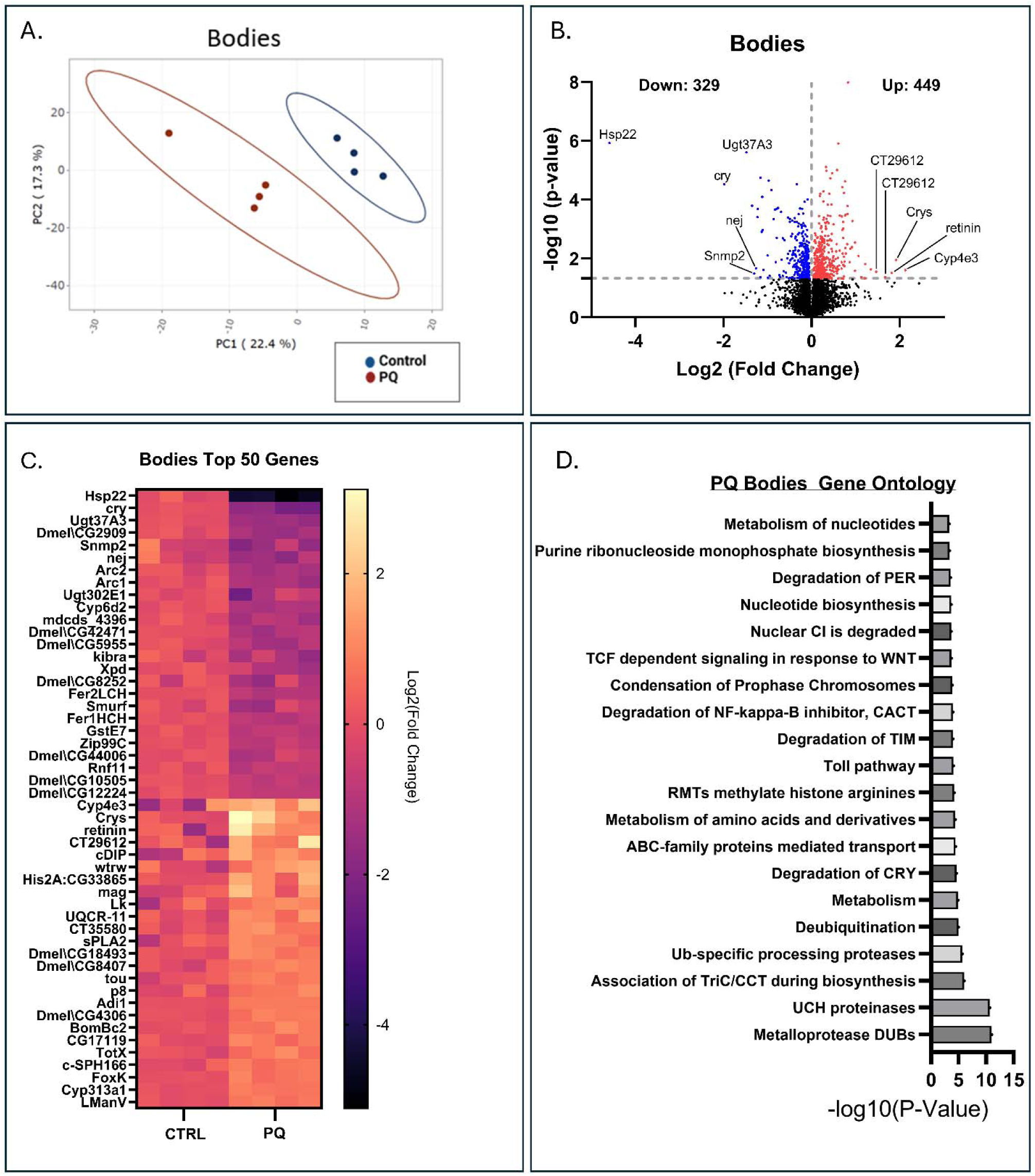
Proteomics analysis of PQ-treated *Drosophila* bodies identified differentially regulated pathways. (A) PCA plot of proteomics data from PQ-treated *Drosophila* bodies and controls, demonstrating separate clustering of groups, ensuring distinct protein expression profiles. (B) Volcano plot showing the most significant up- and down-regulated proteins in the proteomics dataset. (C) Heatmap of the 50 most statistically significant protein expression changes between the PQ-treated *Drosophila* bodies and control bodies, colored based on Log_2_ fold change. The first 25 genes correspond to the most down-regulated proteins, while the next 25 are the most up-regulated. (D) Bar graph of gene ontology analysis of the 20 most statistically significant differentially regulated pathways as identified by the DAVID Functional Annotation Bioinformatics Microarray Analysis.

Gene ontology (GO) analysis of the PQ-exposed *Drosophila* heads showed most of these genes were broadly associated with cellular metabolism, specifically with respiratory electron transport and biogenesis of complex-I of the ETC (Figure 1D), which are well-established primary cellular targets of PQ ^54–56^. One key pathway shown to be altered in the heads was nucleotide biosynthesis, which may relate to declines in uric acid, which have been consistently correlated with an increased risk of PD and with faster disease progression in human patients ^57–59^. Other diverse pathways broadly associated with PD were also altered, including ubiquitin-specific processing proteases and ubiquitin carboxyl-terminal hydrolase (UCH) proteinases, which are both involved in protein degradation and turnover. These functions are notoriously deficient in genetic forms of PD where specific gene mutations in the proteosome can cause or increase the risk of disease incidence (e.g., PARK2/Parkin, PARK5/UCH-L1, PARK15/FBXO7) ^60–63^. Similarly, in idiopathic PD, the ubiquitination of certain lysine residues on α-synuclein can induce structural changes that promote the formation of protein inclusions ^64,65^, and defects in the 26S proteasome are similarly associated with increased formation of inclusions ^66,67^. The GO analysis of the bodies similarly highlighted enrichment in pathways associated with “metabolism of nucleotides”, “nucleotide biosynthesis”, “purine ribonucleoside monophosphate biosynthesis”, and general cellular metabolism (Figure 2D). Proteasomal pathways were also notably dysregulated in the bodies, as indicated by the enrichment of GO terms related to “ubiquitin-specific processing proteases”, “UCH proteinases”, and “deubiquitination”.

### Comparative gene ontology analysis of proteomics data from *Drosophila* α-synuclein and PQ models supports purine metabolism dysregulation as a key overlapping pathway in Parkinsonism pathogenesis

Since α-synuclein aggregation is a hallmark of sporadic PD and most genetic PD cases, to uncover the shared pathogenic mechanisms of the α-synuclein and PQ *Drosophila* models, we performed comparative analysis on the proteomics data sets using ClueGO ^68^, a Cytoscape plugin that creates gene ontology and pathway networks from gene lists using public databases. First, we compared the four data sets to identify similarities and protein co-occurrence patterns throughout all proteomes. Across the four data sets, only 61 proteins overlapped (Figure 3A). Jaccard analysis revealed that the most similar data sets were the α-synuclein heads and bodies, with a coefficient of 0.199, indicating an overlap of ∼20% in their annotated proteomes (Figure 3B). PQ heads and bodies were the second-most similar, with a coefficient of 0.171 (∼17% overlap). Interestingly, α-synuclein heads and PQ heads showed a coefficient of 0.148 (∼15% overlap), suggesting that the pathogenic cellular mechanisms of these models differ but share features. This seemingly low similarity may be because the α-synuclein data sets had a greater number of annotated proteins, with the α-synuclein having ∼60% more hits than the PQ heads, and the bodies had ∼75% more hits than the PQ bodies.

**Figure 3.**
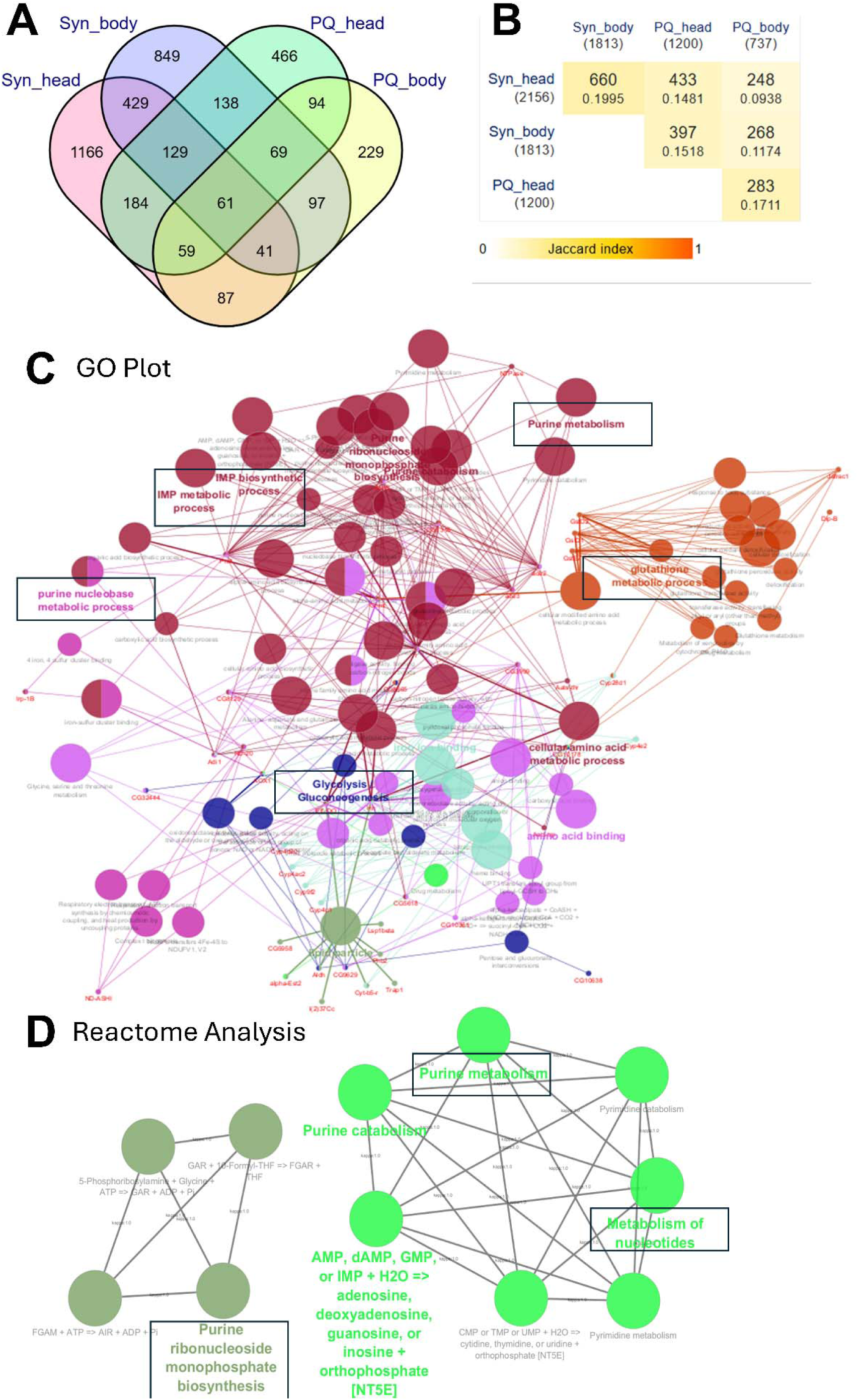
Comparative proteomic gene ontology and Reactome analyses of *Drosophila* α-synuclein and PQ ingestion models of Parkinsonism validate purine metabolism dysregulation as a key overlapping pathway. (A) Venn diagram showing the number of overlapping proteins across all four data sets: heads and bodies of *Drosophila* expressing α-synuclein neuronally and PQ-treated *Drosophila*. (B) Jaccard index analysis of all datasets based on the overlap of identified proteins. The table shows the number of proteins identified for each dataset under each group label. The yellow boxes for each comparison show the number of proteins shared across data set pairs (on top), with the Jaccard index value (below) expressing the fraction of shared proteins as a value between 0 and 1. (C) Comparative gene ontology analysis plot identifying common cellular functions of dysregulated proteins across all four data sets. (D) Comparative Reactome Analysis outlining shared biological pathways across all four data sets.

Comparative GO analysis of these four datasets, examining significantly altered proteins, revealed shared molecular functions, biological processes, and cellular targets. These pathways include “purine metabolism” and “purine nucleobase metabolic process”, “glutathione metabolism”, “inosine monophosphate biosynthesis” and “metabolism”, as well as “glycolysis” and “glucogenesis” (Figure 3C). Consistently, Reactome analysis showed the significant enrichment of “purine metabolism”, “purine ribonucleoside monophosphate biosynthesis”, and “metabolism of nucleotides” across all four data sets (Figure 3D).

To begin investigating metabolic dysfunction in our *Drosophila* models, we employed a metabolic network-based approach to characterize metabolic differences across these models in response to PQ and α-synuclein. We first examined the impact of the differentially abundant proteins (*p*-value <0.05) by detecting statistically significantly enriched FlyBase phenotypes (FDR < 0.05) utilizing the PANGEA tool ^48^. PANGEA identified 382 genes involved in the differential reactions in the PQhead group, demonstrating an enriched fly phenotype of abnormal oxidative stress response (10 overlapped genes and p-value ≌ 0.006).

### Metabolic network analyses further confirm the alteration of nucleotide metabolism in both α-synuclein and PQ models of Parkinsonism

We utilized the iDrosophila1 metabolic network model to further explore PQ- or α-synuclein-induced metabolic disruptions in the brains and bodies of our *Drosophila* models. Proteomics data were first re-annotated to be compatible with iDrosophila1. We assigned FlyBase gene IDs to a substantial proportion of proteins corresponding to the 2,388 genes in the metabolic model (PQ_head_: 65.9%, PQ_body_: 73.2%, αSyn_head_: 71.5%, and αSyn_body_: 76.7%) and ensured its sufficient coverage for downstream network-based modeling. Using the re-annotated proteomics data, two complementary approaches, including ΔFBA and iMAT-based reaction activity analysis, were applied to enhance the predictive power of the metabolic models. This network-based analysis framework is summarized in Figure 4. Through this approach, we identified differential reactions, along with their associated metabolites and genes. The numbers of these differential elements are presented in Table 1.

**Figure 4.**
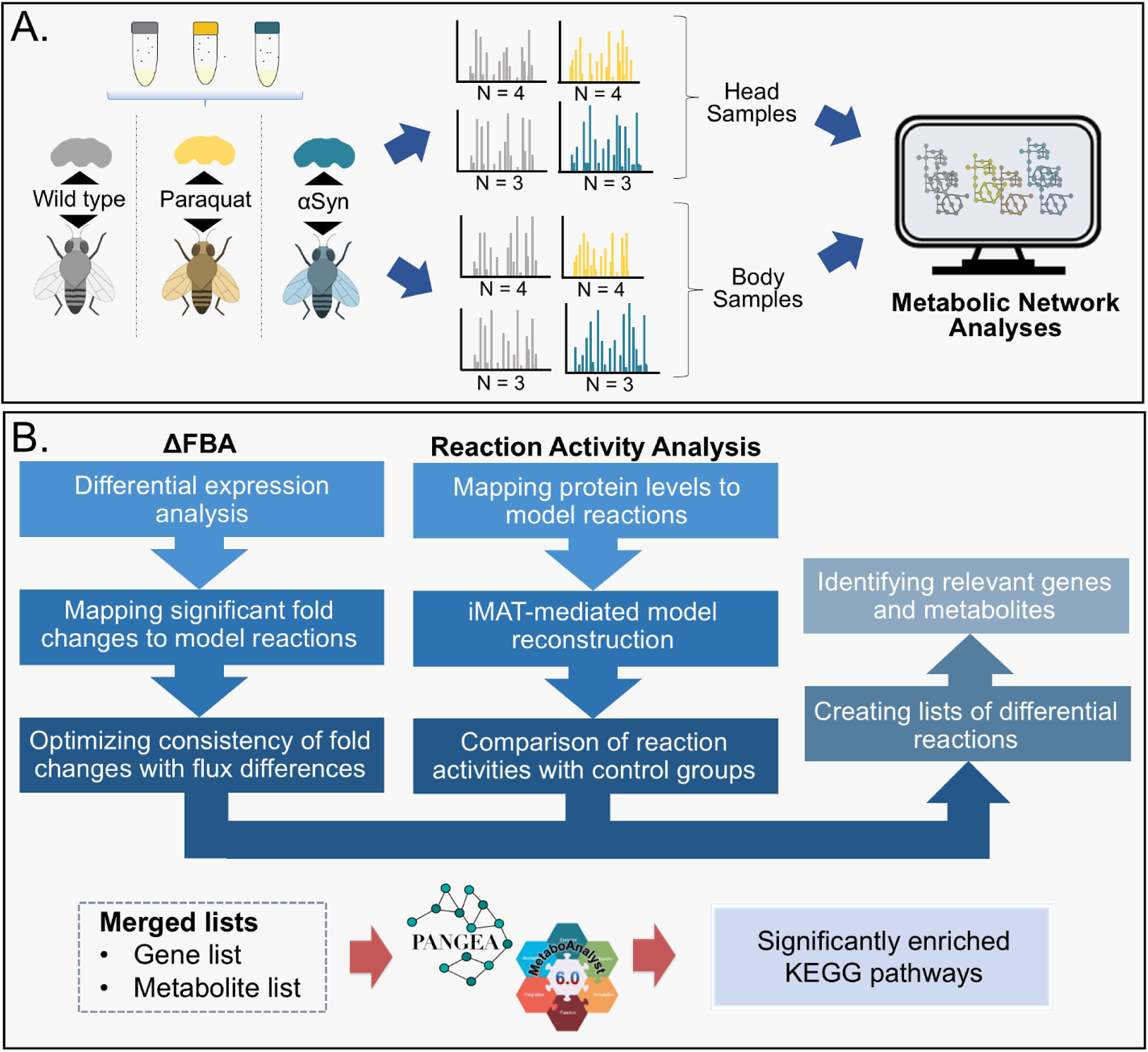
Graphical framework of the iDrosophila1 model-mediated metabolic network analyses. (A) Generation of proteomic data using *Drosophila* models of Parkinsonism (PQ_head_, PQ_body_, αSyn_head_ and αSyn_body_) served as a starting point for metabolic network analyses. (B) iMAT-based reaction activity analysis identifies changes in the activity status of reactions through condition-specific models. The ΔFBA approach aims to optimize the consistency between proteomic changes and predicted metabolic fluxes. Integrating differential reactions identified by both methods for each PD group, along with the analysis of associated genes and metabolites, enables the identification of significantly affected KEGG pathways in response to PQ treatment or α-synuclein expression.

**Table 1.** Overview of differential reactions, their corresponding unique compartment-free metabolites, and genes predicted through metabolic network-based analyses across Parkinson’s disease groups.

| PD Group | Method | No. of: | Metabolites | Genes |
| --- | --- | --- | --- | --- |
|  |  | Reactions |  |  |
| PQ<sub>head</sub> | ΔFBA | 366 | 478 | 351 |
|  | Reaction | 58 | 78 | 44 |
| PQ<sub>body</sub> | ΔFBA | 57 | 76 | 154 |
|  | Reaction | 30 | 56 | 27 |
| αSyn<sub>head</sub> | ΔFBA | 432 | 485 | 307 |
|  | Reaction | 98 | 238 | 81 |
| αSyn<sub>body</sub> | ΔFBA | 259 | 211 | 209 |
|  | Reaction | 82 | 172 | 60 |

Notably, among all PD groups, the PQ_body_ group exhibited the fewest differential reactions across both network-based approaches, accompanied by a correspondingly lower number of total metabolites and associated genes. In agreement with this trend, we observed fewer differential reactions for αSyn_body_ than for αSyn_head_, suggesting that head tissues may undergo more extensive or detectable metabolic changes in response to PD-related perturbations (Table 1). This variation may be attributed to the higher metabolic activity and selective vulnerability of specific brain cell subpopulations in PD, such as dopaminergic neurons in the midbrain ^69,70^.

To investigate tissue- and condition-specific metabolic signatures, we analyzed the gene and metabolite lists derived from the combined set of differential reactions identified by both network-based approaches. To this end, pathway enrichment analysis was performed for each PD group using the gene (Table S1) and metabolite (Table S2) lists. Subsequently, significantly enriched KEGG pathways were quantified. As previously noted, fewer metabolic changes were observed in body samples when compared to head samples (Figure 5A-B, Table S3-4). Specifically, α-synuclein expression and PQ treatment resulted in the enrichment of 107 and 95 pathways for the head samples, respectively, based on the gene lists (FDR < 0.05; Figure 5A). Furthermore, 21 and 27 pathways were over-represented for these groups based on the metabolite lists (*p*-value < 0.05, pathway impact > 0.10; Figure 5B). The total number of significantly enriched KEGG pathways is represented in the upper triangle of each matrix in Figure 5A-B. The lower triangles indicate the number of pathways shared between each pair of PD groups, highlighting the degree of metabolic overlap. Interestingly, a greater number of overlapping metabolic pathways was identified between the PQ_head_ and αSyn groups (αSyn_head_ and αSyn_body_) for both gene and metabolite lists. This pattern could be influenced by the metabolic gene coverage of the underlying proteomics data, and it may shift with more comprehensive datasets.

**Figure 5.**
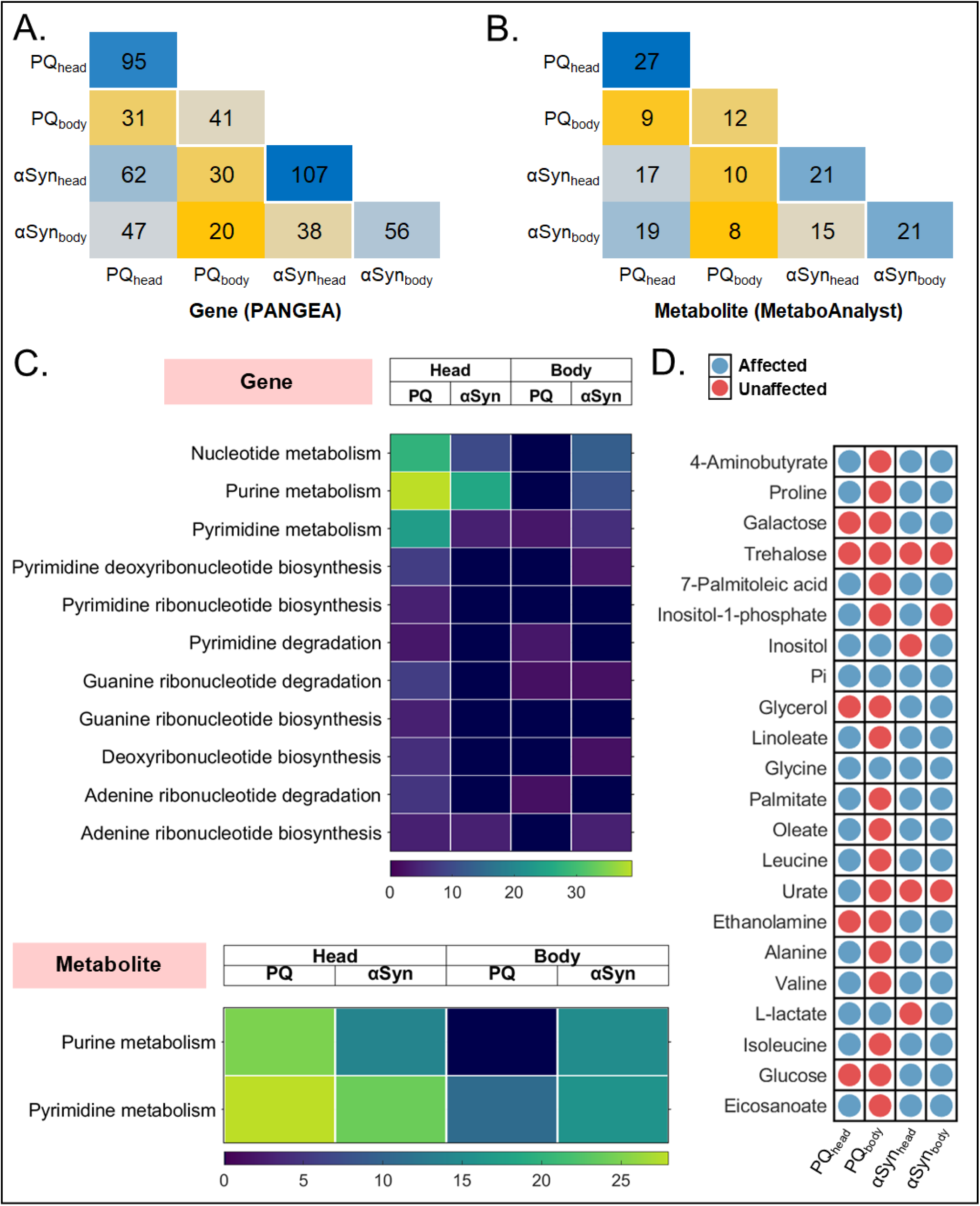
Metabolic network-based analyses of Parkinsonism highlights that differential nucleotide metabolism is one of the commonly affected patterns across diverse PD models. (A) Pathway enrichment analyses were performed for each PD group using gene (A) and metabolite (B) lists derived from differential reactions identified by iMAT-based reaction activity analysis and ΔFBA. The upper triangles in each matrix show the total number of significantly enriched KEGG pathways per group, while the lower triangles indicate the number of pathways shared between PD groups, highlighting metabolic overlap. In both matrices, darker blue indicates a higher number of significantly enriched pathways, while darker orange represents a lower number. C) Heatmaps of the enriched KEGG pathways related to nucleotide metabolism based on the gene (top) and metabolite (bottom) lists. The color gradient reflects the degree of pathway enrichment. It ranges from dark blue to green, with darker blue indicating fewer genes or metabolites involved in that pathway. D) Binary map showing the presence (blue circles) or absence (red circles) of network-based predicted metabolites across different PD groups in the metabolome-based validation list.

Using the gene lists, 11 significantly enriched KEGG pathways were shared by all PQ and αSyn groups, primarily associated with core disruptions in “nucleotide synthesis”, “folate metabolism”, “lipid metabolism”, “amino acid catabolism”, and “cellular transport and signaling”. Several other pathways, particularly those related to energy metabolism (e.g., the TCA cycle, glycolysis, and gluconeogenesis), were specifically over-represented for the head groups (Table S3). Consistently, the metabolite-based enrichment analyses revealed the over-representation of KEGG pathways in similar functional categories, including nucleotide, cofactor, lipid, and amino acid metabolism (Table S4). Of these pathways, “glutathione metabolism”, “folate metabolism”, and “nucleotide metabolism”, emerged as shared metabolic signatures in both PQ-treated and α-synuclein-expressing samples, regardless of tissue type. Specifically, we illustrate the significantly enriched pathways related to nucleotide metabolism for all PD groups. The head samples displayed stronger enrichment than the body samples, particularly for purine metabolism in both the gene and metabolite lists (Figure 5C).

The glutathione (GSH) system is one of the most crucial antioxidant defenses that protects cells from oxidative stress by neutralizing reactive oxygen species ^71^. Therefore, GSH depletion triggers increased oxidative stress, which is suggested to induce neurodegeneration in PD. On the other hand, it remains unclear whether GSH redox imbalance is a consequence of PD-related metabolic changes or a primary cause of its pathophysiology ^72^. ΔFBA analysis revealed a regulation in the flux of the reaction “HMR_4326” in the head following PQ exposure. This reaction, which plays a critical role in the GSH biosynthesis, is catalyzed by the isoenzymes including glutathione synthetase 1 (encoded by *Gss1*; FBgn0030882) and glutathione synthetase 2 (encoded by *Gss2*; FBgn0052495) in the iDrosophila1 model. Glutathione synthetase 1 is absent in the proteomics data, but glutathione synthetase 2 was detected to be significantly decreased in the PQ_head_ group at the proteomic level (FDR ≈ 0.02), thereby suggesting that the suppressed activity of this reaction may contribute to impaired redox homeostasis and increased oxidative vulnerability in the heads of PQ-treated flies.

Folate supplementation was previously shown to transiently reduce mitochondrial hydrogen peroxide levels and partially restore GSH redox equilibrium in *parkin*-null flies; however, this effect was insufficient to alleviate oxidative stress throughout the brain ^73^. This metabolite is also involved in nucleotide synthesis ^74,75^. To date, many researchers have focused on purine metabolism as a target for drug development and nutritional supplementation interventions in PD ^76,77^. This is due to the epidemiological link between low serum urate levels and increased risk for PD, as well as its potential utility in the early detection and prognosis of the disorder ^76^. Notably, numerous studies have reported lower serum urate levels in PD patients compared to healthy controls ^78–81^. In *Drosophila*, the biosynthesis of this antioxidant metabolite is catalyzed by xanthine dehydrogenase (XDH) / oxidase (XOD), encoded by the *rosy* gene (FBgn0003308). Even though urate functions as an antioxidant, the oxidase form (XOD) can also contribute to reactive oxygen species production by generating both urate and hydrogen peroxide under aging conditions ^82^. In the current work, the ΔFBA analysis predicted a changed flux for the XDH/XOD reaction (HMR_4649) in the fly head under PQ exposure. The reduction in the urate production rate was also confirmed by a significant decrease in the protein-level expression of *rosy* (FDR ≈ 0.001). It supports that *rosy* locus mutants lacking urate production were previously shown to be hypersensitive to PQ, highlighting the protective role of urate against oxidative stress ^83^.

Altered urate levels in the PQ-treated *Drosophila* heads were further supported by the findings of Shukla et al., who employed a gas chromatography–mass spectrometry (GC-MS)-based metabolomics approach combined with partial least-square discriminant analysis (PLS-DA) ^84^ (Figure 5D). In their study, flies were exposed to varying concentrations of PQ (5, 10, and 20 mM), and significant shifts in brain metabolite composition were observed. 22 out of 24 differential metabolites from their study are present in the reactions of the iDrosophila1 model. In the current work, over 70% of these differential metabolites were identified to be involved in the differential reactions for PQ_head_ and αSyn groups, while only 4 of them were identified for the PQ_body_ group.

### Behavior assays in a reverse genetic screen of purinergic ectonucleotidases in *Drosophila* treated with PQ reveal disruption in glial purine metabolism

After establishing that disruptions in purine metabolism were present in both genetic and environmental models of PD in flies, we performed a reverse genetic screen of *Drosophila* human orthologs of purinergic ectonucleotidases involved in purine metabolism (Figure 6A). We utilized publicly available UAS RNAi *Drosophila* strains for Veil, Uricase, and CG-16758. The screen used pan-neuronal and pan-glial expression systems (nSyb-GAL4 and SybRepo-GAL4, respectively) to determine whether these cell types are preferentially vulnerable to disruptions in purine metabolism. Locomotor activity assays in these flies showed that PQ-treated glial knockdown of Veil significantly improved the behavioral deficits observed in the PQ-treated genetic vehicle control (Figure 6B). Meanwhile, no statistically significant changes in behavior were observed with PQ treatment in either the neuronal or glial knockdowns of Uricase or CG-16758 (Figure 6B-C).

**Figure 6.**
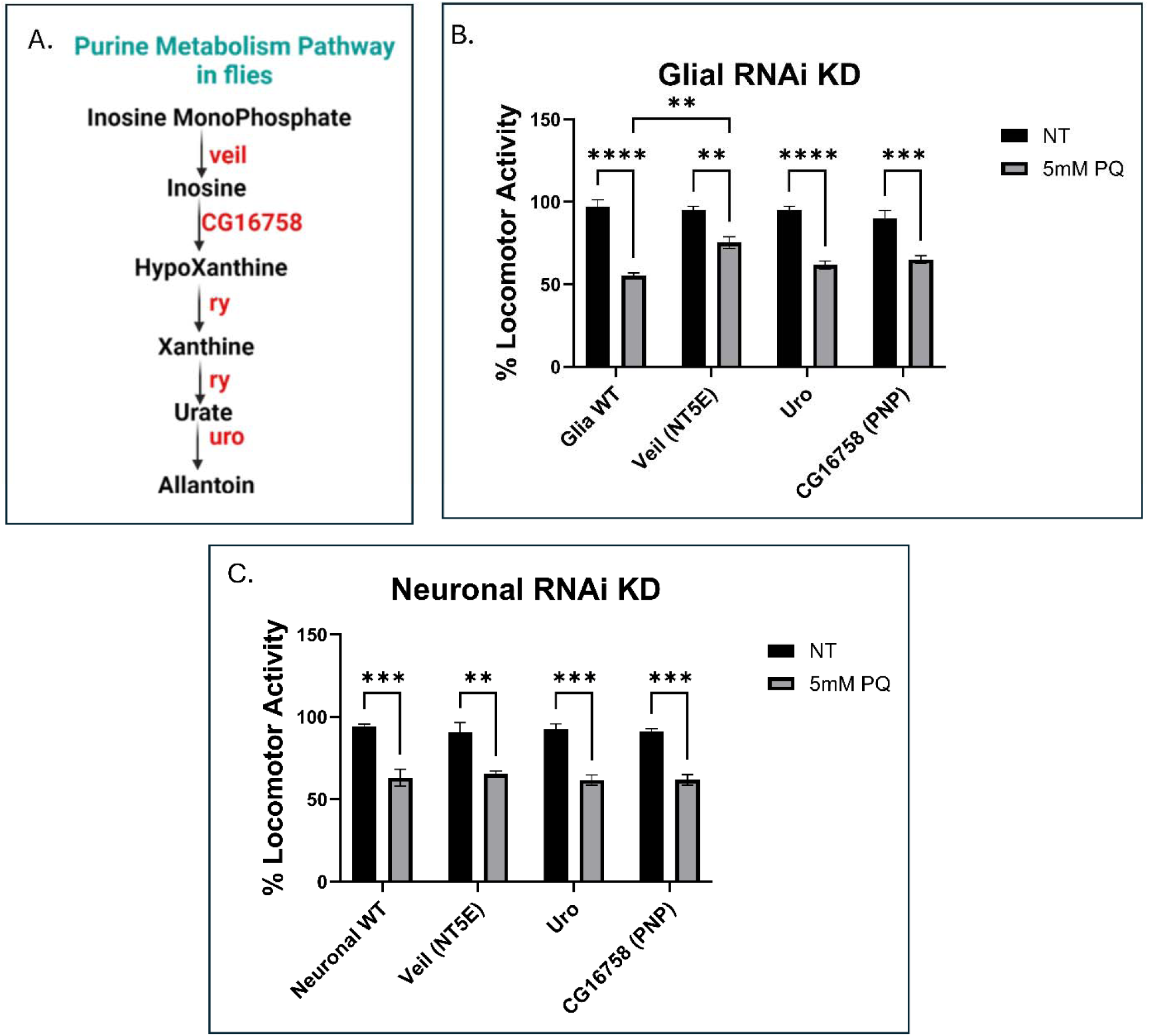
Reverse genetic screen of purine metabolism enzymes in *Drosophila* treated with PQ identifies neuronal NT5E as a potential therapeutic target. (A) Simplified degradation pathway of purine nucleotides in *Drosophila* melanogaster. (B-C) Locomotor assays of transgenic *Drosophila* expressing RNAi constructs in either neurons or pan-glial cells. The purine metabolism enzymes knocked down are the human orthologs Veil, Uricase, and CG-16758. On the x-axis, the gene names for flies are listed first, and the human ortholog is in parentheses (if the name differs between species).

### Metabolomic analysis of the brainstem of mice exposed to aerosolized PQ validates that disruptions in nucleotide metabolism are conserved across vertebrates

Following our comparative proteomic analysis in *Drosophila*, we sought to confirm that the changes in nucleotide metabolism observed in our PQ neurotoxicity ingestion model would translate to a mammalian model. Our lab has previously developed a low-dose (0.2 mg/mL) inhalation model of PQ toxicity in mice to capture a more environmentally relevant route of exposure in humans ^7^. To determine whether similar metabolic pathways are disrupted in a similar manner in mammalian models of PQ toxicity, we performed metabolomic analysis of brainstem microdissections from male and female mice exposed to aerosolized PQ for 3 months (Figure 7A). Both sexes demonstrated statistically significant changes in pyrimidine and purine metabolism, as well as the urea cycle (Figure 7B-C). Other pathways enriched in both sexes included mitochondrial metabolism pathways. These included the oxidation of short-, branched-, and long-chain fatty acids, implicating mitochondrial fatty acid metabolism in PQ-induced neurotoxicity. This enrichment may stem from PQ’s inhibition of Complex I of the ETC, stalling fatty acid β-oxidation in the inner membrane by limiting the availability of NAD+ and FAD+, and subsequently reducing the pool of acetyl-CoA ^85^. Notably, more changes were found in female mice than in males, despite largely sharing similar enriched pathways, indicating a sex-specific effect that may relate to adaptive mechanisms against oxidative stress ^86^. PD has a male bias in diagnoses of about 2:1 in humans ^1^, meaning that these proteomic differences between sexes may underpin this difference in vulnerability.

**Figure 7.**
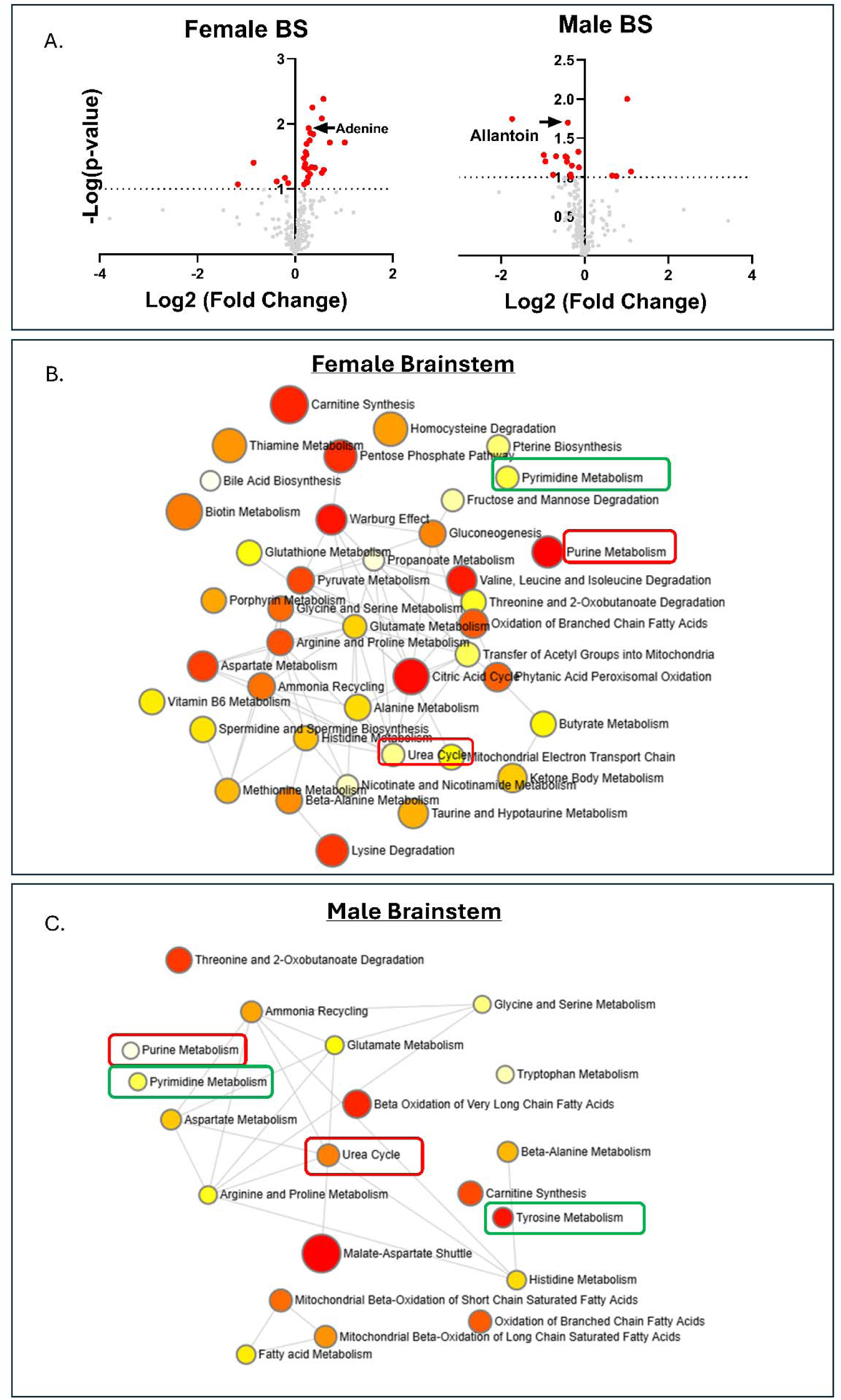
Metabolomic analysis of brainstems from mice exposed to aerosolized PQ validates that disruptions in purine metabolism translate to vertebrates. (A) Volcano plots of metabolite changes in the brainstems of male and female C57 mice exposed to aerosolized PQ. (B-C) Network analyses of statistically significant changes in the metabolomes of female and male brainstems, respectively, highlight purine metabolism changes in red boxes and pathways related to other PD features in green boxes.

## DISCUSSION

Here, we present a comparative proteomic analysis of an environmental neurotoxicant model (paraquat) and a genetic model of PD (neuronal α-synuclein) in *Drosophila*. This was paired with additional metabolic network-based analyses in *Drosophila* and metabolomic validation from a murine model of PQ inhalation. This resulting comparison demonstrated that despite diverse proteomic changes, our PQ ingestion and neuronal α-synuclein models both share key dysregulated pathways. This suggests that both models may act on shared cellular targets through similar mechanisms of action.

Our comparative proteomics analysis of PQ-exposure and neuronal α-synuclein *Drosophila* models confirmed that proteomic changes were distinct and segregated independently in each PD model, exhibiting significant differential expression profiles compared to their respective controls (Figure 1A-B; Supplemental Figure 1A-B). GO enrichment analyses revealed many expected cellular targets for the PQ model, with the heads showing that the dysregulated proteins were predominantly associated with “cellular metabolism”, “respiratory electron transport”, and “biogenesis of complex I of the ETC” (Figure 1D). The bodies exhibited similar metabolic changes, but with greater emphasis on protease-related pathways rather than mitochondrial terms (Figure 2D). Both the heads and bodies had nucleotide biosynthesis and related pathways as dysregulated in their individual analyses. The proteomic data from the neuronal α-synuclein expression flies were consistent with findings reported in previous proteomics studies on this model ^87^. GO analysis revealed that the heads and bodies had broad enrichment across metabolic pathways, with particular emphasis on nucleotide metabolism.

Additionally, we performed metabolic network analyses using the iDrosophila1 model to comprehensively characterize the PQ- and α-synuclein-induced metabolic alterations in the heads and bodies of *Drosophila*. In this regard, the complementary approaches (iMAT-based reaction activity analysis and ΔFBA) were performed to identify differential metabolic reactions. Since each method captures specific aspects of metabolic changes and has its own limitations (e.g., ignoring flux-level changes by the first approach and the dependence of the second method on flux difference constraints), we combined differential reaction lists from each approach for a more nuanced understanding of the metabolic disruptions associated with each PD model. Subsequently, we analyzed the associated genes and metabolites using the Kyoto Encyclopedia of Genes and Genomes (KEGG) database to identify enriched pathways. GSH metabolism, folate metabolism, and nucleotide metabolism were among the most prominent metabolic alterations shared between the *Drosophila* PD models.

Kang et al. reported that PQ administration reduced GSH levels in the substantia nigra of mice, leading to increased oxidative stress and dopaminergic neurotoxicity ^88^. Similar effects on GSH metabolism have also been observed in other systems (e.g., cultured pulmonary endothelial cells, liver, kidney, and lung tissues), where PQ exposure triggered time- and dose-dependent oxidative stress ^89–91^. Overall, modulation of GSH metabolism in both head and body samples may reflect PD-related regulation of the systemic oxidative stress response.

In addition, upregulated nucleotide metabolism has been previously shown to counteract mitochondrial dysfunction in pink1 mutant flies, further supporting a potential mechanistic link among folate-dependent pathways, nucleotide metabolism, and mitochondrial health in PD models ^92,93^. Consistently, the final product of purine degradation, urate, significantly contributes to antioxidant defense by scavenging reactive oxygen and nitrogen species and stabilizing key antioxidant systems ^76^. Thus, we suggest that the dual impairment of urate production and GSH biosynthesis may herein reflect a disrupted redox balance in the PQ-exposed head samples. Furthermore, our results emphasize the potential involvement of nucleotide and urate metabolism, as well as oxidative stress, in the PD-related pathology in both of our *Drosophila* models.

Next, we sought to understand how the purine-related enzymes may be involved in the PQ-induced metabolic dysfunctions observed in our proteomics-based analyses. To this end, we employed the facile genetic manipulation of the *Drosophila* model. Using publicly available RNAi genetic lines, we performed a reverse genetic screen to knock down catabolic purine enzymes separately in neurons or glia (SybSyb-GAL4; SybRepo-GAL4) (Figure 6A). The flies were fed either 5 mM PQ food (or control food) for 7 days post-eclosion. Locomotor activity assays were performed on the knockdown flies on day 7 of treatment. This revealed that the only significant difference in locomotor behavior was found in the glial knockdown of the Veil enzyme (NT5E in humans), that catalyzes the conversion of inosine monophosphate (IMP) to inosine (Figure 6B). These findings suggest that glial conversion of IMP to inosine may prevent scavenging of IMP for conversion to guanosine monophosphate (GMP) or adenine monophosphate (AMP). These molecules are critical for energy metabolism and cell signaling, meaning that preventing the conversion of IMP to inosine increases the labile pool of substrate that can be interchangeably converted to either AMP or GMP. This shift in pathways dynamics may confer a protective benefit to the flies, allowing them to adapt to greater metabolic stress induced by PQ exposure.

Finally, to gain further insight into the metabolic modeling–predicted alterations in urate metabolism and redox balance in the PQ_head_ group, we expanded our analyses with complementary omics approaches. Since constraint-based modeling relies on steady-state assumptions, and *Drosophila* may exhibit metabolic features distinct from those of mammals, we complemented these analyses with metabolomic profiling of brainstem samples from a low-dose PQ inhalation model. Our group has helped develop this model using male and female C57BL/6J mice exposed to aerosolized PQ (0.2 mg/mL) for 4 hours a day, 5 days a week for 3 months (60 days of exposure total) ^7^.

KEGG pathway analysis of this metabolomics data confirmed the enrichment in purine and pyrimidine metabolism, as well as in the urea cycle (Figure 7B-C). However, metabolomics analyses revealed a greater number of differential metabolites in PQ-exposed females than in males, suggesting protective compensatory mechanisms that ameliorate PQ neurotoxicity. Aside from purine and pyrimidine metabolism, females showed broad changes across various metabolic pathways relating to “fatty acid metabolism”, “glutathione metabolism”, and “the urea cycle”. On the other hand, males exhibited significant changes in these same pathways, as well as in “tyrosine metabolism”. Together, these data validate our findings in *Drosophila*, demonstrating dysregulation of purine metabolism following PQ treatment.

In the context of clinical trials involving inosine, these data suggest that targeting inosine may be insufficient to treat PD at early stages. Instead, protein-level modifications of enzymes upstream of inosine metabolism may be more responsible for the protective adaptations conferred onto the glial knockouts of Veil in *Drosophila*. These findings may also suggest that urate’s antioxidant activity is only one of several protective mechanisms that mitigate PQ-neurotoxicity. Instead, other functions of purine nucleotide metabolism, particularly upstream of inosine catabolism, may play a more important role than previously anticipated by the field.

## Acknowledgements

The α-synuclein fly model was a gift from Dr. Mel Feany at Harvard Medical School.

This research was supported by R00ES033723, R21ES037434-01 to SS and T32ES007026, P30ES001247, and T32AG076455-03.

## Data Availability

1. The mass spectrometry proteomics data have been deposited to the ProteomeXchange Consortium via the PRIDE partner repository with the dataset identifier PXD065124.

a. Reviewer access details: Log in to the PRIDE website using the following details: Project accession: PXD065124, Token: jW41e3knNmRd
b. Alternatively, reviewers can access the dataset by logging in to the PRIDE website using the following account details. Username, Password: Bx95XlQPzT49
2. Mass Spectrometry metabolomics data have been deposited to the Metabolomics Workbench Repository ^94^. DOI: PR002672, http://dx.doi.org/10.21228/M88G3B.

a. This data has been embargoed until 12/30/2026. This date is subject to change pending acceptance of the manuscript for publication.
b. This study is available at the NIH Common Fund’s National Metabolomics Data Repository (NMDR) website, the Metabolomics Workbench, https://www.metabolomicsworkbench.org where it has been assigned Study ID ST004235. This work is supported by NIH grants U2C-DK119886 and OT2-OD030544.

---

**Supplemental Figure 1.**
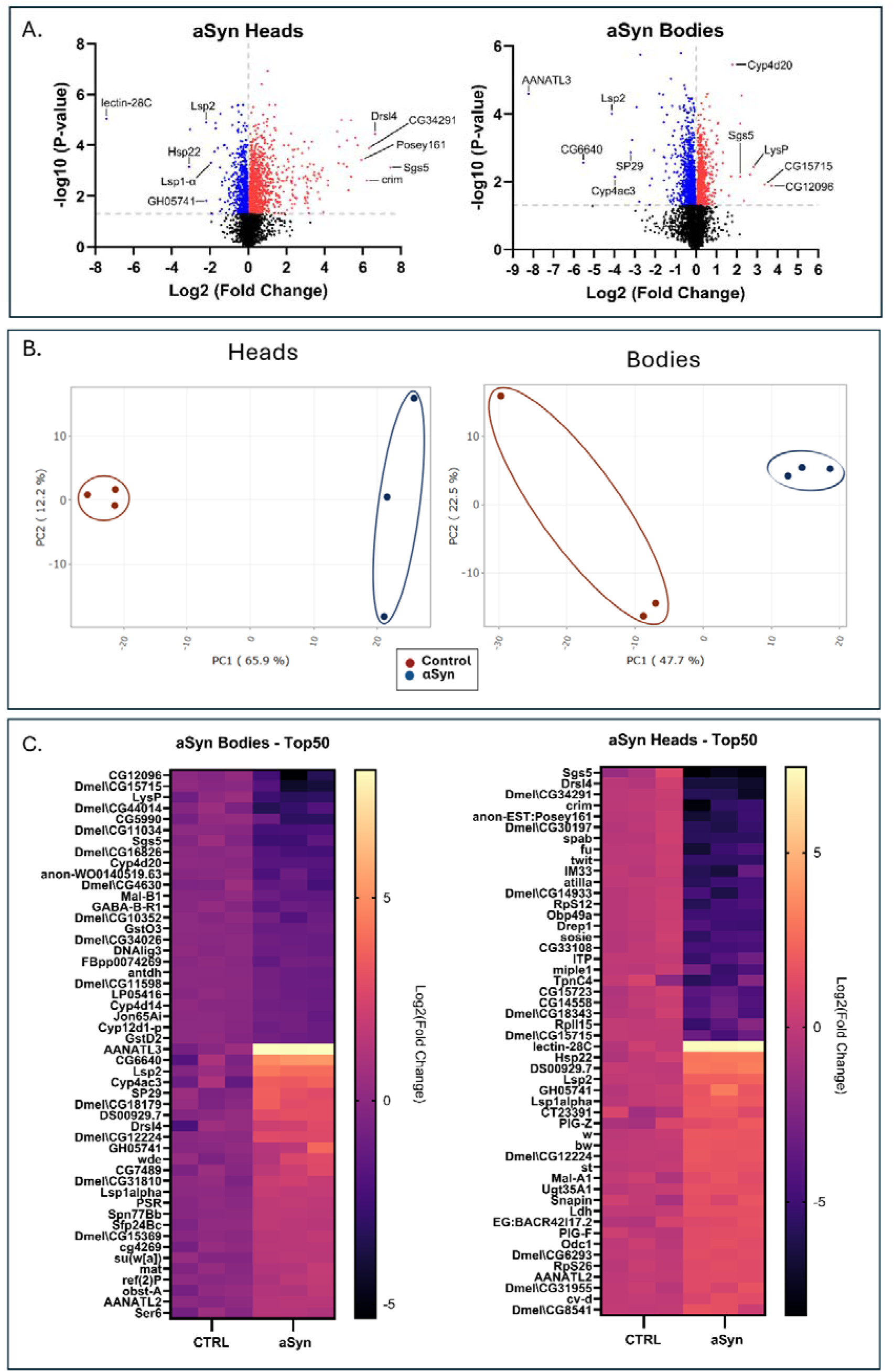
Proteomics analysis of neuronally expressed α-synuclein model of Parkinsonism in *Drosophila*. (A) Volcano plots showing the most significant up- and down-regulated proteins in the proteomics data set by -Log10 p-value. (B) PCA plot of proteomics data from the heads and bodies of neuronal α-synuclein expressing *Drosophila*, demonstrating separate clustering of groups and neat segregation of treatment groups. (C) Heatmap of the 50 genes corresponding to the most statistically significant protein expression changes between the heads and bodies of neuronal α-synuclein expressing *Drosophila and controls*, colored according to their Log_2_ fold change. The first 25 genes are the most down-regulated genes, while the next 25 are the most up-regulated.

## Notes

The authors report no conflict(s) of interest regarding this study.

### Competing Interest Statement

The authors have declared no competing interest.

