## Supplemental Table 1 for "Comparative Proteomic Analysis of Environmental and Genetic Models of Parkinson’s Disease Highlights the Role of Purine Metabolism"

**Table S1 |** List of the genes in the ΔFBA- and reaction activity analysis-predicted differential reactions for paraquat treatment or α-synuclein expression to induce Parkinson's disease-related changes in the *Drosophila* head or body.

| Gene | Gene Symbol | PD Group |
|---|---|---|
| FBgn0000032 | *Acph-1* | $PQ_{head}$ |
| FBgn0000150 | *awd* | $PQ_{head}$ |
| FBgn0000261 | *Cat* | $PQ_{head}$ |
| FBgn0000303 | *ChAT* | $PQ_{head}$ |
| FBgn0000479 | *dnc* | $PQ_{head}$ |
| FBgn0001098 | *Gdh* | $PQ_{head}$ |
| FBgn0001124 | *Got1* | $PQ_{head}$ |
| FBgn0001125 | *Got2* | $PQ_{head}$ |
| FBgn0001128 | *Gpdh1* | $PQ_{head}$ |
| FBgn0001258 | *Ldh* | $PQ_{head}$ |
| FBgn0002069 | *AspRS* | $PQ_{head}$ |
| FBgn0002524 | *lace* | $PQ_{head}$ |
| FBgn0002719 | *Men* | $PQ_{head}$ |
| FBgn0002778 | *mnd* | $PQ_{head}$ |
| FBgn0002937 | *ninaB* | $PQ_{head}$ |
| FBgn0003141 | *pr* | $PQ_{head}$ |
| FBgn0003301 | *rut* | $PQ_{head}$ |
| FBgn0003308 | *ry* | $PQ_{head}$ |
| FBgn0003416 | *sl* | $PQ_{head}$ |
| FBgn0004516 | *Gad1* | $PQ_{head}$ |
| FBgn0004611 | *Plc21C* | $PQ_{head}$ |
| FBgn0004797 | *mdy* | $PQ_{head}$ |
| FBgn0004852 | *Ac76E* | $PQ_{head}$ |
| FBgn0005674 | *GluProRS* | $PQ_{head}$ |
| FBgn0008635 | *βCOP* | $PQ_{head}$ |
| FBgn0010197 | *Gyc32E* | $PQ_{head}$ |
| FBgn0010383 | *Cyp18a1* | $PQ_{head}$ |
| FBgn0010548 | *Aldh-III* | $PQ_{head}$ |
| FBgn0010611 | *Hmgs* | $PQ_{head}$ |
| FBgn0010803 | *TrpRS* | $PQ_{head}$ |
| FBgn0011768 | *Fdh* | $PQ_{head}$ |
| FBgn0012034 | *AcCoAS* | $PQ_{head}$ |
| FBgn0013972 | *Gycα99B* | $PQ_{head}$ |
| FBgn0013973 | *Gycβ100B* | $PQ_{head}$ |
| FBgn0014032 | *Sptr* | $PQ_{head}$ |
| FBgn0014455 | *Ahcy* | $PQ_{head}$ |

| | | |
|---|---|---|
| FBgn0016078 | *wun* | $PQ_{head}$ |
| FBgn0016687 | *Nurf-38* | $PQ_{head}$ |
| FBgn0016694 | *Pdp1* | $PQ_{head}$ |
| FBgn0017482 | *T3dh* | $PQ_{head}$ |
| FBgn0019662 | *qm* | $PQ_{head}$ |
| FBgn0019830 | *colt* | $PQ_{head}$ |
| FBgn0020236 | *ATPCL* | $PQ_{head}$ |
| FBgn0020385 | *pug* | $PQ_{head}$ |
| FBgn0020389 | *Papss* | $PQ_{head}$ |
| FBgn0020653 | *Trxr-1* | $PQ_{head}$ |
| FBgn0021750 | *SerRS-m* | $PQ_{head}$ |
| FBgn0021765 | *scu* | $PQ_{head}$ |
| FBgn0022160 | *Gpo1* | $PQ_{head}$ |
| FBgn0022710 | *Ac13E* | $PQ_{head}$ |
| FBgn0023023 | *CRMP* | $PQ_{head}$ |
| FBgn0023416 | *Ac3* | $PQ_{head}$ |
| FBgn0024150 | *Ac78C* | $PQ_{head}$ |
| FBgn0024947 | *NTPase* | $PQ_{head}$ |
| FBgn0025352 | *Mtpβ* | $PQ_{head}$ |
| FBgn0025373 | *Fpps* | $PQ_{head}$ |
| FBgn0025709 | *CNT2* | $PQ_{head}$ |
| FBgn0025724 | *βCOP* | $PQ_{head}$ |
| FBgn0025725 | *αCOP* | $PQ_{head}$ |
| FBgn0025807 | *Rad9* | $PQ_{head}$ |
| FBgn0025837 | *CG17636* | $PQ_{head}$ |
| FBgn0025885 | *Inos* | $PQ_{head}$ |
| FBgn0026576 | *Pisd* | $PQ_{head}$ |
| FBgn0026778 | *Rad1* | $PQ_{head}$ |
| FBgn0027079 | *ValRS* | $PQ_{head}$ |
| FBgn0027080 | *TyrRS* | $PQ_{head}$ |
| FBgn0027081 | *ThrRS* | $PQ_{head}$ |
| FBgn0027083 | *MetRS-m* | $PQ_{head}$ |
| FBgn0027084 | *LysRS* | $PQ_{head}$ |
| FBgn0027085 | *LeuRS-m* | $PQ_{head}$ |
| FBgn0027086 | *IleRS* | $PQ_{head}$ |
| FBgn0027087 | *HisRS* | $PQ_{head}$ |
| FBgn0027088 | *GlyRS* | $PQ_{head}$ |
| FBgn0027090 | *GlnRS* | $PQ_{head}$ |
| FBgn0027091 | *CysRS* | $PQ_{head}$ |
| FBgn0027093 | *ArgRS* | $PQ_{head}$ |

| | | |
|---|---|---|
| FBgn0027094 | *AlaRS* | $PQ_{head}$ |
| FBgn0027291 | *Idh3a* | $PQ_{head}$ |
| FBgn0027348 | *bgm* | $PQ_{head}$ |
| FBgn0027496 | *εCOP* | $PQ_{head}$ |
| FBgn0028425 | *JhI-21* | $PQ_{head}$ |
| FBgn0028479 | *Mtpα* | $PQ_{head}$ |
| FBgn0028539 | *Eato* | $PQ_{head}$ |
| FBgn0028707 | *Mt2* | $PQ_{head}$ |
| FBgn0028962 | *AlaRS-m* | $PQ_{head}$ |
| FBgn0028968 | *γCOP* | $PQ_{head}$ |
| FBgn0028969 | *δCOP* | $PQ_{head}$ |
| FBgn0028997 | *nmdyn-D7* | $PQ_{head}$ |
| FBgn0029155 | *Men-b* | $PQ_{head}$ |
| FBgn0029502 | *Coq7* | $PQ_{head}$ |
| FBgn0029648 | *CG3603* | $PQ_{head}$ |
| FBgn0029689 | *CG6428* | $PQ_{head}$ |
| FBgn0029848 | *Btnd* | $PQ_{head}$ |
| FBgn0029975 | *spidey* | $PQ_{head}$ |
| FBgn0030007 | *α-PheRS* | $PQ_{head}$ |
| FBgn0030218 | *CG1628* | $PQ_{head}$ |
| FBgn0030245 | *CG1637* | $PQ_{head}$ |
| FBgn0030300 | *Sk1* | $PQ_{head}$ |
| FBgn0030361 | *CG1492* | $PQ_{head}$ |
| FBgn0030407 | *Fpgs* | $PQ_{head}$ |
| FBgn0030421 | *Agpat1* | $PQ_{head}$ |
| FBgn0030431 | *CG4407* | $PQ_{head}$ |
| FBgn0030460 | *Coq5* | $PQ_{head}$ |
| FBgn0030465 | *CG15743* | $PQ_{head}$ |
| FBgn0030558 | *CG1461* | $PQ_{head}$ |
| FBgn0030573 | *nmdyn-D6* | $PQ_{head}$ |
| FBgn0030653 | *CG7860* | $PQ_{head}$ |
| FBgn0030683 | *Mvd* | $PQ_{head}$ |
| FBgn0030796 | *CG4829* | $PQ_{head}$ |
| FBgn0030816 | *CG16700* | $PQ_{head}$ |
| FBgn0030817 | *CG4991* | $PQ_{head}$ |
| FBgn0030882 | *Gss1* | $PQ_{head}$ |
| FBgn0030932 | *Ggt-1* | $PQ_{head}$ |
| FBgn0031048 | *CG12237* | $PQ_{head}$ |
| FBgn0031148 | *Cbs* | $PQ_{head}$ |
| FBgn0031170 | *Abca3* | $PQ_{head}$ |

| | | |
|---|---|---|
| FBgn0031250 | *Ent1* | $PQ_{head}$ |
| FBgn0031397 | *CG15385* | $PQ_{head}$ |
| FBgn0031484 | *CG3165* | $PQ_{head}$ |
| FBgn0031497 | *SerRS* | $PQ_{head}$ |
| FBgn0031630 | *CG15629* | $PQ_{head}$ |
| FBgn0031663 | *CG8891* | $PQ_{head}$ |
| FBgn0031713 | *Coq6* | $PQ_{head}$ |
| FBgn0031877 | *Hmgcl* | $PQ_{head}$ |
| FBgn0032059 | *PrBP* | $PQ_{head}$ |
| FBgn0032083 | *CG9541* | $PQ_{head}$ |
| FBgn0032160 | *CG4598* | $PQ_{head}$ |
| FBgn0032219 | *CG4995* | $PQ_{head}$ |
| FBgn0032449 | *CG17036* | $PQ_{head}$ |
| FBgn0032811 | *Pmvk* | $PQ_{head}$ |
| FBgn0032845 | *CG10747* | $PQ_{head}$ |
| FBgn0032910 | *CG9265* | $PQ_{head}$ |
| FBgn0032922 | *Coq3* | $PQ_{head}$ |
| FBgn0032923 | *Pld3* | $PQ_{head}$ |
| FBgn0033214 | *CG1941* | $PQ_{head}$ |
| FBgn0033215 | *Dgat2* | $PQ_{head}$ |
| FBgn0033216 | *CG1946* | $PQ_{head}$ |
| FBgn0033226 | *puml* | $PQ_{head}$ |
| FBgn0033476 | *oys* | $PQ_{head}$ |
| FBgn0033679 | *CG8888* | $PQ_{head}$ |
| FBgn0033761 | *CG8778* | $PQ_{head}$ |
| FBgn0033879 | *Echs1* | $PQ_{head}$ |
| FBgn0033900 | *CysRS-m* | $PQ_{head}$ |
| FBgn0033911 | *VGAT* | $PQ_{head}$ |
| FBgn0034177 | *AsnRS-m* | $PQ_{head}$ |
| FBgn0034225 | *veil* | $PQ_{head}$ |
| FBgn0034299 | *CG5757* | $PQ_{head}$ |
| FBgn0034401 | *MetRS* | $PQ_{head}$ |
| FBgn0034432 | *CG7461* | $PQ_{head}$ |
| FBgn0034493 | *CG8908* | $PQ_{head}$ |
| FBgn0034497 | *Mpcp1* | $PQ_{head}$ |
| FBgn0034716 | *Oatp58Dc* | $PQ_{head}$ |
| FBgn0034800 | *CG3788* | $PQ_{head}$ |
| FBgn0034988 | *cN-IIIB* | $PQ_{head}$ |
| FBgn0035064 | *TyrRS-m* | $PQ_{head}$ |
| FBgn0035169 | *Dci* | $PQ_{head}$ |

| | | |
|---|---|---|
| FBgn0035203 | *Acat2* | $PQ_{head}$ |
| FBgn0035348 | *CG16758* | $PQ_{head}$ |
| FBgn0035371 | *AhcyL1* | $PQ_{head}$ |
| FBgn0035383 | *CPT2* | $PQ_{head}$ |
| FBgn0035471 | *Sc2* | $PQ_{head}$ |
| FBgn0035811 | *Mcad* | $PQ_{head}$ |
| FBgn0035839 | *CG7550* | $PQ_{head}$ |
| FBgn0035942 | *ValRS-m* | $PQ_{head}$ |
| FBgn0035964 | *Dhpr* | $PQ_{head}$ |
| FBgn0036337 | *Adk2* | $PQ_{head}$ |
| FBgn0036569 | *IleRS-m* | $PQ_{head}$ |
| FBgn0036622 | *Agpat4* | $PQ_{head}$ |
| FBgn0036623 | *Agpat3* | $PQ_{head}$ |
| FBgn0036629 | *GluRS-m* | $PQ_{head}$ |
| FBgn0036691 | *beg* | $PQ_{head}$ |
| FBgn0036732 | *Oatp74D* | $PQ_{head}$ |
| FBgn0036747 | *CG6052* | $PQ_{head}$ |
| FBgn0036763 | *TrpRS-m* | $PQ_{head}$ |
| FBgn0036770 | *Prestin* | $PQ_{head}$ |
| FBgn0036821 | *CG3961* | $PQ_{head}$ |
| FBgn0036875 | *CG9449* | $PQ_{head}$ |
| FBgn0036876 | *CG9451* | $PQ_{head}$ |
| FBgn0037044 | *Pdss2* | $PQ_{head}$ |
| FBgn0037063 | *CG9391* | $PQ_{head}$ |
| FBgn0037138 | *P5CDh1* | $PQ_{head}$ |
| FBgn0037163 | *laza* | $PQ_{head}$ |
| FBgn0037165 | *CG11437* | $PQ_{head}$ |
| FBgn0037166 | *CG11426* | $PQ_{head}$ |
| FBgn0037170 | *Trxr-2* | $PQ_{head}$ |
| FBgn0037203 | *slif* | $PQ_{head}$ |
| FBgn0037341 | *CG12746* | $PQ_{head}$ |
| FBgn0037440 | *CRAT* | $PQ_{head}$ |
| FBgn0037513 | *pyd3* | $PQ_{head}$ |
| FBgn0037526 | *ArgRS-m* | $PQ_{head}$ |
| FBgn0037574 | *Coq2* | $PQ_{head}$ |
| FBgn0037759 | *CG8526* | $PQ_{head}$ |
| FBgn0037845 | *CG14694* | $PQ_{head}$ |
| FBgn0037846 | *CG6574* | $PQ_{head}$ |
| FBgn0037852 | *Tpc1* | $PQ_{head}$ |
| FBgn0037900 | *CG5276* | $PQ_{head}$ |

| | | |
|---|---|---|
| FBgn0037913 | *fabp* | $PQ_{head}$ |
| FBgn0038237 | *Pde6* | $PQ_{head}$ |
| FBgn0038302 | *CG4210* | $PQ_{head}$ |
| FBgn0038376 | *Hmt-1* | $PQ_{head}$ |
| FBgn0038516 | *P5cr-2* | $PQ_{head}$ |
| FBgn0038876 | *Idi* | $PQ_{head}$ |
| FBgn0038912 | *CG6656* | $PQ_{head}$ |
| FBgn0038922 | *Idh3b* | $PQ_{head}$ |
| FBgn0038925 | *Cchl* | $PQ_{head}$ |
| FBgn0039175 | *β-PheRS* | $PQ_{head}$ |
| FBgn0039184 | *CG6432* | $PQ_{head}$ |
| FBgn0039244 | *CG11069* | $PQ_{head}$ |
| FBgn0039304 | *CG10425* | $PQ_{head}$ |
| FBgn0039357 | *CG4743* | $PQ_{head}$ |
| FBgn0039358 | *CG5028* | $PQ_{head}$ |
| FBgn0039464 | *CG6330* | $PQ_{head}$ |
| FBgn0039698 | *CG7789* | $PQ_{head}$ |
| FBgn0039774 | *CDase* | $PQ_{head}$ |
| FBgn0039809 | *CG15547* | $PQ_{head}$ |
| FBgn0039844 | *CG1607* | $PQ_{head}$ |
| FBgn0040064 | *yip2* | $PQ_{head}$ |
| FBgn0040077 | *primo-1* | $PQ_{head}$ |
| FBgn0040319 | *Gclc* | $PQ_{head}$ |
| FBgn0041087 | *wun2* | $PQ_{head}$ |
| FBgn0041150 | *hoe1* | $PQ_{head}$ |
| FBgn0042627 | *FASN2* | $PQ_{head}$ |
| FBgn0045064 | *bwa* | $PQ_{head}$ |
| FBgn0046114 | *Gclm* | $PQ_{head}$ |
| FBgn0050035 | *Tret1-1* | $PQ_{head}$ |
| FBgn0050104 | *NT5E-2* | $PQ_{head}$ |
| FBgn0050345 | *CG30345* | $PQ_{head}$ |
| FBgn0051005 | *qless* | $PQ_{head}$ |
| FBgn0051075 | *CG31075* | $PQ_{head}$ |
| FBgn0051183 | *CG31183* | $PQ_{head}$ |
| FBgn0051739 | *AspRS-m* | $PQ_{head}$ |
| FBgn0052099 | *CG32099* | $PQ_{head}$ |
| FBgn0052380 | *SMSr* | $PQ_{head}$ |
| FBgn0052484 | *Sk2* | $PQ_{head}$ |
| FBgn0052549 | *CG32549* | $PQ_{head}$ |
| FBgn0052750 | *CG32750* | $PQ_{head}$ |

| | | |
|---|---|---|
| FBgn0061359 | *Mvk* | $PQ_{head}$ |
| FBgn0067783 | *DPCoAC* | $PQ_{head}$ |
| FBgn0083956 | *CG34120* | $PQ_{head}$ |
| FBgn0085223 | *CG34194* | $PQ_{head}$ |
| FBgn0085322 | *CG34293* | $PQ_{head}$ |
| FBgn0085370 | *Pde11* | $PQ_{head}$ |
| FBgn0085386 | *CG34357* | $PQ_{head}$ |
| FBgn0086443 | *AsnRS* | $PQ_{head}$ |
| FBgn0086450 | *su(r)* | $PQ_{head}$ |
| FBgn0086532 | *Spt-I* | $PQ_{head}$ |
| FBgn0250837 | *dUTPase* | $PQ_{head}$ |
| FBgn0259171 | *Pde9* | $PQ_{head}$ |
| FBgn0260795 | *NaPi-III* | $PQ_{head}$ |
| FBgn0260960 | *Baldspot* | $PQ_{head}$ |
| FBgn0261244 | *inaE* | $PQ_{head}$ |
| FBgn0261266 | *zuc* | $PQ_{head}$ |
| FBgn0261436 | *DhpD* | $PQ_{head}$ |
| FBgn0261862 | *whd* | $PQ_{head}$ |
| FBgn0262112 | *sro* | $PQ_{head}$ |
| FBgn0262738 | *norpA* | $PQ_{head}$ |
| FBgn0263120 | *Acsl* | $PQ_{head}$ |
| FBgn0263131 | *CG43373* | $PQ_{head}$ |
| FBgn0263593 | *Lpin* | $PQ_{head}$ |
| FBgn0263607 | *l(3)72Dp* | $PQ_{head}$ |
| FBgn0263782 | *Hmgcr* | $PQ_{head}$ |
| FBgn0263916 | *Ent2* | $PQ_{head}$ |
| FBgn0264711 | *CG43980* | $PQ_{head}$ |
| FBgn0264815 | *Pde1c* | $PQ_{head}$ |
| FBgn0265178 | *CG44243* | $PQ_{head}$ |
| FBgn0265187 | *Fatp2* | $PQ_{head}$ |
| FBgn0266136 | *Gyc76C* | $PQ_{head}$ |
| FBgn0266377 | *Pde8* | $PQ_{head}$ |
| FBgn0267385 | *PyK* | $PQ_{head}$ |
| FBgn0270926 | *AsnS* | $PQ_{head}$ |
| FBgn0270928 | *VAChT* | $PQ_{head}$ |
| FBgn0275436 | *PheRS-m* | $PQ_{head}$ |
| FBgn0283427 | *FASN1* | $PQ_{head}$ |
| FBgn0284253 | *LeuRS* | $PQ_{head}$ |
| FBgn0286222 | *Fum1* | $PQ_{head}$ |
| FBgn0286511 | *Pld* | $PQ_{head}$ |

| | | |
|---|---|---|
| FBgn0015011 | *AhcyL2* | $PQ_{head}$ |
| FBgn0034898 | *CG18128* | $PQ_{head}$ |
| FBgn0026428 | *HDAC6* | $PQ_{head}$ |
| FBgn0038734 | *CG11453* | $PQ_{head}$ |
| FBgn0027601 | *pdgy* | $PQ_{head}$ |
| FBgn0026602 | *Adk3* | $PQ_{head}$ |
| FBgn0037995 | *Adk1* | $PQ_{head}$ |
| FBgn0037534 | *Elovl7* | $PQ_{head}$ |
| FBgn0037761 | *CG8534* | $PQ_{head}$ |
| FBgn0037762 | *eloF* | $PQ_{head}$ |
| FBgn0037765 | *CG9458* | $PQ_{head}$ |
| FBgn0038983 | *CG5326* | $PQ_{head}$ |
| FBgn0038986 | *sit* | $PQ_{head}$ |
| FBgn0051522 | *CG31522* | $PQ_{head}$ |
| FBgn0051523 | *CG31523* | $PQ_{head}$ |
| FBgn0053110 | *CG33110* | $PQ_{head}$ |
| FBgn0260942 | *bond* | $PQ_{head}$ |
| FBgn0052495 | *Gss2* | $PQ_{head}$ |
| FBgn0032522 | *CG16848* | $PQ_{head}$ |
| FBgn0051924 | *CG31924* | $PQ_{head}$ |
| FBgn0034105 | *CG7755* | $PQ_{head}$ |
| FBgn0035005 | *CG3483* | $PQ_{head}$ |
| FBgn0052026 | *CG32026* | $PQ_{head}$ |
| FBgn0032889 | *CG9331* | $PQ_{head}$ |
| FBgn0036403 | *CG6661* | $PQ_{head}$ |
| FBgn0005670 | *Cyp4d1* | $PQ_{head}$ |
| FBgn0010019 | *Cyp4g1* | $PQ_{head}$ |
| FBgn0015032 | *Cyp4c3* | $PQ_{head}$ |
| FBgn0015035 | *Cyp4e3* | $PQ_{head}$ |
| FBgn0015037 | *Cyp4p1* | $PQ_{head}$ |
| FBgn0031925 | *Cyp4d21* | $PQ_{head}$ |
| FBgn0011576 | *Cyp4d2* | $PQ_{head}$ |
| FBgn0014469 | *Cyp4e2* | $PQ_{head}$ |
| FBgn0015034 | *Cyp4e1* | $PQ_{head}$ |
| FBgn0015036 | *Cyp4ae1* | $PQ_{head}$ |
| FBgn0023541 | *Cyp4d14* | $PQ_{head}$ |
| FBgn0030304 | *Cyp4g15* | $PQ_{head}$ |
| FBgn0030367 | *Cyp311a1* | $PQ_{head}$ |
| FBgn0030615 | *Cyp4s3* | $PQ_{head}$ |
| FBgn0031693 | *Cyp4ac1* | $PQ_{head}$ |

| | | |
|---|---|---|
| FBgn0031694 | *Cyp4ac2* | $PQ_{head}$ |
| FBgn0031695 | *Cyp4ac3* | $PQ_{head}$ |
| FBgn0033292 | *Cyp4ad1* | $PQ_{head}$ |
| FBgn0033395 | *Cyp4p2* | $PQ_{head}$ |
| FBgn0033397 | *Cyp4p3* | $PQ_{head}$ |
| FBgn0034053 | *Cyp4aa1* | $PQ_{head}$ |
| FBgn0035344 | *Cyp4d20* | $PQ_{head}$ |
| FBgn0036778 | *Cyp312a1* | $PQ_{head}$ |
| FBgn0038006 | *Cyp313a2* | $PQ_{head}$ |
| FBgn0038076 | *Cyp313a4* | $PQ_{head}$ |
| FBgn0015781 | *P5cr* | $PQ_{head}$ |
| FBgn0029153 | *Menl-2* | $PQ_{head}$ |
| FBgn0029154 | *Menl-1* | $PQ_{head}$ |
| FBgn0034127 | *CG7848* | $PQ_{head}$ |
| FBgn0033538 | *CG11883* | $PQ_{head}$ |
| FBgn0286723 | *hll* | $PQ_{head}$ |
| FBgn0032061 | *CG9314* | $PQ_{head}$ |
| FBgn0029994 | *Ldsdh1* | $PQ_{head}$ |
| FBgn0032405 | *firl* | $PQ_{head}$ |
| FBgn0000055 | *Adh* | $PQ_{head}$ |
| FBgn0025454 | *Cyp6g1* | $PQ_{head}$ |
| FBgn0000473 | *Cyp6a2* | $PQ_{head}$ |
| FBgn0015714 | *Cyp6a17* | $PQ_{head}$ |
| FBgn0033302 | *Cyp6a14* | $PQ_{head}$ |
| FBgn0033978 | *Cyp6a23* | $PQ_{head}$ |
| FBgn0038037 | *Cyp9f2* | $PQ_{head}$ |
| FBgn0031489 | *CG17224* | $PQ_{head}$ |
| FBgn0030411 | *CG2540* | $PQ_{head}$ |
| FBgn0039109 | *CG10365* | $PQ_{head}$ |
| FBgn0028848 | *Gpo3* | $PQ_{head}$ |
| FBgn0033190 | *Gpo2* | $PQ_{head}$ |
| FBgn0034387 | *Cyp12b2* | $PQ_{head}$ |
| FBgn0015623 | *Cpr* | $PQ_{head}$ |
| FBgn0040512 | *ζCOP* | $PQ_{head}$ |
| FBgn0010348 | *Arf79F* | $PQ_{head}$ |
| FBgn0011703 | *RnrL* | $PQ_{head}$ |
| FBgn0011704 | *RnrS* | $PQ_{head}$ |
| FBgn0022029 | *l(2)k01209* | $PQ_{head}$ |
| FBgn0027945 | *ppl* | $PQ_{head}$ |
| FBgn0029950 | *CG9657* | $PQ_{head}$ |

| | | |
|---|---|---|
| FBgn0030482 | *CG1673* | $PQ_{head}$ |
| FBgn0031998 | *SLC5A11* | $PQ_{head}$ |
| FBgn0032123 | *Oatp30B* | $PQ_{head}$ |
| FBgn0032287 | *CG6415* | $PQ_{head}$ |
| FBgn0033524 | *Cyp49a1* | $PQ_{head}$ |
| FBgn0033543 | *Daao1* | $PQ_{head}$ |
| FBgn0034356 | *Pepck2* | $PQ_{head}$ |
| FBgn0034715 | *Oatp58Db* | $PQ_{head}$ |
| FBgn0036208 | *CG10361* | $PQ_{head}$ |
| FBgn0036666 | *TSG101* | $PQ_{head}$ |
| FBgn0036762 | *CG7430* | $PQ_{head}$ |
| FBgn0036806 | *Cyp12c1* | $PQ_{head}$ |
| FBgn0037801 | *CG3999* | $PQ_{head}$ |
| FBgn0040070 | *Trx-2* | $PQ_{head}$ |
| FBgn0043043 | *Desat2* | $PQ_{head}$ |
| FBgn0050277 | *Oatp58Da* | $PQ_{head}$ |
| FBgn0051262 | *CG31262* | $PQ_{head}$ |
| FBgn0051634 | *Oatp26F* | $PQ_{head}$ |
| FBgn0051668 | *CG31668* | $PQ_{head}$ |
| FBgn0053124 | *CG33124* | $PQ_{head}$ |
| FBgn0086687 | *Desat1* | $PQ_{head}$ |
| FBgn0250757 | *CG42235* | $PQ_{head}$ |
| FBgn0263398 | *Uck* | $PQ_{head}$ |
| FBgn0039071 | *bb8* | $PQ_{head}$ |
| FBgn0003067 | *Pepck1* | $PQ_{head}$ |
| FBgn0019982 | *Gs1l* | $PQ_{head}$ |
| FBgn0033856 | *CG13334* | $PQ_{head}$ |
| FBgn0031265 | *CG2794* | $PQ_{head}$ |
| FBgn0031860 | *Daao2* | $PQ_{head}$ |
| FBgn0036997 | *CG5955* | $PQ_{head}$ |
| FBgn0001128 | *Gpdh1* | $PQ_{body}$ |
| FBgn0010217 | *ATPsynβ* | $PQ_{body}$ |
| FBgn0010226 | *GstS1* | $PQ_{body}$ |
| FBgn0010383 | *Cyp18a1* | $PQ_{body}$ |
| FBgn0010612 | *ATPsynG* | $PQ_{body}$ |
| FBgn0011211 | *blw* | $PQ_{body}$ |
| FBgn0014391 | *sun* | $PQ_{body}$ |
| FBgn0016119 | *ATPsynCF6* | $PQ_{body}$ |
| FBgn0016120 | *ATPsynD* | $PQ_{body}$ |
| FBgn0016691 | *ATPsynO* | $PQ_{body}$ |

| | | |
|---|---|---|
| FBgn0019644 | *ATPsynB* | $PQ_{body}$ |
| FBgn0019830 | *colt* | $PQ_{body}$ |
| FBgn0020235 | *ATPsynγ* | $PQ_{body}$ |
| FBgn0021765 | *scu* | $PQ_{body}$ |
| FBgn0022160 | *Gpo1* | $PQ_{body}$ |
| FBgn0023023 | *CRMP* | $PQ_{body}$ |
| FBgn0025352 | *Mtpβ* | $PQ_{body}$ |
| FBgn0027348 | *bgm* | $PQ_{body}$ |
| FBgn0027572 | *ACOX1* | $PQ_{body}$ |
| FBgn0027844 | *CAH1* | $PQ_{body}$ |
| FBgn0028342 | *ATPsynδ* | $PQ_{body}$ |
| FBgn0028479 | *Mtpα* | $PQ_{body}$ |
| FBgn0030731 | *Mfe2* | $PQ_{body}$ |
| FBgn0031250 | *Ent1* | $PQ_{body}$ |
| FBgn0031813 | *Acox3* | $PQ_{body}$ |
| FBgn0032160 | *CG4598* | $PQ_{body}$ |
| FBgn0032219 | *CG4995* | $PQ_{body}$ |
| FBgn0033101 | *CG9436* | $PQ_{body}$ |
| FBgn0033246 | *ACC* | $PQ_{body}$ |
| FBgn0033542 | *CAH13* | $PQ_{body}$ |
| FBgn0033879 | *Echs1* | $PQ_{body}$ |
| FBgn0034432 | *CG7461* | $PQ_{body}$ |
| FBgn0034554 | *CAH14* | $PQ_{body}$ |
| FBgn0034560 | *CAH15* | $PQ_{body}$ |
| FBgn0034883 | *Eglp2* | $PQ_{body}$ |
| FBgn0034884 | *Eglp3* | $PQ_{body}$ |
| FBgn0035032 | *ATPsynF* | $PQ_{body}$ |
| FBgn0035169 | *Dci* | $PQ_{body}$ |
| FBgn0035383 | *CPT2* | $PQ_{body}$ |
| FBgn0035471 | *Sc2* | $PQ_{body}$ |
| FBgn0035476 | *CG12766* | $PQ_{body}$ |
| FBgn0035811 | *Mcad* | $PQ_{body}$ |
| FBgn0035915 | *S-Lap1* | $PQ_{body}$ |
| FBgn0036732 | *Oatp74D* | $PQ_{body}$ |
| FBgn0036821 | *CG3961* | $PQ_{body}$ |
| FBgn0037440 | *CRAT* | $PQ_{body}$ |
| FBgn0037513 | *pyd3* | $PQ_{body}$ |
| FBgn0037973 | *CG18547* | $PQ_{body}$ |
| FBgn0037974 | *CG12224* | $PQ_{body}$ |
| FBgn0037975 | *CG3397* | $PQ_{body}$ |

| | | |
|---|---|---|
| FBgn0038224 | *ATPsynE* | $PQ_{body}$ |
| FBgn0038469 | *CG4009* | $PQ_{body}$ |
| FBgn0039830 | *ATPsynC* | $PQ_{body}$ |
| FBgn0039858 | *CycG* | $PQ_{body}$ |
| FBgn0039890 | *Abcd1* | $PQ_{body}$ |
| FBgn0040064 | *yip2* | $PQ_{body}$ |
| FBgn0051477 | *ATPsynεL* | $PQ_{body}$ |
| FBgn0052351 | *S-Lap2* | $PQ_{body}$ |
| FBgn0086254 | *Akr1B* | $PQ_{body}$ |
| FBgn0260960 | *Baldspot* | $PQ_{body}$ |
| FBgn0261112 | *APP-BP1* | $PQ_{body}$ |
| FBgn0261397 | *didum* | $PQ_{body}$ |
| FBgn0261862 | *whd* | $PQ_{body}$ |
| FBgn0263120 | *Acsl* | $PQ_{body}$ |
| FBgn0263916 | *Ent2* | $PQ_{body}$ |
| FBgn0265187 | *Fatp2* | $PQ_{body}$ |
| FBgn0283531 | *Duox* | $PQ_{body}$ |
| FBgn0028935 | *CG7653* | $PQ_{body}$ |
| FBgn0030222 | *CG9806* | $PQ_{body}$ |
| FBgn0033860 | *S-Lap5* | $PQ_{body}$ |
| FBgn0033868 | *S-Lap7* | $PQ_{body}$ |
| FBgn0034132 | *S-Lap8* | $PQ_{body}$ |
| FBgn0038135 | *CG8773* | $PQ_{body}$ |
| FBgn0038136 | *CG8774* | $PQ_{body}$ |
| FBgn0038897 | *CG5849* | $PQ_{body}$ |
| FBgn0039640 | *superdeath* | $PQ_{body}$ |
| FBgn0039656 | *CG11951* | $PQ_{body}$ |
| FBgn0040493 | *grsm* | $PQ_{body}$ |
| FBgn0045770 | *S-Lap3* | $PQ_{body}$ |
| FBgn0046253 | *CG3502* | $PQ_{body}$ |
| FBgn0051198 | *CG31198* | $PQ_{body}$ |
| FBgn0051233 | *CG31233* | $PQ_{body}$ |
| FBgn0051343 | *CG31343* | $PQ_{body}$ |
| FBgn0051445 | *CG31445* | $PQ_{body}$ |
| FBgn0052064 | *S-Lap4* | $PQ_{body}$ |
| FBgn0052473 | *CG32473* | $PQ_{body}$ |
| FBgn0259237 | *CG42335* | $PQ_{body}$ |
| FBgn0259795 | *loopin-1* | $PQ_{body}$ |
| FBgn0261243 | *Psa* | $PQ_{body}$ |
| FBgn0263236 | *SP1029* | $PQ_{body}$ |

| | | |
|---|---|---|
| FBgn0285963 | *CG46339* | $PQ_{body}$ |
| FBgn0036910 | *Cyp305a1* | $PQ_{body}$ |
| FBgn0034825 | *Gpdh2* | $PQ_{body}$ |
| FBgn0263048 | *Gpdh3* | $PQ_{body}$ |
| FBgn0037534 | *Elovl7* | $PQ_{body}$ |
| FBgn0037761 | *CG8534* | $PQ_{body}$ |
| FBgn0037762 | *eloF* | $PQ_{body}$ |
| FBgn0037765 | *CG9458* | $PQ_{body}$ |
| FBgn0038983 | *CG5326* | $PQ_{body}$ |
| FBgn0038986 | *sit* | $PQ_{body}$ |
| FBgn0051522 | *CG31522* | $PQ_{body}$ |
| FBgn0051523 | *CG31523* | $PQ_{body}$ |
| FBgn0053110 | *CG33110* | $PQ_{body}$ |
| FBgn0260942 | *bond* | $PQ_{body}$ |
| FBgn0003486 | *spo* | $PQ_{body}$ |
| FBgn0011576 | *Cyp4d2* | $PQ_{body}$ |
| FBgn0030615 | *Cyp4s3* | $PQ_{body}$ |
| FBgn0035344 | *Cyp4d20* | $PQ_{body}$ |
| FBgn0025454 | *Cyp6g1* | $PQ_{body}$ |
| FBgn0033304 | *Cyp6a13* | $PQ_{body}$ |
| FBgn0000473 | *Cyp6a2* | $PQ_{body}$ |
| FBgn0013771 | *Cyp6a9* | $PQ_{body}$ |
| FBgn0013772 | *Cyp6a8* | $PQ_{body}$ |
| FBgn0013773 | *Cyp6a22* | $PQ_{body}$ |
| FBgn0015038 | *Cyp9b1* | $PQ_{body}$ |
| FBgn0015039 | *Cyp9b2* | $PQ_{body}$ |
| FBgn0015040 | *Cyp9c1* | $PQ_{body}$ |
| FBgn0015714 | *Cyp6a17* | $PQ_{body}$ |
| FBgn0028940 | *Cyp28a5* | $PQ_{body}$ |
| FBgn0031126 | *Cyp6v1* | $PQ_{body}$ |
| FBgn0031182 | *Cyp6t1* | $PQ_{body}$ |
| FBgn0031432 | *Cyp309a1* | $PQ_{body}$ |
| FBgn0032693 | *Cyp310a1* | $PQ_{body}$ |
| FBgn0033065 | *Cyp6w1* | $PQ_{body}$ |
| FBgn0033121 | *Cyp6u1* | $PQ_{body}$ |
| FBgn0033302 | *Cyp6a14* | $PQ_{body}$ |
| FBgn0033696 | *Cyp6g2* | $PQ_{body}$ |
| FBgn0033697 | *Cyp6t3* | $PQ_{body}$ |
| FBgn0033775 | *Cyp9h1* | $PQ_{body}$ |
| FBgn0033978 | *Cyp6a23* | $PQ_{body}$ |
| FBgn0033979 | *Cyp6a19* | $PQ_{body}$ |
| FBgn0033980 | *Cyp6a20* | $PQ_{body}$ |

| | | |
|---|---|---|
| FBgn0288232 | *Cyp6a21* | $PQ_{body}$ |
| FBgn0033982 | *Cyp317a1* | $PQ_{body}$ |
| FBgn0034756 | *Cyp6d2* | $PQ_{body}$ |
| FBgn0038037 | *Cyp9f2* | $PQ_{body}$ |
| FBgn0038194 | *Cyp6d5* | $PQ_{body}$ |
| FBgn0039006 | *Cyp6d4* | $PQ_{body}$ |
| FBgn0039519 | *Cyp6a18* | $PQ_{body}$ |
| FBgn0041337 | *Cyp309a2* | $PQ_{body}$ |
| FBgn0036568 | *ATPsynβL* | $PQ_{body}$ |
| FBgn0030411 | *CG2540* | $PQ_{body}$ |
| FBgn0039109 | *CG10365* | $PQ_{body}$ |
| FBgn0034387 | *Cyp12b2* | $PQ_{body}$ |
| FBgn0030223 | *CG2111* | $PQ_{body}$ |
| FBgn0013672 | *mt:ATPase6* | $PQ_{body}$ |
| FBgn0013673 | *mt:ATPase8* | $PQ_{body}$ |
| FBgn0031108 | *CG15459* | $PQ_{body}$ |
| FBgn0031941 | *ATPsynGL* | $PQ_{body}$ |
| FBgn0035585 | *ATPsynCF6L* | $PQ_{body}$ |
| FBgn0036345 | *CG17300* | $PQ_{body}$ |
| FBgn0040651 | *CG15458* | $PQ_{body}$ |
| FBgn0083167 | *Neb-cGP* | $PQ_{body}$ |
| FBgn0285943 | *knon* | $PQ_{body}$ |
| FBgn0016984 | *sktl* | $PQ_{body}$ |
| FBgn0028539 | *Eato* | $PQ_{body}$ |
| FBgn0030465 | *CG15743* | $PQ_{body}$ |
| FBgn0031170 | *Abca3* | $PQ_{body}$ |
| FBgn0031998 | *SLC5A11* | $PQ_{body}$ |
| FBgn0034493 | *CG8908* | $PQ_{body}$ |
| FBgn0034789 | *PIP5K59B* | $PQ_{body}$ |
| FBgn0035348 | *CG16758* | $PQ_{body}$ |
| FBgn0036058 | *CG6707* | $PQ_{body}$ |
| FBgn0036747 | *CG6052* | $PQ_{body}$ |
| FBgn0037852 | *Tpc1* | $PQ_{body}$ |
| FBgn0038074 | *Gnmt* | $PQ_{body}$ |
| FBgn0038302 | *CG4210* | $PQ_{body}$ |
| FBgn0039698 | *CG7789* | $PQ_{body}$ |
| FBgn0051262 | *CG31262* | $PQ_{body}$ |
| FBgn0051668 | *CG31668* | $PQ_{body}$ |
| FBgn0052099 | *CG32099* | $PQ_{body}$ |
| FBgn0053124 | *CG33124* | $PQ_{body}$ |
| FBgn0083956 | *CG34120* | $PQ_{body}$ |
| FBgn0086450 | *su(r)* | $PQ_{body}$ |

| | | |
|---|---|---|
| FBgn0250757 | *CG42235* | $PQ_{body}$ |
| FBgn0034898 | *CG18128* | $PQ_{body}$ |
| FBgn0026428 | *HDAC6* | $PQ_{body}$ |
| FBgn0032424 | *CG17010* | $PQ_{body}$ |
| FBgn0030510 | *CG12177* | $PQ_{body}$ |
| FBgn0032436 | *CG5418* | $PQ_{body}$ |
| FBgn0000406 | *Cyt-b5-r* | $\alpha Syn_{head}$ |
| FBgn0000422 | *Ddc* | $\alpha Syn_{head}$ |
| FBgn0000566 | *Eip55E* | $\alpha Syn_{head}$ |
| FBgn0002069 | *AspRS* | $\alpha Syn_{head}$ |
| FBgn0002778 | *mnd* | $\alpha Syn_{head}$ |
| FBgn0003076 | *Pgm1* | $\alpha Syn_{head}$ |
| FBgn0003141 | *pr* | $\alpha Syn_{head}$ |
| FBgn0003301 | *rut* | $\alpha Syn_{head}$ |
| FBgn0003416 | *sl* | $\alpha Syn_{head}$ |
| FBgn0003656 | *sws* | $\alpha Syn_{head}$ |
| FBgn0003965 | *v* | $\alpha Syn_{head}$ |
| FBgn0004516 | *Gad1* | $\alpha Syn_{head}$ |
| FBgn0004611 | *Plc21C* | $\alpha Syn_{head}$ |
| FBgn0004852 | *Ac76E* | $\alpha Syn_{head}$ |
| FBgn0005674 | *GluProRS* | $\alpha Syn_{head}$ |
| FBgn0010197 | *Gyc32E* | $\alpha Syn_{head}$ |
| FBgn0010217 | *ATPsynβ* | $\alpha Syn_{head}$ |
| FBgn0010352 | *Nc73EF* | $\alpha Syn_{head}$ |
| FBgn0010383 | *Cyp18a1* | $\alpha Syn_{head}$ |
| FBgn0010611 | *Hmgs* | $\alpha Syn_{head}$ |
| FBgn0010612 | *ATPsynG* | $\alpha Syn_{head}$ |
| FBgn0010803 | *TrpRS* | $\alpha Syn_{head}$ |
| FBgn0011211 | *blw* | $\alpha Syn_{head}$ |
| FBgn0012034 | *AcCoAS* | $\alpha Syn_{head}$ |
| FBgn0013972 | *Gycα99B* | $\alpha Syn_{head}$ |
| FBgn0013973 | *Gycβ100B* | $\alpha Syn_{head}$ |
| FBgn0014032 | *Sptr* | $\alpha Syn_{head}$ |
| FBgn0014391 | *sun* | $\alpha Syn_{head}$ |
| FBgn0016119 | *ATPsynCF6* | $\alpha Syn_{head}$ |
| FBgn0016120 | *ATPsynD* | $\alpha Syn_{head}$ |
| FBgn0016126 | *CaMKI* | $\alpha Syn_{head}$ |
| FBgn0016687 | *Nurf-38* | $\alpha Syn_{head}$ |
| FBgn0016691 | *ATPsynO* | $\alpha Syn_{head}$ |
| FBgn0016694 | *Pdp1* | $\alpha Syn_{head}$ |
| FBgn0019644 | *ATPsynB* | $\alpha Syn_{head}$ |
| FBgn0019830 | *colt* | $\alpha Syn_{head}$ |

| | | |
|---|---|---|
| FBgn0020235 | *ATPsynγ* | $\alpha Syn_{head}$ |
| FBgn0020545 | *kraken* | $\alpha Syn_{head}$ |
| FBgn0021750 | *SerRS-m* | $\alpha Syn_{head}$ |
| FBgn0021765 | *scu* | $\alpha Syn_{head}$ |
| FBgn0021795 | *Tapδ* | $\alpha Syn_{head}$ |
| FBgn0022709 | *Ak1* | $\alpha Syn_{head}$ |
| FBgn0022710 | *Ac13E* | $\alpha Syn_{head}$ |
| FBgn0023416 | *Ac3* | $\alpha Syn_{head}$ |
| FBgn0024150 | *Ac78C* | $\alpha Syn_{head}$ |
| FBgn0024320 | *Npc1a* | $\alpha Syn_{head}$ |
| FBgn0024728 | *Slip1* | $\alpha Syn_{head}$ |
| FBgn0024947 | *NTPase* | $\alpha Syn_{head}$ |
| FBgn0025352 | *Mtpβ* | $\alpha Syn_{head}$ |
| FBgn0025592 | *Gk1* | $\alpha Syn_{head}$ |
| FBgn0025740 | *PlexB* | $\alpha Syn_{head}$ |
| FBgn0025807 | *Rad9* | $\alpha Syn_{head}$ |
| FBgn0025885 | *Inos* | $\alpha Syn_{head}$ |
| FBgn0026630 | *nes* | $\alpha Syn_{head}$ |
| FBgn0026778 | *Rad1* | $\alpha Syn_{head}$ |
| FBgn0027079 | *ValRS* | $\alpha Syn_{head}$ |
| FBgn0027080 | *TyrRS* | $\alpha Syn_{head}$ |
| FBgn0027081 | *ThrRS* | $\alpha Syn_{head}$ |
| FBgn0027083 | *MetRS-m* | $\alpha Syn_{head}$ |
| FBgn0027084 | *LysRS* | $\alpha Syn_{head}$ |
| FBgn0027085 | *LeuRS-m* | $\alpha Syn_{head}$ |
| FBgn0027086 | *IleRS* | $\alpha Syn_{head}$ |
| FBgn0027087 | *HisRS* | $\alpha Syn_{head}$ |
| FBgn0027088 | *GlyRS* | $\alpha Syn_{head}$ |
| FBgn0027090 | *GlnRS* | $\alpha Syn_{head}$ |
| FBgn0027091 | *CysRS* | $\alpha Syn_{head}$ |
| FBgn0027093 | *ArgRS* | $\alpha Syn_{head}$ |
| FBgn0027094 | *AlaRS* | $\alpha Syn_{head}$ |
| FBgn0027348 | *bgm* | $\alpha Syn_{head}$ |
| FBgn0027579 | *mino* | $\alpha Syn_{head}$ |
| FBgn0027844 | *CAH1* | $\alpha Syn_{head}$ |
| FBgn0028342 | *ATPsynδ* | $\alpha Syn_{head}$ |
| FBgn0028425 | *JhI-21* | $\alpha Syn_{head}$ |
| FBgn0028479 | *Mtpα* | $\alpha Syn_{head}$ |
| FBgn0028539 | *Eato* | $\alpha Syn_{head}$ |
| FBgn0028707 | *Mt2* | $\alpha Syn_{head}$ |
| FBgn0028916 | *CG33090* | $\alpha Syn_{head}$ |
| FBgn0028962 | *AlaRS-m* | $\alpha Syn_{head}$ |

| | | |
|---|---|---|
| FBgn0029689 | *CG6428* | $\alpha Syn_{head}$ |
| FBgn0029706 | *CG3626* | $\alpha Syn_{head}$ |
| FBgn0029969 | *CG10932* | $\alpha Syn_{head}$ |
| FBgn0029975 | *spidey* | $\alpha Syn_{head}$ |
| FBgn0030007 | *α-PheRS* | $\alpha Syn_{head}$ |
| FBgn0030013 | *GIIIspla2* | $\alpha Syn_{head}$ |
| FBgn0030300 | *Sk1* | $\alpha Syn_{head}$ |
| FBgn0030407 | *Fpgs* | $\alpha Syn_{head}$ |
| FBgn0030421 | *Agpat1* | $\alpha Syn_{head}$ |
| FBgn0030638 | *CG11655* | $\alpha Syn_{head}$ |
| FBgn0030683 | *Mvd* | $\alpha Syn_{head}$ |
| FBgn0030743 | *CG9921* | $\alpha Syn_{head}$ |
| FBgn0030756 | *CG9903* | $\alpha Syn_{head}$ |
| FBgn0030816 | *CG16700* | $\alpha Syn_{head}$ |
| FBgn0030817 | *CG4991* | $\alpha Syn_{head}$ |
| FBgn0031048 | *CG12237* | $\alpha Syn_{head}$ |
| FBgn0031148 | *Cbs* | $\alpha Syn_{head}$ |
| FBgn0031170 | *Abca3* | $\alpha Syn_{head}$ |
| FBgn0031213 | *galectin* | $\alpha Syn_{head}$ |
| FBgn0031214 | *CG11374* | $\alpha Syn_{head}$ |
| FBgn0031250 | *Ent1* | $\alpha Syn_{head}$ |
| FBgn0031289 | *CG13950* | $\alpha Syn_{head}$ |
| FBgn0031484 | *CG3165* | $\alpha Syn_{head}$ |
| FBgn0031497 | *SerRS* | $\alpha Syn_{head}$ |
| FBgn0031735 | *CG11029* | $\alpha Syn_{head}$ |
| FBgn0032083 | *CG9541* | $\alpha Syn_{head}$ |
| FBgn0032160 | *CG4598* | $\alpha Syn_{head}$ |
| FBgn0032219 | *CG4995* | $\alpha Syn_{head}$ |
| FBgn0032394 | *Hacd1* | $\alpha Syn_{head}$ |
| FBgn0032524 | *Hacd2* | $\alpha Syn_{head}$ |
| FBgn0032603 | *CG17928* | $\alpha Syn_{head}$ |
| FBgn0032726 | *CG10621* | $\alpha Syn_{head}$ |
| FBgn0032727 | *CG10623* | $\alpha Syn_{head}$ |
| FBgn0032811 | *Pmvk* | $\alpha Syn_{head}$ |
| FBgn0032845 | *CG10747* | $\alpha Syn_{head}$ |
| FBgn0032923 | *Pld3* | $\alpha Syn_{head}$ |
| FBgn0033101 | *CG9436* | $\alpha Syn_{head}$ |
| FBgn0033226 | *puml* | $\alpha Syn_{head}$ |
| FBgn0033246 | *ACC* | $\alpha Syn_{head}$ |
| FBgn0033377 | *Pgm2a* | $\alpha Syn_{head}$ |
| FBgn0033476 | *oys* | $\alpha Syn_{head}$ |
| FBgn0033542 | *CAH13* | $\alpha Syn_{head}$ |

| | | |
|---|---|---|
| FBgn0033754 | *Ak6* | $\alpha Syn_{head}$ |
| FBgn0033836 | *CG18278* | $\alpha Syn_{head}$ |
| FBgn0033844 | *bbc* | $\alpha Syn_{head}$ |
| FBgn0033879 | *Echs1* | $\alpha Syn_{head}$ |
| FBgn0033900 | *CysRS-m* | $\alpha Syn_{head}$ |
| FBgn0033911 | *VGAT* | $\alpha Syn_{head}$ |
| FBgn0034117 | *CG7997* | $\alpha Syn_{head}$ |
| FBgn0034177 | *AsnRS-m* | $\alpha Syn_{head}$ |
| FBgn0034276 | *Sardh* | $\alpha Syn_{head}$ |
| FBgn0034364 | *CG5493* | $\alpha Syn_{head}$ |
| FBgn0034401 | *MetRS* | $\alpha Syn_{head}$ |
| FBgn0034419 | *CG15111* | $\alpha Syn_{head}$ |
| FBgn0034432 | *CG7461* | $\alpha Syn_{head}$ |
| FBgn0034491 | *Hsl* | $\alpha Syn_{head}$ |
| FBgn0034493 | *CG8908* | $\alpha Syn_{head}$ |
| FBgn0034554 | *CAH14* | $\alpha Syn_{head}$ |
| FBgn0034560 | *CAH15* | $\alpha Syn_{head}$ |
| FBgn0034884 | *Eglp3* | $\alpha Syn_{head}$ |
| FBgn0034887 | *St1* | $\alpha Syn_{head}$ |
| FBgn0034971 | *Gpat4* | $\alpha Syn_{head}$ |
| FBgn0034997 | *CG3376* | $\alpha Syn_{head}$ |
| FBgn0035032 | *ATPsynF* | $\alpha Syn_{head}$ |
| FBgn0035064 | *TyrRS-m* | $\alpha Syn_{head}$ |
| FBgn0035147 | *Gale* | $\alpha Syn_{head}$ |
| FBgn0035165 | *CG13887* | $\alpha Syn_{head}$ |
| FBgn0035169 | *Dci* | $\alpha Syn_{head}$ |
| FBgn0035203 | *Acat2* | $\alpha Syn_{head}$ |
| FBgn0035383 | *CPT2* | $\alpha Syn_{head}$ |
| FBgn0035392 | *CG1271* | $\alpha Syn_{head}$ |
| FBgn0035421 | *nSMase* | $\alpha Syn_{head}$ |
| FBgn0035471 | *Sc2* | $\alpha Syn_{head}$ |
| FBgn0035476 | *CG12766* | $\alpha Syn_{head}$ |
| FBgn0035811 | *Mcad* | $\alpha Syn_{head}$ |
| FBgn0035942 | *ValRS-m* | $\alpha Syn_{head}$ |
| FBgn0036030 | *Prps* | $\alpha Syn_{head}$ |
| FBgn0036053 | *iPLA2-VIA* | $\alpha Syn_{head}$ |
| FBgn0036099 | *CG11811* | $\alpha Syn_{head}$ |
| FBgn0036319 | *Ent3* | $\alpha Syn_{head}$ |
| FBgn0036366 | *JMJD7* | $\alpha Syn_{head}$ |
| FBgn0036449 | *bmm* | $\alpha Syn_{head}$ |
| FBgn0036545 | *GXIVsPLA2* | $\alpha Syn_{head}$ |
| FBgn0036569 | *IleRS-m* | $\alpha Syn_{head}$ |

| | | |
|---|---|---|
| FBgn0036622 | *Agpat4* | $\alpha Syn_{head}$ |
| FBgn0036623 | *Agpat3* | $\alpha Syn_{head}$ |
| FBgn0036629 | *GluRS-m* | $\alpha Syn_{head}$ |
| FBgn0036659 | *CG9701* | $\alpha Syn_{head}$ |
| FBgn0036691 | *beg* | $\alpha Syn_{head}$ |
| FBgn0036698 | *CG7724* | $\alpha Syn_{head}$ |
| FBgn0036747 | *CG6052* | $\alpha Syn_{head}$ |
| FBgn0036762 | *CG7430* | $\alpha Syn_{head}$ |
| FBgn0036763 | *TrpRS-m* | $\alpha Syn_{head}$ |
| FBgn0036816 | *Indy* | $\alpha Syn_{head}$ |
| FBgn0036821 | *CG3961* | $\alpha Syn_{head}$ |
| FBgn0036844 | *Mkp3* | $\alpha Syn_{head}$ |
| FBgn0036927 | *Gabat* | $\alpha Syn_{head}$ |
| FBgn0036939 | *CG7365* | $\alpha Syn_{head}$ |
| FBgn0037203 | *slif* | $\alpha Syn_{head}$ |
| FBgn0037440 | *CRAT* | $\alpha Syn_{head}$ |
| FBgn0037526 | *ArgRS-m* | $\alpha Syn_{head}$ |
| FBgn0037684 | *Srr* | $\alpha Syn_{head}$ |
| FBgn0037759 | *CG8526* | $\alpha Syn_{head}$ |
| FBgn0037852 | *Tpc1* | $\alpha Syn_{head}$ |
| FBgn0037891 | *CG5214* | $\alpha Syn_{head}$ |
| FBgn0037900 | *CG5276* | $\alpha Syn_{head}$ |
| FBgn0037958 | *CG6962* | $\alpha Syn_{head}$ |
| FBgn0038224 | *ATPsynE* | $\alpha Syn_{head}$ |
| FBgn0039175 | *β-PheRS* | $\alpha Syn_{head}$ |
| FBgn0039184 | *CG6432* | $\alpha Syn_{head}$ |
| FBgn0039357 | *CG4743* | $\alpha Syn_{head}$ |
| FBgn0039358 | *CG5028* | $\alpha Syn_{head}$ |
| FBgn0039475 | *CG6277* | $\alpha Syn_{head}$ |
| FBgn0039655 | *CG14507* | $\alpha Syn_{head}$ |
| FBgn0039774 | *CDase* | $\alpha Syn_{head}$ |
| FBgn0039799 | *CG15543* | $\alpha Syn_{head}$ |
| FBgn0039830 | *ATPsynC* | $\alpha Syn_{head}$ |
| FBgn0039844 | *CG1607* | $\alpha Syn_{head}$ |
| FBgn0039858 | *CycG* | $\alpha Syn_{head}$ |
| FBgn0039890 | *Abcd1* | $\alpha Syn_{head}$ |
| FBgn0040064 | *yip2* | $\alpha Syn_{head}$ |
| FBgn0040212 | *Gnpat* | $\alpha Syn_{head}$ |
| FBgn0040271 | *Sulf1* | $\alpha Syn_{head}$ |
| FBgn0040918 | *schlank* | $\alpha Syn_{head}$ |
| FBgn0041150 | *hoe1* | $\alpha Syn_{head}$ |
| FBgn0041342 | *Pcyt1* | $\alpha Syn_{head}$ |

| | | |
|---|---|---|
| FBgn0042094 | *Ak3* | αSyn$_{head}$ |
| FBgn0042138 | *Apt1* | αSyn$_{head}$ |
| FBgn0042175 | *CG18858* | αSyn$_{head}$ |
| FBgn0042627 | *FASN2* | αSyn$_{head}$ |
| FBgn0043043 | *Desat2* | αSyn$_{head}$ |
| FBgn0045064 | *bwa* | αSyn$_{head}$ |
| FBgn0051148 | *Gba1a* | αSyn$_{head}$ |
| FBgn0051183 | *CG31183* | αSyn$_{head}$ |
| FBgn0051414 | *Gba1b* | αSyn$_{head}$ |
| FBgn0051477 | *ATPsynεL* | αSyn$_{head}$ |
| FBgn0051739 | *AspRS-m* | αSyn$_{head}$ |
| FBgn0052191 | *CG32191* | αSyn$_{head}$ |
| FBgn0052380 | *SMSr* | αSyn$_{head}$ |
| FBgn0052484 | *Sk2* | αSyn$_{head}$ |
| FBgn0052699 | *LPCAT* | αSyn$_{head}$ |
| FBgn0061359 | *Mvk* | αSyn$_{head}$ |
| FBgn0083956 | *CG34120* | αSyn$_{head}$ |
| FBgn0085386 | *CG34357* | αSyn$_{head}$ |
| FBgn0086443 | *AsnRS* | αSyn$_{head}$ |
| FBgn0086687 | *Desat1* | αSyn$_{head}$ |
| FBgn0260475 | *CG30059* | αSyn$_{head}$ |
| FBgn0260750 | *Mulk* | αSyn$_{head}$ |
| FBgn0260960 | *Baldspot* | αSyn$_{head}$ |
| FBgn0261112 | *APP-BP1* | αSyn$_{head}$ |
| FBgn0261244 | *inaE* | αSyn$_{head}$ |
| FBgn0261266 | *zuc* | αSyn$_{head}$ |
| FBgn0261862 | *whd* | αSyn$_{head}$ |
| FBgn0262738 | *norpA* | αSyn$_{head}$ |
| FBgn0263120 | *Acsl* | αSyn$_{head}$ |
| FBgn0263131 | *CG43373* | αSyn$_{head}$ |
| FBgn0263607 | *l(3)72Dp* | αSyn$_{head}$ |
| FBgn0263782 | *Hmgcr* | αSyn$_{head}$ |
| FBgn0263916 | *Ent2* | αSyn$_{head}$ |
| FBgn0264975 | *Nrg* | αSyn$_{head}$ |
| FBgn0265187 | *Fatp2* | αSyn$_{head}$ |
| FBgn0266136 | *Gyc76C* | αSyn$_{head}$ |
| FBgn0275436 | *PheRS-m* | αSyn$_{head}$ |
| FBgn0283427 | *FASN1* | αSyn$_{head}$ |
| FBgn0283494 | *Ak2* | αSyn$_{head}$ |
| FBgn0284253 | *LeuRS* | αSyn$_{head}$ |
| FBgn0286511 | *Pld* | αSyn$_{head}$ |
| FBgn0286813 | *SRPK* | αSyn$_{head}$ |

| | | |
|---|---|---|
| FBgn0287585 | *Pss* | αSyn$_{head}$ |
| FBgn0287695 | *KFase* | αSyn$_{head}$ |
| FBgn0033673 | *CG8298* | αSyn$_{head}$ |
| FBgn0035266 | *Gk2* | αSyn$_{head}$ |
| FBgn0030339 | *Cyp28c1* | αSyn$_{head}$ |
| FBgn0031688 | *Cyp28d2* | αSyn$_{head}$ |
| FBgn0031689 | *Cyp28d1* | αSyn$_{head}$ |
| FBgn0038236 | *Cyp313a1* | αSyn$_{head}$ |
| FBgn0050446 | *Tdc2* | αSyn$_{head}$ |
| FBgn0259977 | *Tdc1* | αSyn$_{head}$ |
| FBgn0033969 | *Pgm2b* | αSyn$_{head}$ |
| FBgn0037534 | *Elovl7* | αSyn$_{head}$ |
| FBgn0037761 | *CG8534* | αSyn$_{head}$ |
| FBgn0037762 | *eloF* | αSyn$_{head}$ |
| FBgn0037765 | *CG9458* | αSyn$_{head}$ |
| FBgn0038983 | *CG5326* | αSyn$_{head}$ |
| FBgn0038986 | *sit* | αSyn$_{head}$ |
| FBgn0051522 | *CG31522* | αSyn$_{head}$ |
| FBgn0051523 | *CG31523* | αSyn$_{head}$ |
| FBgn0053110 | *CG33110* | αSyn$_{head}$ |
| FBgn0260942 | *bond* | αSyn$_{head}$ |
| FBgn0003486 | *spo* | αSyn$_{head}$ |
| FBgn0039737 | *CG7920* | αSyn$_{head}$ |
| FBgn0031759 | *Kdm5* | αSyn$_{head}$ |
| FBgn0011576 | *Cyp4d2* | αSyn$_{head}$ |
| FBgn0030615 | *Cyp4s3* | αSyn$_{head}$ |
| FBgn0035344 | *Cyp4d20* | αSyn$_{head}$ |
| FBgn0001624 | *dlg1* | αSyn$_{head}$ |
| FBgn0050021 | *metro* | αSyn$_{head}$ |
| FBgn0250785 | *vari* | αSyn$_{head}$ |
| FBgn0261873 | *sdt* | αSyn$_{head}$ |
| FBgn0286723 | *hll* | αSyn$_{head}$ |
| FBgn0039790 | *CG2246* | αSyn$_{head}$ |
| FBgn0035240 | *CG33791* | αSyn$_{head}$ |
| FBgn0031860 | *Daao2* | αSyn$_{head}$ |
| FBgn0032424 | *CG17010* | αSyn$_{head}$ |
| FBgn0025454 | *Cyp6g1* | αSyn$_{head}$ |
| FBgn0000473 | *Cyp6a2* | αSyn$_{head}$ |
| FBgn0015714 | *Cyp6a17* | αSyn$_{head}$ |
| FBgn0033302 | *Cyp6a14* | αSyn$_{head}$ |
| FBgn0033978 | *Cyp6a23* | αSyn$_{head}$ |
| FBgn0038037 | *Cyp9f2* | αSyn$_{head}$ |

| | | |
|---|---|---|
| FBgn0036568 | *ATPsynβL* | $\alpha Syn_{head}$ |
| FBgn0039094 | *CG10184* | $\alpha Syn_{head}$ |
| FBgn0031592 | *Art2* | $\alpha Syn_{head}$ |
| FBgn0038189 | *Art6* | $\alpha Syn_{head}$ |
| FBgn0052152 | *CG32152* | $\alpha Syn_{head}$ |
| FBgn0038295 | *Gyc88E* | $\alpha Syn_{head}$ |
| FBgn0038435 | *Gyc89Da* | $\alpha Syn_{head}$ |
| FBgn0038436 | *Gyc89Db* | $\alpha Syn_{head}$ |
| FBgn0261363 | *PPO3* | $\alpha Syn_{head}$ |
| FBgn0283437 | *PPO1* | $\alpha Syn_{head}$ |
| FBgn0013672 | *mt:ATPase6* | $\alpha Syn_{head}$ |
| FBgn0013673 | *mt:ATPase8* | $\alpha Syn_{head}$ |
| FBgn0031108 | *CG15459* | $\alpha Syn_{head}$ |
| FBgn0031941 | *ATPsynGL* | $\alpha Syn_{head}$ |
| FBgn0035585 | *ATPsynCF6L* | $\alpha Syn_{head}$ |
| FBgn0036345 | *CG17300* | $\alpha Syn_{head}$ |
| FBgn0040651 | *CG15458* | $\alpha Syn_{head}$ |
| FBgn0083167 | *Neb-cGP* | $\alpha Syn_{head}$ |
| FBgn0285943 | *knon* | $\alpha Syn_{head}$ |
| FBgn0000221 | *brn* | $\alpha Syn_{head}$ |
| FBgn0001970 | *Pgant35A* | $\alpha Syn_{head}$ |
| FBgn0004087 | *Dhfr* | $\alpha Syn_{head}$ |
| FBgn0004797 | *mdy* | $\alpha Syn_{head}$ |
| FBgn0010329 | *Tbh* | $\alpha Syn_{head}$ |
| FBgn0011205 | *fbl* | $\alpha Syn_{head}$ |
| FBgn0011336 | *Stt3B* | $\alpha Syn_{head}$ |
| FBgn0014075 | *Uggt* | $\alpha Syn_{head}$ |
| FBgn0015277 | *Pi3K59F* | $\alpha Syn_{head}$ |
| FBgn0020764 | *Alas* | $\alpha Syn_{head}$ |
| FBgn0024994 | *Ugalt* | $\alpha Syn_{head}$ |
| FBgn0026379 | *Pten* | $\alpha Syn_{head}$ |
| FBgn0027538 | *β4GalNAcTA* | $\alpha Syn_{head}$ |
| FBgn0028741 | *fab1* | $\alpha Syn_{head}$ |
| FBgn0030558 | *CG1461* | $\alpha Syn_{head}$ |
| FBgn0030574 | *sbm* | $\alpha Syn_{head}$ |
| FBgn0030930 | *Pgant7* | $\alpha Syn_{head}$ |
| FBgn0031149 | *Stt3A* | $\alpha Syn_{head}$ |
| FBgn0031491 | *α4GT1* | $\alpha Syn_{head}$ |
| FBgn0031663 | *CG8891* | $\alpha Syn_{head}$ |
| FBgn0031682 | *CG5828* | $\alpha Syn_{head}$ |
| FBgn0031765 | *CG9109* | $\alpha Syn_{head}$ |
| FBgn0031986 | *CG8673* | $\alpha Syn_{head}$ |

| | | |
|---|---|---|
| FBgn0032001 | *CG8360* | $\alpha Syn_{head}$ |
| FBgn0032002 | *CG8353* | $\alpha Syn_{head}$ |
| FBgn0032014 | *CG7840* | $\alpha Syn_{head}$ |
| FBgn0032117 | *FucTB* | $\alpha Syn_{head}$ |
| FBgn0032449 | *CG17036* | $\alpha Syn_{head}$ |
| FBgn0033214 | *CG1941* | $\alpha Syn_{head}$ |
| FBgn0033215 | *Dgat2* | $\alpha Syn_{head}$ |
| FBgn0033216 | *CG1946* | $\alpha Syn_{head}$ |
| FBgn0033524 | *Cyp49a1* | $\alpha Syn_{head}$ |
| FBgn0034649 | *PIG-M* | $\alpha Syn_{head}$ |
| FBgn0034715 | *Oatp58Db* | $\alpha Syn_{head}$ |
| FBgn0034716 | *Oatp58Dc* | $\alpha Syn_{head}$ |
| FBgn0035464 | *PIG-B* | $\alpha Syn_{head}$ |
| FBgn0036058 | *CG6707* | $\alpha Syn_{head}$ |
| FBgn0036157 | *CG7560* | $\alpha Syn_{head}$ |
| FBgn0036446 | *Mgat4a* | $\alpha Syn_{head}$ |
| FBgn0036485 | *FucTA* | $\alpha Syn_{head}$ |
| FBgn0036732 | *Oatp74D* | $\alpha Syn_{head}$ |
| FBgn0036806 | *Cyp12c1* | $\alpha Syn_{head}$ |
| FBgn0037170 | *Trxr-2* | $\alpha Syn_{head}$ |
| FBgn0037845 | *CG14694* | $\alpha Syn_{head}$ |
| FBgn0037846 | *CG6574* | $\alpha Syn_{head}$ |
| FBgn0038552 | *Alg1* | $\alpha Syn_{head}$ |
| FBgn0039258 | *β4GalT7* | $\alpha Syn_{head}$ |
| FBgn0039378 | *α4GT2* | $\alpha Syn_{head}$ |
| FBgn0039580 | *Gfat2* | $\alpha Syn_{head}$ |
| FBgn0039625 | *β4GalNAcTB* | $\alpha Syn_{head}$ |
| FBgn0050036 | *CG30036* | $\alpha Syn_{head}$ |
| FBgn0050037 | *CG30037* | $\alpha Syn_{head}$ |
| FBgn0050277 | *Oatp58Da* | $\alpha Syn_{head}$ |
| FBgn0050290 | *Ppcdc* | $\alpha Syn_{head}$ |
| FBgn0053145 | *GalT1* | $\alpha Syn_{head}$ |
| FBgn0086253 | *rumi* | $\alpha Syn_{head}$ |
| FBgn0259166 | *CG42271* | $\alpha Syn_{head}$ |
| FBgn0261403 | *sxc* | $\alpha Syn_{head}$ |
| FBgn0265174 | *PIG-V* | $\alpha Syn_{head}$ |
| FBgn0266438 | *PIG-Z* | $\alpha Syn_{head}$ |
| FBgn0287209 | *Gfat1* | $\alpha Syn_{head}$ |
| FBgn0035619 | *Alp10* | $\alpha Syn_{head}$ |
| FBgn0035620 | *Alp9* | $\alpha Syn_{head}$ |
| FBgn0038845 | *Alp5* | $\alpha Syn_{head}$ |
| FBgn0026314 | *Ugt35B1* | $\alpha Syn_{head}$ |

| | | |
|---|---|---|
| FBgn0027073 | *Ugt49B1* | αSyn$_{head}$ |
| FBgn0027074 | *Ugt36F1* | αSyn$_{head}$ |
| FBgn0032684 | *Ugt301D1* | αSyn$_{head}$ |
| FBgn0051002 | *Ugt35D1* | αSyn$_{head}$ |
| FBgn0027070 | *Ugt36E1* | αSyn$_{head}$ |
| FBgn0270927 | *βGlu* | αSyn$_{head}$ |
| FBgn0040257 | *Ugt302E1* | αSyn$_{head}$ |
| FBgn0001098 | *Gdh* | αSyn$_{body}$ |
| FBgn0001124 | *Got1* | αSyn$_{body}$ |
| FBgn0001128 | *Gpdh1* | αSyn$_{body}$ |
| FBgn0002069 | *AspRS* | αSyn$_{body}$ |
| FBgn0002778 | *mnd* | αSyn$_{body}$ |
| FBgn0003116 | *pn* | αSyn$_{body}$ |
| FBgn0003141 | *pr* | αSyn$_{body}$ |
| FBgn0004516 | *Gad1* | αSyn$_{body}$ |
| FBgn0005674 | *GluProRS* | αSyn$_{body}$ |
| FBgn0010383 | *Cyp18a1* | αSyn$_{body}$ |
| FBgn0010548 | *Aldh-III* | αSyn$_{body}$ |
| FBgn0010611 | *Hmgs* | αSyn$_{body}$ |
| FBgn0011205 | *fbl* | αSyn$_{body}$ |
| FBgn0011768 | *Fdh* | αSyn$_{body}$ |
| FBgn0014930 | *CG2846* | αSyn$_{body}$ |
| FBgn0015872 | *Drip* | αSyn$_{body}$ |
| FBgn0016687 | *Nurf-38* | αSyn$_{body}$ |
| FBgn0017482 | *T3dh* | αSyn$_{body}$ |
| FBgn0019662 | *qm* | αSyn$_{body}$ |
| FBgn0020236 | *ATPCL* | αSyn$_{body}$ |
| FBgn0020930 | *Dgkε* | αSyn$_{body}$ |
| FBgn0021750 | *SerRS-m* | αSyn$_{body}$ |
| FBgn0022160 | *Gpo1* | αSyn$_{body}$ |
| FBgn0022359 | *Sodh-2* | αSyn$_{body}$ |
| FBgn0022709 | *Ak1* | αSyn$_{body}$ |
| FBgn0024289 | *Sodh-1* | αSyn$_{body}$ |
| FBgn0024995 | *CG2680* | αSyn$_{body}$ |
| FBgn0025373 | *Fpps* | αSyn$_{body}$ |
| FBgn0025592 | *Gk1* | αSyn$_{body}$ |
| FBgn0025807 | *Rad9* | αSyn$_{body}$ |
| FBgn0025837 | *CG17636* | αSyn$_{body}$ |
| FBgn0026778 | *Rad1* | αSyn$_{body}$ |
| FBgn0027079 | *ValRS* | αSyn$_{body}$ |
| FBgn0027080 | *TyrRS* | αSyn$_{body}$ |
| FBgn0027083 | *MetRS-m* | αSyn$_{body}$ |

| | | |
|---|---|---|
| FBgn0027084 | *LysRS* | $\alpha Syn_{body}$ |
| FBgn0027085 | *LeuRS-m* | $\alpha Syn_{body}$ |
| FBgn0027088 | *GlyRS* | $\alpha Syn_{body}$ |
| FBgn0027093 | *ArgRS* | $\alpha Syn_{body}$ |
| FBgn0027094 | *AlaRS* | $\alpha Syn_{body}$ |
| FBgn0027583 | *CG7601* | $\alpha Syn_{body}$ |
| FBgn0027610 | *Dic1* | $\alpha Syn_{body}$ |
| FBgn0028425 | *JhI-21* | $\alpha Syn_{body}$ |
| FBgn0028962 | *AlaRS-m* | $\alpha Syn_{body}$ |
| FBgn0029502 | *Coq7* | $\alpha Syn_{body}$ |
| FBgn0029848 | *Btnd* | $\alpha Syn_{body}$ |
| FBgn0030361 | *CG1492* | $\alpha Syn_{body}$ |
| FBgn0030407 | *Fpgs* | $\alpha Syn_{body}$ |
| FBgn0030431 | *CG4407* | $\alpha Syn_{body}$ |
| FBgn0030460 | *Coq5* | $\alpha Syn_{body}$ |
| FBgn0030465 | *CG15743* | $\alpha Syn_{body}$ |
| FBgn0030683 | *Mvd* | $\alpha Syn_{body}$ |
| FBgn0030796 | *CG4829* | $\alpha Syn_{body}$ |
| FBgn0030816 | *CG16700* | $\alpha Syn_{body}$ |
| FBgn0030817 | *CG4991* | $\alpha Syn_{body}$ |
| FBgn0030932 | *Ggt-1* | $\alpha Syn_{body}$ |
| FBgn0031250 | *Ent1* | $\alpha Syn_{body}$ |
| FBgn0031484 | *CG3165* | $\alpha Syn_{body}$ |
| FBgn0031497 | *SerRS* | $\alpha Syn_{body}$ |
| FBgn0031630 | *CG15629* | $\alpha Syn_{body}$ |
| FBgn0031663 | *CG8891* | $\alpha Syn_{body}$ |
| FBgn0031682 | *CG5828* | $\alpha Syn_{body}$ |
| FBgn0031713 | *Coq6* | $\alpha Syn_{body}$ |
| FBgn0031877 | *Hmgcl* | $\alpha Syn_{body}$ |
| FBgn0032083 | *CG9541* | $\alpha Syn_{body}$ |
| FBgn0032449 | *CG17036* | $\alpha Syn_{body}$ |
| FBgn0032811 | *Pmvk* | $\alpha Syn_{body}$ |
| FBgn0032922 | *Coq3* | $\alpha Syn_{body}$ |
| FBgn0033101 | *CG9436* | $\alpha Syn_{body}$ |
| FBgn0033203 | *CG2070* | $\alpha Syn_{body}$ |
| FBgn0033204 | *CG2065* | $\alpha Syn_{body}$ |
| FBgn0033205 | *CG2064* | $\alpha Syn_{body}$ |
| FBgn0033368 | *CG13743* | $\alpha Syn_{body}$ |
| FBgn0033373 | *CG8080* | $\alpha Syn_{body}$ |
| FBgn0033679 | *CG8888* | $\alpha Syn_{body}$ |
| FBgn0033754 | *Ak6* | $\alpha Syn_{body}$ |
| FBgn0033761 | *CG8778* | $\alpha Syn_{body}$ |

| | | |
|---|---|---|
| FBgn0033853 | *CG6145* | αSyn$_{body}$ |
| FBgn0033911 | *VGAT* | αSyn$_{body}$ |
| FBgn0034401 | *MetRS* | αSyn$_{body}$ |
| FBgn0034497 | *Mpcp1* | αSyn$_{body}$ |
| FBgn0034715 | *Oatp58Db* | αSyn$_{body}$ |
| FBgn0034733 | *CG4752* | αSyn$_{body}$ |
| FBgn0034883 | *Eglp2* | αSyn$_{body}$ |
| FBgn0034884 | *Eglp3* | αSyn$_{body}$ |
| FBgn0034887 | *St1* | αSyn$_{body}$ |
| FBgn0034997 | *CG3376* | αSyn$_{body}$ |
| FBgn0035064 | *TyrRS-m* | αSyn$_{body}$ |
| FBgn0035082 | *CG2811* | αSyn$_{body}$ |
| FBgn0035083 | *Tina-1* | αSyn$_{body}$ |
| FBgn0035203 | *Acat2* | αSyn$_{body}$ |
| FBgn0035348 | *CG16758* | αSyn$_{body}$ |
| FBgn0035392 | *CG1271* | αSyn$_{body}$ |
| FBgn0035421 | *nSMase* | αSyn$_{body}$ |
| FBgn0035839 | *CG7550* | αSyn$_{body}$ |
| FBgn0035915 | *S-Lap1* | αSyn$_{body}$ |
| FBgn0035942 | *ValRS-m* | αSyn$_{body}$ |
| FBgn0036381 | *CG8745* | αSyn$_{body}$ |
| FBgn0036428 | *Gbs-70E* | αSyn$_{body}$ |
| FBgn0036629 | *GluRS-m* | αSyn$_{body}$ |
| FBgn0036698 | *CG7724* | αSyn$_{body}$ |
| FBgn0036732 | *Oatp74D* | αSyn$_{body}$ |
| FBgn0036770 | *Prestin* | αSyn$_{body}$ |
| FBgn0036787 | *CG4306* | αSyn$_{body}$ |
| FBgn0037044 | *Pdss2* | αSyn$_{body}$ |
| FBgn0037138 | *P5CDh1* | αSyn$_{body}$ |
| FBgn0037203 | *slif* | αSyn$_{body}$ |
| FBgn0037370 | *CG1236* | αSyn$_{body}$ |
| FBgn0037526 | *ArgRS-m* | αSyn$_{body}$ |
| FBgn0037574 | *Coq2* | αSyn$_{body}$ |
| FBgn0037845 | *CG14694* | αSyn$_{body}$ |
| FBgn0037846 | *CG6574* | αSyn$_{body}$ |
| FBgn0037852 | *Tpc1* | αSyn$_{body}$ |
| FBgn0037958 | *CG6962* | αSyn$_{body}$ |
| FBgn0038302 | *CG4210* | αSyn$_{body}$ |
| FBgn0038376 | *Hmt-1* | αSyn$_{body}$ |
| FBgn0038516 | *P5cr-2* | αSyn$_{body}$ |
| FBgn0038876 | *Idi* | αSyn$_{body}$ |
| FBgn0038925 | *Cchl* | αSyn$_{body}$ |

| | | |
|---|---|---|
| FBgn0039357 | *CG4743* | $\alpha Syn_{body}$ |
| FBgn0039698 | *CG7789* | $\alpha Syn_{body}$ |
| FBgn0039799 | *CG15543* | $\alpha Syn_{body}$ |
| FBgn0039844 | *CG1607* | $\alpha Syn_{body}$ |
| FBgn0039846 | *PNPase* | $\alpha Syn_{body}$ |
| FBgn0039858 | *CycG* | $\alpha Syn_{body}$ |
| FBgn0041150 | *hoe1* | $\alpha Syn_{body}$ |
| FBgn0042094 | *Ak3* | $\alpha Syn_{body}$ |
| FBgn0050345 | *CG30345* | $\alpha Syn_{body}$ |
| FBgn0050394 | *CG30394* | $\alpha Syn_{body}$ |
| FBgn0051005 | *qless* | $\alpha Syn_{body}$ |
| FBgn0051075 | *CG31075* | $\alpha Syn_{body}$ |
| FBgn0051140 | *CG31140* | $\alpha Syn_{body}$ |
| FBgn0051739 | *AspRS-m* | $\alpha Syn_{body}$ |
| FBgn0052099 | *CG32099* | $\alpha Syn_{body}$ |
| FBgn0052351 | *S-Lap2* | $\alpha Syn_{body}$ |
| FBgn0052380 | *SMSr* | $\alpha Syn_{body}$ |
| FBgn0052750 | *CG32750* | $\alpha Syn_{body}$ |
| FBgn0053156 | *CG33156* | $\alpha Syn_{body}$ |
| FBgn0061359 | *Mvk* | $\alpha Syn_{body}$ |
| FBgn0067783 | *DPCoAC* | $\alpha Syn_{body}$ |
| FBgn0085390 | *Dgk* | $\alpha Syn_{body}$ |
| FBgn0085413 | *CG34384* | $\alpha Syn_{body}$ |
| FBgn0086254 | *Akr1B* | $\alpha Syn_{body}$ |
| FBgn0261549 | *rdgA* | $\alpha Syn_{body}$ |
| FBgn0263607 | *l(3)72Dp* | $\alpha Syn_{body}$ |
| FBgn0263782 | *Hmgcr* | $\alpha Syn_{body}$ |
| FBgn0263916 | *Ent2* | $\alpha Syn_{body}$ |
| FBgn0265178 | *CG44243* | $\alpha Syn_{body}$ |
| FBgn0266064 | *GlyS* | $\alpha Syn_{body}$ |
| FBgn0270926 | *AsnS* | $\alpha Syn_{body}$ |
| FBgn0283494 | *Ak2* | $\alpha Syn_{body}$ |
| FBgn0283531 | *Duox* | $\alpha Syn_{body}$ |
| FBgn0284253 | *LeuRS* | $\alpha Syn_{body}$ |
| FBgn0285912 | *mah* | $\alpha Syn_{body}$ |
| FBgn0028935 | *CG7653* | $\alpha Syn_{body}$ |
| FBgn0030222 | *CG9806* | $\alpha Syn_{body}$ |
| FBgn0033860 | *S-Lap5* | $\alpha Syn_{body}$ |
| FBgn0033868 | *S-Lap7* | $\alpha Syn_{body}$ |
| FBgn0034132 | *S-Lap8* | $\alpha Syn_{body}$ |
| FBgn0038135 | *CG8773* | $\alpha Syn_{body}$ |
| FBgn0038136 | *CG8774* | $\alpha Syn_{body}$ |

| | | |
|---|---|---|
| FBgn0038897 | *CG5849* | αSyn$_{body}$ |
| FBgn0039640 | *superdeath* | αSyn$_{body}$ |
| FBgn0039656 | *CG11951* | αSyn$_{body}$ |
| FBgn0040493 | *grsm* | αSyn$_{body}$ |
| FBgn0045770 | *S-Lap3* | αSyn$_{body}$ |
| FBgn0046253 | *CG3502* | αSyn$_{body}$ |
| FBgn0051198 | *CG31198* | αSyn$_{body}$ |
| FBgn0051233 | *CG31233* | αSyn$_{body}$ |
| FBgn0051343 | *CG31343* | αSyn$_{body}$ |
| FBgn0051445 | *CG31445* | αSyn$_{body}$ |
| FBgn0052064 | *S-Lap4* | αSyn$_{body}$ |
| FBgn0052473 | *CG32473* | αSyn$_{body}$ |
| FBgn0259237 | *CG42335* | αSyn$_{body}$ |
| FBgn0259795 | *loopin-1* | αSyn$_{body}$ |
| FBgn0261243 | *Psa* | αSyn$_{body}$ |
| FBgn0263236 | *SP1029* | αSyn$_{body}$ |
| FBgn0285963 | *CG46339* | αSyn$_{body}$ |
| FBgn0033673 | *CG8298* | αSyn$_{body}$ |
| FBgn0035266 | *Gk2* | αSyn$_{body}$ |
| FBgn0030339 | *Cyp28c1* | αSyn$_{body}$ |
| FBgn0031688 | *Cyp28d2* | αSyn$_{body}$ |
| FBgn0031689 | *Cyp28d1* | αSyn$_{body}$ |
| FBgn0038236 | *Cyp313a1* | αSyn$_{body}$ |
| FBgn0034898 | *CG18128* | αSyn$_{body}$ |
| FBgn0026428 | *HDAC6* | αSyn$_{body}$ |
| FBgn0038734 | *CG11453* | αSyn$_{body}$ |
| FBgn0027601 | *pdgy* | αSyn$_{body}$ |
| FBgn0058064 | *ARY* | αSyn$_{body}$ |
| FBgn0003486 | *spo* | αSyn$_{body}$ |
| FBgn0032522 | *CG16848* | αSyn$_{body}$ |
| FBgn0051924 | *CG31924* | αSyn$_{body}$ |
| FBgn0036403 | *CG6661* | αSyn$_{body}$ |
| FBgn0011576 | *Cyp4d2* | αSyn$_{body}$ |
| FBgn0030615 | *Cyp4s3* | αSyn$_{body}$ |
| FBgn0035344 | *Cyp4d20* | αSyn$_{body}$ |
| FBgn0015781 | *P5cr* | αSyn$_{body}$ |
| FBgn0038610 | *CG7675* | αSyn$_{body}$ |
| FBgn0050495 | *CG30495* | αSyn$_{body}$ |
| FBgn0025454 | *Cyp6g1* | αSyn$_{body}$ |
| FBgn0000473 | *Cyp6a2* | αSyn$_{body}$ |
| FBgn0015714 | *Cyp6a17* | αSyn$_{body}$ |
| FBgn0033302 | *Cyp6a14* | αSyn$_{body}$ |

| | | |
|---|---|---|
| FBgn0033978 | *Cyp6a23* | $\alpha Syn_{body}$ |
| FBgn0038037 | *Cyp9f2* | $\alpha Syn_{body}$ |
| FBgn0027552 | *CG10863* | $\alpha Syn_{body}$ |
| FBgn0036183 | *CG6083* | $\alpha Syn_{body}$ |
| FBgn0036290 | *CG10638* | $\alpha Syn_{body}$ |
| FBgn0030223 | *CG2111* | $\alpha Syn_{body}$ |
| FBgn0000303 | *ChAT* | $\alpha Syn_{body}$ |
| FBgn0010241 | *Mdr50* | $\alpha Syn_{body}$ |
| FBgn0011703 | *RnrL* | $\alpha Syn_{body}$ |
| FBgn0011704 | *RnrS* | $\alpha Syn_{body}$ |
| FBgn0013307 | *Odc1* | $\alpha Syn_{body}$ |
| FBgn0014032 | *Sptr* | $\alpha Syn_{body}$ |
| FBgn0016672 | *Ipp* | $\alpha Syn_{body}$ |
| FBgn0022029 | *l(2)k01209* | $\alpha Syn_{body}$ |
| FBgn0022338 | *dnk* | $\alpha Syn_{body}$ |
| FBgn0024920 | *Ts* | $\alpha Syn_{body}$ |
| FBgn0025709 | *CNT2* | $\alpha Syn_{body}$ |
| FBgn0026630 | *nes* | $\alpha Syn_{body}$ |
| FBgn0028539 | *Eato* | $\alpha Syn_{body}$ |
| FBgn0030013 | *GIIIspla2* | $\alpha Syn_{body}$ |
| FBgn0030574 | *sbm* | $\alpha Syn_{body}$ |
| FBgn0031170 | *Abca3* | $\alpha Syn_{body}$ |
| FBgn0031735 | *CG11029* | $\alpha Syn_{body}$ |
| FBgn0033371 | *CNT1* | $\alpha Syn_{body}$ |
| FBgn0034493 | *CG8908* | $\alpha Syn_{body}$ |
| FBgn0034716 | *Oatp58Dc* | $\alpha Syn_{body}$ |
| FBgn0036053 | *iPLA2-VIA* | $\alpha Syn_{body}$ |
| FBgn0036208 | *CG10361* | $\alpha Syn_{body}$ |
| FBgn0036366 | *JMJD7* | $\alpha Syn_{body}$ |
| FBgn0036545 | *GXIVsPLA2* | $\alpha Syn_{body}$ |
| FBgn0036747 | *CG6052* | $\alpha Syn_{body}$ |
| FBgn0036834 | *CG6836* | $\alpha Syn_{body}$ |
| FBgn0036939 | *CG7365* | $\alpha Syn_{body}$ |
| FBgn0037063 | *CG9391* | $\alpha Syn_{body}$ |
| FBgn0039655 | *CG14507* | $\alpha Syn_{body}$ |
| FBgn0040070 | *Trx-2* | $\alpha Syn_{body}$ |
| FBgn0050277 | *Oatp58Da* | $\alpha Syn_{body}$ |
| FBgn0051634 | *Oatp26F* | $\alpha Syn_{body}$ |
| FBgn0052699 | *LPCAT* | $\alpha Syn_{body}$ |
| FBgn0083956 | *CG34120* | $\alpha Syn_{body}$ |
| FBgn0261625 | *GLS* | $\alpha Syn_{body}$ |
| FBgn0263398 | *Uck* | $\alpha Syn_{body}$ |

| | | |
|---|---|---|
| FBgn0287585 | *Pss* | $\alpha Syn_{body}$ |
| FBgn0035619 | *Alp10* | $\alpha Syn_{body}$ |
| FBgn0035620 | *Alp9* | $\alpha Syn_{body}$ |
| FBgn0038845 | *Alp5* | $\alpha Syn_{body}$ |
| FBgn0000024 | *Ace* | $\alpha Syn_{body}$ |
| FBgn0001987 | *Gli* | $\alpha Syn_{body}$ |
| FBgn0031327 | *CG5397* | $\alpha Syn_{body}$ |
| FBgn0032132 | *CG4382* | $\alpha Syn_{body}$ |
| FBgn0030949 | *Cyp308a1* | $\alpha Syn_{body}$ |
| FBgn0034387 | *Cyp12b2* | $\alpha Syn_{body}$ |
| FBgn0036997 | *CG5955* | $\alpha Syn_{body}$ |
