## Supplemental Table 2 for "Comparative Proteomic Analysis of Environmental and Genetic Models of Parkinson’s Disease Highlights the Role of Purine Metabolism"

**Table S2 |** List of the metabolites in the ΔFBA- and reaction activity analysis-predicted differential reactions for paraquat treatment or α-synuclein expression to induce Parkinson's disease-related changes in the

| Metabolite | Metabolite Name | KEGG ID | PD Group |
|---|---|---|---|
| m02819[p] | pyruvate | C00022 | $PQ_{head}$ |
| m02403[p] | L-lactate | C00186 | $PQ_{head}$ |
| m03161[c] | glycogen | C00182 | $PQ_{head}$ |
| m02148[c] | hydroxyacetone | C05235 | $PQ_{head}$ |
| m01256[c] | acetone | C00207 | $PQ_{head}$ |
| m00671[m] | 2-oxobutyrate | C00109 | $PQ_{head}$ |
| m01253[m] | acetoacetate | C00164 | $PQ_{head}$ |
| m01256[m] | acetone | C00207 | $PQ_{head}$ |
| m01255[c] | acetoacetyl-CoA | C00332 | $PQ_{head}$ |
| m01253[c] | acetoacetate | C00164 | $PQ_{head}$ |
| m00157[m] | (R)-3-hydroxybutanoate | C01089 | $PQ_{head}$ |
| m01419[c] | cAMP | C00575 | $PQ_{head}$ |
| m01280[m] | adenosine | C00212 | $PQ_{head}$ |
| m01639[c] | dAMP | C00360 | $PQ_{head}$ |
| m01645[c] | dCTP | C00458 | $PQ_{head}$ |
| m01756[c] | dUTP | C00460 | $PQ_{head}$ |
| m01643[c] | dCDP | C00705 | $PQ_{head}$ |
| m01754[c] | dUDP | C01346 | $PQ_{head}$ |
| m00266[c] | 10-formyl-THF | C00234 | $PQ_{head}$ |
| m02147[c] | hydroxide | C01328 | $PQ_{head}$ |
| m01642[m] | dATP | C00131 | $PQ_{head}$ |
| m01637[m] | dADP | C00206 | $PQ_{head}$ |
| m01688[m] | dGTP | C00286 | $PQ_{head}$ |
| m01754[m] | dUDP | C01346 | $PQ_{head}$ |
| m01756[m] | dUTP | C00460 | $PQ_{head}$ |
| m01747[c] | dTDP | C00363 | $PQ_{head}$ |
| m01753[m] | dTTP | C00459 | $PQ_{head}$ |
| m01747[m] | dTDP | C00363 | $PQ_{head}$ |
| m02046[c] | HCO3- | C00288 | $PQ_{head}$ |
| m01052[c] | 5,6-Dihydrouracil | C00429 | $PQ_{head}$ |
| m00923[c] | 3-ureidopropionate | C02642 | $PQ_{head}$ |
| m02997[c] | thymine | C00178 | $PQ_{head}$ |
| m01705[c] | dihydrothymine | C00906 | $PQ_{head}$ |
| m01644[c] | dCMP | C00239 | $PQ_{head}$ |
| m01755[m] | dUMP | C00365 | $PQ_{head}$ |
| m01045[c] | 5,10-methylene-THF | C00143 | $PQ_{head}$ |
| m00922[c] | 3-ureidoisobutyrate | C05100 | $PQ_{head}$ |
| m01633[c] | D-3-amino-isobutanoate | C01205 | $PQ_{head}$ |
| m01721[c] | DNA | C00039 | $PQ_{head}$ |
| m02751[l] | Pi | C00009 | $PQ_{head}$ |

| | | | |
|---|---|---|---|
| m01645[m] | dCTP | C00458 | $PQ_{head}$ |
| m01643[m] | dCDP | C00705 | $PQ_{head}$ |
| m02578[m] | NH3 | C00014 | $PQ_{head}$ |
| m01588[c] | citrulline | C00327 | $PQ_{head}$ |
| m01369[c] | asparagine | C00152 | $PQ_{head}$ |
| m01588[m] | citrulline | C00327 | $PQ_{head}$ |
| m02125[c] | histidine | C00135 | $PQ_{head}$ |
| m02981[c] | THF-hexaglutamate | | $PQ_{head}$ |
| m02184[c] | isoleucine | C00407 | $PQ_{head}$ |
| m02360[c] | leucine | C00123 | $PQ_{head}$ |
| m01698[c] | dihydrobiopterin | C00268 | $PQ_{head}$ |
| m01005[c] | 4-hydroxyphenylpyruvate | C01179 | $PQ_{head}$ |
| m01004[c] | 4-hydroxyphenyllactate | C03672 | $PQ_{head}$ |
| m00996[c] | 4-hydroxybenzoyl-CoA | C02949 | $PQ_{head}$ |
| m00995[c] | 4-hydroxybenzoate | C00156 | $PQ_{head}$ |
| m01115[c] | 5-methyl-THF | C00440 | $PQ_{head}$ |
| m01721[n] | DNA | C00039 | $PQ_{head}$ |
| m01974[s] | glutamate | C00025 | $PQ_{head}$ |
| m00097[s] | (5-L-glutamyl)-L-amino acid | C03740 | $PQ_{head}$ |
| m01383[c] | beta-alanine | C00099 | $PQ_{head}$ |
| m03075[c] | tRNA(met) | C01647 | $PQ_{head}$ |
| m02131[m] | HMG-CoA | C00356 | $PQ_{head}$ |
| m03081[c] | tRNA(tyr) | C00787 | $PQ_{head}$ |
| m02421[c] | L-tyrosyl-tRNA(tyr) | C02839 | $PQ_{head}$ |
| m03063[c] | tRNA(ala) | C01635 | $PQ_{head}$ |
| m02335[c] | L-alanyl-tRNA(ala) | C00886 | $PQ_{head}$ |
| m03064[c] | tRNA(arg) | C01636 | $PQ_{head}$ |
| m02340[c] | L-arginyl-tRNA(arg) | C02163 | $PQ_{head}$ |
| m03065[c] | tRNA(asn) | C01637 | $PQ_{head}$ |
| m02341[c] | L-asparaginyl-tRNA(asn) | C03402 | $PQ_{head}$ |
| m03066[c] | tRNA(asp) | C01638 | $PQ_{head}$ |
| m02342[c] | L-aspartyl-tRNA(asp) | C02984 | $PQ_{head}$ |
| m03067[c] | tRNA(cys) | C01639 | $PQ_{head}$ |
| m02351[c] | L-cysteinyl-tRNA(cys) | C03125 | $PQ_{head}$ |
| m03068[c] | tRNA(gln) | C01640 | $PQ_{head}$ |
| m02376[c] | L-glutaminyl-tRNA(gln) | C02282 | $PQ_{head}$ |
| m03069[c] | tRNA(glu) | C01641 | $PQ_{head}$ |
| m02377[c] | L-glutamyl-tRNA(glu) | C02987 | $PQ_{head}$ |
| m03070[c] | tRNA(gly) | C01642 | $PQ_{head}$ |
| m02006[c] | glycyl-tRNA(gly) | C02412 | $PQ_{head}$ |
| m03071[c] | tRNA(his) | C01643 | $PQ_{head}$ |
| m02380[c] | L-histidyl-tRNA(his) | C02988 | $PQ_{head}$ |
| m03072[c] | tRNA(ile) | C01644 | $PQ_{head}$ |

| | | | |
|---|---|---|---|
| m02401[c] | L-isoleucyl-tRNA(ile) | C03127 | $PQ_{head}$ |
| m03073[c] | tRNA(leu) | C01645 | $PQ_{head}$ |
| m02404[c] | L-leucyl-tRNA(leu) | C02047 | $PQ_{head}$ |
| m03074[c] | tRNA(lys) | C01646 | $PQ_{head}$ |
| m02405[c] | L-lysyl-tRNA(lys) | C01931 | $PQ_{head}$ |
| m02408[c] | L-methionyl-tRNA(met) | C02430 | $PQ_{head}$ |
| m03076[c] | tRNA(phe) | C01648 | $PQ_{head}$ |
| m02412[c] | L-phenylalanyl-tRNA(phe) | C03511 | $PQ_{head}$ |
| m03077[c] | tRNA(pro) | C01649 | $PQ_{head}$ |
| m02415[c] | L-prolyl-tRNA(pro) | C02702 | $PQ_{head}$ |
| m03078[c] | tRNA(ser) | C01650 | $PQ_{head}$ |
| m02416[c] | L-seryl-tRNA(ser) | C02553 | $PQ_{head}$ |
| m03079[c] | tRNA(thr) | C01651 | $PQ_{head}$ |
| m02419[c] | L-threonyl-tRNA(thr) | C02992 | $PQ_{head}$ |
| m03080[c] | tRNA(trp) | C01652 | $PQ_{head}$ |
| m02420[c] | L-tryptophanyl-tRNA(trp) | C03512 | $PQ_{head}$ |
| m03082[c] | tRNA(val) | C01653 | $PQ_{head}$ |
| m02423[c] | L-valyl-tRNA(val) | C02554 | $PQ_{head}$ |
| m00919[c] | 3-sulfinylpyruvic acid | C05527 | $PQ_{head}$ |
| m02183[m] | isocitrate | C00311 | $PQ_{head}$ |
| m02344[c] | lauric acid | C02679 | $PQ_{head}$ |
| m03051[c] | tridecylic acid | C17076 | $PQ_{head}$ |
| m03050[c] | tridecanoyl-CoA | | $PQ_{head}$ |
| m00128[c] | (9E)-tetradecenoic acid | | $PQ_{head}$ |
| m00129[c] | (9E)-tetradecenoyl-CoA | | $PQ_{head}$ |
| m00117[c] | (7Z)-tetradecenoic acid | | $PQ_{head}$ |
| m00118[c] | (7Z)-tetradecenoyl-CoA | | $PQ_{head}$ |
| m02745[c] | physeteric acid | | $PQ_{head}$ |
| m01141[c] | 5-tetradecenoyl-CoA | | $PQ_{head}$ |
| m02690[c] | pentadecylic acid | C16537 | $PQ_{head}$ |
| m02689[c] | pentadecanoyl-CoA | | $PQ_{head}$ |
| m02675[c] | palmitolate | C08362 | $PQ_{head}$ |
| m01197[c] | 7-palmitoleic acid | | $PQ_{head}$ |
| m01191[c] | 7-hexadecenoyl-CoA | | $PQ_{head}$ |
| m02456[c] | margaric acid | | $PQ_{head}$ |
| m02101[c] | heptadecanoyl-CoA | | $PQ_{head}$ |
| m00003[c] | (10Z)-heptadecenoic acid | | $PQ_{head}$ |
| m00004[c] | (10Z)-heptadecenoyl-CoA | | $PQ_{head}$ |
| m01238[c] | 9-heptadecylenic acid | C16536 | $PQ_{head}$ |
| m01237[c] | 9-heptadecenoyl-CoA | | $PQ_{head}$ |
| m02938[c] | stearate | C01530 | $PQ_{head}$ |
| m00019[c] | (13Z)-octadecenoic acid | | $PQ_{head}$ |
| m00020[c] | (13Z)-octadecenoyl-CoA | | $PQ_{head}$ |

| | | | |
|---|---|---|---|
| m01585[c] | cis-vaccenic acid | C08367 | $PQ_{head}$ |
| m01586[c] | cis-vaccenoyl-CoA | C21945 | $PQ_{head}$ |
| m02646[c] | oleate | C00712 | $PQ_{head}$ |
| m01778[c] | elaidate | C00712 | $PQ_{head}$ |
| m00127[c] | (9E)-octadecenoyl-CoA | | $PQ_{head}$ |
| m00115[c] | (7Z)-octadecenoic acid | | $PQ_{head}$ |
| m00104[c] | (6Z,9Z)-octadecadienoic acid | | $PQ_{head}$ |
| m00106[c] | (6Z,9Z)-octadecadienoyl-CoA | | $PQ_{head}$ |
| m02613[c] | nonadecylic acid | C16535 | $PQ_{head}$ |
| m02612[c] | nonadecanoyl-CoA | | $PQ_{head}$ |
| m01771[c] | eicosanoate | C06425 | $PQ_{head}$ |
| m01773[c] | eicosanoyl-CoA | C02041 | $PQ_{head}$ |
| m00017[c] | (13Z)-eicosenoic acid | | $PQ_{head}$ |
| m00018[c] | (13Z)-eicosenoyl-CoA | | $PQ_{head}$ |
| m01584[c] | cis-gondoic acid | C16526 | $PQ_{head}$ |
| m00007[c] | (11Z)-eicosenoyl-CoA | C16530 | $PQ_{head}$ |
| m01235[c] | 9-eicosenoic acid | | $PQ_{head}$ |
| m01236[c] | 9-eicosenoyl-CoA | | $PQ_{head}$ |
| m01207[c] | 8,11-eicosadienoic acid | | $PQ_{head}$ |
| m00123[c] | (8Z,11Z)-eicosadienoyl-CoA | C21937 | $PQ_{head}$ |
| m02457[c] | mead acid | | $PQ_{head}$ |
| m00101[c] | (5Z,8Z,11Z)-eicosatrienoyl-CoA | C21939 | $PQ_{head}$ |
| m02053[c] | henicosanoic acid | | $PQ_{head}$ |
| m02052[c] | heneicosanoyl-CoA | | $PQ_{head}$ |
| m01373[c] | behenic acid | C08281 | $PQ_{head}$ |
| m01725[c] | docosanoyl-CoA | C16528 | $PQ_{head}$ |
| m01583[c] | cis-erucic acid | C08316 | $PQ_{head}$ |
| m00016[c] | (13Z)-docosenoyl-CoA | C16531 | $PQ_{head}$ |
| m01582[c] | cis-cetoleic acid | | $PQ_{head}$ |
| m00006[c] | (11Z)-docosenoyl-CoA | | $PQ_{head}$ |
| m03045[c] | tricosanoic acid | | $PQ_{head}$ |
| m03047[c] | tricosanoyl-CoA | | $PQ_{head}$ |
| m02385[c] | lignocerate | C08320 | $PQ_{head}$ |
| m02971[c] | tetracosanoyl-CoA | C16529 | $PQ_{head}$ |
| m02564[c] | nervonic acid | C08323 | $PQ_{head}$ |
| m01432[c] | cerotic acid | | $PQ_{head}$ |
| m03153[c] | ximenic acid | | $PQ_{head}$ |
| m02112[c] | hexacosenoyl-CoA | | $PQ_{head}$ |
| m02389[c] | linolenate | C06427 | $PQ_{head}$ |
| m02939[c] | stearidonic acid | C16300 | $PQ_{head}$ |
| m00108[c] | (6Z,9Z,12Z,15Z)-octadecatetraenoyl-CoA | C16163 | $PQ_{head}$ |
| m02648[c] | omega-3-arachidonic acid | | $PQ_{head}$ |
| m00125[c] | (8Z,11Z,14Z,17Z)-eicosatetraenoyl-CoA | C16164 | $PQ_{head}$ |

| | | | |
|---|---|---|---|
| m01784[c] | EPA | C06428 | $PQ_{head}$ |
| m00103[c] | (5Z,8Z,11Z,14Z,17Z)-eicosapentaenoyl-CoA | C16165 | $PQ_{head}$ |
| m01741[c] | DPA | C16513 | $PQ_{head}$ |
| m00121[c] | (7Z,10Z,13Z,16Z,19Z)-docosapentaenoyl-CoA | C16166 | $PQ_{head}$ |
| m00135[c] | (9Z,12Z,15Z,18Z,21Z)-TPA | | $PQ_{head}$ |
| m00134[c] | (9Z,12Z,15Z,18Z,21Z)-tetracosapentaenoyl-CoA | C16167 | $PQ_{head}$ |
| m00114[c] | (6Z,9Z,12Z,15Z,18Z,21Z)-THA | | $PQ_{head}$ |
| m01689[c] | DHA | C06429 | $PQ_{head}$ |
| m00010[c] | (11Z,14Z,17Z)-eicosatrienoic acid | C16522 | $PQ_{head}$ |
| m00012[c] | (11Z,14Z,17Z)-eicosatrienoyl-CoA | C16179 | $PQ_{head}$ |
| m00341[c] | 13,16,19-docosatrienoic acid | C16534 | $PQ_{head}$ |
| m00343[c] | 13,16,19-docosatrienoyl-CoA | | $PQ_{head}$ |
| m00260[c] | 10,13,16,19-docosatetraenoic acid | | $PQ_{head}$ |
| m00262[c] | 10,13,16,19-docosatetraenoyl-CoA | | $PQ_{head}$ |
| m00315[c] | 12,15,18,21-tetracosatetraenoic acid | | $PQ_{head}$ |
| m00317[c] | 12,15,18,21-tetracosatetraenoyl-CoA | | $PQ_{head}$ |
| m02387[c] | linoleate | C01595 | $PQ_{head}$ |
| m01932[c] | gamma-linolenate | C06426 | $PQ_{head}$ |
| m01696[c] | dihomo-gamma-linolenate | C03242 | $PQ_{head}$ |
| m01697[c] | dihomo-gamma-linolenoyl-CoA | C03595 | $PQ_{head}$ |
| m01291[c] | adrenic acid | C16527 | $PQ_{head}$ |
| m00119[c] | (7Z,10Z,13Z,16Z)-docosatetraenoyl-CoA | C16170 | $PQ_{head}$ |
| m00132[c] | (9Z,12Z,15Z,18Z)-TTA | | $PQ_{head}$ |
| m00111[c] | (6Z,9Z,12Z,15Z,18Z)-TPA | | $PQ_{head}$ |
| m00094[c] | (4Z,7Z,10Z,13Z,16Z)-DPA | | $PQ_{head}$ |
| m00008[c] | (11Z,14Z)-eicosadienoic acid | C16525 | $PQ_{head}$ |
| m00009[c] | (11Z,14Z)-eicosadienoyl-CoA | C16180 | $PQ_{head}$ |
| m00021[c] | (13Z,16Z)-docosadienoic acid | C16533 | $PQ_{head}$ |
| m00023[c] | (13Z,16Z)-docosadienoyl-CoA | C16645 | $PQ_{head}$ |
| m00265[c] | 10,13,16-docosatriynoic acid | | $PQ_{head}$ |
| m00264[c] | 10,13,16-docosatrienoyl-CoA | | $PQ_{head}$ |
| m00116[c] | (7Z)-octadecenoyl-CoA | | $PQ_{head}$ |
| m02394[c] | lipoic acid | C00725 | $PQ_{head}$ |
| m02398[c] | lipoyl-AMP | C16238 | $PQ_{head}$ |
| m00209[c] | [protein]-N6-(lipoyl)lysine | C16237 | $PQ_{head}$ |
| m00710[c] | 3(S)-hydroxy-dihomo-gamma-linolenoyl-CoA | | $PQ_{head}$ |
| m03029[c] | trans-2-cis,cis,cis-8,11,14-eicosatetraenoyl-CoA | | $PQ_{head}$ |
| m00107[c] | (6Z,9Z,12Z,15Z)-octadecatetraenoylcarnitine | | $PQ_{head}$ |
| m00102[c] | (5Z,8Z,11Z,14Z,17Z)-eicosapentaenoylcarnitine | | $PQ_{head}$ |
| m00107[m] | (6Z,9Z,12Z,15Z)-octadecatetraenoylcarnitine | | $PQ_{head}$ |
| m00108[m] | (6Z,9Z,12Z,15Z)-octadecatetraenoyl-CoA | C16163 | $PQ_{head}$ |
| m00102[m] | (5Z,8Z,11Z,14Z,17Z)-eicosapentaenoylcarnitine | | $PQ_{head}$ |
| m00103[m] | (5Z,8Z,11Z,14Z,17Z)-eicosapentaenoyl-CoA | C16165 | $PQ_{head}$ |

| | | | |
|---|---|---|---|
| m00240[c] | 1,2-diacylglycerol-LD-TAG pool | C00641 | $PQ_{head}$ |
| m02733[c] | phosphatidate-LD-TAG pool | C00416 | $PQ_{head}$ |
| m01277[c] | acyl-CoA-LD-TG3 pool | | $PQ_{head}$ |
| m02958[c] | TAG-LD pool | C00422 | $PQ_{head}$ |
| m01812[c] | fatty acid-LD-SM pool | C00162 | $PQ_{head}$ |
| m02187[c] | isopentenyl-pPP | C00129 | $PQ_{head}$ |
| m01316[c] | all-trans-decaprenyl-diphosphate | C17432 | $PQ_{head}$ |
| m00995[m] | 4-hydroxybenzoate | C00156 | $PQ_{head}$ |
| m01316[m] | all-trans-decaprenyl-diphosphate | C17432 | $PQ_{head}$ |
| m00767[m] | 3-decaprenyl-4-hydroxybenzoate | | $PQ_{head}$ |
| m00725[m] | 3,4-dihydroxy-5-all-trans-decaprenylbenzoate | | $PQ_{head}$ |
| m00817[m] | 3-methoxy-4-hydroxy-5-all-trans-decaprenylbenzoate | | $PQ_{head}$ |
| m00657[m] | 2-methoxy-6-(all-trans-decaprenyl)phenol | | $PQ_{head}$ |
| m00658[m] | 2-methoxy-6-all trans-decaprenyl-2-methoxy-1,4-benzoquinol | | $PQ_{head}$ |
| m01165[m] | 6-methoxy-3-methyl-2-all-trans-decaprenyl-1,4-benzoquinol | | $PQ_{head}$ |
| m00770[m] | 3-demethylubiquinol-10 | | $PQ_{head}$ |
| m01450[c] | cholesterol | C00187 | $PQ_{head}$ |
| m02131[c] | HMG-CoA | C00356 | $PQ_{head}$ |
| m00167[c] | (R)-mevalonate | C00418 | $PQ_{head}$ |
| m00165[c] | (R)-5-phosphomevalonate | C01107 | $PQ_{head}$ |
| m00164[c] | (R)-5-diphosphomevalonate | C01143 | $PQ_{head}$ |
| m01706[c] | dimethylallyl-PP | C00235 | $PQ_{head}$ |
| m01953[c] | geranyl-PP | C00341 | $PQ_{head}$ |
| m01451[r] | cholesterol-ester pool | C02530 | $PQ_{head}$ |
| m01699[c] | dihydroceramide pool | | $PQ_{head}$ |
| m01430[c] | ceramide pool | C00195 | $PQ_{head}$ |
| m02684[c] | PC-LD pool | C00157 | $PQ_{head}$ |
| m00769[c] | 3-dehydrosphinganine | C02934 | $PQ_{head}$ |
| m02927[c] | sphinganine | C00836 | $PQ_{head}$ |
| m02928[c] | sphinganine-1-phosphate | C01120 | $PQ_{head}$ |
| m01798[c] | ethanolamine-phosphate | C00346 | $PQ_{head}$ |
| m02908[r] | SM pool | C00550 | $PQ_{head}$ |
| m02684[r] | PC-LD pool | C00157 | $PQ_{head}$ |
| m00484[c] | 1-acylglycerol-3P-laur | | $PQ_{head}$ |
| m00502[c] | 1-acylglycerol-3P-tridec | | $PQ_{head}$ |
| m00493[c] | 1-acylglycerol-3P-myrist | | $PQ_{head}$ |
| m00471[c] | 1-acylglycerol-3P-9-tetrade | | $PQ_{head}$ |
| m00463[c] | 1-acylglycerol-3P-7-tetrade | | $PQ_{head}$ |
| m00454[c] | 1-acylglycerol-3P-5-tetrade | | $PQ_{head}$ |
| m00498[c] | 1-acylglycerol-3P-pentade | | $PQ_{head}$ |
| m00496[c] | 1-acylglycerol-3P-palm | | $PQ_{head}$ |
| m00497[c] | 1-acylglycerol-3P-palmn | | $PQ_{head}$ |
| m00461[c] | 1-acylglycerol-3P-7-hexade | | $PQ_{head}$ |

| | | | |
|---|---|---|---|
| m00481[c] | 1-acylglycerol-3P-heptade | | $PQ_{head}$ |
| m00438[c] | 1-acylglycerol-3P-10-heptade | | $PQ_{head}$ |
| m00469[c] | 1-acylglycerol-3P-9-heptade | | $PQ_{head}$ |
| m00499[c] | 1-acylglycerol-3P-stea | | $PQ_{head}$ |
| m00448[c] | 1-acylglycerol-3P-13-octade | | $PQ_{head}$ |
| m00474[c] | 1-acylglycerol-3P-cis-vac | | $PQ_{head}$ |
| m00495[c] | 1-acylglycerol-3P-ol | | $PQ_{head}$ |
| m00470[c] | 1-acylglycerol-3P-9-octade | | $PQ_{head}$ |
| m00462[c] | 1-acylglycerol-3P-7-octade | | $PQ_{head}$ |
| m00458[c] | 1-acylglycerol-3P-6,9-octa | | $PQ_{head}$ |
| m00494[c] | 1-acylglycerol-3P-nanode | | $PQ_{head}$ |
| m00478[c] | 1-acylglycerol-3P-eico | | $PQ_{head}$ |
| m00447[c] | 1-acylglycerol-3P-13-eicose | | $PQ_{head}$ |
| m00442[c] | 1-acylglycerol-3P-11-eico | | $PQ_{head}$ |
| m00468[c] | 1-acylglycerol-3P-9-eicose | | $PQ_{head}$ |
| m00465[c] | 1-acylglycerol-3P-8,11-eico | | $PQ_{head}$ |
| m00453[c] | 1-acylglycerol-3P-5,8,11-eico | | $PQ_{head}$ |
| m00480[c] | 1-acylglycerol-3P-heneico | | $PQ_{head}$ |
| m00477[c] | 1-acylglycerol-3P-docosa | | $PQ_{head}$ |
| m00446[c] | 1-acylglycerol-3P-13-docose | | $PQ_{head}$ |
| m00441[c] | 1-acylglycerol-3P-11-docose | | $PQ_{head}$ |
| m00501[c] | 1-acylglycerol-3P-trico | | $PQ_{head}$ |
| m00500[c] | 1-acylglycerol-3P-tetraco | | $PQ_{head}$ |
| m00449[c] | 1-acylglycerol-3P-15-tetra | | $PQ_{head}$ |
| m00482[c] | 1-acylglycerol-3P-hexacosa | | $PQ_{head}$ |
| m00483[c] | 1-acylglycerol-3P-hexecose | | $PQ_{head}$ |
| m00492[c] | 1-acylglycerol-3P-linolen | | $PQ_{head}$ |
| m00457[c] | 1-acylglycerol-3P-6,9,12,15-octa | | $PQ_{head}$ |
| m00464[c] | 1-acylglycerol-3P-8,11,14,17-eico | | $PQ_{head}$ |
| m00452[c] | 1-acylglycerol-3P-5,8,11,14,17-eico | | $PQ_{head}$ |
| m00459[c] | 1-acylglycerol-3P-7,10,13,16,19-docosa | | $PQ_{head}$ |
| m00466[c] | 1-acylglycerol-3P-9,12,15,18,21-tetra | | $PQ_{head}$ |
| m00455[c] | 1-acylglycerol-3P-6,9,12,15,18,21-tetra | | $PQ_{head}$ |
| m00450[c] | 1-acylglycerol-3P-4,7,10,13,16,19-doco | | $PQ_{head}$ |
| m00439[c] | 1-acylglycerol-3P-11,14,17-eico | | $PQ_{head}$ |
| m00444[c] | 1-acylglycerol-3P-13,16,19-doco | | $PQ_{head}$ |
| m00436[c] | 1-acylglycerol-3P-10,13,16,19-doco | | $PQ_{head}$ |
| m00443[c] | 1-acylglycerol-3P-12,15,18,21-tetra | | $PQ_{head}$ |
| m00491[c] | 1-acylglycerol-3P-lin | | $PQ_{head}$ |
| m00479[c] | 1-acylglycerol-3P-gamma-lin | | $PQ_{head}$ |
| m00476[c] | 1-acylglycerol-3P-dihomo-gamma | | $PQ_{head}$ |
| m00472[c] | 1-acylglycerol-3P-arach | | $PQ_{head}$ |
| m00460[c] | 1-acylglycerol-3P-7,10,13,16-docosa | | $PQ_{head}$ |

| | | | |
|---|---|---|---|
| m00467[c] | 1-acylglycerol-3P-9,12,15,18-tetraco | | $PQ_{head}$ |
| m00456[c] | 1-acylglycerol-3P-6,9,12,15,18-tetraco | | $PQ_{head}$ |
| m00451[c] | 1-acylglycerol-3P-4,7,10,13,16-docosa | | $PQ_{head}$ |
| m00440[c] | 1-acylglycerol-3P-11,14-eicosa | | $PQ_{head}$ |
| m00445[c] | 1-acylglycerol-3P-13,16-docosa | | $PQ_{head}$ |
| m00437[c] | 1-acylglycerol-3P-10,13,16-docosa | | $PQ_{head}$ |
| m00490[c] | 1-acylglycerol-3P-LD-TG1 pool | C00681 | $PQ_{head}$ |
| m00488[c] | 1-acylglycerol-3P-LD-PS pool | C00681 | $PQ_{head}$ |
| m02728[c] | phosphatidate-LD-PC pool | C00416 | $PQ_{head}$ |
| m01274[c] | acyl-CoA-LD-PS pool | | $PQ_{head}$ |
| m02731[c] | phosphatidate-LD-PS pool | C00416 | $PQ_{head}$ |
| m02171[c] | inositol | C00137 | $PQ_{head}$ |
| m02750[c] | PI pool | C01194 | $PQ_{head}$ |
| m02685[c] | PE-LD pool | C00350 | $PQ_{head}$ |
| m02808[c] | PS-LD pool | C02737 | $PQ_{head}$ |
| m00656[c] | 2-lysolecithin pool | C04230 | $PQ_{head}$ |
| m02600[c] | N-methylethanolamine-phosphate | C01210 | $PQ_{head}$ |
| m02739[c] | phosphodimethylethanolamine | C13482 | $PQ_{head}$ |
| m01513[r] | choline | C00114 | $PQ_{head}$ |
| m02728[r] | phosphatidate-LD-PC pool | C00416 | $PQ_{head}$ |
| m02786[c] | prostaglandin E2 | C00584 | $PQ_{head}$ |
| m01627[c] | cysteamine | C01678 | $PQ_{head}$ |
| m02519[s] | Na+ | C01330 | $PQ_{head}$ |
| m02519[c] | Na+ | C01330 | $PQ_{head}$ |
| m01349[c] | apo-[ACP] | C03688 | $PQ_{head}$ |
| m02741[c] | phosphopantetheine | C01134 | $PQ_{head}$ |
| m02679[c] | pantetheine | C00831 | $PQ_{head}$ |
| m02680[c] | pantothenate | C00864 | $PQ_{head}$ |
| m01044[c] | 5,10-methenyl-THF | C00445 | $PQ_{head}$ |
| m00267[c] | 10-formyl-THF-glu(5) | | $PQ_{head}$ |
| m00267[l] | 10-formyl-THF-glu(5) | | $PQ_{head}$ |
| m00266[l] | 10-formyl-THF | C00234 | $PQ_{head}$ |
| m01974[l] | glutamate | C00025 | $PQ_{head}$ |
| m02980[s] | THF | C00101 | $PQ_{head}$ |
| m01401[c] | biotin | C00120 | $PQ_{head}$ |
| m01401[s] | biotin | C00120 | $PQ_{head}$ |
| m02426[s] | lysine | C00047 | $PQ_{head}$ |
| m01155[c] | 6-[(1S,2R)-1,2-dihydroxy-3-triphosphooxypropyl]-7,8-dihydropterin | C04895 | $PQ_{head}$ |
| m03057[c] | triphosphate | C00536 | $PQ_{head}$ |
| m01704[c] | dihydroneopterin | C04874 | $PQ_{head}$ |
| m02049[m] | heme | C00032 | $PQ_{head}$ |
| m01631[m] | cytochrome-C | C00524 | $PQ_{head}$ |
| m02049[c] | heme | C00032 | $PQ_{head}$ |

| | | | |
|---|---|---|---|
| m01385[c] | beta-carotene | C02094 | $PQ_{head}$ |
| m03057[m] | triphosphate | C00536 | $PQ_{head}$ |
| m01600[m] | cobamide-coenzyme | C00194 | $PQ_{head}$ |
| m01442[c] | chloride | C00698 | $PQ_{head}$ |
| m02360[s] | leucine | C00123 | $PQ_{head}$ |
| m02834[s] | retinol | C00473 | $PQ_{head}$ |
| m02392[c] | lipid droplet | | $PQ_{head}$ |
| m02685[r] | PE-LD pool | C00350 | $PQ_{head}$ |
| m02808[r] | PS-LD pool | C02737 | $PQ_{head}$ |
| m02958[r] | TAG-LD pool | C00422 | $PQ_{head}$ |
| m02958[l] | TAG-LD pool | C00422 | $PQ_{head}$ |
| m01450[s] | cholesterol | C00187 | $PQ_{head}$ |
| m01115[s] | 5-methyl-THF | C00440 | $PQ_{head}$ |
| m02680[s] | pantothenate | C00864 | $PQ_{head}$ |
| m02171[s] | inositol | C00137 | $PQ_{head}$ |
| m01383[s] | beta-alanine | C00099 | $PQ_{head}$ |
| m01442[s] | chloride | C00698 | $PQ_{head}$ |
| m01253[s] | acetoacetate | C00164 | $PQ_{head}$ |
| m00157[s] | (R)-3-hydroxybutanoate | C01089 | $PQ_{head}$ |
| m00157[c] | (R)-3-hydroxybutanoate | C01089 | $PQ_{head}$ |
| m01633[s] | D-3-amino-isobutanoate | C01205 | $PQ_{head}$ |
| m02125[s] | histidine | C00135 | $PQ_{head}$ |
| m03089[s] | tryptophan | C00078 | $PQ_{head}$ |
| m02724[s] | phenylalanine | C00079 | $PQ_{head}$ |
| m01369[s] | asparagine | C00152 | $PQ_{head}$ |
| m03135[s] | valine | C00183 | $PQ_{head}$ |
| m02184[s] | isoleucine | C00407 | $PQ_{head}$ |
| m02961[s] | taurine | C00245 | $PQ_{head}$ |
| m02772[s] | propanoate | C00163 | $PQ_{head}$ |
| m02786[s] | prostaglandin E2 | C00584 | $PQ_{head}$ |
| m02394[s] | lipoic acid | C00725 | $PQ_{head}$ |
| m02147[s] | hydroxide | C01328 | $PQ_{head}$ |
| m02049[s] | heme | C00032 | $PQ_{head}$ |
| m02928[s] | sphinganine-1-phosphate | C01120 | $PQ_{head}$ |
| m01361[s] | aquacob(III)alamin | C00992 | $PQ_{head}$ |
| m02348[s] | L-carnitine | C15025 | $PQ_{head}$ |
| m01385[s] | beta-carotene | C02094 | $PQ_{head}$ |
| m03161[s] | glycogen | C00182 | $PQ_{head}$ |
| m01682[s] | D-glucitol | C00794 | $PQ_{head}$ |
| m01285[l] | ADP | C00008 | $PQ_{head}$ |
| m01371[l] | ATP | C00002 | $PQ_{head}$ |
| m01365[l] | arginine | C00062 | $PQ_{head}$ |
| m01369[l] | asparagine | C00152 | $PQ_{head}$ |

| | | | |
|---|---|---|---|
| m01628[l] | cysteine | C00097 | $PQ_{head}$ |
| m01975[l] | glutamine | C00064 | $PQ_{head}$ |
| m02125[l] | histidine | C00135 | $PQ_{head}$ |
| m02360[l] | leucine | C00123 | $PQ_{head}$ |
| m02426[l] | lysine | C00047 | $PQ_{head}$ |
| m02471[l] | methionine | C00073 | $PQ_{head}$ |
| m02724[l] | phenylalanine | C00079 | $PQ_{head}$ |
| m02770[l] | proline | C00148 | $PQ_{head}$ |
| m02896[l] | serine | C00065 | $PQ_{head}$ |
| m03089[l] | tryptophan | C00078 | $PQ_{head}$ |
| m03101[l] | tyrosine | C00082 | $PQ_{head}$ |
| m03135[l] | valine | C00183 | $PQ_{head}$ |
| m01307[l] | alanine | C00041 | $PQ_{head}$ |
| m02184[l] | isoleucine | C00407 | $PQ_{head}$ |
| m02993[l] | threonine | C00188 | $PQ_{head}$ |
| m03139[c] | vitamin A derivatives | | $PQ_{head}$ |
| m03143[c] | vitamin E derivatives | | $PQ_{head}$ |
| m01602[c] | cofactors and vitamins | | $PQ_{head}$ |
| m02040[c] | H2O | C00001 | $PQ_{head}$ |
| m01998[c] | glycolate | C00160 | $PQ_{head}$ |
| m02877[c] | SAM | C00019 | $PQ_{head}$ |
| m02871[c] | SAH | C00021 | $PQ_{head}$ |
| m02039[c] | H+ | C00080 | $PQ_{head}$ |
| m02877[m] | SAM | C00019 | $PQ_{head}$ |
| m02871[m] | SAH | C00021 | $PQ_{head}$ |
| m02039[m] | H+ | C00080 | $PQ_{head}$ |
| m02877[n] | SAM | C00019 | $PQ_{head}$ |
| m02871[n] | SAH | C00021 | $PQ_{head}$ |
| m02039[n] | H+ | C00080 | $PQ_{head}$ |
| m01371[c] | ATP | C00002 | $PQ_{head}$ |
| m01285[c] | ADP | C00008 | $PQ_{head}$ |
| m01626[c] | cys-gly | C01419 | $PQ_{head}$ |
| m01628[c] | cysteine | C00097 | $PQ_{head}$ |
| m01986[c] | glycine | C00037 | $PQ_{head}$ |
| m01626[s] | cys-gly | C01419 | $PQ_{head}$ |
| m01986[s] | glycine | C00037 | $PQ_{head}$ |
| m02040[m] | H2O | C00001 | $PQ_{head}$ |
| m02914[c] | sn-glycerol-3-phosphate | C00093 | $PQ_{head}$ |
| m01371[m] | ATP | C00002 | $PQ_{head}$ |
| m01285[m] | ADP | C00008 | $PQ_{head}$ |
| m03106[c] | UDP | C00015 | $PQ_{head}$ |
| m02039[r] | H+ | C00080 | $PQ_{head}$ |
| m02039[s] | H+ | C00080 | $PQ_{head}$ |

| | | | |
|---|---|---|---|
| m02041[c] | H2O2 | C00027 | $PQ_{head}$ |
| m01597[c] | CoA | C00010 | $PQ_{head}$ |
| m01597[m] | CoA | C00010 | $PQ_{head}$ |
| m02630[r] | O2 | C00007 | $PQ_{head}$ |
| km00013[r] | Red-NADPH-Hemoprotein-Reductases | | $PQ_{head}$ |
| km00014[r] | Ox-NADPH-Hemoprotein-Reductases | | $PQ_{head}$ |
| m02040[r] | H2O | C00001 | $PQ_{head}$ |
| m02555[c] | NADPH | C00005 | $PQ_{head}$ |
| m02554[c] | NADP+ | C00006 | $PQ_{head}$ |
| m02555[m] | NADPH | C00005 | $PQ_{head}$ |
| m02554[m] | NADP+ | C00006 | $PQ_{head}$ |
| m01974[c] | glutamate | C00025 | $PQ_{head}$ |
| m01306[c] | AKG | C00026 | $PQ_{head}$ |
| m01974[m] | glutamate | C00025 | $PQ_{head}$ |
| m01306[m] | AKG | C00026 | $PQ_{head}$ |
| m02630[c] | O2 | C00007 | $PQ_{head}$ |
| m02555[r] | NADPH | C00005 | $PQ_{head}$ |
| m02554[r] | NADP+ | C00006 | $PQ_{head}$ |
| m01280[c] | adenosine | C00212 | $PQ_{head}$ |
| m02133[c] | homocysteine | C00155 | $PQ_{head}$ |
| m02552[s] | NAD+ | C00003 | $PQ_{head}$ |
| m01362[c] | arachidonate | C00219 | $PQ_{head}$ |
| m02547[c] | N-acetylputrescine | C02714 | $PQ_{head}$ |
| m01252[c] | acetate | C00033 | $PQ_{head}$ |
| m02812[c] | putrescine | C00134 | $PQ_{head}$ |
| m02630[m] | O2 | C00007 | $PQ_{head}$ |
| km00046[m] | Reduced-adrenal-ferredoxins | | $PQ_{head}$ |
| km00048[m] | Oxidized-adrenal-ferredoxins | C00667 | $PQ_{head}$ |
| m02819[m] | pyruvate | C00022 | $PQ_{head}$ |
| m01334[c] | AMP | C00020 | $PQ_{head}$ |
| m02759[c] | PPi | C00013 | $PQ_{head}$ |
| m02774[c] | propanoyl-CoA | C00100 | $PQ_{head}$ |
| m02552[c] | NAD+ | C00003 | $PQ_{head}$ |
| m02553[c] | NADH | C00004 | $PQ_{head}$ |
| m02552[p] | NAD+ | C00003 | $PQ_{head}$ |
| m02553[p] | NADH | C00004 | $PQ_{head}$ |
| m02039[p] | H+ | C00080 | $PQ_{head}$ |
| m02552[m] | NAD+ | C00003 | $PQ_{head}$ |
| m02553[m] | NADH | C00004 | $PQ_{head}$ |
| m02444[c] | malonyl-CoA | C00083 | $PQ_{head}$ |
| m01596[c] | CO2 | C00011 | $PQ_{head}$ |
| m02040[l] | H2O | C00001 | $PQ_{head}$ |
| m01753[c] | dTTP | C00459 | $PQ_{head}$ |

| | | | |
|---|---|---|---|
| m01752[c] | dTMP | C00364 | $PQ_{head}$ |
| m02759[n] | PPi | C00013 | $PQ_{head}$ |
| m03148[c] | xanthine | C00385 | $PQ_{head}$ |
| m02630[p] | O2 | C00007 | $PQ_{head}$ |
| m02040[p] | H2O | C00001 | $PQ_{head}$ |
| m02041[p] | H2O2 | C00027 | $PQ_{head}$ |
| m00981[c] | 4-coumarate | C00811 | $PQ_{head}$ |
| m00982[c] | 4-coumaroyl-CoA | C00223 | $PQ_{head}$ |
| m01334[m] | AMP | C00020 | $PQ_{head}$ |
| m02759[m] | PPi | C00013 | $PQ_{head}$ |
| m01261[c] | acetyl-CoA | C00024 | $PQ_{head}$ |
| m02633[c] | OAA | C00036 | $PQ_{head}$ |
| m02696[c] | PEP | C00074 | $PQ_{head}$ |
| m01596[m] | CO2 | C00011 | $PQ_{head}$ |
| m03101[c] | tyrosine | C00082 | $PQ_{head}$ |
| m01690[c] | DHAP | C00111 | $PQ_{head}$ |
| m01170[c] | 6-pyruvoyltetrahydropterin | C03684 | $PQ_{head}$ |
| m02026[c] | GSH | C00051 | $PQ_{head}$ |
| m00639[c] | 2-deoxy-D-ribose-1-phosphate | C00672 | $PQ_{head}$ |
| m01590[c] | CMP | C00055 | $PQ_{head}$ |
| m02647[c] | oleoyl-CoA | C00510 | $PQ_{head}$ |
| m00918[c] | 3-sulfinoalanine | C00606 | $PQ_{head}$ |
| m02157[c] | hypotaurine | C00519 | $PQ_{head}$ |
| m00131[c] | (9Z,12Z,15Z,18Z)-tetracosatetraenoyl-CoA | C16171 | $PQ_{head}$ |
| m01669[s] | deoxyguanosine | C00330 | $PQ_{head}$ |
| m01828[c] | FMN | C00061 | $PQ_{head}$ |
| m01802[c] | FAD | C00016 | $PQ_{head}$ |
| m01802[m] | FAD | C00016 | $PQ_{head}$ |
| m01668[s] | deoxycytidine | C00881 | $PQ_{head}$ |
| m01285[s] | ADP | C00008 | $PQ_{head}$ |
| m02471[c] | methionine | C00073 | $PQ_{head}$ |
| m02772[c] | propanoate | C00163 | $PQ_{head}$ |
| m02774[m] | propanoyl-CoA | C00100 | $PQ_{head}$ |
| m02007[c] | glyoxalate | C00048 | $PQ_{head}$ |
| m02961[c] | taurine | C00245 | $PQ_{head}$ |
| m02553[r] | NADH | C00004 | $PQ_{head}$ |
| m02552[r] | NAD+ | C00003 | $PQ_{head}$ |
| m01927[c] | gamma-glutamyl-cysteine | C00669 | $PQ_{head}$ |
| m01968[c] | glucose-6-phosphate | C00092 | $PQ_{head}$ |
| m02042[c] | H2S | C00283 | $PQ_{head}$ |
| m01587[c] | citrate | C00158 | $PQ_{head}$ |
| m02946[c] | sulfate | C00059 | $PQ_{head}$ |

| | | | |
|---|---|---|---|
| m01283[c] | adenylyl sulfate | C00224 | $PQ_{head}$ |
| m02391[c] | linoleoyl-CoA | C02050 | $PQ_{head}$ |
| m02896[s] | serine | C00065 | $PQ_{head}$ |
| m03089[c] | tryptophan | C00078 | $PQ_{head}$ |
| m01669[c] | deoxyguanosine | C00330 | $PQ_{head}$ |
| m02037[c] | guanine | C00242 | $PQ_{head}$ |
| m03120[c] | urate | C00366 | $PQ_{head}$ |
| m03120[p] | urate | C00366 | $PQ_{head}$ |
| m02770[c] | proline | C00148 | $PQ_{head}$ |
| m00559[m] | 1-pyrroline-5-carboxylate | C03912 | $PQ_{head}$ |
| m02770[m] | proline | C00148 | $PQ_{head}$ |
| m02819[c] | pyruvate | C00022 | $PQ_{head}$ |
| m02439[m] | malate | C00711 | $PQ_{head}$ |
| m02941[c] | stearoyl-CoA | C00412 | $PQ_{head}$ |
| m01370[c] | aspartate | C00049 | $PQ_{head}$ |
| m01370[l] | aspartate | C00049 | $PQ_{head}$ |
| m01934[c] | gamma-linolenoyl-CoA | C03035 | $PQ_{head}$ |
| m00859[c] | 3-oxo-dihomo-gamma-linolenoyl-CoA | | $PQ_{head}$ |
| m01806[c] | farnesyl-PP | C00448 | $PQ_{head}$ |
| m01975[c] | glutamine | C00064 | $PQ_{head}$ |
| m01975[m] | glutamine | C00064 | $PQ_{head}$ |
| m02159[c] | hypoxanthine | C00262 | $PQ_{head}$ |
| m02678[c] | palmitoyl-CoA | C00154 | $PQ_{head}$ |
| m01686[c] | dGMP | C00362 | $PQ_{head}$ |
| m01680[c] | dGDP | C00361 | $PQ_{head}$ |
| m01686[m] | dGMP | C00362 | $PQ_{head}$ |
| m01680[m] | dGDP | C00361 | $PQ_{head}$ |
| m01307[c] | alanine | C00041 | $PQ_{head}$ |
| m01668[c] | deoxycytidine | C00881 | $PQ_{head}$ |
| m01673[c] | deoxyuridine | C00526 | $PQ_{head}$ |
| m02834[c] | retinol | C00473 | $PQ_{head}$ |
| m02832[c] | retinal | C00376 | $PQ_{head}$ |
| m02834[r] | retinol | C00473 | $PQ_{head}$ |
| m02832[r] | retinal | C00376 | $PQ_{head}$ |
| m01249[c] | acetaldehyde | C00084 | $PQ_{head}$ |
| m02403[c] | L-lactate | C00186 | $PQ_{head}$ |
| m00110[c] | (6Z,9Z,12Z,15Z,18Z)-tetracosapentaenoyl-CoA | C16172 | $PQ_{head}$ |
| m02949[c] | sulfite | C00094 | $PQ_{head}$ |
| m02110[c] | hexacosanoyl-CoA | | $PQ_{head}$ |
| m02016[c] | GMP | C00144 | $PQ_{head}$ |
| m01365[c] | arginine | C00062 | $PQ_{head}$ |
| m02426[c] | lysine | C00047 | $PQ_{head}$ |

| | | | |
|---|---|---|---|
| m02896[c] | serine | C00065 | $PQ_{head}$ |
| m02495[c] | myristoyl-CoA | C02593 | $PQ_{head}$ |
| m01365[s] | arginine | C00062 | $PQ_{head}$ |
| m01671[c] | deoxyinosine | C05512 | $PQ_{head}$ |
| m01327[r] | alpha-tocopherol | C02477 | $PQ_{head}$ |
| m00356[r] | 13-hydroxy-alpha-tocopherol | | $PQ_{head}$ |
| m02666[c] | oxidized thioredoxin | C00343 | $PQ_{head}$ |
| m00184[c] | [ACP] | C00229 | $PQ_{head}$ |
| m01260[c] | acetylcholine | C01996 | $PQ_{head}$ |
| m01513[c] | choline | C00114 | $PQ_{head}$ |
| m01260[s] | acetylcholine | C01996 | $PQ_{head}$ |
| m01261[m] | acetyl-CoA | C00024 | $PQ_{head}$ |
| m02847[c] | RNA | C00046 | $PQ_{head}$ |
| m01666[s] | deoxyadenosine | C00559 | $PQ_{head}$ |
| m02682[c] | PAPS | C00053 | $PQ_{head}$ |
| m02681[c] | PAP | C00054 | $PQ_{head}$ |
| m02046[s] | HCO3- | C00288 | $PQ_{head}$ |
| m02842[s] | riboflavin | C00255 | $PQ_{head}$ |
| m01736[s] | dopamine | C03758 | $PQ_{head}$ |
| m02677[c] | palmitoleoyl-CoA | C21072 | $PQ_{head}$ |
| m01862[c] | fumarate | C00122 | $PQ_{head}$ |
| m02724[c] | phenylalanine | C00079 | $PQ_{head}$ |
| m00113[c] | (6Z,9Z,12Z,15Z,18Z,21Z)-tetracosahexaenoyl-CoA | C16168 | $PQ_{head}$ |
| m01127[c] | 5-oxoproline | C01879 | $PQ_{head}$ |
| m03102[m] | ubiquinol | C00390 | $PQ_{head}$ |
| m02348[c] | L-carnitine | C15025 | $PQ_{head}$ |
| m02348[m] | L-carnitine | C15025 | $PQ_{head}$ |
| m01364[c] | arachidonyl-CoA | C02249 | $PQ_{head}$ |
| m02038[c] | guanosine | C00387 | $PQ_{head}$ |
| m02993[s] | threonine | C00188 | $PQ_{head}$ |
| m03118[c] | uracil | C00106 | $PQ_{head}$ |
| m02390[c] | linolenoyl-CoA | C16162 | $PQ_{head}$ |
| m02993[c] | threonine | C00188 | $PQ_{head}$ |
| m00025[c] | (15Z)-tetracosenoyl-CoA | C16532 | $PQ_{head}$ |
| m02842[c] | riboflavin | C00255 | $PQ_{head}$ |
| m02908[c] | SM pool | C00550 | $PQ_{head}$ |
| m02738[c] | phosphocholine | C00588 | $PQ_{head}$ |
| m02173[c] | inositol-1-phosphate | C01177 | $PQ_{head}$ |
| m03130[c] | UTP | C00075 | $PQ_{head}$ |
| m03114[c] | UMP | C00105 | $PQ_{head}$ |
| m01986[l] | glycine | C00037 | $PQ_{head}$ |
| m02026[s] | GSH | C00051 | $PQ_{head}$ |

| | | | |
|---|---|---|---|
| m02007[p] | glyoxalate | C00048 | $PQ_{head}$ |
| m01637[c] | dADP | C00206 | $PQ_{head}$ |
| m01642[c] | dATP | C00131 | $PQ_{head}$ |
| m02770[s] | proline | C00148 | $PQ_{head}$ |
| m02674[c] | palmitate | C00249 | $PQ_{head}$ |
| m02996[c] | thymidine | C00214 | $PQ_{head}$ |
| m02996[s] | thymidine | C00214 | $PQ_{head}$ |
| m02159[p] | hypoxanthine | C00262 | $PQ_{head}$ |
| m01688[c] | dGTP | C00286 | $PQ_{head}$ |
| m01755[c] | dUMP | C00365 | $PQ_{head}$ |
| m01666[c] | deoxyadenosine | C00559 | $PQ_{head}$ |
| m02345[c] | lauroyl-CoA | C01832 | $PQ_{head}$ |
| m01307[s] | alanine | C00041 | $PQ_{head}$ |
| m02494[c] | myristic acid | C06424 | $PQ_{head}$ |
| m02578[c] | NH3 | C00014 | $PQ_{head}$ |
| m00606[m] | 22beta-hydroxycholesterol | C05502 | $PQ_{head}$ |
| m00579[m] | 20alpha,22beta dihydroxycholesterol | C05501 | $PQ_{head}$ |
| m02182[m] | isocaproic-aldehyde | C02373 | $PQ_{head}$ |
| m02763[m] | pregnenolone | C01953 | $PQ_{head}$ |
| m03135[c] | valine | C00183 | $PQ_{head}$ |
| m01450[r] | cholesterol | C00187 | $PQ_{head}$ |
| m02751[c] | Pi | C00009 | $PQ_{head}$ |
| m03101[s] | tyrosine | C00082 | $PQ_{head}$ |
| m01803[c] | FADH2 | C01352 | $PQ_{head}$ |
| m01803[m] | FADH2 | C01352 | $PQ_{head}$ |
| m02980[c] | THF | C00101 | $PQ_{head}$ |
| m02751[m] | Pi | C00009 | $PQ_{head}$ |
| m02751[s] | Pi | C00009 | $PQ_{head}$ |
| m00671[c] | 2-oxobutyrate | C00109 | $PQ_{head}$ |
| m01682[c] | D-glucitol | C00794 | $PQ_{head}$ |
| m01790[r] | estrone | C00468 | $PQ_{head}$ |
| m02578[p] | NH3 | C00014 | $PQ_{head}$ |
| m02475[c] | methylglyoxal | C00546 | $PQ_{head}$ |
| m02833[c] | retinoate | C00777 | $PQ_{head}$ |
| m02833[r] | retinoate | C00777 | $PQ_{head}$ |
| m01787[r] | estradiol-17beta | C00951 | $PQ_{head}$ |
| m02990[c] | thioredoxin | C00342 | $PQ_{head}$ |
| m02442[c] | malonyl-[ACP] | C01209 | $PQ_{head}$ |
| m01704[s] | dihydroneopterin | C04874 | $PQ_{head}$ |
| m00095[c] | (4Z,7Z,10Z,13Z,16Z,19Z)-docosahexaenoyl-CoA | C16169 | $PQ_{head}$ |
| m00093[c] | (4Z,7Z,10Z,13Z,16Z)-docosapentaenoyl-CoA | C16173 | $PQ_{head}$ |
| m01232[c] | 9-cis-retinol | C16682 | $PQ_{head}$ |

| | | | |
|---|---|---|---|
| m02978[c] | tetrahydrobiopterin | C00272 | $PQ_{head}$ |
| m01361[c] | aquacob(III)alamin | C00992 | $PQ_{head}$ |
| m01599[c] | cob(II)alamin | C00541 | $PQ_{head}$ |
| m01599[m] | cob(II)alamin | C00541 | $PQ_{head}$ |
| km00877[r] | [Reduced NADPH---hemoprotein reductase] | C03024 | $PQ_{head}$ |
| km00878[r] | [Oxidized NADPH---hemoprotein reductase] | C03161 | $PQ_{head}$ |
| m00427[r] | 18-hydroxy-all-trans-retinoate | C16679 | $PQ_{head}$ |
| m01093[m] | 5beta-cholestane-3alpha,7alpha,26-triol | C05444 | $PQ_{head}$ |
| m00756[m] | 3alpha,7alpha-dihydroxy-5beta-cholestan-26-al | C05445 | $PQ_{head}$ |
| m01722[n] | DNA-5-methylcytosine | C02967 | $PQ_{head}$ |
| m01598[m] | cob(I)alamin | C00853 | $PQ_{head}$ |
| m01358[c] | apocytochrome-C | C02248 | $PQ_{head}$ |
| m01358[l] | apocytochrome-C | C02248 | $PQ_{head}$ |
| m01358[m] | apocytochrome-C | C02248 | $PQ_{head}$ |
| m02034[c] | GTP | C00044 | $PQ_{head}$ |
| m01948[c] | GDP | C00035 | $PQ_{head}$ |
| m02034[m] | GTP | C00044 | $PQ_{head}$ |
| m01948[m] | GDP | C00035 | $PQ_{head}$ |
| m02633[m] | OAA | C00036 | $PQ_{head}$ |
| m02696[m] | PEP | C00074 | $PQ_{head}$ |
| m01948[s] | GDP | C00035 | $PQ_{head}$ |
| m02016[s] | GMP | C00144 | $PQ_{head}$ |
| m02040[s] | H2O | C00001 | $PQ_{head}$ |
| m03114[s] | UMP | C00105 | $PQ_{head}$ |
| m03106[s] | UDP | C00015 | $PQ_{head}$ |
| m01752[m] | dTMP | C00364 | $PQ_{head}$ |
| m01630[c] | cytidine | C00475 | $PQ_{head}$ |
| m02161[s] | IDP | C00104 | $PQ_{head}$ |
| m02167[s] | IMP | C00130 | $PQ_{head}$ |
| m01371[s] | ATP | C00002 | $PQ_{head}$ |
| m01334[s] | AMP | C00020 | $PQ_{head}$ |
| m01641[p] | D-aspartate | C00402 | $PQ_{head}$ |
| m02633[p] | OAA | C00036 | $PQ_{head}$ |
| m01986[m] | glycine | C00037 | $PQ_{head}$ |
| m02321[m] | L-2-amino-3-oxobutanoic acid | C03508 | $PQ_{head}$ |
| m01986[p] | glycine | C00037 | $PQ_{head}$ |
| m02393[m] | lipoamide | C00248 | $PQ_{head}$ |
| m02878[m] | S-aminomethyldihydrolipoamide | | $PQ_{head}$ |
| m01701[m] | dihydrolipoamide | C00579 | $PQ_{head}$ |
| m01045[m] | 5,10-methylene-THF | C00143 | $PQ_{head}$ |
| m02980[m] | THF | C00101 | $PQ_{head}$ |
| m02400[m] | lipoylprotein | C02051 | $PQ_{head}$ |

| | | | |
|---|---|---|---|
| m02879[m] | S-aminomethyldihydrolipoylprotein | C01242 | $PQ_{head}$ |
| m01703[m] | dihydrolipoylprotein | C02972 | $PQ_{head}$ |
| m00669[c] | 2-oxo-3-methylvalerate | C03465 | $PQ_{head}$ |
| m02184[m] | isoleucine | C00407 | $PQ_{head}$ |
| m00669[m] | 2-oxo-3-methylvalerate | C03465 | $PQ_{head}$ |
| m00815[c] | 3-mercaptolactate | C05823 | $PQ_{head}$ |
| m02464[c] | mercaptopyruvate | C00957 | $PQ_{head}$ |
| m00816[c] | 3-mercaptolactate-cysteine-disulfide | | $PQ_{head}$ |
| m01638[p] | D-alanine | C00133 | $PQ_{head}$ |
| m01141[c] | 5-tetradecenoyl-CoA | | $PQ_{head}$ |
| m01450[m] | cholesterol | C00187 | $PQ_{head}$ |
| m02453[s] | mannose | C00159 | $PQ_{head}$ |
| m02453[c] | mannose | C00159 | $PQ_{head}$ |
| m02998[c] | thyroxine | C01829 | $PQ_{head}$ |
| m02998[s] | thyroxine | C01829 | $PQ_{head}$ |
| m02900[c] | S-glutathionyl-2-4-dinitrobenzene | | $PQ_{head}$ |
| m02900[s] | S-glutathionyl-2-4-dinitrobenzene | | $PQ_{head}$ |
| m03052[s] | triiodothyronine | C02465 | $PQ_{head}$ |
| m03052[c] | triiodothyronine | C02465 | $PQ_{head}$ |
| m00816[s] | 3-mercaptolactate-cysteine-disulfide | | $PQ_{head}$ |
| m02993[m] | threonine | C00188 | $PQ_{head}$ |
| m01641[c] | D-aspartate | C00402 | $PQ_{head}$ |
| m01638[c] | D-alanine | C00133 | $PQ_{head}$ |
| m02736[c] | phosphatidylinositol-4,5-bisphosphate | C04637 | $PQ_{head}$ |
| m02736[n] | phosphatidylinositol-4,5-bisphosphate | C04637 | $PQ_{head}$ |
| m00970[s] | 4-aminobutyrate | C00334 | $PQ_{head}$ |
| m02579[c] | NH4+ | C00014 | $PQ_{head}$ |
| m02579[m] | NH4+ | C00014 | $PQ_{head}$ |
| km00377[m] | pseudouridine 5'-phosphate | C01168 | $PQ_{head}$ |
| km00626[m] | pseudouridine | C02067 | $PQ_{head}$ |
| m02827[m] | reduced adrenal ferredoxin | C00662 | $PQ_{head}$ |
| m00592[m] | 20-hydroxycholesterol | C05500 | $PQ_{head}$ |
| m02663[m] | oxidized adrenal ferredoxin | C00667 | $PQ_{head}$ |
| m02148[c] | hydroxyacetone | C05235 | $PQ_{body}$ |
| m02046[c] | HCO3- | C00288 | $PQ_{body}$ |
| m02997[c] | thymine | C00178 | $PQ_{body}$ |
| m01705[c] | dihydrothymine | C00906 | $PQ_{body}$ |
| m00922[c] | 3-ureidoisobutyrate | C05100 | $PQ_{body}$ |
| m01633[c] | D-3-amino-isobutanoate | C01205 | $PQ_{body}$ |
| m02039[i] | H+ | C00080 | $PQ_{body}$ |
| m00101[c] | (5Z,8Z,11Z)-eicosatrienoyl-CoA | C21939 | $PQ_{body}$ |
| m00103[c] | (5Z,8Z,11Z,14Z,17Z)-eicosapentaenoyl-CoA | C16165 | $PQ_{body}$ |

| | | | |
|---|---|---|---|
| m00121[c] | (7Z,10Z,13Z,16Z,19Z)-docosapentaenoyl-CoA | C16166 | $PQ_{body}$ |
| m01697[c] | dihomo-gamma-linolenoyl-CoA | C03595 | $PQ_{body}$ |
| m01664[p] | delta2-THA-CoA | | $PQ_{body}$ |
| m00801[p] | 3-hydroxy-THA-CoA | C16375 | $PQ_{body}$ |
| m00814[p] | 3-keto-THA-CoA | | $PQ_{body}$ |
| m00710[c] | 3(S)-hydroxy-dihomo-gamma-linolenoyl-CoA | | $PQ_{body}$ |
| m03029[c] | trans-2-cis,cis,cis-8,11,14-eicosatetraenoyl-CoA | | $PQ_{body}$ |
| m02794[c] | prostaglandin H2 | C00427 | $PQ_{body}$ |
| m00100[c] | (5Z,8Z,11Z)-eicosatrienoylcarnitine | | $PQ_{body}$ |
| m00102[c] | (5Z,8Z,11Z,14Z,17Z)-eicosapentaenoylcarnitine | | $PQ_{body}$ |
| m00100[m] | (5Z,8Z,11Z)-eicosatrienoylcarnitine | | $PQ_{body}$ |
| m00101[m] | (5Z,8Z,11Z)-eicosatrienoyl-CoA | C21939 | $PQ_{body}$ |
| m00102[m] | (5Z,8Z,11Z,14Z,17Z)-eicosapentaenoylcarnitine | | $PQ_{body}$ |
| m00103[m] | (5Z,8Z,11Z,14Z,17Z)-eicosapentaenoyl-CoA | C16165 | $PQ_{body}$ |
| m02783[c] | prostaglandin D2 | C00696 | $PQ_{body}$ |
| m02997[s] | thymine | C00178 | $PQ_{body}$ |
| m01633[s] | D-3-amino-isobutanoate | C01205 | $PQ_{body}$ |
| m02783[s] | prostaglandin D2 | C00696 | $PQ_{body}$ |
| m02040[c] | H2O | C00001 | $PQ_{body}$ |
| m02039[c] | H+ | C00080 | $PQ_{body}$ |
| m02039[m] | H+ | C00080 | $PQ_{body}$ |
| m01371[c] | ATP | C00002 | $PQ_{body}$ |
| m01285[c] | ADP | C00008 | $PQ_{body}$ |
| m01626[c] | cys-gly | C01419 | $PQ_{body}$ |
| m01628[c] | cysteine | C00097 | $PQ_{body}$ |
| m01986[c] | glycine | C00037 | $PQ_{body}$ |
| m02040[m] | H2O | C00001 | $PQ_{body}$ |
| m02914[c] | sn-glycerol-3-phosphate | C00093 | $PQ_{body}$ |
| m01371[m] | ATP | C00002 | $PQ_{body}$ |
| m01285[m] | ADP | C00008 | $PQ_{body}$ |
| m02039[r] | H+ | C00080 | $PQ_{body}$ |
| m02041[c] | H2O2 | C00027 | $PQ_{body}$ |
| m01597[c] | CoA | C00010 | $PQ_{body}$ |
| m01597[m] | CoA | C00010 | $PQ_{body}$ |
| m02630[r] | O2 | C00007 | $PQ_{body}$ |
| km00013[r] | Red-NADPH-Hemoprotein-Reductases | | $PQ_{body}$ |
| km00014[r] | Ox-NADPH-Hemoprotein-Reductases | | $PQ_{body}$ |
| m02040[r] | H2O | C00001 | $PQ_{body}$ |
| m02555[c] | NADPH | C00005 | $PQ_{body}$ |
| m02554[c] | NADP+ | C00006 | $PQ_{body}$ |
| m02555[m] | NADPH | C00005 | $PQ_{body}$ |
| m02554[m] | NADP+ | C00006 | $PQ_{body}$ |

| | | | |
|---|---|---|---|
| m02630[c] | O2 | C00007 | $PQ_{body}$ |
| m02555[r] | NADPH | C00005 | $PQ_{body}$ |
| m02554[r] | NADP+ | C00006 | $PQ_{body}$ |
| m01362[c] | arachidonate | C00219 | $PQ_{body}$ |
| km00013[c] | Red-NADPH-Hemoprotein-Reductases | | $PQ_{body}$ |
| km00014[c] | Ox-NADPH-Hemoprotein-Reductases | | $PQ_{body}$ |
| m02630[m] | O2 | C00007 | $PQ_{body}$ |
| km00046[m] | Reduced-adrenal-ferredoxins | | $PQ_{body}$ |
| km00048[m] | Oxidized-adrenal-ferredoxins | C00667 | $PQ_{body}$ |
| m01334[c] | AMP | C00020 | $PQ_{body}$ |
| m02759[c] | PPi | C00013 | $PQ_{body}$ |
| m01597[p] | CoA | C00010 | $PQ_{body}$ |
| m02552[c] | NAD+ | C00003 | $PQ_{body}$ |
| m02553[c] | NADH | C00004 | $PQ_{body}$ |
| m02552[p] | NAD+ | C00003 | $PQ_{body}$ |
| m02553[p] | NADH | C00004 | $PQ_{body}$ |
| m02039[p] | H+ | C00080 | $PQ_{body}$ |
| m02552[m] | NAD+ | C00003 | $PQ_{body}$ |
| m02553[m] | NADH | C00004 | $PQ_{body}$ |
| m02444[c] | malonyl-CoA | C00083 | $PQ_{body}$ |
| m01596[c] | CO2 | C00011 | $PQ_{body}$ |
| m02630[p] | O2 | C00007 | $PQ_{body}$ |
| m02040[p] | H2O | C00001 | $PQ_{body}$ |
| m02041[p] | H2O2 | C00027 | $PQ_{body}$ |
| m01261[c] | acetyl-CoA | C00024 | $PQ_{body}$ |
| m01690[c] | DHAP | C00111 | $PQ_{body}$ |
| m02026[c] | GSH | C00051 | $PQ_{body}$ |
| m03142[c] | vitamin D3 | C05443 | $PQ_{body}$ |
| m01415[c] | calcidiol | C01561 | $PQ_{body}$ |
| m01802[m] | FAD | C00016 | $PQ_{body}$ |
| m01596[s] | CO2 | C00011 | $PQ_{body}$ |
| m02041[s] | H2O2 | C00027 | $PQ_{body}$ |
| m01934[c] | gamma-linolenoyl-CoA | C03035 | $PQ_{body}$ |
| m00859[c] | 3-oxo-dihomo-gamma-linolenoyl-CoA | | $PQ_{body}$ |
| m01261[m] | acetyl-CoA | C00024 | $PQ_{body}$ |
| m00113[c] | (6Z,9Z,12Z,15Z,18Z,21Z)-tetracosahexaenoyl-CoA | C16168 | $PQ_{body}$ |
| m01127[c] | 5-oxoproline | C01879 | $PQ_{body}$ |
| m03102[m] | ubiquinol | C00390 | $PQ_{body}$ |
| m03103[m] | ubiquinone | C00399 | $PQ_{body}$ |
| m02348[c] | L-carnitine | C15025 | $PQ_{body}$ |
| m02348[m] | L-carnitine | C15025 | $PQ_{body}$ |
| m01261[p] | acetyl-CoA | C00024 | $PQ_{body}$ |

| | | | |
|---|---|---|---|
| m01364[c] | arachidonyl-CoA | C02249 | $PQ_{body}$ |
| km00504[r] | methyl (2E,6E)-farnesoate | C16503 | $PQ_{body}$ |
| km00182[r] | juvenile hormone III | C09694 | $PQ_{body}$ |
| m02578[c] | NH3 | C00014 | $PQ_{body}$ |
| m00606[m] | 22beta-hydroxycholesterol | C05502 | $PQ_{body}$ |
| m00579[m] | 20alpha,22beta dihydroxycholesterol | C05501 | $PQ_{body}$ |
| m02182[m] | isocaproic-aldehyde | C02373 | $PQ_{body}$ |
| m02763[m] | pregnenolone | C01953 | $PQ_{body}$ |
| m01078[m] | 5beta-cholestan-3alpha,7alpha,12alpha-triol | C05454 | $PQ_{body}$ |
| m01090[m] | 5beta-cholestane-3alpha,7alpha,12alpha,26-tetrol | C05446 | $PQ_{body}$ |
| m02751[c] | Pi | C00009 | $PQ_{body}$ |
| m01803[m] | FADH2 | C01352 | $PQ_{body}$ |
| m02751[m] | Pi | C00009 | $PQ_{body}$ |
| m02475[c] | methylglyoxal | C00546 | $PQ_{body}$ |
| m02833[r] | retinoate | C00777 | $PQ_{body}$ |
| m00113[p] | (6Z,9Z,12Z,15Z,18Z,21Z)-tetracosahexaenoyl-CoA | C16168 | $PQ_{body}$ |
| m00095[c] | (4Z,7Z,10Z,13Z,16Z,19Z)-docosahexaenoyl-CoA | C16169 | $PQ_{body}$ |
| m00095[p] | (4Z,7Z,10Z,13Z,16Z,19Z)-docosahexaenoyl-CoA | C16169 | $PQ_{body}$ |
| km00877[r] | [Reduced NADPH---hemoprotein reductase] | C03024 | $PQ_{body}$ |
| km00878[r] | [Oxidized NADPH---hemoprotein reductase] | C03161 | $PQ_{body}$ |
| m00427[r] | 18-hydroxy-all-trans-retinoate | C16679 | $PQ_{body}$ |
| m00752[m] | 3alpha,7alpha,12alpha-trihydroxy-5beta-cholestanate | C04722 | $PQ_{body}$ |
| m00750[m] | 3alpha,7alpha,12alpha-trihydroxy-5beta-cholestan-26-al | C01301 | $PQ_{body}$ |
| m00671[m] | 2-oxobutyrate | C00109 | $PQ_{body}$ |
| m01596[m] | CO2 | C00011 | $PQ_{body}$ |
| m02774[m] | propanoyl-CoA | C00100 | $PQ_{body}$ |
| m00639[c] | 2-deoxy-D-ribose-1-phosphate | C00672 | $PQ_{body}$ |
| m02159[c] | hypoxanthine | C00262 | $PQ_{body}$ |
| m01671[c] | deoxyinosine | C05512 | $PQ_{body}$ |
| m02547[c] | N-acetylputrescine | C02714 | $PQ_{body}$ |
| m02812[c] | putrescine | C00134 | $PQ_{body}$ |
| m02880[c] | sarcosine | C00213 | $PQ_{body}$ |
| m02877[c] | SAM | C00019 | $PQ_{body}$ |
| m02871[c] | SAH | C00021 | $PQ_{body}$ |
| m02681[c] | PAP | C00054 | $PQ_{body}$ |
| m00432[c] | 19-hydroxytestosterone | C05294 | $PQ_{body}$ |
| m00434[c] | 19-oxo-testosterone | C05295 | $PQ_{body}$ |
| m01833[c] | formate | C00058 | $PQ_{body}$ |
| m01787[c] | estradiol-17beta | C00951 | $PQ_{body}$ |
| m02585[c] | nicotinate ribonucleotide | C01185 | $PQ_{body}$ |
| m02584[c] | nicotinate D-ribonucleoside | C05841 | $PQ_{body}$ |
| m02844[c] | ribose-1-phosphate | C00620 | $PQ_{body}$ |

| | | | |
|---|---|---|---|
| m02586[c] | nicotinate | C00253 | $PQ_{body}$ |
| m01349[c] | apo-[ACP] | C03688 | $PQ_{body}$ |
| m02741[c] | phosphopantetheine | C01134 | $PQ_{body}$ |
| m00184[c] | [ACP] | C00229 | $PQ_{body}$ |
| m02679[c] | pantetheine | C00831 | $PQ_{body}$ |
| m00554[c] | 1-phosphatidyl-1D-myo-inositol-5-phosphate | C11557 | $PQ_{body}$ |
| m02736[c] | phosphatidylinositol-4,5-bisphosphate | C04637 | $PQ_{body}$ |
| m02750[c] | PI pool | C01194 | $PQ_{body}$ |
| m02675[c] | palmitolate | C08362 | $PQ_{body}$ |
| m02675[s] | palmitolate | C08362 | $PQ_{body}$ |
| m00432[s] | 19-hydroxytestosterone | C05294 | $PQ_{body}$ |
| m02171[c] | inositol | C00137 | $PQ_{body}$ |
| m02519[s] | Na+ | C01330 | $PQ_{body}$ |
| m02519[c] | Na+ | C01330 | $PQ_{body}$ |
| m02171[s] | inositol | C00137 | $PQ_{body}$ |
| m00671[c] | 2-oxobutyrate | C00109 | $PQ_{body}$ |
| m01645[c] | dCTP | C00458 | $PQ_{body}$ |
| m01645[m] | dCTP | C00458 | $PQ_{body}$ |
| m01680[c] | dGDP | C00361 | $PQ_{body}$ |
| m01680[m] | dGDP | C00361 | $PQ_{body}$ |
| m00656[r] | 2-lysolecithin pool | C04230 | $PQ_{body}$ |
| m00656[l] | 2-lysolecithin pool | C04230 | $PQ_{body}$ |
| m00656[c] | 2-lysolecithin pool | C04230 | $PQ_{body}$ |
| m02403[s] | L-lactate | C00186 | $PQ_{body}$ |
| m01671[s] | deoxyinosine | C05512 | $PQ_{body}$ |
| m02452[s] | maltotriose | C01835 | $PQ_{body}$ |
| m01252[c] | acetate | C00033 | $PQ_{body}$ |
| m02845[c] | ribose-5-phosphate | C00117 | $PQ_{body}$ |
| km00615[c] | D-Ribofuranose | | $PQ_{body}$ |
| m03149[c] | xanthosine | C01762 | $PQ_{body}$ |
| m03148[c] | xanthine | C00385 | $PQ_{body}$ |
| m03161[c] | glycogen | C00182 | $\alpha Syn_{head}$ |
| m01824[m] | ferricytochrome C | C00125 | $\alpha Syn_{head}$ |
| m01826[m] | ferrocytochrome C | C00126 | $\alpha Syn_{head}$ |
| m00671[m] | 2-oxobutyrate | C00109 | $\alpha Syn_{head}$ |
| m01255[c] | acetoacetyl-CoA | C00332 | $\alpha Syn_{head}$ |
| m01639[c] | dAMP | C00360 | $\alpha Syn_{head}$ |
| m00266[c] | 10-formyl-THF | C00234 | $\alpha Syn_{head}$ |
| m02046[c] | HCO3- | C00288 | $\alpha Syn_{head}$ |
| m01644[c] | dCMP | C00239 | $\alpha Syn_{head}$ |
| m01045[c] | 5,10-methylene-THF | C00143 | $\alpha Syn_{head}$ |
| m01721[c] | DNA | C00039 | $\alpha Syn_{head}$ |

| | | | |
|---|---|---|---|
| m02751[l] | Pi | C00009 | $\alpha Syn_{head}$ |
| m02578[m] | NH3 | C00014 | $\alpha Syn_{head}$ |
| m01369[c] | asparagine | C00152 | $\alpha Syn_{head}$ |
| m02942[m] | succinate semialdehyde | C00232 | $\alpha Syn_{head}$ |
| m00970[c] | 4-aminobutyrate | C00334 | $\alpha Syn_{head}$ |
| m00970[m] | 4-aminobutyrate | C00334 | $\alpha Syn_{head}$ |
| m02125[c] | histidine | C00135 | $\alpha Syn_{head}$ |
| m02880[m] | sarcosine | C00213 | $\alpha Syn_{head}$ |
| m01045[m] | 5,10-methylene-THF | C00143 | $\alpha Syn_{head}$ |
| m02880[c] | sarcosine | C00213 | $\alpha Syn_{head}$ |
| m01654[c] | dehydroalanine | C02218 | $\alpha Syn_{head}$ |
| m01393[c] | betaine | C00719 | $\alpha Syn_{head}$ |
| m01708[c] | dimethylglycine | C01026 | $\alpha Syn_{head}$ |
| m02981[c] | THF-hexaglutamate | | $\alpha Syn_{head}$ |
| m01255[m] | acetoacetyl-CoA | C00332 | $\alpha Syn_{head}$ |
| m02319[c] | kynurenine | C00328 | $\alpha Syn_{head}$ |
| m02184[c] | isoleucine | C00407 | $\alpha Syn_{head}$ |
| m02360[c] | leucine | C00123 | $\alpha Syn_{head}$ |
| m02371[c] | L-formylkynurenine | C02700 | $\alpha Syn_{head}$ |
| m02349[c] | L-cystathionine | C02291 | $\alpha Syn_{head}$ |
| m01721[n] | DNA | C00039 | $\alpha Syn_{head}$ |
| m01974[s] | glutamate | C00025 | $\alpha Syn_{head}$ |
| m00097[s] | (5-L-glutamyl)-L-amino acid | C03740 | $\alpha Syn_{head}$ |
| m03075[c] | tRNA(met) | C01647 | $\alpha Syn_{head}$ |
| m02131[m] | HMG-CoA | C00356 | $\alpha Syn_{head}$ |
| m03081[c] | tRNA(tyr) | C00787 | $\alpha Syn_{head}$ |
| m02421[c] | L-tyrosyl-tRNA(tyr) | C02839 | $\alpha Syn_{head}$ |
| m03063[c] | tRNA(ala) | C01635 | $\alpha Syn_{head}$ |
| m02335[c] | L-alanyl-tRNA(ala) | C00886 | $\alpha Syn_{head}$ |
| m03064[c] | tRNA(arg) | C01636 | $\alpha Syn_{head}$ |
| m02340[c] | L-arginyl-tRNA(arg) | C02163 | $\alpha Syn_{head}$ |
| m03065[c] | tRNA(asn) | C01637 | $\alpha Syn_{head}$ |
| m02341[c] | L-asparaginyl-tRNA(asn) | C03402 | $\alpha Syn_{head}$ |
| m03066[c] | tRNA(asp) | C01638 | $\alpha Syn_{head}$ |
| m02342[c] | L-aspartyl-tRNA(asp) | C02984 | $\alpha Syn_{head}$ |
| m03067[c] | tRNA(cys) | C01639 | $\alpha Syn_{head}$ |
| m02351[c] | L-cysteinyl-tRNA(cys) | C03125 | $\alpha Syn_{head}$ |
| m03068[c] | tRNA(gln) | C01640 | $\alpha Syn_{head}$ |
| m02376[c] | L-glutaminyl-tRNA(gln) | C02282 | $\alpha Syn_{head}$ |
| m03069[c] | tRNA(glu) | C01641 | $\alpha Syn_{head}$ |
| m02377[c] | L-glutamyl-tRNA(glu) | C02987 | $\alpha Syn_{head}$ |
| m03070[c] | tRNA(gly) | C01642 | $\alpha Syn_{head}$ |

| | | | |
|---|---|---|---|
| m02006[c] | glycyl-tRNA(gly) | C02412 | αSyn$_{head}$ |
| m03071[c] | tRNA(his) | C01643 | αSyn$_{head}$ |
| m02380[c] | L-histidyl-tRNA(his) | C02988 | αSyn$_{head}$ |
| m03072[c] | tRNA(ile) | C01644 | αSyn$_{head}$ |
| m02401[c] | L-isoleucyl-tRNA(ile) | C03127 | αSyn$_{head}$ |
| m03073[c] | tRNA(leu) | C01645 | αSyn$_{head}$ |
| m02404[c] | L-leucyl-tRNA(leu) | C02047 | αSyn$_{head}$ |
| m03074[c] | tRNA(lys) | C01646 | αSyn$_{head}$ |
| m02405[c] | L-lysyl-tRNA(lys) | C01931 | αSyn$_{head}$ |
| m02408[c] | L-methionyl-tRNA(met) | C02430 | αSyn$_{head}$ |
| m03076[c] | tRNA(phe) | C01648 | αSyn$_{head}$ |
| m02412[c] | L-phenylalanyl-tRNA(phe) | C03511 | αSyn$_{head}$ |
| m03077[c] | tRNA(pro) | C01649 | αSyn$_{head}$ |
| m02415[c] | L-prolyl-tRNA(pro) | C02702 | αSyn$_{head}$ |
| m03078[c] | tRNA(ser) | C01650 | αSyn$_{head}$ |
| m02416[c] | L-seryl-tRNA(ser) | C02553 | αSyn$_{head}$ |
| m03079[c] | tRNA(thr) | C01651 | αSyn$_{head}$ |
| m02419[c] | L-threonyl-tRNA(thr) | C02992 | αSyn$_{head}$ |
| m03080[c] | tRNA(trp) | C01652 | αSyn$_{head}$ |
| m02420[c] | L-tryptophanyl-tRNA(trp) | C03512 | αSyn$_{head}$ |
| m03082[c] | tRNA(val) | C01653 | αSyn$_{head}$ |
| m02423[c] | L-valyl-tRNA(val) | C02554 | αSyn$_{head}$ |
| m02039[i] | H+ | C00080 | αSyn$_{head}$ |
| m02631[n] | O2- | C00704 | αSyn$_{head}$ |
| m02344[c] | lauric acid | C02679 | αSyn$_{head}$ |
| m03051[c] | tridecylic acid | C17076 | αSyn$_{head}$ |
| m03050[c] | tridecanoyl-CoA | | αSyn$_{head}$ |
| m00128[c] | (9E)-tetradecenoic acid | | αSyn$_{head}$ |
| m00129[c] | (9E)-tetradecenoyl-CoA | | αSyn$_{head}$ |
| m00117[c] | (7Z)-tetradecenoic acid | | αSyn$_{head}$ |
| m00118[c] | (7Z)-tetradecenoyl-CoA | | αSyn$_{head}$ |
| m02745[c] | physeteric acid | | αSyn$_{head}$ |
| m01141[c] | 5-tetradecenoyl-CoA | | αSyn$_{head}$ |
| m02690[c] | pentadecylic acid | C16537 | αSyn$_{head}$ |
| m02689[c] | pentadecanoyl-CoA | | αSyn$_{head}$ |
| m02675[c] | palmitolate | C08362 | αSyn$_{head}$ |
| m01197[c] | 7-palmitoleic acid | | αSyn$_{head}$ |
| m01191[c] | 7-hexadecenoyl-CoA | | αSyn$_{head}$ |
| m02456[c] | margaric acid | | αSyn$_{head}$ |
| m02101[c] | heptadecanoyl-CoA | | αSyn$_{head}$ |
| m00003[c] | (10Z)-heptadecenoic acid | | αSyn$_{head}$ |
| m00004[c] | (10Z)-heptadecenoyl-CoA | | αSyn$_{head}$ |

| | | | |
|---|---|---|---|
| m01238[c] | 9-heptadecylenic acid | C16536 | $\alpha Syn_{head}$ |
| m01237[c] | 9-heptadecenoyl-CoA | | $\alpha Syn_{head}$ |
| m02938[c] | stearate | C01530 | $\alpha Syn_{head}$ |
| m00019[c] | (13Z)-octadecenoic acid | | $\alpha Syn_{head}$ |
| m00020[c] | (13Z)-octadecenoyl-CoA | | $\alpha Syn_{head}$ |
| m01585[c] | cis-vaccenic acid | C08367 | $\alpha Syn_{head}$ |
| m01586[c] | cis-vaccenoyl-CoA | C21945 | $\alpha Syn_{head}$ |
| m02646[c] | oleate | C00712 | $\alpha Syn_{head}$ |
| m01778[c] | elaidate | C00712 | $\alpha Syn_{head}$ |
| m00127[c] | (9E)-octadecenoyl-CoA | | $\alpha Syn_{head}$ |
| m00115[c] | (7Z)-octadecenoic acid | | $\alpha Syn_{head}$ |
| m00104[c] | (6Z,9Z)-octadecadienoic acid | | $\alpha Syn_{head}$ |
| m00106[c] | (6Z,9Z)-octadecadienoyl-CoA | | $\alpha Syn_{head}$ |
| m02613[c] | nonadecylic acid | C16535 | $\alpha Syn_{head}$ |
| m02612[c] | nonadecanoyl-CoA | | $\alpha Syn_{head}$ |
| m01771[c] | eicosanoate | C06425 | $\alpha Syn_{head}$ |
| m01773[c] | eicosanoyl-CoA | C02041 | $\alpha Syn_{head}$ |
| m00017[c] | (13Z)-eicosenoic acid | | $\alpha Syn_{head}$ |
| m00018[c] | (13Z)-eicosenoyl-CoA | | $\alpha Syn_{head}$ |
| m01584[c] | cis-gondoic acid | C16526 | $\alpha Syn_{head}$ |
| m00007[c] | (11Z)-eicosenoyl-CoA | C16530 | $\alpha Syn_{head}$ |
| m01235[c] | 9-eicosenoic acid | | $\alpha Syn_{head}$ |
| m01236[c] | 9-eicosenoyl-CoA | | $\alpha Syn_{head}$ |
| m01207[c] | 8,11-eicosadienoic acid | | $\alpha Syn_{head}$ |
| m00123[c] | (8Z,11Z)-eicosadienoyl-CoA | C21937 | $\alpha Syn_{head}$ |
| m02457[c] | mead acid | | $\alpha Syn_{head}$ |
| m00101[c] | (5Z,8Z,11Z)-eicosatrienoyl-CoA | C21939 | $\alpha Syn_{head}$ |
| m02053[c] | henicosanoic acid | | $\alpha Syn_{head}$ |
| m02052[c] | heneicosanoyl-CoA | | $\alpha Syn_{head}$ |
| m01373[c] | behenic acid | C08281 | $\alpha Syn_{head}$ |
| m01725[c] | docosanoyl-CoA | C16528 | $\alpha Syn_{head}$ |
| m01583[c] | cis-erucic acid | C08316 | $\alpha Syn_{head}$ |
| m00016[c] | (13Z)-docosenoyl-CoA | C16531 | $\alpha Syn_{head}$ |
| m01582[c] | cis-cetoleic acid | | $\alpha Syn_{head}$ |
| m00006[c] | (11Z)-docosenoyl-CoA | | $\alpha Syn_{head}$ |
| m03045[c] | tricosanoic acid | | $\alpha Syn_{head}$ |
| m03047[c] | tricosanoyl-CoA | | $\alpha Syn_{head}$ |
| m02385[c] | lignocerate | C08320 | $\alpha Syn_{head}$ |
| m02971[c] | tetracosanoyl-CoA | C16529 | $\alpha Syn_{head}$ |
| m02564[c] | nervonic acid | C08323 | $\alpha Syn_{head}$ |
| m01432[c] | cerotic acid | | $\alpha Syn_{head}$ |
| m03153[c] | ximenic acid | | $\alpha Syn_{head}$ |

| | | | |
|---|---|---|---|
| m02112[c] | hexacosenoyl-CoA | | αSyn$_{head}$ |
| m02389[c] | linolenate | C06427 | αSyn$_{head}$ |
| m02939[c] | stearidonic acid | C16300 | αSyn$_{head}$ |
| m00108[c] | (6Z,9Z,12Z,15Z)-octadecatetraenoyl-CoA | C16163 | αSyn$_{head}$ |
| m02648[c] | omega-3-arachidonic acid | | αSyn$_{head}$ |
| m00125[c] | (8Z,11Z,14Z,17Z)-eicosatetraenoyl-CoA | C16164 | αSyn$_{head}$ |
| m01784[c] | EPA | C06428 | αSyn$_{head}$ |
| m00103[c] | (5Z,8Z,11Z,14Z,17Z)-eicosapentaenoyl-CoA | C16165 | αSyn$_{head}$ |
| m01741[c] | DPA | C16513 | αSyn$_{head}$ |
| m00121[c] | (7Z,10Z,13Z,16Z,19Z)-docosapentaenoyl-CoA | C16166 | αSyn$_{head}$ |
| m00135[c] | (9Z,12Z,15Z,18Z,21Z)-TPA | | αSyn$_{head}$ |
| m00134[c] | (9Z,12Z,15Z,18Z,21Z)-tetracosapentaenoyl-CoA | C16167 | αSyn$_{head}$ |
| m00114[c] | (6Z,9Z,12Z,15Z,18Z,21Z)-THA | | αSyn$_{head}$ |
| m01689[c] | DHA | C06429 | αSyn$_{head}$ |
| m00010[c] | (11Z,14Z,17Z)-eicosatrienoic acid | C16522 | αSyn$_{head}$ |
| m00012[c] | (11Z,14Z,17Z)-eicosatrienoyl-CoA | C16179 | αSyn$_{head}$ |
| m00341[c] | 13,16,19-docosatrienoic acid | C16534 | αSyn$_{head}$ |
| m00343[c] | 13,16,19-docosatrienoyl-CoA | | αSyn$_{head}$ |
| m00260[c] | 10,13,16,19-docosatetraenoic acid | | αSyn$_{head}$ |
| m00262[c] | 10,13,16,19-docosatetraenoyl-CoA | | αSyn$_{head}$ |
| m00315[c] | 12,15,18,21-tetracosatetraenoic acid | | αSyn$_{head}$ |
| m00317[c] | 12,15,18,21-tetracosatetraenoyl-CoA | | αSyn$_{head}$ |
| m02387[c] | linoleate | C01595 | αSyn$_{head}$ |
| m01932[c] | gamma-linolenate | C06426 | αSyn$_{head}$ |
| m01696[c] | dihomo-gamma-linolenate | C03242 | αSyn$_{head}$ |
| m01697[c] | dihomo-gamma-linolenoyl-CoA | C03595 | αSyn$_{head}$ |
| m01291[c] | adrenic acid | C16527 | αSyn$_{head}$ |
| m00119[c] | (7Z,10Z,13Z,16Z)-docosatetraenoyl-CoA | C16170 | αSyn$_{head}$ |
| m00132[c] | (9Z,12Z,15Z,18Z)-TTA | | αSyn$_{head}$ |
| m00111[c] | (6Z,9Z,12Z,15Z,18Z)-TPA | | αSyn$_{head}$ |
| m00094[c] | (4Z,7Z,10Z,13Z,16Z)-DPA | | αSyn$_{head}$ |
| m00008[c] | (11Z,14Z)-eicosadienoic acid | C16525 | αSyn$_{head}$ |
| m00009[c] | (11Z,14Z)-eicosadienoyl-CoA | C16180 | αSyn$_{head}$ |
| m00021[c] | (13Z,16Z)-docosadienoic acid | C16533 | αSyn$_{head}$ |
| m00023[c] | (13Z,16Z)-docosadienoyl-CoA | C16645 | αSyn$_{head}$ |
| m00265[c] | 10,13,16-docosatriynoic acid | | αSyn$_{head}$ |
| m00264[c] | 10,13,16-docosatrienoyl-CoA | | αSyn$_{head}$ |
| m03048[c] | tridecanoyl-[ACP] | | αSyn$_{head}$ |
| m00896[c] | 3-oxopentadecanoyl-[ACP] | | αSyn$_{head}$ |
| m00794[c] | 3-hydroxypentadecanoyl-[ACP] | | αSyn$_{head}$ |
| m00060[c] | (2E)-pentadecenoyl-[ACP] | | αSyn$_{head}$ |
| m02687[c] | pentadecanoyl-[ACP] | | αSyn$_{head}$ |

| | | | |
|---|---|---|---|
| m00874[c] | 3-oxoheptadecanoyl-[ACP] | | αSyn$_{head}$ |
| m00779[c] | 3-hydroxyheptadecanoyl-[ACP] | | αSyn$_{head}$ |
| m00045[c] | (2E)-heptadecenoyl-[ACP] | | αSyn$_{head}$ |
| m02099[c] | heptadecanoyl-[ACP] | | αSyn$_{head}$ |
| m00840[c] | 3-oxo-11-cis-eicosenoyl-CoA | | αSyn$_{head}$ |
| m00700[c] | 3(S)-hydroxy-11-cis-eicosenoyl-CoA | | αSyn$_{head}$ |
| m03012[c] | trans,cis-2,11-eicosadienoyl-CoA | | αSyn$_{head}$ |
| m00856[c] | 3-oxo-cis-15-tetracosaenoyl-CoA | | αSyn$_{head}$ |
| m00709[c] | 3(S)-hydroxy-cis-15-tetracosaenoyl-CoA | | αSyn$_{head}$ |
| m03015[c] | trans,cis-2,15-tetracosadienoyl-CoA | | αSyn$_{head}$ |
| m00116[c] | (7Z)-octadecenoyl-CoA | | αSyn$_{head}$ |
| m00871[c] | 3-oxo-eicosadienoyl-CoA | | αSyn$_{head}$ |
| m00704[c] | 3(S)-hydroxy-3-oxo-eicosadienoyl-CoA | | αSyn$_{head}$ |
| m03017[c] | trans,cis-2,8,11-eicosatrienoyl-CoA | | αSyn$_{head}$ |
| m00872[c] | 3-oxoeicosanoyl-CoA | | αSyn$_{head}$ |
| m00777[c] | 3-hydroxyeicosanoyl-CoA | | αSyn$_{head}$ |
| m00043[c] | (2E)-eicosenoyl-CoA | | αSyn$_{head}$ |
| m00800[c] | 3-hydroxytetracosanoyl-CoA | | αSyn$_{head}$ |
| m00064[c] | (2E)-tetracosenoyl-CoA | | αSyn$_{head}$ |
| m00887[c] | 3-oxononadecanoyl-CoA | | αSyn$_{head}$ |
| m00790[c] | 3-hydroxynonadecanoyl-CoA | | αSyn$_{head}$ |
| m00054[c] | (2E)-nonadecenoyl-CoA | | αSyn$_{head}$ |
| m02674[p] | palmitate | C00249 | αSyn$_{head}$ |
| m00811[c] | 3-keto-eicosa-8,11,14,17-all-cis-tetraenoyl-CoA | | αSyn$_{head}$ |
| m00708[c] | 3(S)-hydroxy-all-cis-8,11,14,17-eicosatetraenoyl-CoA | | αSyn$_{head}$ |
| m01769[c] | eicosa-(2E,8Z,11Z,14Z,17Z)-pentaenoyl-CoA | | αSyn$_{head}$ |
| m00716[c] | 3(S)-hydroxy-tetracosa-12,15,18,21-all-cis-tetraenoyl-CoA | | αSyn$_{head}$ |
| m03003[c] | trans,cis,cis,cis,cis-2,12,15,18,21-tetracosapentaenoyl-CoA | | αSyn$_{head}$ |
| m00869[c] | 3-oxoeicosa-cis,cis,cis-11,14,17-trienoyl-CoA | | αSyn$_{head}$ |
| m00086[c] | (3S)-hydroxy-eicosa-cis,cis,cis-11,14,17-trienoyl-CoA | | αSyn$_{head}$ |
| m03005[c] | trans,cis,cis,cis-2,11,14,17-eicosatetraenoyl-CoA | | αSyn$_{head}$ |
| m00710[c] | 3(S)-hydroxy-dihomo-gamma-linolenoyl-CoA | | αSyn$_{head}$ |
| m03029[c] | trans-2-cis,cis,cis-8,11,14-eicosatetraenoyl-CoA | | αSyn$_{head}$ |
| m00903[c] | 3-oxo-tetracosa-9,12,15,18-all-cis-tetraenoyl-CoA | | αSyn$_{head}$ |
| m00681[c] | 2-trans-9,12,15,18-all-cis-tetracosapentaenoyl-CoA | | αSyn$_{head}$ |
| m00870[c] | 3-oxoeicosa-cis,cis-11,14-dienoyl-CoA | | αSyn$_{head}$ |
| m00087[c] | (3S)-hydroxy-eicosa-cis,cis-11,14-dienoyl-CoA | | αSyn$_{head}$ |
| m03007[c] | trans,cis,cis-2,11,14-eicosatrienoyl-CoA | | αSyn$_{head}$ |
| m02103[c] | heptadecenoylcarnitine(8) | | αSyn$_{head}$ |
| m02940[c] | stearoylcarnitine | | αSyn$_{head}$ |
| m02410[c] | L-oleoylcarnitine | | αSyn$_{head}$ |
| m00105[c] | (6Z,9Z)-octadecadienoylcarnitine | | αSyn$_{head}$ |

| | | | |
|---|---|---|---|
| m01724[c] | docosanoylcarnitine | | $\alpha Syn_{head}$ |
| m01727[c] | docosenoylcarnitine | | $\alpha Syn_{head}$ |
| m00107[c] | (6Z,9Z,12Z,15Z)-octadecatetraenoylcarnitine | | $\alpha Syn_{head}$ |
| m00120[c] | (7Z,10Z,13Z,16Z,19Z)-docosapentaenoylcarnitine | | $\alpha Syn_{head}$ |
| m00092[c] | (4Z,7Z,10Z,13Z,16Z)-docosapentaenoylcarnitine | | $\alpha Syn_{head}$ |
| m02103[m] | heptadecenoylcarnitine(8) | | $\alpha Syn_{head}$ |
| m01237[m] | 9-heptadecenoyl-CoA | | $\alpha Syn_{head}$ |
| m02940[m] | stearoylcarnitine | | $\alpha Syn_{head}$ |
| m02941[m] | stearoyl-CoA | C00412 | $\alpha Syn_{head}$ |
| m02410[m] | L-oleoylcarnitine | | $\alpha Syn_{head}$ |
| m02647[m] | oleoyl-CoA | C00510 | $\alpha Syn_{head}$ |
| m00105[m] | (6Z,9Z)-octadecadienoylcarnitine | | $\alpha Syn_{head}$ |
| m00106[m] | (6Z,9Z)-octadecadienoyl-CoA | | $\alpha Syn_{head}$ |
| m01773[m] | eicosanoyl-CoA | C02041 | $\alpha Syn_{head}$ |
| m00007[m] | (11Z)-eicosenoyl-CoA | C16530 | $\alpha Syn_{head}$ |
| m01724[m] | docosanoylcarnitine | | $\alpha Syn_{head}$ |
| m01725[m] | docosanoyl-CoA | C16528 | $\alpha Syn_{head}$ |
| m01727[m] | docosenoylcarnitine | | $\alpha Syn_{head}$ |
| m00016[m] | (13Z)-docosenoyl-CoA | C16531 | $\alpha Syn_{head}$ |
| m00107[m] | (6Z,9Z,12Z,15Z)-octadecatetraenoylcarnitine | | $\alpha Syn_{head}$ |
| m00108[m] | (6Z,9Z,12Z,15Z)-octadecatetraenoyl-CoA | C16163 | $\alpha Syn_{head}$ |
| m00120[m] | (7Z,10Z,13Z,16Z,19Z)-docosapentaenoylcarnitine | | $\alpha Syn_{head}$ |
| m00121[m] | (7Z,10Z,13Z,16Z,19Z)-docosapentaenoyl-CoA | C16166 | $\alpha Syn_{head}$ |
| m00092[m] | (4Z,7Z,10Z,13Z,16Z)-docosapentaenoylcarnitine | | $\alpha Syn_{head}$ |
| m00093[m] | (4Z,7Z,10Z,13Z,16Z)-docosapentaenoyl-CoA | C16173 | $\alpha Syn_{head}$ |
| m01813[c] | fatty acid-LD-TG1 pool | C00162 | $\alpha Syn_{head}$ |
| m00240[c] | 1,2-diacylglycerol-LD-TAG pool | C00641 | $\alpha Syn_{head}$ |
| m02733[c] | phosphatidate-LD-TAG pool | C00416 | $\alpha Syn_{head}$ |
| m00509[c] | 1-acylglycerol-LD-TG1 pool | C01885 | $\alpha Syn_{head}$ |
| m01814[c] | fatty acid-LD-TG2 pool | C00162 | $\alpha Syn_{head}$ |
| m01808[c] | fatty acid-LD-PC pool | C00162 | $\alpha Syn_{head}$ |
| m00236[c] | 1,2-diacylglycerol-LD-PE pool | C00641 | $\alpha Syn_{head}$ |
| m00505[c] | 1-acylglycerol-LD-PE pool | C01885 | $\alpha Syn_{head}$ |
| m01809[c] | fatty acid-LD-PE pool | C00162 | $\alpha Syn_{head}$ |
| m00508[c] | 1-acylglycerol-LD-SM pool | C01885 | $\alpha Syn_{head}$ |
| m01812[c] | fatty acid-LD-SM pool | C00162 | $\alpha Syn_{head}$ |
| m01276[c] | acyl-CoA-LD-TG2 pool | | $\alpha Syn_{head}$ |
| m01278[c] | acylglycerone-phosphate | C03372 | $\alpha Syn_{head}$ |
| m00040[m] | (2E)-docosenoyl-CoA | | $\alpha Syn_{head}$ |
| m00776[m] | 3-hydroxydocosanoyl-CoA | | $\alpha Syn_{head}$ |
| m00043[m] | (2E)-eicosenoyl-CoA | | $\alpha Syn_{head}$ |
| m00777[m] | 3-hydroxyeicosanoyl-CoA | | $\alpha Syn_{head}$ |

| | | | |
|---|---|---|---|
| m00872[m] | 3-oxoeicosanoyl-CoA | | $\alpha Syn_{head}$ |
| m03013[m] | trans,cis-2,13-docosadienoyl-CoA | | $\alpha Syn_{head}$ |
| m00701[m] | 3(S)-hydroxy-13-cis-docosenoyl-CoA | | $\alpha Syn_{head}$ |
| m00842[m] | 3-oxo-13cis-docosenoyl-CoA | | $\alpha Syn_{head}$ |
| m03012[m] | trans,cis-2,11-eicosadienoyl-CoA | | $\alpha Syn_{head}$ |
| m00700[m] | 3(S)-hydroxy-11-cis-eicosenoyl-CoA | | $\alpha Syn_{head}$ |
| m00840[m] | 3-oxo-11-cis-eicosenoyl-CoA | | $\alpha Syn_{head}$ |
| m02187[c] | isopentenyl-pPP | C00129 | $\alpha Syn_{head}$ |
| m01450[c] | cholesterol | C00187 | $\alpha Syn_{head}$ |
| m00971[r] | 4-androstene-3,17-dione | C00280 | $\alpha Syn_{head}$ |
| m01659[c] | dehydroepiandrosterone sulfate | C04555 | $\alpha Syn_{head}$ |
| m02131[c] | HMG-CoA | C00356 | $\alpha Syn_{head}$ |
| m00167[c] | (R)-mevalonate | C00418 | $\alpha Syn_{head}$ |
| m00165[c] | (R)-5-phosphomevalonate | C01107 | $\alpha Syn_{head}$ |
| m00164[c] | (R)-5-diphosphomevalonate | C01143 | $\alpha Syn_{head}$ |
| m00432[c] | 19-hydroxytestosterone | C05294 | $\alpha Syn_{head}$ |
| m01789[c] | estrone 3-sulfate | C02538 | $\alpha Syn_{head}$ |
| m00986[c] | 4-hydroxy-17beta-estradiol | C14209 | $\alpha Syn_{head}$ |
| m00986[r] | 4-hydroxy-17beta-estradiol | C14209 | $\alpha Syn_{head}$ |
| m01451[r] | cholesterol-ester pool | C02530 | $\alpha Syn_{head}$ |
| m01699[c] | dihydroceramide pool | | $\alpha Syn_{head}$ |
| m01430[c] | ceramide pool | C00195 | $\alpha Syn_{head}$ |
| m02684[c] | PC-LD pool | C00157 | $\alpha Syn_{head}$ |
| m02927[c] | sphinganine | C00836 | $\alpha Syn_{head}$ |
| m02928[c] | sphinganine-1-phosphate | C01120 | $\alpha Syn_{head}$ |
| m01275[c] | acyl-CoA-LD-SM pool | | $\alpha Syn_{head}$ |
| m02929[c] | sphingosine | C00319 | $\alpha Syn_{head}$ |
| m02930[c] | sphingosine-1-phosphate | C06124 | $\alpha Syn_{head}$ |
| m01679[l] | D-galactosyl-N-acylsphingosine | C02686 | $\alpha Syn_{head}$ |
| m01694[l] | digalactosylceramide | C06126 | $\alpha Syn_{head}$ |
| m02931[c] | sphingosylphosphorylcholine | C03640 | $\alpha Syn_{head}$ |
| m00484[c] | 1-acylglycerol-3P-laur | | $\alpha Syn_{head}$ |
| m00502[c] | 1-acylglycerol-3P-tridec | | $\alpha Syn_{head}$ |
| m00493[c] | 1-acylglycerol-3P-myrist | | $\alpha Syn_{head}$ |
| m00471[c] | 1-acylglycerol-3P-9-tetrade | | $\alpha Syn_{head}$ |
| m00463[c] | 1-acylglycerol-3P-7-tetrade | | $\alpha Syn_{head}$ |
| m00454[c] | 1-acylglycerol-3P-5-tetrade | | $\alpha Syn_{head}$ |
| m00498[c] | 1-acylglycerol-3P-pentade | | $\alpha Syn_{head}$ |
| m00496[c] | 1-acylglycerol-3P-palm | | $\alpha Syn_{head}$ |
| m00497[c] | 1-acylglycerol-3P-palmn | | $\alpha Syn_{head}$ |
| m00461[c] | 1-acylglycerol-3P-7-hexade | | $\alpha Syn_{head}$ |
| m00481[c] | 1-acylglycerol-3P-heptade | | $\alpha Syn_{head}$ |

| | | | |
|---|---|---|---|
| m00438[c] | 1-acylglycerol-3P-10-heptade | | $\alpha Syn_{head}$ |
| m00469[c] | 1-acylglycerol-3P-9-heptade | | $\alpha Syn_{head}$ |
| m00499[c] | 1-acylglycerol-3P-stea | | $\alpha Syn_{head}$ |
| m00448[c] | 1-acylglycerol-3P-13-octade | | $\alpha Syn_{head}$ |
| m00474[c] | 1-acylglycerol-3P-cis-vac | | $\alpha Syn_{head}$ |
| m00495[c] | 1-acylglycerol-3P-ol | | $\alpha Syn_{head}$ |
| m00470[c] | 1-acylglycerol-3P-9-octade | | $\alpha Syn_{head}$ |
| m00462[c] | 1-acylglycerol-3P-7-octade | | $\alpha Syn_{head}$ |
| m00458[c] | 1-acylglycerol-3P-6,9-octa | | $\alpha Syn_{head}$ |
| m00494[c] | 1-acylglycerol-3P-nanode | | $\alpha Syn_{head}$ |
| m00478[c] | 1-acylglycerol-3P-eico | | $\alpha Syn_{head}$ |
| m00447[c] | 1-acylglycerol-3P-13-eicose | | $\alpha Syn_{head}$ |
| m00442[c] | 1-acylglycerol-3P-11-eico | | $\alpha Syn_{head}$ |
| m00468[c] | 1-acylglycerol-3P-9-eicose | | $\alpha Syn_{head}$ |
| m00465[c] | 1-acylglycerol-3P-8,11-eico | | $\alpha Syn_{head}$ |
| m00453[c] | 1-acylglycerol-3P-5,8,11-eico | | $\alpha Syn_{head}$ |
| m00480[c] | 1-acylglycerol-3P-heneico | | $\alpha Syn_{head}$ |
| m00477[c] | 1-acylglycerol-3P-docosa | | $\alpha Syn_{head}$ |
| m00446[c] | 1-acylglycerol-3P-13-docose | | $\alpha Syn_{head}$ |
| m00441[c] | 1-acylglycerol-3P-11-docose | | $\alpha Syn_{head}$ |
| m00501[c] | 1-acylglycerol-3P-trico | | $\alpha Syn_{head}$ |
| m00500[c] | 1-acylglycerol-3P-tetraco | | $\alpha Syn_{head}$ |
| m00449[c] | 1-acylglycerol-3P-15-tetra | | $\alpha Syn_{head}$ |
| m00482[c] | 1-acylglycerol-3P-hexacosa | | $\alpha Syn_{head}$ |
| m00483[c] | 1-acylglycerol-3P-hexecose | | $\alpha Syn_{head}$ |
| m00492[c] | 1-acylglycerol-3P-linolen | | $\alpha Syn_{head}$ |
| m00457[c] | 1-acylglycerol-3P-6,9,12,15-octa | | $\alpha Syn_{head}$ |
| m00464[c] | 1-acylglycerol-3P-8,11,14,17-eico | | $\alpha Syn_{head}$ |
| m00452[c] | 1-acylglycerol-3P-5,8,11,14,17-eico | | $\alpha Syn_{head}$ |
| m00459[c] | 1-acylglycerol-3P-7,10,13,16,19-docosa | | $\alpha Syn_{head}$ |
| m00466[c] | 1-acylglycerol-3P-9,12,15,18,21-tetra | | $\alpha Syn_{head}$ |
| m00455[c] | 1-acylglycerol-3P-6,9,12,15,18,21-tetra | | $\alpha Syn_{head}$ |
| m00450[c] | 1-acylglycerol-3P-4,7,10,13,16,19-doco | | $\alpha Syn_{head}$ |
| m00439[c] | 1-acylglycerol-3P-11,14,17-eico | | $\alpha Syn_{head}$ |
| m00444[c] | 1-acylglycerol-3P-13,16,19-doco | | $\alpha Syn_{head}$ |
| m00436[c] | 1-acylglycerol-3P-10,13,16,19-doco | | $\alpha Syn_{head}$ |
| m00443[c] | 1-acylglycerol-3P-12,15,18,21-tetra | | $\alpha Syn_{head}$ |
| m00491[c] | 1-acylglycerol-3P-lin | | $\alpha Syn_{head}$ |
| m00479[c] | 1-acylglycerol-3P-gamma-lin | | $\alpha Syn_{head}$ |
| m00476[c] | 1-acylglycerol-3P-dihomo-gamma | | $\alpha Syn_{head}$ |
| m00472[c] | 1-acylglycerol-3P-arach | | $\alpha Syn_{head}$ |
| m00460[c] | 1-acylglycerol-3P-7,10,13,16-docosa | | $\alpha Syn_{head}$ |

| | | | |
|---|---|---|---|
| m00467[c] | 1-acylglycerol-3P-9,12,15,18-tetraco | | $\alpha Syn_{head}$ |
| m00456[c] | 1-acylglycerol-3P-6,9,12,15,18-tetraco | | $\alpha Syn_{head}$ |
| m00451[c] | 1-acylglycerol-3P-4,7,10,13,16-docosa | | $\alpha Syn_{head}$ |
| m00440[c] | 1-acylglycerol-3P-11,14-eicosa | | $\alpha Syn_{head}$ |
| m00445[c] | 1-acylglycerol-3P-13,16-docosa | | $\alpha Syn_{head}$ |
| m00437[c] | 1-acylglycerol-3P-10,13,16-docosa | | $\alpha Syn_{head}$ |
| m00490[c] | 1-acylglycerol-3P-LD-TG1 pool | C00681 | $\alpha Syn_{head}$ |
| m00485[c] | 1-acylglycerol-3P-LD-PC pool | C00681 | $\alpha Syn_{head}$ |
| m00486[c] | 1-acylglycerol-3P-LD-PE pool | C00681 | $\alpha Syn_{head}$ |
| m00489[c] | 1-acylglycerol-3P-LD-SM pool | C00681 | $\alpha Syn_{head}$ |
| m01425[c] | CDP-choline | C00307 | $\alpha Syn_{head}$ |
| m01271[c] | acyl-CoA-LD-PC pool | | $\alpha Syn_{head}$ |
| m02728[c] | phosphatidate-LD-PC pool | C00416 | $\alpha Syn_{head}$ |
| m01272[c] | acyl-CoA-LD-PE pool | | $\alpha Syn_{head}$ |
| m02729[c] | phosphatidate-LD-PE pool | C00416 | $\alpha Syn_{head}$ |
| m02750[c] | PI pool | C01194 | $\alpha Syn_{head}$ |
| m02685[c] | PE-LD pool | C00350 | $\alpha Syn_{head}$ |
| m02808[c] | PS-LD pool | C02737 | $\alpha Syn_{head}$ |
| m01797[c] | ethanolamine | C00189 | $\alpha Syn_{head}$ |
| m00656[c] | 2-lysolecithin pool | C04230 | $\alpha Syn_{head}$ |
| m02912[c] | sn-glycerol-3-PC | C00670 | $\alpha Syn_{head}$ |
| m00511[c] | 1-acyl-PE pool | C04438 | $\alpha Syn_{head}$ |
| m00549[c] | 1-organyl-2-lyso-sn-glycero-3-phosphocholine | C04317 | $\alpha Syn_{head}$ |
| m00560[c] | 1-radyl-2-acyl-sn-glycero-3-phosphocholine | C05212 | $\alpha Syn_{head}$ |
| m01694[c] | digalactosylceramide | C06126 | $\alpha Syn_{head}$ |
| m02519[s] | Na+ | C01330 | $\alpha Syn_{head}$ |
| m02519[c] | Na+ | C01330 | $\alpha Syn_{head}$ |
| m00267[c] | 10-formyl-THF-glu(5) | | $\alpha Syn_{head}$ |
| m00267[l] | 10-formyl-THF-glu(5) | | $\alpha Syn_{head}$ |
| m00266[l] | 10-formyl-THF | C00234 | $\alpha Syn_{head}$ |
| m01974[l] | glutamate | C00025 | $\alpha Syn_{head}$ |
| m02426[s] | lysine | C00047 | $\alpha Syn_{head}$ |
| m01155[c] | 6-[(1S,2R)-1,2-dihydroxy-3-triphosphooxypropyl]-7,8-dihydropterin | C04895 | $\alpha Syn_{head}$ |
| m01155[n] | 6-[(1S,2R)-1,2-dihydroxy-3-triphosphooxypropyl]-7,8-dihydropterin | C04895 | $\alpha Syn_{head}$ |
| m01833[n] | formate | C00058 | $\alpha Syn_{head}$ |
| m03057[c] | triphosphate | C00536 | $\alpha Syn_{head}$ |
| m01704[c] | dihydroneopterin | C04874 | $\alpha Syn_{head}$ |
| m03057[n] | triphosphate | C00536 | $\alpha Syn_{head}$ |
| m02978[n] | tetrahydrobiopterin | C00272 | $\alpha Syn_{head}$ |
| m03057[m] | triphosphate | C00536 | $\alpha Syn_{head}$ |
| m00325[c] | 12-hydroxydodecanoic acid | C08317 | $\alpha Syn_{head}$ |
| m00371[c] | 14-hydroxytetradecanoic acid | | $\alpha Syn_{head}$ |

| | | | |
|---|---|---|---|
| m00403[c] | 16-hydroxyhexadecanoic acid | C18218 | $\alpha Syn_{head}$ |
| m01442[c] | chloride | C00698 | $\alpha Syn_{head}$ |
| m02360[s] | leucine | C00123 | $\alpha Syn_{head}$ |
| m02392[c] | lipid droplet | | $\alpha Syn_{head}$ |
| m01450[s] | cholesterol | C00187 | $\alpha Syn_{head}$ |
| m01442[s] | chloride | C00698 | $\alpha Syn_{head}$ |
| m02125[s] | histidine | C00135 | $\alpha Syn_{head}$ |
| m02471[s] | methionine | C00073 | $\alpha Syn_{head}$ |
| m01975[s] | glutamine | C00064 | $\alpha Syn_{head}$ |
| m02724[s] | phenylalanine | C00079 | $\alpha Syn_{head}$ |
| m01369[s] | asparagine | C00152 | $\alpha Syn_{head}$ |
| m03135[s] | valine | C00183 | $\alpha Syn_{head}$ |
| m02184[s] | isoleucine | C00407 | $\alpha Syn_{head}$ |
| m01370[s] | aspartate | C00049 | $\alpha Syn_{head}$ |
| m01833[s] | formate | C00058 | $\alpha Syn_{head}$ |
| m02772[s] | propanoate | C00163 | $\alpha Syn_{head}$ |
| m02833[s] | retinoate | C00777 | $\alpha Syn_{head}$ |
| m01789[s] | estrone 3-sulfate | C02538 | $\alpha Syn_{head}$ |
| m01659[s] | dehydroepiandrosterone sulfate | C04555 | $\alpha Syn_{head}$ |
| m01433[s] | cGMP | C00942 | $\alpha Syn_{head}$ |
| m01158[c] | 6beta-hydroxytestosterone | C14497 | $\alpha Syn_{head}$ |
| m01158[s] | 6beta-hydroxytestosterone | C14497 | $\alpha Syn_{head}$ |
| m02928[s] | sphinganine-1-phosphate | C01120 | $\alpha Syn_{head}$ |
| m01338[s] | androsterone | C00523 | $\alpha Syn_{head}$ |
| m02991[s] | thiosulfate | C00320 | $\alpha Syn_{head}$ |
| m00325[s] | 12-hydroxydodecanoic acid | C08317 | $\alpha Syn_{head}$ |
| m00371[s] | 14-hydroxytetradecanoic acid | | $\alpha Syn_{head}$ |
| m00403[s] | 16-hydroxyhexadecanoic acid | C18218 | $\alpha Syn_{head}$ |
| m00986[s] | 4-hydroxy-17beta-estradiol | C14209 | $\alpha Syn_{head}$ |
| m01285[l] | ADP | C00008 | $\alpha Syn_{head}$ |
| m01371[l] | ATP | C00002 | $\alpha Syn_{head}$ |
| m01365[l] | arginine | C00062 | $\alpha Syn_{head}$ |
| m01369[l] | asparagine | C00152 | $\alpha Syn_{head}$ |
| m01628[l] | cysteine | C00097 | $\alpha Syn_{head}$ |
| m01975[l] | glutamine | C00064 | $\alpha Syn_{head}$ |
| m02125[l] | histidine | C00135 | $\alpha Syn_{head}$ |
| m02360[l] | leucine | C00123 | $\alpha Syn_{head}$ |
| m02426[l] | lysine | C00047 | $\alpha Syn_{head}$ |
| m02471[l] | methionine | C00073 | $\alpha Syn_{head}$ |
| m02724[l] | phenylalanine | C00079 | $\alpha Syn_{head}$ |
| m02770[l] | proline | C00148 | $\alpha Syn_{head}$ |
| m02896[l] | serine | C00065 | $\alpha Syn_{head}$ |

| | | | |
|---|---|---|---|
| m03089[l] | tryptophan | C00078 | $\alpha Syn_{head}$ |
| m03101[l] | tyrosine | C00082 | $\alpha Syn_{head}$ |
| m03135[l] | valine | C00183 | $\alpha Syn_{head}$ |
| m01307[l] | alanine | C00041 | $\alpha Syn_{head}$ |
| m02184[l] | isoleucine | C00407 | $\alpha Syn_{head}$ |
| m02993[l] | threonine | C00188 | $\alpha Syn_{head}$ |
| m01602[c] | cofactors and vitamins | | $\alpha Syn_{head}$ |
| m02319[s] | kynurenine | C00328 | $\alpha Syn_{head}$ |
| m02040[c] | H2O | C00001 | $\alpha Syn_{head}$ |
| m02040[s] | H2O | C00001 | $\alpha Syn_{head}$ |
| m02877[c] | SAM | C00019 | $\alpha Syn_{head}$ |
| m02871[c] | SAH | C00021 | $\alpha Syn_{head}$ |
| m02039[c] | H+ | C00080 | $\alpha Syn_{head}$ |
| m02877[m] | SAM | C00019 | $\alpha Syn_{head}$ |
| m02871[m] | SAH | C00021 | $\alpha Syn_{head}$ |
| m02039[m] | H+ | C00080 | $\alpha Syn_{head}$ |
| m02877[n] | SAM | C00019 | $\alpha Syn_{head}$ |
| m02871[n] | SAH | C00021 | $\alpha Syn_{head}$ |
| m02039[n] | H+ | C00080 | $\alpha Syn_{head}$ |
| m01371[c] | ATP | C00002 | $\alpha Syn_{head}$ |
| m01285[c] | ADP | C00008 | $\alpha Syn_{head}$ |
| m01306[n] | AKG | C00026 | $\alpha Syn_{head}$ |
| m02630[n] | O2 | C00007 | $\alpha Syn_{head}$ |
| m01596[n] | CO2 | C00011 | $\alpha Syn_{head}$ |
| m01831[n] | formaldehyde | C00067 | $\alpha Syn_{head}$ |
| m02943[n] | succinate | C00042 | $\alpha Syn_{head}$ |
| m01628[c] | cysteine | C00097 | $\alpha Syn_{head}$ |
| m01986[c] | glycine | C00037 | $\alpha Syn_{head}$ |
| m01986[s] | glycine | C00037 | $\alpha Syn_{head}$ |
| m02040[m] | H2O | C00001 | $\alpha Syn_{head}$ |
| m01986[m] | glycine | C00037 | $\alpha Syn_{head}$ |
| m01983[c] | glycerol | C00116 | $\alpha Syn_{head}$ |
| m02914[c] | sn-glycerol-3-phosphate | C00093 | $\alpha Syn_{head}$ |
| m01371[m] | ATP | C00002 | $\alpha Syn_{head}$ |
| m01285[m] | ADP | C00008 | $\alpha Syn_{head}$ |
| m02039[r] | H+ | C00080 | $\alpha Syn_{head}$ |
| m03106[r] | UDP | C00015 | $\alpha Syn_{head}$ |
| m02555[n] | NADPH | C00005 | $\alpha Syn_{head}$ |
| m02554[n] | NADP+ | C00006 | $\alpha Syn_{head}$ |
| m02041[c] | H2O2 | C00027 | $\alpha Syn_{head}$ |
| m01597[c] | CoA | C00010 | $\alpha Syn_{head}$ |
| m01597[m] | CoA | C00010 | $\alpha Syn_{head}$ |

| | | | |
|---|---|---|---|
| m02630[r] | O2 | C00007 | $\alpha Syn_{head}$ |
| km00013[r] | Red-NADPH-Hemoprotein-Reductases | | $\alpha Syn_{head}$ |
| km00014[r] | Ox-NADPH-Hemoprotein-Reductases | | $\alpha Syn_{head}$ |
| m02040[r] | H2O | C00001 | $\alpha Syn_{head}$ |
| m02555[c] | NADPH | C00005 | $\alpha Syn_{head}$ |
| m02554[c] | NADP+ | C00006 | $\alpha Syn_{head}$ |
| m02555[m] | NADPH | C00005 | $\alpha Syn_{head}$ |
| m02554[m] | NADP+ | C00006 | $\alpha Syn_{head}$ |
| m01974[c] | glutamate | C00025 | $\alpha Syn_{head}$ |
| m02579[c] | NH4+ | C00014 | $\alpha Syn_{head}$ |
| m01974[m] | glutamate | C00025 | $\alpha Syn_{head}$ |
| m01306[m] | AKG | C00026 | $\alpha Syn_{head}$ |
| m02034[c] | GTP | C00044 | $\alpha Syn_{head}$ |
| m01948[c] | GDP | C00035 | $\alpha Syn_{head}$ |
| m02630[c] | O2 | C00007 | $\alpha Syn_{head}$ |
| m02555[r] | NADPH | C00005 | $\alpha Syn_{head}$ |
| m02554[r] | NADP+ | C00006 | $\alpha Syn_{head}$ |
| m01280[c] | adenosine | C00212 | $\alpha Syn_{head}$ |
| m02133[c] | homocysteine | C00155 | $\alpha Syn_{head}$ |
| m02552[s] | NAD+ | C00003 | $\alpha Syn_{head}$ |
| m01362[c] | arachidonate | C00219 | $\alpha Syn_{head}$ |
| km00013[c] | Red-NADPH-Hemoprotein-Reductases | | $\alpha Syn_{head}$ |
| km00014[c] | Ox-NADPH-Hemoprotein-Reductases | | $\alpha Syn_{head}$ |
| m02844[c] | ribose-1-phosphate | C00620 | $\alpha Syn_{head}$ |
| m01252[m] | acetate | C00033 | $\alpha Syn_{head}$ |
| m02040[n] | H2O | C00001 | $\alpha Syn_{head}$ |
| m02630[m] | O2 | C00007 | $\alpha Syn_{head}$ |
| m01334[c] | AMP | C00020 | $\alpha Syn_{head}$ |
| m02759[c] | PPi | C00013 | $\alpha Syn_{head}$ |
| m02774[c] | propanoyl-CoA | C00100 | $\alpha Syn_{head}$ |
| m02552[c] | NAD+ | C00003 | $\alpha Syn_{head}$ |
| m02552[m] | NAD+ | C00003 | $\alpha Syn_{head}$ |
| m02553[m] | NADH | C00004 | $\alpha Syn_{head}$ |
| m01368[c] | ascorbate | C00072 | $\alpha Syn_{head}$ |
| m01368[s] | ascorbate | C00072 | $\alpha Syn_{head}$ |
| m02630[s] | O2 | C00007 | $\alpha Syn_{head}$ |
| m02444[c] | malonyl-CoA | C00083 | $\alpha Syn_{head}$ |
| m01596[c] | CO2 | C00011 | $\alpha Syn_{head}$ |
| m02040[l] | H2O | C00001 | $\alpha Syn_{head}$ |
| m01910[l] | galactose | C00984 | $\alpha Syn_{head}$ |
| m01752[c] | dTMP | C00364 | $\alpha Syn_{head}$ |
| m02630[p] | O2 | C00007 | $\alpha Syn_{head}$ |

| | | | |
|---|---|---|---|
| m02034[n] | GTP | C00044 | αSyn$_{head}$ |
| m01334[m] | AMP | C00020 | αSyn$_{head}$ |
| m02759[m] | PPi | C00013 | αSyn$_{head}$ |
| m01261[c] | acetyl-CoA | C00024 | αSyn$_{head}$ |
| m01596[m] | CO2 | C00011 | αSyn$_{head}$ |
| m03101[c] | tyrosine | C00082 | αSyn$_{head}$ |
| m03099[c] | tyramine | C00483 | αSyn$_{head}$ |
| m01690[c] | DHAP | C00111 | αSyn$_{head}$ |
| m00514[c] | 1-alkyl-2-acylglycerol | C03201 | αSyn$_{head}$ |
| m01590[c] | CMP | C00055 | αSyn$_{head}$ |
| m02944[m] | succinyl-CoA | C00091 | αSyn$_{head}$ |
| m02943[m] | succinate | C00042 | αSyn$_{head}$ |
| m02647[c] | oleoyl-CoA | C00510 | αSyn$_{head}$ |
| m02555[p] | NADPH | C00005 | αSyn$_{head}$ |
| m02554[p] | NADP+ | C00006 | αSyn$_{head}$ |
| m00918[c] | 3-sulfinoalanine | C00606 | αSyn$_{head}$ |
| m00131[c] | (9Z,12Z,15Z,18Z)-tetracosatetraenoyl-CoA | C16171 | αSyn$_{head}$ |
| m03142[c] | vitamin D3 | C05443 | αSyn$_{head}$ |
| m01415[c] | calcidiol | C01561 | αSyn$_{head}$ |
| m01802[m] | FAD | C00016 | αSyn$_{head}$ |
| m01371[s] | ATP | C00002 | αSyn$_{head}$ |
| m01280[s] | adenosine | C00212 | αSyn$_{head}$ |
| m01334[s] | AMP | C00020 | αSyn$_{head}$ |
| m02471[c] | methionine | C00073 | αSyn$_{head}$ |
| m02943[c] | succinate | C00042 | αSyn$_{head}$ |
| m02772[c] | propanoate | C00163 | αSyn$_{head}$ |
| m02774[m] | propanoyl-CoA | C00100 | αSyn$_{head}$ |
| m02007[c] | glyoxalate | C00048 | αSyn$_{head}$ |
| m01831[c] | formaldehyde | C00067 | αSyn$_{head}$ |
| m02553[r] | NADH | C00004 | αSyn$_{head}$ |
| m02552[r] | NAD+ | C00003 | αSyn$_{head}$ |
| m02350[c] | L-cysteate | C00506 | αSyn$_{head}$ |
| m01968[c] | glucose-6-phosphate | C00092 | αSyn$_{head}$ |
| m01965[c] | glucose | C00031 | αSyn$_{head}$ |
| m02943[s] | succinate | C00042 | αSyn$_{head}$ |
| m00904[c] | 3-oxotetracosanoyl-CoA | | αSyn$_{head}$ |
| m02946[c] | sulfate | C00059 | αSyn$_{head}$ |
| m02631[c] | O2- | C00704 | αSyn$_{head}$ |
| m02631[s] | O2- | C00704 | αSyn$_{head}$ |
| m02631[p] | O2- | C00704 | αSyn$_{head}$ |
| m02391[c] | linoleoyl-CoA | C02050 | αSyn$_{head}$ |
| m02896[s] | serine | C00065 | αSyn$_{head}$ |

| | | | |
|---|---|---|---|
| m03089[c] | tryptophan | C00078 | $\alpha Syn_{head}$ |
| m02770[c] | proline | C00148 | $\alpha Syn_{head}$ |
| m02941[c] | stearoyl-CoA | C00412 | $\alpha Syn_{head}$ |
| m01370[c] | aspartate | C00049 | $\alpha Syn_{head}$ |
| m01370[l] | aspartate | C00049 | $\alpha Syn_{head}$ |
| m02354[c] | L-dopa | C00355 | $\alpha Syn_{head}$ |
| m01934[c] | gamma-linolenoyl-CoA | C03035 | $\alpha Syn_{head}$ |
| m00859[c] | 3-oxo-dihomo-gamma-linolenoyl-CoA | | $\alpha Syn_{head}$ |
| km00239[n] | Protein-L-arginine | C00613 | $\alpha Syn_{head}$ |
| m01975[c] | glutamine | C00064 | $\alpha Syn_{head}$ |
| m02678[c] | palmitoyl-CoA | C00154 | $\alpha Syn_{head}$ |
| m01686[c] | dGMP | C00362 | $\alpha Syn_{head}$ |
| m01307[c] | alanine | C00041 | $\alpha Syn_{head}$ |
| m03108[c] | UDP-glucose | C00029 | $\alpha Syn_{head}$ |
| km00274[n] | Protein-N-omega-dimethyl-arginine | | $\alpha Syn_{head}$ |
| m02845[c] | ribose-5-phosphate | C00117 | $\alpha Syn_{head}$ |
| m01249[c] | acetaldehyde | C00084 | $\alpha Syn_{head}$ |
| m00110[c] | (6Z,9Z,12Z,15Z,18Z)-tetracosapentaenoyl-CoA | C16172 | $\alpha Syn_{head}$ |
| m02949[c] | sulfite | C00094 | $\alpha Syn_{head}$ |
| m02991[c] | thiosulfate | C00320 | $\alpha Syn_{head}$ |
| m02110[c] | hexacosanoyl-CoA | | $\alpha Syn_{head}$ |
| m02016[c] | GMP | C00144 | $\alpha Syn_{head}$ |
| m01365[c] | arginine | C00062 | $\alpha Syn_{head}$ |
| m02426[c] | lysine | C00047 | $\alpha Syn_{head}$ |
| m02896[c] | serine | C00065 | $\alpha Syn_{head}$ |
| m02495[c] | myristoyl-CoA | C02593 | $\alpha Syn_{head}$ |
| m01365[s] | arginine | C00062 | $\alpha Syn_{head}$ |
| m01327[c] | alpha-tocopherol | C02477 | $\alpha Syn_{head}$ |
| m00356[c] | 13-hydroxy-alpha-tocopherol | | $\alpha Syn_{head}$ |
| m02806[c] | PRPP | C00119 | $\alpha Syn_{head}$ |
| m00866[m] | 3-oxodocosanoyl-CoA | | $\alpha Syn_{head}$ |
| m00184[c] | [ACP] | C00229 | $\alpha Syn_{head}$ |
| m01513[c] | choline | C00114 | $\alpha Syn_{head}$ |
| m01261[m] | acetyl-CoA | C00024 | $\alpha Syn_{head}$ |
| m02969[r] | testosterone | C00535 | $\alpha Syn_{head}$ |
| m01158[r] | 6beta-hydroxytestosterone | C14497 | $\alpha Syn_{head}$ |
| m02847[c] | RNA | C00046 | $\alpha Syn_{head}$ |
| m02682[c] | PAPS | C00053 | $\alpha Syn_{head}$ |
| m02681[c] | PAP | C00054 | $\alpha Syn_{head}$ |
| m02046[s] | HCO3- | C00288 | $\alpha Syn_{head}$ |
| m02842[s] | riboflavin | C00255 | $\alpha Syn_{head}$ |
| m02677[c] | palmitoleoyl-CoA | C21072 | $\alpha Syn_{head}$ |

| | | | |
|---|---|---|---|
| m02678[p] | palmitoyl-CoA | C00154 | $\alpha Syn_{head}$ |
| m02724[c] | phenylalanine | C00079 | $\alpha Syn_{head}$ |
| m00113[c] | (6Z,9Z,12Z,15Z,18Z,21Z)-tetracosahexaenoyl-CoA | C16168 | $\alpha Syn_{head}$ |
| m02348[c] | L-carnitine | C15025 | $\alpha Syn_{head}$ |
| m02348[m] | L-carnitine | C15025 | $\alpha Syn_{head}$ |
| m01261[p] | acetyl-CoA | C00024 | $\alpha Syn_{head}$ |
| m01364[c] | arachidonyl-CoA | C02249 | $\alpha Syn_{head}$ |
| m02390[c] | linolenoyl-CoA | C16162 | $\alpha Syn_{head}$ |
| m01983[s] | glycerol | C00116 | $\alpha Syn_{head}$ |
| m02751[r] | Pi | C00009 | $\alpha Syn_{head}$ |
| m02993[c] | threonine | C00188 | $\alpha Syn_{head}$ |
| m00025[c] | (15Z)-tetracosenoyl-CoA | C16532 | $\alpha Syn_{head}$ |
| m02842[c] | riboflavin | C00255 | $\alpha Syn_{head}$ |
| m02908[c] | SM pool | C00550 | $\alpha Syn_{head}$ |
| m02738[c] | phosphocholine | C00588 | $\alpha Syn_{head}$ |
| m02173[c] | inositol-1-phosphate | C01177 | $\alpha Syn_{head}$ |
| m01986[l] | glycine | C00037 | $\alpha Syn_{head}$ |
| m01433[c] | cGMP | C00942 | $\alpha Syn_{head}$ |
| m01623[c] | CTP | C00063 | $\alpha Syn_{head}$ |
| km00615[c] | D-Ribofuranose | | $\alpha Syn_{head}$ |
| m02770[s] | proline | C00148 | $\alpha Syn_{head}$ |
| m02674[c] | palmitate | C00249 | $\alpha Syn_{head}$ |
| m02345[c] | lauroyl-CoA | C01832 | $\alpha Syn_{head}$ |
| m01307[s] | alanine | C00041 | $\alpha Syn_{head}$ |
| m02320[m] | L-1-pyrroline-3-hydroxy-5-carboxylate | C04281 | $\alpha Syn_{head}$ |
| m02494[c] | myristic acid | C06424 | $\alpha Syn_{head}$ |
| m02578[c] | NH3 | C00014 | $\alpha Syn_{head}$ |
| m03135[c] | valine | C00183 | $\alpha Syn_{head}$ |
| m01833[c] | formate | C00058 | $\alpha Syn_{head}$ |
| m01833[r] | formate | C00058 | $\alpha Syn_{head}$ |
| m02751[c] | Pi | C00009 | $\alpha Syn_{head}$ |
| m02354[s] | L-dopa | C00355 | $\alpha Syn_{head}$ |
| m01803[m] | FADH2 | C01352 | $\alpha Syn_{head}$ |
| m02980[c] | THF | C00101 | $\alpha Syn_{head}$ |
| m02980[m] | THF | C00101 | $\alpha Syn_{head}$ |
| m02751[m] | Pi | C00009 | $\alpha Syn_{head}$ |
| m02751[s] | Pi | C00009 | $\alpha Syn_{head}$ |
| m00671[c] | 2-oxobutyrate | C00109 | $\alpha Syn_{head}$ |
| m01790[c] | estrone | C00468 | $\alpha Syn_{head}$ |
| m01790[r] | estrone | C00468 | $\alpha Syn_{head}$ |
| m01338[c] | androsterone | C00523 | $\alpha Syn_{head}$ |
| m02969[c] | testosterone | C00535 | $\alpha Syn_{head}$ |

| | | | |
|---|---|---|---|
| m02833[c] | retinoate | C00777 | αSyn$_{head}$ |
| m02833[r] | retinoate | C00777 | αSyn$_{head}$ |
| m01787[c] | estradiol-17beta | C00951 | αSyn$_{head}$ |
| m01787[r] | estradiol-17beta | C00951 | αSyn$_{head}$ |
| m03037[m] | trans-4-hydroxy-L-proline | C01157 | αSyn$_{head}$ |
| m01972[c] | glucosylceramide pool | C01190 | αSyn$_{head}$ |
| m02442[c] | malonyl-[ACP] | C01209 | αSyn$_{head}$ |
| m01704[s] | dihydroneopterin | C04874 | αSyn$_{head}$ |
| m03107[c] | UDP-galactose | C00052 | αSyn$_{head}$ |
| m03114[r] | UMP | C00105 | αSyn$_{head}$ |
| m02387[s] | linoleate | C01595 | αSyn$_{head}$ |
| m00549[s] | 1-organyl-2-lyso-sn-glycero-3-phosphocholine | C04317 | αSyn$_{head}$ |
| m02389[s] | linolenate | C06427 | αSyn$_{head}$ |
| m00095[c] | (4Z,7Z,10Z,13Z,16Z,19Z)-docosahexaenoyl-CoA | C16169 | αSyn$_{head}$ |
| m00093[c] | (4Z,7Z,10Z,13Z,16Z)-docosapentaenoyl-CoA | C16173 | αSyn$_{head}$ |
| m01170[n] | 6-pyruvoyltetrahydropterin | C03684 | αSyn$_{head}$ |
| m02978[c] | tetrahydrobiopterin | C00272 | αSyn$_{head}$ |
| m01660[c] | dehydroepiandrosterone | C01227 | αSyn$_{head}$ |
| m01660[r] | dehydroepiandrosterone | C01227 | αSyn$_{head}$ |
| km00877[r] | [Reduced NADPH---hemoprotein reductase] | C03024 | αSyn$_{head}$ |
| km00878[r] | [Oxidized NADPH---hemoprotein reductase] | C03161 | αSyn$_{head}$ |
| m01679[c] | D-galactosyl-N-acylsphingosine | C02686 | αSyn$_{head}$ |
| m00427[r] | 18-hydroxy-all-trans-retinoate | C16679 | αSyn$_{head}$ |
| m01722[n] | DNA-5-methylcytosine | C02967 | αSyn$_{head}$ |
| m03130[s] | UTP | C00075 | αSyn$_{head}$ |
| m02759[s] | PPi | C00013 | αSyn$_{head}$ |
| m03114[s] | UMP | C00105 | αSyn$_{head}$ |
| m01668[c] | deoxycytidine | C00881 | αSyn$_{head}$ |
| m01673[c] | deoxyuridine | C00526 | αSyn$_{head}$ |
| m01630[c] | cytidine | C00475 | αSyn$_{head}$ |
| m03123[c] | uridine | C00299 | αSyn$_{head}$ |
| m03150[c] | xanthosine-5-phosphate | C00655 | αSyn$_{head}$ |
| m03154[c] | XTP | C00700 | αSyn$_{head}$ |
| m02167[c] | IMP | C00130 | αSyn$_{head}$ |
| m02193[c] | ITP | C00081 | αSyn$_{head}$ |
| m02026[c] | GSH | C00051 | αSyn$_{head}$ |
| m02027[c] | GSSG | C00127 | αSyn$_{head}$ |
| m01016[c] | 4-methylthio-2-oxobutanoic acid | C01180 | αSyn$_{head}$ |
| m00630[m] | 2-amino-3-oxoadipate | C05520 | αSyn$_{head}$ |
| m01074[m] | 5-aminolevulinate | C00430 | αSyn$_{head}$ |
| m01655[c] | dehydroascorbic acid | C00425 | αSyn$_{head}$ |
| m01736[c] | dopamine | C03758 | αSyn$_{head}$ |

| | | | |
|---|---|---|---|
| m02617[c] | noradrenaline | C00547 | $\alpha Syn_{head}$ |
| m01306[c] | AKG | C00026 | $\alpha Syn_{head}$ |
| m02553[c] | NADH | C00004 | $\alpha Syn_{head}$ |
| m01963[c] | glucosamine-6-phosphate | C00352 | $\alpha Syn_{head}$ |
| m01845[c] | fructose-6-phosphate | C00085 | $\alpha Syn_{head}$ |
| m02526[c] | N-acetylgalactosamine-1-phosphate | | $\alpha Syn_{head}$ |
| m02525[c] | N-acetylgalactosamine | C01074 | $\alpha Syn_{head}$ |
| m02161[c] | IDP | C00104 | $\alpha Syn_{head}$ |
| m01277[c] | acyl-CoA-LD-TG3 pool | | $\alpha Syn_{head}$ |
| m02958[c] | TAG-LD pool | C00422 | $\alpha Syn_{head}$ |
| m01815[c] | fatty acid-LD-TG3 pool | C00162 | $\alpha Syn_{head}$ |
| m00579[m] | 20alpha,22beta dihydroxycholesterol | C05501 | $\alpha Syn_{head}$ |
| m02182[m] | isocaproic-aldehyde | C02373 | $\alpha Syn_{head}$ |
| m02763[m] | pregnenolone | C01953 | $\alpha Syn_{head}$ |
| m01450[m] | cholesterol | C00187 | $\alpha Syn_{head}$ |
| m01069[c] | 5-alpha-dihydrotestosterone | C03917 | $\alpha Syn_{head}$ |
| m01430[g] | ceramide pool | C00195 | $\alpha Syn_{head}$ |
| m02684[g] | PC-LD pool | C00157 | $\alpha Syn_{head}$ |
| m00240[g] | 1,2-diacylglycerol-LD-TAG pool | C00641 | $\alpha Syn_{head}$ |
| m02908[g] | SM pool | C00550 | $\alpha Syn_{head}$ |
| m01972[r] | glucosylceramide pool | C01190 | $\alpha Syn_{head}$ |
| m02328[r] | LacCer pool | C01290 | $\alpha Syn_{head}$ |
| m03107[r] | UDP-galactose | C00052 | $\alpha Syn_{head}$ |
| m01435[s] | chenodiol | C02528 | $\alpha Syn_{head}$ |
| m01980[c] | glutathionyl-leukotriene C4 | | $\alpha Syn_{head}$ |
| m01980[s] | glutathionyl-leukotriene C4 | | $\alpha Syn_{head}$ |
| m01435[c] | chenodiol | C02528 | $\alpha Syn_{head}$ |
| m02900[c] | S-glutathionyl-2-4-dinitrobenzene | | $\alpha Syn_{head}$ |
| m02900[s] | S-glutathionyl-2-4-dinitrobenzene | | $\alpha Syn_{head}$ |
| m02741[c] | phosphopantetheine | C01134 | $\alpha Syn_{head}$ |
| m02679[c] | pantetheine | C00831 | $\alpha Syn_{head}$ |
| m00163[c] | (R)-4-phosphopantothenoyl-cysteine | C04352 | $\alpha Syn_{head}$ |
| m00552[c] | 1-phosphatidyl-1D-myo-inositol-3-phosphate | C04549 | $\alpha Syn_{head}$ |
| m00555[c] | 1-phosphatidyl-myo-inositol-3,5-bisphosphate | C11556 | $\alpha Syn_{head}$ |
| m00551[c] | 1-phosphatidyl-1D-myo-inositol-3,4-bisphosphate | C11554 | $\alpha Syn_{head}$ |
| m00553[c] | 1-phosphatidyl-1D-myo-inositol-4-phosphate | C01277 | $\alpha Syn_{head}$ |
| m00554[c] | 1-phosphatidyl-1D-myo-inositol-5-phosphate | C11557 | $\alpha Syn_{head}$ |
| m02736[c] | phosphatidylinositol-4,5-bisphosphate | C04637 | $\alpha Syn_{head}$ |
| m01700[m] | dihydrofolate | C00415 | $\alpha Syn_{head}$ |
| m01115[c] | 5-methyl-THF | C00440 | $\alpha Syn_{head}$ |
| m01830[c] | folate | C00504 | $\alpha Syn_{head}$ |
| m01700[c] | dihydrofolate | C00415 | $\alpha Syn_{head}$ |

| | | | |
|---|---|---|---|
| m01830[m] | folate | C00504 | $\alpha Syn_{head}$ |
| m01802[c] | FAD | C00016 | $\alpha Syn_{head}$ |
| m01803[c] | FADH2 | C01352 | $\alpha Syn_{head}$ |
| m01330[r] | alpha-tocotrienol | C14153 | $\alpha Syn_{head}$ |
| m00357[r] | 13-hydroxy-alpha-tocotrienol | | $\alpha Syn_{head}$ |
| m00345[r] | 13-carboxy-alpha-tocotrienol | | $\alpha Syn_{head}$ |
| m03050[c] | tridecanoyl-CoA | | $\alpha Syn_{head}$ |
| m00129[c] | (9E)-tetradecenoyl-CoA | | $\alpha Syn_{head}$ |
| m00118[c] | (7Z)-tetradecenoyl-CoA | | $\alpha Syn_{head}$ |
| m01141[c] | 5-tetradecenoyl-CoA | | $\alpha Syn_{head}$ |
| m02689[c] | pentadecanoyl-CoA | | $\alpha Syn_{head}$ |
| m01191[c] | 7-hexadecenoyl-CoA | | $\alpha Syn_{head}$ |
| m02101[c] | heptadecanoyl-CoA | | $\alpha Syn_{head}$ |
| m00004[c] | (10Z)-heptadecenoyl-CoA | | $\alpha Syn_{head}$ |
| m01237[c] | 9-heptadecenoyl-CoA | | $\alpha Syn_{head}$ |
| m00020[c] | (13Z)-octadecenoyl-CoA | | $\alpha Syn_{head}$ |
| m00127[c] | (9E)-octadecenoyl-CoA | | $\alpha Syn_{head}$ |
| m00106[c] | (6Z,9Z)-octadecadienoyl-CoA | | $\alpha Syn_{head}$ |
| m02612[c] | nonadecanoyl-CoA | | $\alpha Syn_{head}$ |
| m00018[c] | (13Z)-eicosenoyl-CoA | | $\alpha Syn_{head}$ |
| m01236[c] | 9-eicosenoyl-CoA | | $\alpha Syn_{head}$ |
| m02052[c] | heneicosanoyl-CoA | | $\alpha Syn_{head}$ |
| m00006[c] | (11Z)-docosenoyl-CoA | | $\alpha Syn_{head}$ |
| m03047[c] | tricosanoyl-CoA | | $\alpha Syn_{head}$ |
| m02112[c] | hexacosenoyl-CoA | | $\alpha Syn_{head}$ |
| m00343[c] | 13,16,19-docosatrienoyl-CoA | | $\alpha Syn_{head}$ |
| m00262[c] | 10,13,16,19-docosatetraenoyl-CoA | | $\alpha Syn_{head}$ |
| m00317[c] | 12,15,18,21-tetracosatetraenoyl-CoA | | $\alpha Syn_{head}$ |
| m00264[c] | 10,13,16-docosatrienoyl-CoA | | $\alpha Syn_{head}$ |
| m00116[c] | (7Z)-octadecenoyl-CoA | | $\alpha Syn_{head}$ |
| m02110[c] | hexacosanoyl-CoA | | $\alpha Syn_{head}$ |
| m00128[c] | (9E)-tetradecenoic acid | | $\alpha Syn_{head}$ |
| m00117[c] | (7Z)-tetradecenoic acid | | $\alpha Syn_{head}$ |
| m02745[c] | physeteric acid | | $\alpha Syn_{head}$ |
| m01197[c] | 7-palmitoleic acid | | $\alpha Syn_{head}$ |
| m02456[c] | margaric acid | | $\alpha Syn_{head}$ |
| m00003[c] | (10Z)-heptadecenoic acid | | $\alpha Syn_{head}$ |
| m00019[c] | (13Z)-octadecenoic acid | | $\alpha Syn_{head}$ |
| m00115[c] | (7Z)-octadecenoic acid | | $\alpha Syn_{head}$ |
| m00104[c] | (6Z,9Z)-octadecadienoic acid | | $\alpha Syn_{head}$ |
| m00017[c] | (13Z)-eicosenoic acid | | $\alpha Syn_{head}$ |
| m01235[c] | 9-eicosenoic acid | | $\alpha Syn_{head}$ |

| | | | |
|---|---|---|---|
| m01207[c] | 8,11-eicosadienoic acid | | $\alpha Syn_{head}$ |
| m02457[c] | mead acid | | $\alpha Syn_{head}$ |
| m02053[c] | henicosanoic acid | | $\alpha Syn_{head}$ |
| m01582[c] | cis-cetoleic acid | | $\alpha Syn_{head}$ |
| m03045[c] | tricosanoic acid | | $\alpha Syn_{head}$ |
| m01432[c] | cerotic acid | | $\alpha Syn_{head}$ |
| m03153[c] | ximenic acid | | $\alpha Syn_{head}$ |
| m02648[c] | omega-3-arachidonic acid | | $\alpha Syn_{head}$ |
| m00135[c] | (9Z,12Z,15Z,18Z,21Z)-TPA | | $\alpha Syn_{head}$ |
| m00114[c] | (6Z,9Z,12Z,15Z,18Z,21Z)-THA | | $\alpha Syn_{head}$ |
| m00260[c] | 10,13,16,19-docosatetraenoic acid | | $\alpha Syn_{head}$ |
| m00315[c] | 12,15,18,21-tetracosatetraenoic acid | | $\alpha Syn_{head}$ |
| m00132[c] | (9Z,12Z,15Z,18Z)-TTA | | $\alpha Syn_{head}$ |
| m00111[c] | (6Z,9Z,12Z,15Z,18Z)-TPA | | $\alpha Syn_{head}$ |
| m00094[c] | (4Z,7Z,10Z,13Z,16Z)-DPA | | $\alpha Syn_{head}$ |
| m00265[c] | 10,13,16-docosatriynoic acid | | $\alpha Syn_{head}$ |
| m02366[c] | leukotriene C4 | C02166 | $\alpha Syn_{head}$ |
| m02366[s] | leukotriene C4 | C02166 | $\alpha Syn_{head}$ |
| m02843[c] | ribose | C00121 | $\alpha Syn_{head}$ |
| m02843[s] | ribose | C00121 | $\alpha Syn_{head}$ |
| m02038[s] | guanosine | C00387 | $\alpha Syn_{head}$ |
| m02038[c] | guanosine | C00387 | $\alpha Syn_{head}$ |
| m02950[c] | sulfochenodeoxycholate | | $\alpha Syn_{head}$ |
| m02950[s] | sulfochenodeoxycholate | | $\alpha Syn_{head}$ |
| m02998[c] | thyroxine | C01829 | $\alpha Syn_{head}$ |
| m02901[c] | S-glutathionyl-ethacrynic acid | | $\alpha Syn_{head}$ |
| m02901[s] | S-glutathionyl-ethacrynic acid | | $\alpha Syn_{head}$ |
| m02998[s] | thyroxine | C01829 | $\alpha Syn_{head}$ |
| m03052[s] | triiodothyronine | C02465 | $\alpha Syn_{head}$ |
| m03052[c] | triiodothyronine | C02465 | $\alpha Syn_{head}$ |
| m00591[s] | 20-hydroxy-arachidonate | C14748 | $\alpha Syn_{head}$ |
| m00591[c] | 20-hydroxy-arachidonate | C14748 | $\alpha Syn_{head}$ |
| m01398[s] | bilirubin-monoglucuronoside | C03374 | $\alpha Syn_{head}$ |
| m01398[c] | bilirubin-monoglucuronoside | C03374 | $\alpha Syn_{head}$ |
| m02781[c] | prostaglandin C2 | C05955 | $\alpha Syn_{head}$ |
| m02781[s] | prostaglandin C2 | C05955 | $\alpha Syn_{head}$ |
| m02784[s] | prostaglandin D3 | C13802 | $\alpha Syn_{head}$ |
| m02784[c] | prostaglandin D3 | C13802 | $\alpha Syn_{head}$ |
| m02792[s] | prostaglandin G2 | C05956 | $\alpha Syn_{head}$ |
| m02792[c] | prostaglandin G2 | C05956 | $\alpha Syn_{head}$ |
| m02794[c] | prostaglandin H2 | C00427 | $\alpha Syn_{head}$ |
| m02794[s] | prostaglandin H2 | C00427 | $\alpha Syn_{head}$ |

| | | | |
|---|---|---|---|
| m02795[c] | prostaglandin I2 | C01312 | $\alpha Syn_{head}$ |
| m02795[s] | prostaglandin I2 | C01312 | $\alpha Syn_{head}$ |
| m02787[c] | prostaglandin E3 | C06439 | $\alpha Syn_{head}$ |
| m02787[s] | prostaglandin E3 | C06439 | $\alpha Syn_{head}$ |
| m02026[s] | GSH | C00051 | $\alpha Syn_{head}$ |
| m02780[c] | prostaglandin C1 | C04686 | $\alpha Syn_{head}$ |
| m02780[s] | prostaglandin C1 | C04686 | $\alpha Syn_{head}$ |
| m02782[s] | prostaglandin D1 | C06438 | $\alpha Syn_{head}$ |
| m02782[c] | prostaglandin D1 | C06438 | $\alpha Syn_{head}$ |
| m02790[s] | prostaglandin F2beta | C02314 | $\alpha Syn_{head}$ |
| m02790[c] | prostaglandin F2beta | C02314 | $\alpha Syn_{head}$ |
| m01668[s] | deoxycytidine | C00881 | $\alpha Syn_{head}$ |
| m02147[c] | hydroxide | C01328 | $\alpha Syn_{head}$ |
| m01830[s] | folate | C00504 | $\alpha Syn_{head}$ |
| m02147[s] | hydroxide | C01328 | $\alpha Syn_{head}$ |
| m02193[s] | ITP | C00081 | $\alpha Syn_{head}$ |
| m00648[s] | 2-hydroxybutyrate | C05984 | $\alpha Syn_{head}$ |
| m02039[s] | H+ | C00080 | $\alpha Syn_{head}$ |
| m00648[c] | 2-hydroxybutyrate | C05984 | $\alpha Syn_{head}$ |
| m03154[s] | XTP | C00700 | $\alpha Syn_{head}$ |
| m01688[m] | dGTP | C00286 | $\alpha Syn_{head}$ |
| m01747[c] | dTDP | C00363 | $\alpha Syn_{head}$ |
| m01747[m] | dTDP | C00363 | $\alpha Syn_{head}$ |
| m01688[c] | dGTP | C00286 | $\alpha Syn_{head}$ |
| m02552[p] | NAD+ | C00003 | $\alpha Syn_{head}$ |
| m02553[p] | NADH | C00004 | $\alpha Syn_{head}$ |
| m00553[n] | 1-phosphatidyl-1D-myo-inositol-4-phosphate | C01277 | $\alpha Syn_{head}$ |
| m03114[c] | UMP | C00105 | $\alpha Syn_{head}$ |
| m01330[c] | alpha-tocotrienol | C14153 | $\alpha Syn_{head}$ |
| m00345[c] | 13-carboxy-alpha-tocotrienol | | $\alpha Syn_{head}$ |
| m02328[c] | LacCer pool | C01290 | $\alpha Syn_{head}$ |
| m02370[s] | leukotriene F4 | C06462 | $\alpha Syn_{head}$ |
| m03109[c] | UDP-glucuronate | C00167 | $\alpha Syn_{head}$ |
| m03106[c] | UDP | C00015 | $\alpha Syn_{head}$ |
| km00800[c] | SN-38 | C11173 | $\alpha Syn_{head}$ |
| km00804[c] | SN-38G | C11376 | $\alpha Syn_{head}$ |
| m01973[c] | glucuronate | C00191 | $\alpha Syn_{head}$ |
| m03161[c] | glycogen | C00182 | $\alpha Syn_{body}$ |
| m01910[s] | galactose | C00984 | $\alpha Syn_{body}$ |
| m02148[c] | hydroxyacetone | C05235 | $\alpha Syn_{body}$ |
| m01256[c] | acetone | C00207 | $\alpha Syn_{body}$ |
| m01715[c] | D-lactaldehyde | C00937 | $\alpha Syn_{body}$ |

| | | | |
|---|---|---|---|
| m01253[m] | acetoacetate | C00164 | $\alpha Syn_{body}$ |
| m01256[m] | acetone | C00207 | $\alpha Syn_{body}$ |
| m01255[c] | acetoacetyl-CoA | C00332 | $\alpha Syn_{body}$ |
| m01253[c] | acetoacetate | C00164 | $\alpha Syn_{body}$ |
| m01639[c] | dAMP | C00360 | $\alpha Syn_{body}$ |
| m00266[c] | 10-formyl-THF | C00234 | $\alpha Syn_{body}$ |
| m03121[c] | urea | C00086 | $\alpha Syn_{body}$ |
| m03121[s] | urea | C00086 | $\alpha Syn_{body}$ |
| m02147[c] | hydroxide | C01328 | $\alpha Syn_{body}$ |
| m01644[c] | dCMP | C00239 | $\alpha Syn_{body}$ |
| m01045[c] | 5,10-methylene-THF | C00143 | $\alpha Syn_{body}$ |
| m01721[c] | DNA | C00039 | $\alpha Syn_{body}$ |
| m02751[l] | Pi | C00009 | $\alpha Syn_{body}$ |
| m02578[m] | NH3 | C00014 | $\alpha Syn_{body}$ |
| m01369[c] | asparagine | C00152 | $\alpha Syn_{body}$ |
| m00970[c] | 4-aminobutyrate | C00334 | $\alpha Syn_{body}$ |
| m02125[c] | histidine | C00135 | $\alpha Syn_{body}$ |
| m01045[m] | 5,10-methylene-THF | C00143 | $\alpha Syn_{body}$ |
| m02184[c] | isoleucine | C00407 | $\alpha Syn_{body}$ |
| m02360[c] | leucine | C00123 | $\alpha Syn_{body}$ |
| m01005[c] | 4-hydroxyphenylpyruvate | C01179 | $\alpha Syn_{body}$ |
| m01004[c] | 4-hydroxyphenyllactate | C03672 | $\alpha Syn_{body}$ |
| m00996[c] | 4-hydroxybenzoyl-CoA | C02949 | $\alpha Syn_{body}$ |
| m00995[c] | 4-hydroxybenzoate | C00156 | $\alpha Syn_{body}$ |
| m01721[n] | DNA | C00039 | $\alpha Syn_{body}$ |
| m01974[s] | glutamate | C00025 | $\alpha Syn_{body}$ |
| m00097[s] | (5-L-glutamyl)-L-amino acid | C03740 | $\alpha Syn_{body}$ |
| m01383[c] | beta-alanine | C00099 | $\alpha Syn_{body}$ |
| m03075[c] | tRNA(met) | C01647 | $\alpha Syn_{body}$ |
| m02131[m] | HMG-CoA | C00356 | $\alpha Syn_{body}$ |
| m03081[c] | tRNA(tyr) | C00787 | $\alpha Syn_{body}$ |
| m02421[c] | L-tyrosyl-tRNA(tyr) | C02839 | $\alpha Syn_{body}$ |
| m03063[c] | tRNA(ala) | C01635 | $\alpha Syn_{body}$ |
| m02335[c] | L-alanyl-tRNA(ala) | C00886 | $\alpha Syn_{body}$ |
| m03064[c] | tRNA(arg) | C01636 | $\alpha Syn_{body}$ |
| m02340[c] | L-arginyl-tRNA(arg) | C02163 | $\alpha Syn_{body}$ |
| m03066[c] | tRNA(asp) | C01638 | $\alpha Syn_{body}$ |
| m02342[c] | L-aspartyl-tRNA(asp) | C02984 | $\alpha Syn_{body}$ |
| m03069[c] | tRNA(glu) | C01641 | $\alpha Syn_{body}$ |
| m02377[c] | L-glutamyl-tRNA(glu) | C02987 | $\alpha Syn_{body}$ |
| m03070[c] | tRNA(gly) | C01642 | $\alpha Syn_{body}$ |
| m02006[c] | glycyl-tRNA(gly) | C02412 | $\alpha Syn_{body}$ |

| | | | |
|---|---|---|---|
| m03073[c] | tRNA(leu) | C01645 | $\alpha Syn_{body}$ |
| m02404[c] | L-leucyl-tRNA(leu) | C02047 | $\alpha Syn_{body}$ |
| m03074[c] | tRNA(lys) | C01646 | $\alpha Syn_{body}$ |
| m02405[c] | L-lysyl-tRNA(lys) | C01931 | $\alpha Syn_{body}$ |
| m02408[c] | L-methionyl-tRNA(met) | C02430 | $\alpha Syn_{body}$ |
| m03077[c] | tRNA(pro) | C01649 | $\alpha Syn_{body}$ |
| m02415[c] | L-prolyl-tRNA(pro) | C02702 | $\alpha Syn_{body}$ |
| m03078[c] | tRNA(ser) | C01650 | $\alpha Syn_{body}$ |
| m02416[c] | L-seryl-tRNA(ser) | C02553 | $\alpha Syn_{body}$ |
| m03082[c] | tRNA(val) | C01653 | $\alpha Syn_{body}$ |
| m02423[c] | L-valyl-tRNA(val) | C02554 | $\alpha Syn_{body}$ |
| m02394[c] | lipoic acid | C00725 | $\alpha Syn_{body}$ |
| m02398[c] | lipoyl-AMP | C16238 | $\alpha Syn_{body}$ |
| m00209[c] | [protein]-N6-(lipoyl)lysine | C16237 | $\alpha Syn_{body}$ |
| m00240[c] | 1,2-diacylglycerol-LD-TAG pool | C00641 | $\alpha Syn_{body}$ |
| m02733[c] | phosphatidate-LD-TAG pool | C00416 | $\alpha Syn_{body}$ |
| m02187[c] | isopentenyl-pPP | C00129 | $\alpha Syn_{body}$ |
| m01316[c] | all-trans-decaprenyl-diphosphate | C17432 | $\alpha Syn_{body}$ |
| m00995[m] | 4-hydroxybenzoate | C00156 | $\alpha Syn_{body}$ |
| m01316[m] | all-trans-decaprenyl-diphosphate | C17432 | $\alpha Syn_{body}$ |
| m00767[m] | 3-decaprenyl-4-hydroxybenzoate | | $\alpha Syn_{body}$ |
| m00725[m] | 3,4-dihydroxy-5-all-trans-decaprenylbenzoate | | $\alpha Syn_{body}$ |
| m00817[m] | 3-methoxy-4-hydroxy-5-all-trans-decaprenylbenzoate | | $\alpha Syn_{body}$ |
| m00657[m] | 2-methoxy-6-(all-trans-decaprenyl)phenol | | $\alpha Syn_{body}$ |
| m00658[m] | 2-methoxy-6-all trans-decaprenyl-2-methoxy-1,4-benzoquinol | | $\alpha Syn_{body}$ |
| m01165[m] | 6-methoxy-3-methyl-2-all-trans-decaprenyl-1,4-benzoquinol | | $\alpha Syn_{body}$ |
| m00770[m] | 3-demethylubiquinol-10 | | $\alpha Syn_{body}$ |
| m00971[r] | 4-androstene-3,17-dione | C00280 | $\alpha Syn_{body}$ |
| m01075[r] | 5-androstene-3,17-dione | C20252 | $\alpha Syn_{body}$ |
| m01659[c] | dehydroepiandrosterone sulfate | C04555 | $\alpha Syn_{body}$ |
| m00971[c] | 4-androstene-3,17-dione | C00280 | $\alpha Syn_{body}$ |
| m01064[c] | 5alpha-androstane-3,17-dione | C00674 | $\alpha Syn_{body}$ |
| m02131[c] | HMG-CoA | C00356 | $\alpha Syn_{body}$ |
| m00167[c] | (R)-mevalonate | C00418 | $\alpha Syn_{body}$ |
| m00165[c] | (R)-5-phosphomevalonate | C01107 | $\alpha Syn_{body}$ |
| m00164[c] | (R)-5-diphosphomevalonate | C01143 | $\alpha Syn_{body}$ |
| m01706[c] | dimethylallyl-PP | C00235 | $\alpha Syn_{body}$ |
| m01953[c] | geranyl-PP | C00341 | $\alpha Syn_{body}$ |
| m01789[c] | estrone 3-sulfate | C02538 | $\alpha Syn_{body}$ |
| m00986[c] | 4-hydroxy-17beta-estradiol | C14209 | $\alpha Syn_{body}$ |
| m00986[r] | 4-hydroxy-17beta-estradiol | C14209 | $\alpha Syn_{body}$ |
| m01430[c] | ceramide pool | C00195 | $\alpha Syn_{body}$ |

| | | | |
|---|---|---|---|
| m02684[c] | PC-LD pool | C00157 | $\alpha Syn_{body}$ |
| m01798[c] | ethanolamine-phosphate | C00346 | $\alpha Syn_{body}$ |
| m02600[c] | N-methylethanolamine-phosphate | C01210 | $\alpha Syn_{body}$ |
| m02739[c] | phosphodimethylethanolamine | C13482 | $\alpha Syn_{body}$ |
| m01627[c] | cysteamine | C01678 | $\alpha Syn_{body}$ |
| m02519[s] | Na+ | C01330 | $\alpha Syn_{body}$ |
| m02519[c] | Na+ | C01330 | $\alpha Syn_{body}$ |
| m01349[c] | apo-[ACP] | C03688 | $\alpha Syn_{body}$ |
| m02741[c] | phosphopantetheine | C01134 | $\alpha Syn_{body}$ |
| m02679[c] | pantetheine | C00831 | $\alpha Syn_{body}$ |
| m02680[c] | pantothenate | C00864 | $\alpha Syn_{body}$ |
| m00267[c] | 10-formyl-THF-glu(5) | | $\alpha Syn_{body}$ |
| m00267[l] | 10-formyl-THF-glu(5) | | $\alpha Syn_{body}$ |
| m00266[l] | 10-formyl-THF | C00234 | $\alpha Syn_{body}$ |
| m01974[l] | glutamate | C00025 | $\alpha Syn_{body}$ |
| m02692[s] | pentaglutamyl-folate(THF) | | $\alpha Syn_{body}$ |
| m02980[s] | THF | C00101 | $\alpha Syn_{body}$ |
| m01401[c] | biotin | C00120 | $\alpha Syn_{body}$ |
| m01401[s] | biotin | C00120 | $\alpha Syn_{body}$ |
| m02426[s] | lysine | C00047 | $\alpha Syn_{body}$ |
| m01155[c] | 6-[(1S,2R)-1,2-dihydroxy-3-triphosphooxypropyl]-7,8-dihydropterin | C04895 | $\alpha Syn_{body}$ |
| m03057[c] | triphosphate | C00536 | $\alpha Syn_{body}$ |
| m01704[c] | dihydroneopterin | C04874 | $\alpha Syn_{body}$ |
| m02049[m] | heme | C00032 | $\alpha Syn_{body}$ |
| m01631[m] | cytochrome-C | C00524 | $\alpha Syn_{body}$ |
| m02049[c] | heme | C00032 | $\alpha Syn_{body}$ |
| m03057[m] | triphosphate | C00536 | $\alpha Syn_{body}$ |
| m01600[m] | cobamide-coenzyme | C00194 | $\alpha Syn_{body}$ |
| m01442[c] | chloride | C00698 | $\alpha Syn_{body}$ |
| m02360[s] | leucine | C00123 | $\alpha Syn_{body}$ |
| m02834[s] | retinol | C00473 | $\alpha Syn_{body}$ |
| m00970[s] | 4-aminobutyrate | C00334 | $\alpha Syn_{body}$ |
| m02680[s] | pantothenate | C00864 | $\alpha Syn_{body}$ |
| m01383[s] | beta-alanine | C00099 | $\alpha Syn_{body}$ |
| m01442[s] | chloride | C00698 | $\alpha Syn_{body}$ |
| m01253[s] | acetoacetate | C00164 | $\alpha Syn_{body}$ |
| m02125[s] | histidine | C00135 | $\alpha Syn_{body}$ |
| m02471[s] | methionine | C00073 | $\alpha Syn_{body}$ |
| m01975[s] | glutamine | C00064 | $\alpha Syn_{body}$ |
| m03089[s] | tryptophan | C00078 | $\alpha Syn_{body}$ |
| m02724[s] | phenylalanine | C00079 | $\alpha Syn_{body}$ |
| m01369[s] | asparagine | C00152 | $\alpha Syn_{body}$ |

| | | | |
|---|---|---|---|
| m03135[s] | valine | C00183 | $\alpha Syn_{body}$ |
| m02184[s] | isoleucine | C00407 | $\alpha Syn_{body}$ |
| m02961[s] | taurine | C00245 | $\alpha Syn_{body}$ |
| m01833[s] | formate | C00058 | $\alpha Syn_{body}$ |
| m01789[s] | estrone 3-sulfate | C02538 | $\alpha Syn_{body}$ |
| m01659[s] | dehydroepiandrosterone sulfate | C04555 | $\alpha Syn_{body}$ |
| m02394[s] | lipoic acid | C00725 | $\alpha Syn_{body}$ |
| m02147[s] | hydroxide | C01328 | $\alpha Syn_{body}$ |
| m02049[s] | heme | C00032 | $\alpha Syn_{body}$ |
| m01158[c] | 6beta-hydroxytestosterone | C14497 | $\alpha Syn_{body}$ |
| m01158[s] | 6beta-hydroxytestosterone | C14497 | $\alpha Syn_{body}$ |
| m01338[s] | androsterone | C00523 | $\alpha Syn_{body}$ |
| m01361[s] | aquacob(III)alamin | C00992 | $\alpha Syn_{body}$ |
| m02969[s] | testosterone | C00535 | $\alpha Syn_{body}$ |
| m00986[s] | 4-hydroxy-17beta-estradiol | C14209 | $\alpha Syn_{body}$ |
| m01285[l] | ADP | C00008 | $\alpha Syn_{body}$ |
| m01371[l] | ATP | C00002 | $\alpha Syn_{body}$ |
| m01365[l] | arginine | C00062 | $\alpha Syn_{body}$ |
| m01369[l] | asparagine | C00152 | $\alpha Syn_{body}$ |
| m01628[l] | cysteine | C00097 | $\alpha Syn_{body}$ |
| m01975[l] | glutamine | C00064 | $\alpha Syn_{body}$ |
| m02125[l] | histidine | C00135 | $\alpha Syn_{body}$ |
| m02360[l] | leucine | C00123 | $\alpha Syn_{body}$ |
| m02426[l] | lysine | C00047 | $\alpha Syn_{body}$ |
| m02471[l] | methionine | C00073 | $\alpha Syn_{body}$ |
| m02724[l] | phenylalanine | C00079 | $\alpha Syn_{body}$ |
| m02770[l] | proline | C00148 | $\alpha Syn_{body}$ |
| m02896[l] | serine | C00065 | $\alpha Syn_{body}$ |
| m03089[l] | tryptophan | C00078 | $\alpha Syn_{body}$ |
| m03101[l] | tyrosine | C00082 | $\alpha Syn_{body}$ |
| m03135[l] | valine | C00183 | $\alpha Syn_{body}$ |
| m01307[l] | alanine | C00041 | $\alpha Syn_{body}$ |
| m02184[l] | isoleucine | C00407 | $\alpha Syn_{body}$ |
| m02993[l] | threonine | C00188 | $\alpha Syn_{body}$ |
| m03139[c] | vitamin A derivatives | | $\alpha Syn_{body}$ |
| m03143[c] | vitamin E derivatives | | $\alpha Syn_{body}$ |
| m01602[c] | cofactors and vitamins | | $\alpha Syn_{body}$ |
| m02040[c] | H2O | C00001 | $\alpha Syn_{body}$ |
| m02040[s] | H2O | C00001 | $\alpha Syn_{body}$ |
| m02877[c] | SAM | C00019 | $\alpha Syn_{body}$ |
| m02871[c] | SAH | C00021 | $\alpha Syn_{body}$ |
| m02039[c] | H+ | C00080 | $\alpha Syn_{body}$ |

| | | | |
|---|---|---|---|
| m02877[m] | SAM | C00019 | $\alpha Syn_{body}$ |
| m02871[m] | SAH | C00021 | $\alpha Syn_{body}$ |
| m02039[m] | H+ | C00080 | $\alpha Syn_{body}$ |
| m01371[c] | ATP | C00002 | $\alpha Syn_{body}$ |
| m01285[c] | ADP | C00008 | $\alpha Syn_{body}$ |
| m01626[c] | cys-gly | C01419 | $\alpha Syn_{body}$ |
| m01628[c] | cysteine | C00097 | $\alpha Syn_{body}$ |
| m01986[c] | glycine | C00037 | $\alpha Syn_{body}$ |
| m01626[s] | cys-gly | C01419 | $\alpha Syn_{body}$ |
| m01986[s] | glycine | C00037 | $\alpha Syn_{body}$ |
| m02040[m] | H2O | C00001 | $\alpha Syn_{body}$ |
| m01986[m] | glycine | C00037 | $\alpha Syn_{body}$ |
| m01983[c] | glycerol | C00116 | $\alpha Syn_{body}$ |
| m02914[c] | sn-glycerol-3-phosphate | C00093 | $\alpha Syn_{body}$ |
| m01371[m] | ATP | C00002 | $\alpha Syn_{body}$ |
| m03106[c] | UDP | C00015 | $\alpha Syn_{body}$ |
| m02039[r] | H+ | C00080 | $\alpha Syn_{body}$ |
| m02039[s] | H+ | C00080 | $\alpha Syn_{body}$ |
| m02041[c] | H2O2 | C00027 | $\alpha Syn_{body}$ |
| m01597[c] | CoA | C00010 | $\alpha Syn_{body}$ |
| m01597[m] | CoA | C00010 | $\alpha Syn_{body}$ |
| m02630[r] | O2 | C00007 | $\alpha Syn_{body}$ |
| km00013[r] | Red-NADPH-Hemoprotein-Reductases | | $\alpha Syn_{body}$ |
| km00014[r] | Ox-NADPH-Hemoprotein-Reductases | | $\alpha Syn_{body}$ |
| m02040[r] | H2O | C00001 | $\alpha Syn_{body}$ |
| m02555[c] | NADPH | C00005 | $\alpha Syn_{body}$ |
| m02554[c] | NADP+ | C00006 | $\alpha Syn_{body}$ |
| m02555[m] | NADPH | C00005 | $\alpha Syn_{body}$ |
| m02554[m] | NADP+ | C00006 | $\alpha Syn_{body}$ |
| m01974[c] | glutamate | C00025 | $\alpha Syn_{body}$ |
| m01306[c] | AKG | C00026 | $\alpha Syn_{body}$ |
| m01974[m] | glutamate | C00025 | $\alpha Syn_{body}$ |
| m01306[m] | AKG | C00026 | $\alpha Syn_{body}$ |
| m01948[c] | GDP | C00035 | $\alpha Syn_{body}$ |
| m02630[c] | O2 | C00007 | $\alpha Syn_{body}$ |
| m02555[r] | NADPH | C00005 | $\alpha Syn_{body}$ |
| m02554[r] | NADP+ | C00006 | $\alpha Syn_{body}$ |
| m01280[c] | adenosine | C00212 | $\alpha Syn_{body}$ |
| m02552[s] | NAD+ | C00003 | $\alpha Syn_{body}$ |
| km00013[c] | Red-NADPH-Hemoprotein-Reductases | | $\alpha Syn_{body}$ |
| km00014[c] | Ox-NADPH-Hemoprotein-Reductases | | $\alpha Syn_{body}$ |
| m02844[c] | ribose-1-phosphate | C00620 | $\alpha Syn_{body}$ |

| | | | |
|---|---|---|---|
| m02547[c] | N-acetylputrescine | C02714 | $\alpha Syn_{body}$ |
| m01252[c] | acetate | C00033 | $\alpha Syn_{body}$ |
| m02812[c] | putrescine | C00134 | $\alpha Syn_{body}$ |
| m02630[m] | O2 | C00007 | $\alpha Syn_{body}$ |
| m01334[c] | AMP | C00020 | $\alpha Syn_{body}$ |
| m02759[c] | PPi | C00013 | $\alpha Syn_{body}$ |
| m02552[c] | NAD+ | C00003 | $\alpha Syn_{body}$ |
| m02553[c] | NADH | C00004 | $\alpha Syn_{body}$ |
| m02552[m] | NAD+ | C00003 | $\alpha Syn_{body}$ |
| m02553[m] | NADH | C00004 | $\alpha Syn_{body}$ |
| m01368[c] | ascorbate | C00072 | $\alpha Syn_{body}$ |
| m01368[s] | ascorbate | C00072 | $\alpha Syn_{body}$ |
| m01596[c] | CO2 | C00011 | $\alpha Syn_{body}$ |
| m02040[l] | H2O | C00001 | $\alpha Syn_{body}$ |
| m01752[c] | dTMP | C00364 | $\alpha Syn_{body}$ |
| m00981[c] | 4-coumarate | C00811 | $\alpha Syn_{body}$ |
| m00982[c] | 4-coumaroyl-CoA | C00223 | $\alpha Syn_{body}$ |
| m01334[m] | AMP | C00020 | $\alpha Syn_{body}$ |
| m02759[m] | PPi | C00013 | $\alpha Syn_{body}$ |
| m01261[c] | acetyl-CoA | C00024 | $\alpha Syn_{body}$ |
| m02633[c] | OAA | C00036 | $\alpha Syn_{body}$ |
| m01596[m] | CO2 | C00011 | $\alpha Syn_{body}$ |
| m03101[c] | tyrosine | C00082 | $\alpha Syn_{body}$ |
| m01690[c] | DHAP | C00111 | $\alpha Syn_{body}$ |
| m01170[c] | 6-pyruvoyltetrahydropterin | C03684 | $\alpha Syn_{body}$ |
| m02026[c] | GSH | C00051 | $\alpha Syn_{body}$ |
| m02157[c] | hypotaurine | C00519 | $\alpha Syn_{body}$ |
| m03142[c] | vitamin D3 | C05443 | $\alpha Syn_{body}$ |
| m01415[c] | calcidiol | C01561 | $\alpha Syn_{body}$ |
| m01669[s] | deoxyguanosine | C00330 | $\alpha Syn_{body}$ |
| m01828[c] | FMN | C00061 | $\alpha Syn_{body}$ |
| m01802[c] | FAD | C00016 | $\alpha Syn_{body}$ |
| m01802[m] | FAD | C00016 | $\alpha Syn_{body}$ |
| m01285[s] | ADP | C00008 | $\alpha Syn_{body}$ |
| m02471[c] | methionine | C00073 | $\alpha Syn_{body}$ |
| m02961[c] | taurine | C00245 | $\alpha Syn_{body}$ |
| m01831[c] | formaldehyde | C00067 | $\alpha Syn_{body}$ |
| m02553[r] | NADH | C00004 | $\alpha Syn_{body}$ |
| m02552[r] | NAD+ | C00003 | $\alpha Syn_{body}$ |
| m01965[c] | glucose | C00031 | $\alpha Syn_{body}$ |
| m01587[c] | citrate | C00158 | $\alpha Syn_{body}$ |
| m01587[s] | citrate | C00158 | $\alpha Syn_{body}$ |

| | | | |
|---|---|---|---|
| m02631[s] | O2- | C00704 | $\alpha Syn_{body}$ |
| m02170[s] | inosine | C00294 | $\alpha Syn_{body}$ |
| m02896[s] | serine | C00065 | $\alpha Syn_{body}$ |
| m03089[c] | tryptophan | C00078 | $\alpha Syn_{body}$ |
| m01669[c] | deoxyguanosine | C00330 | $\alpha Syn_{body}$ |
| m02770[c] | proline | C00148 | $\alpha Syn_{body}$ |
| m00559[m] | 1-pyrroline-5-carboxylate | C03912 | $\alpha Syn_{body}$ |
| m02770[m] | proline | C00148 | $\alpha Syn_{body}$ |
| m01370[c] | aspartate | C00049 | $\alpha Syn_{body}$ |
| m01370[l] | aspartate | C00049 | $\alpha Syn_{body}$ |
| m01806[c] | farnesyl-PP | C00448 | $\alpha Syn_{body}$ |
| m01975[c] | glutamine | C00064 | $\alpha Syn_{body}$ |
| m01279[c] | adenine | C00147 | $\alpha Syn_{body}$ |
| m01686[c] | dGMP | C00362 | $\alpha Syn_{body}$ |
| m01307[c] | alanine | C00041 | $\alpha Syn_{body}$ |
| m03108[c] | UDP-glucose | C00029 | $\alpha Syn_{body}$ |
| m02834[c] | retinol | C00473 | $\alpha Syn_{body}$ |
| m02832[c] | retinal | C00376 | $\alpha Syn_{body}$ |
| m01249[c] | acetaldehyde | C00084 | $\alpha Syn_{body}$ |
| m01365[c] | arginine | C00062 | $\alpha Syn_{body}$ |
| m02426[c] | lysine | C00047 | $\alpha Syn_{body}$ |
| m02896[c] | serine | C00065 | $\alpha Syn_{body}$ |
| m01365[s] | arginine | C00062 | $\alpha Syn_{body}$ |
| m00184[c] | [ACP] | C00229 | $\alpha Syn_{body}$ |
| m01260[s] | acetylcholine | C01996 | $\alpha Syn_{body}$ |
| m01261[m] | acetyl-CoA | C00024 | $\alpha Syn_{body}$ |
| m02969[r] | testosterone | C00535 | $\alpha Syn_{body}$ |
| m01158[r] | 6beta-hydroxytestosterone | C14497 | $\alpha Syn_{body}$ |
| m02847[c] | RNA | C00046 | $\alpha Syn_{body}$ |
| m02682[c] | PAPS | C00053 | $\alpha Syn_{body}$ |
| m02681[c] | PAP | C00054 | $\alpha Syn_{body}$ |
| m02842[s] | riboflavin | C00255 | $\alpha Syn_{body}$ |
| m01736[s] | dopamine | C03758 | $\alpha Syn_{body}$ |
| m02724[c] | phenylalanine | C00079 | $\alpha Syn_{body}$ |
| m01127[c] | 5-oxoproline | C01879 | $\alpha Syn_{body}$ |
| m03102[m] | ubiquinol | C00390 | $\alpha Syn_{body}$ |
| m03103[m] | ubiquinone | C00399 | $\alpha Syn_{body}$ |
| m02348[c] | L-carnitine | C15025 | $\alpha Syn_{body}$ |
| m02993[s] | threonine | C00188 | $\alpha Syn_{body}$ |
| m02993[c] | threonine | C00188 | $\alpha Syn_{body}$ |
| m02842[c] | riboflavin | C00255 | $\alpha Syn_{body}$ |
| m02908[c] | SM pool | C00550 | $\alpha Syn_{body}$ |

| | | | |
|---|---|---|---|
| m02738[c] | phosphocholine | C00588 | $\alpha Syn_{body}$ |
| m01986[l] | glycine | C00037 | $\alpha Syn_{body}$ |
| m01424[c] | CDP | C00112 | $\alpha Syn_{body}$ |
| m02770[s] | proline | C00148 | $\alpha Syn_{body}$ |
| m01307[s] | alanine | C00041 | $\alpha Syn_{body}$ |
| m02578[c] | NH3 | C00014 | $\alpha Syn_{body}$ |
| m03135[c] | valine | C00183 | $\alpha Syn_{body}$ |
| m01833[c] | formate | C00058 | $\alpha Syn_{body}$ |
| m02751[c] | Pi | C00009 | $\alpha Syn_{body}$ |
| m03101[s] | tyrosine | C00082 | $\alpha Syn_{body}$ |
| m01803[c] | FADH2 | C01352 | $\alpha Syn_{body}$ |
| m01803[m] | FADH2 | C01352 | $\alpha Syn_{body}$ |
| m01840[c] | fructose | C02336 | $\alpha Syn_{body}$ |
| m02980[c] | THF | C00101 | $\alpha Syn_{body}$ |
| m02980[m] | THF | C00101 | $\alpha Syn_{body}$ |
| m02751[m] | Pi | C00009 | $\alpha Syn_{body}$ |
| m02751[s] | Pi | C00009 | $\alpha Syn_{body}$ |
| m01682[c] | D-glucitol | C00794 | $\alpha Syn_{body}$ |
| m01790[c] | estrone | C00468 | $\alpha Syn_{body}$ |
| m01790[r] | estrone | C00468 | $\alpha Syn_{body}$ |
| m01338[c] | androsterone | C00523 | $\alpha Syn_{body}$ |
| m02969[c] | testosterone | C00535 | $\alpha Syn_{body}$ |
| m02475[c] | methylglyoxal | C00546 | $\alpha Syn_{body}$ |
| m02771[c] | propane-1,2-diol | C00583 | $\alpha Syn_{body}$ |
| m02833[r] | retinoate | C00777 | $\alpha Syn_{body}$ |
| m01787[r] | estradiol-17beta | C00951 | $\alpha Syn_{body}$ |
| m01704[s] | dihydroneopterin | C04874 | $\alpha Syn_{body}$ |
| m02978[c] | tetrahydrobiopterin | C00272 | $\alpha Syn_{body}$ |
| m01361[c] | aquacob(III)alamin | C00992 | $\alpha Syn_{body}$ |
| m01599[c] | cob(II)alamin | C00541 | $\alpha Syn_{body}$ |
| m01599[m] | cob(II)alamin | C00541 | $\alpha Syn_{body}$ |
| m01660[c] | dehydroepiandrosterone | C01227 | $\alpha Syn_{body}$ |
| m01660[r] | dehydroepiandrosterone | C01227 | $\alpha Syn_{body}$ |
| km00877[r] | [Reduced NADPH---hemoprotein reductase] | C03024 | $\alpha Syn_{body}$ |
| km00878[r] | [Oxidized NADPH---hemoprotein reductase] | C03161 | $\alpha Syn_{body}$ |
| m00427[r] | 18-hydroxy-all-trans-retinoate | C16679 | $\alpha Syn_{body}$ |
| m01598[m] | cob(I)alamin | C00853 | $\alpha Syn_{body}$ |
| m01358[c] | apocytochrome-C | C02248 | $\alpha Syn_{body}$ |
| m01358[m] | apocytochrome-C | C02248 | $\alpha Syn_{body}$ |
| m01645[c] | dCTP | C00458 | $\alpha Syn_{body}$ |
| m02666[c] | oxidized thioredoxin | C00343 | $\alpha Syn_{body}$ |
| m01623[c] | CTP | C00063 | $\alpha Syn_{body}$ |

| | | | |
|---|---|---|---|
| m02990[c] | thioredoxin | C00342 | $\alpha Syn_{body}$ |
| m01643[c] | dCDP | C00705 | $\alpha Syn_{body}$ |
| m01700[c] | dihydrofolate | C00415 | $\alpha Syn_{body}$ |
| m01755[c] | dUMP | C00365 | $\alpha Syn_{body}$ |
| m01756[c] | dUTP | C00460 | $\alpha Syn_{body}$ |
| m01754[c] | dUDP | C01346 | $\alpha Syn_{body}$ |
| m01590[c] | CMP | C00055 | $\alpha Syn_{body}$ |
| m01630[c] | cytidine | C00475 | $\alpha Syn_{body}$ |
| m01752[m] | dTMP | C00364 | $\alpha Syn_{body}$ |
| m02996[m] | thymidine | C00214 | $\alpha Syn_{body}$ |
| m01285[m] | ADP | C00008 | $\alpha Syn_{body}$ |
| m01975[m] | glutamine | C00064 | $\alpha Syn_{body}$ |
| m02658[s] | ornithine | C00077 | $\alpha Syn_{body}$ |
| m02812[s] | putrescine | C00134 | $\alpha Syn_{body}$ |
| m01596[s] | CO2 | C00011 | $\alpha Syn_{body}$ |
| m02321[m] | L-2-amino-3-oxobutanoic acid | C03508 | $\alpha Syn_{body}$ |
| m01997[c] | glycolaldehyde | C00266 | $\alpha Syn_{body}$ |
| m02154[c] | hydroxypyruvate | C00168 | $\alpha Syn_{body}$ |
| m02763[c] | pregnenolone | C01953 | $\alpha Syn_{body}$ |
| m02769[c] | progesterone | C00410 | $\alpha Syn_{body}$ |
| m02685[c] | PE-LD pool | C00350 | $\alpha Syn_{body}$ |
| m02808[c] | PS-LD pool | C02737 | $\alpha Syn_{body}$ |
| m01797[c] | ethanolamine | C00189 | $\alpha Syn_{body}$ |
| m01808[c] | fatty acid-LD-PC pool | C00162 | $\alpha Syn_{body}$ |
| m00656[c] | 2-lysolecithin pool | C04230 | $\alpha Syn_{body}$ |
| m01260[c] | acetylcholine | C01996 | $\alpha Syn_{body}$ |
| m01513[c] | choline | C00114 | $\alpha Syn_{body}$ |
| m01513[s] | choline | C00114 | $\alpha Syn_{body}$ |
| m01252[s] | acetate | C00033 | $\alpha Syn_{body}$ |
| m02760[c] | pregn-5-ene-3,20-dione | | $\alpha Syn_{body}$ |
| m02794[r] | prostaglandin H2 | C00427 | $\alpha Syn_{body}$ |
| m02994[r] | thromboxane A2 | C02198 | $\alpha Syn_{body}$ |
| m01276[c] | acyl-CoA-LD-TG2 pool | | $\alpha Syn_{body}$ |
| m00223[c] | 1-(1-alkenyl)-sn-glycero-3-phosphoethanolamine | C04635 | $\alpha Syn_{body}$ |
| m02629[c] | O-1-alk-1-enyl-2-acyl-sn-glycero-3-phosphoethanolamine | C04756 | $\alpha Syn_{body}$ |
| m01814[c] | fatty acid-LD-TG2 pool | C00162 | $\alpha Syn_{body}$ |
| m00549[c] | 1-organyl-2-lyso-sn-glycero-3-phosphocholine | C04317 | $\alpha Syn_{body}$ |
| m00560[c] | 1-radyl-2-acyl-sn-glycero-3-phosphocholine | C05212 | $\alpha Syn_{body}$ |
| m01445[s] | cholate | C00695 | $\alpha Syn_{body}$ |
| m01445[c] | cholate | C00695 | $\alpha Syn_{body}$ |
| m01988[c] | glycocholate | C01921 | $\alpha Syn_{body}$ |
| m01988[s] | glycocholate | C01921 | $\alpha Syn_{body}$ |

| | | | |
|---|---|---|---|
| m02357[c] | lepidimoide | C08241 | $\alpha Syn_{body}$ |
| m02357[s] | lepidimoide | C08241 | $\alpha Syn_{body}$ |
| m01987[c] | glycochenodeoxycholate | C05466 | $\alpha Syn_{body}$ |
| m01987[s] | glycochenodeoxycholate | C05466 | $\alpha Syn_{body}$ |
| m02963[c] | taurocholate | C05122 | $\alpha Syn_{body}$ |
| m02963[s] | taurocholate | C05122 | $\alpha Syn_{body}$ |
| m02962[c] | taurochenodeoxycholate | C05465 | $\alpha Syn_{body}$ |
| m02962[s] | taurochenodeoxycholate | C05465 | $\alpha Syn_{body}$ |
| m00526[c] | 1D-myo-inositol-1,4-bisphosphate | C01220 | $\alpha Syn_{body}$ |
| m00530[c] | 1D-myo-inositol-4-phosphate | C03546 | $\alpha Syn_{body}$ |
| m02171[c] | inositol | C00137 | $\alpha Syn_{body}$ |
| m01164[c] | 6-lactoyl-5,6,7,8-tetrahydropterin | C04244 | $\alpha Syn_{body}$ |
| m01698[c] | dihydrobiopterin | C00268 | $\alpha Syn_{body}$ |
| m01115[c] | 5-methyl-THF | C00440 | $\alpha Syn_{body}$ |
| m00003[s] | (10Z)-heptadecenoic acid | | $\alpha Syn_{body}$ |
| m00008[s] | (11Z,14Z)-eicosadienoic acid | C16525 | $\alpha Syn_{body}$ |
| m00010[s] | (11Z,14Z,17Z)-eicosatrienoic acid | C16522 | $\alpha Syn_{body}$ |
| m00017[s] | (13Z)-eicosenoic acid | | $\alpha Syn_{body}$ |
| m00019[s] | (13Z)-octadecenoic acid | | $\alpha Syn_{body}$ |
| m00021[s] | (13Z,16Z)-docosadienoic acid | C16533 | $\alpha Syn_{body}$ |
| m00094[s] | (4Z,7Z,10Z,13Z,16Z)-DPA | | $\alpha Syn_{body}$ |
| m00104[s] | (6Z,9Z)-octadecadienoic acid | | $\alpha Syn_{body}$ |
| m00111[s] | (6Z,9Z,12Z,15Z,18Z)-TPA | | $\alpha Syn_{body}$ |
| m00114[s] | (6Z,9Z,12Z,15Z,18Z,21Z)-THA | | $\alpha Syn_{body}$ |
| m00115[s] | (7Z)-octadecenoic acid | | $\alpha Syn_{body}$ |
| m00117[s] | (7Z)-tetradecenoic acid | | $\alpha Syn_{body}$ |
| m00128[s] | (9E)-tetradecenoic acid | | $\alpha Syn_{body}$ |
| m00132[s] | (9Z,12Z,15Z,18Z)-TTA | | $\alpha Syn_{body}$ |
| m00135[s] | (9Z,12Z,15Z,18Z,21Z)-TPA | | $\alpha Syn_{body}$ |
| m00260[s] | 10,13,16,19-docosatetraenoic acid | | $\alpha Syn_{body}$ |
| m00265[s] | 10,13,16-docosatriynoic acid | | $\alpha Syn_{body}$ |
| m00315[s] | 12,15,18,21-tetracosatetraenoic acid | | $\alpha Syn_{body}$ |
| m00341[s] | 13,16,19-docosatrienoic acid | C16534 | $\alpha Syn_{body}$ |
| m01197[s] | 7-palmitoleic acid | | $\alpha Syn_{body}$ |
| m01207[s] | 8,11-eicosadienoic acid | | $\alpha Syn_{body}$ |
| m01235[s] | 9-eicosenoic acid | | $\alpha Syn_{body}$ |
| m01238[s] | 9-heptadecylenic acid | C16536 | $\alpha Syn_{body}$ |
| m01291[s] | adrenic acid | C16527 | $\alpha Syn_{body}$ |
| m01373[s] | behenic acid | C08281 | $\alpha Syn_{body}$ |
| m01432[s] | cerotic acid | | $\alpha Syn_{body}$ |
| m01582[s] | cis-cetoleic acid | | $\alpha Syn_{body}$ |
| m01583[s] | cis-erucic acid | C08316 | $\alpha Syn_{body}$ |

| | | | |
|---|---|---|---|
| m01584[s] | cis-gondoic acid | C16526 | αSyn$_{body}$ |
| m01585[s] | cis-vaccenic acid | C08367 | αSyn$_{body}$ |
| m01689[s] | DHA | C06429 | αSyn$_{body}$ |
| m01696[s] | dihomo-gamma-linolenate | C03242 | αSyn$_{body}$ |
| m01741[s] | DPA | C16513 | αSyn$_{body}$ |
| m01771[s] | eicosanoate | C06425 | αSyn$_{body}$ |
| m01778[s] | elaidate | C00712 | αSyn$_{body}$ |
| m01784[s] | EPA | C06428 | αSyn$_{body}$ |
| m01932[s] | gamma-linolenate | C06426 | αSyn$_{body}$ |
| m02053[s] | henicosanoic acid | | αSyn$_{body}$ |
| m02344[s] | lauric acid | C02679 | αSyn$_{body}$ |
| m02385[s] | lignocerate | C08320 | αSyn$_{body}$ |
| m02456[s] | margaric acid | | αSyn$_{body}$ |
| m02457[s] | mead acid | | αSyn$_{body}$ |
| m02494[s] | myristic acid | C06424 | αSyn$_{body}$ |
| m02564[s] | nervonic acid | C08323 | αSyn$_{body}$ |
| m02613[s] | nonadecylic acid | C16535 | αSyn$_{body}$ |
| m02648[s] | omega-3-arachidonic acid | | αSyn$_{body}$ |
| m02690[s] | pentadecylic acid | C16537 | αSyn$_{body}$ |
| m02745[s] | physeteric acid | | αSyn$_{body}$ |
| m02938[s] | stearate | C01530 | αSyn$_{body}$ |
| m02939[s] | stearidonic acid | C16300 | αSyn$_{body}$ |
| m03045[s] | tricosanoic acid | | αSyn$_{body}$ |
| m03051[s] | tridecylic acid | C17076 | αSyn$_{body}$ |
| m03153[s] | ximenic acid | | αSyn$_{body}$ |
| m02560[s] | NEFA blood pool in | | αSyn$_{body}$ |
| m02646[s] | oleate | C00712 | αSyn$_{body}$ |
| m02674[s] | palmitate | C00249 | αSyn$_{body}$ |
| m01362[s] | arachidonate | C00219 | αSyn$_{body}$ |
| m02387[s] | linoleate | C01595 | αSyn$_{body}$ |
| m02389[s] | linolenate | C06427 | αSyn$_{body}$ |
| m02675[s] | palmitolate | C08362 | αSyn$_{body}$ |
| m02646[c] | oleate | C00712 | αSyn$_{body}$ |
| m02046[c] | HCO3- | C00288 | αSyn$_{body}$ |
| m01980[c] | glutathionyl-leukotriene C4 | | αSyn$_{body}$ |
| m01980[s] | glutathionyl-leukotriene C4 | | αSyn$_{body}$ |
| m02897[s] | serotonin | C00780 | αSyn$_{body}$ |
| m02897[c] | serotonin | C00780 | αSyn$_{body}$ |
| m02046[s] | HCO3- | C00288 | αSyn$_{body}$ |
| m01280[s] | adenosine | C00212 | αSyn$_{body}$ |
| m02843[c] | ribose | C00121 | αSyn$_{body}$ |
| m02843[s] | ribose | C00121 | αSyn$_{body}$ |

| | | | |
|---|---|---|---|
| m02403[s] | L-lactate | C00186 | $\alpha Syn_{body}$ |
| m02833[s] | retinoate | C00777 | $\alpha Syn_{body}$ |
| m02403[c] | L-lactate | C00186 | $\alpha Syn_{body}$ |
| m02833[c] | retinoate | C00777 | $\alpha Syn_{body}$ |
| m03052[s] | triiodothyronine | C02465 | $\alpha Syn_{body}$ |
| m03052[c] | triiodothyronine | C02465 | $\alpha Syn_{body}$ |
| m02998[c] | thyroxine | C01829 | $\alpha Syn_{body}$ |
| m02998[s] | thyroxine | C01829 | $\alpha Syn_{body}$ |
| m02026[s] | GSH | C00051 | $\alpha Syn_{body}$ |
| m01397[c] | bilirubin-bisglucuronoside | C05787 | $\alpha Syn_{body}$ |
| m01397[s] | bilirubin-bisglucuronoside | C05787 | $\alpha Syn_{body}$ |
| m02779[c] | prostaglandin B2 | C05954 | $\alpha Syn_{body}$ |
| m02779[s] | prostaglandin B2 | C05954 | $\alpha Syn_{body}$ |
| m02900[c] | S-glutathionyl-2-4-dinitrobenzene | | $\alpha Syn_{body}$ |
| m02900[s] | S-glutathionyl-2-4-dinitrobenzene | | $\alpha Syn_{body}$ |
| m02789[c] | prostaglandin F2alpha | C00639 | $\alpha Syn_{body}$ |
| m02789[s] | prostaglandin F2alpha | C00639 | $\alpha Syn_{body}$ |
| m02901[c] | S-glutathionyl-ethacrynic acid | | $\alpha Syn_{body}$ |
| m02901[s] | S-glutathionyl-ethacrynic acid | | $\alpha Syn_{body}$ |
| m02792[s] | prostaglandin G2 | C05956 | $\alpha Syn_{body}$ |
| m02792[c] | prostaglandin G2 | C05956 | $\alpha Syn_{body}$ |
| m02781[c] | prostaglandin C2 | C05955 | $\alpha Syn_{body}$ |
| m02781[s] | prostaglandin C2 | C05955 | $\alpha Syn_{body}$ |
| m02839[c] | reverse triiodthyronine | C07639 | $\alpha Syn_{body}$ |
| m02839[s] | reverse triiodthyronine | C07639 | $\alpha Syn_{body}$ |
| m02777[c] | prostaglandin A2 | C05953 | $\alpha Syn_{body}$ |
| m02777[s] | prostaglandin A2 | C05953 | $\alpha Syn_{body}$ |
| m02784[s] | prostaglandin D3 | C13802 | $\alpha Syn_{body}$ |
| m02784[c] | prostaglandin D3 | C13802 | $\alpha Syn_{body}$ |
| m02794[c] | prostaglandin H2 | C00427 | $\alpha Syn_{body}$ |
| m02794[s] | prostaglandin H2 | C00427 | $\alpha Syn_{body}$ |
| m01256[s] | acetone | C00207 | $\alpha Syn_{body}$ |
| m02994[c] | thromboxane A2 | C02198 | $\alpha Syn_{body}$ |
| m02994[s] | thromboxane A2 | C02198 | $\alpha Syn_{body}$ |
| m01306[s] | AKG | C00026 | $\alpha Syn_{body}$ |
| m02993[m] | threonine | C00188 | $\alpha Syn_{body}$ |
| m01645[m] | dCTP | C00458 | $\alpha Syn_{body}$ |
| m01643[m] | dCDP | C00705 | $\alpha Syn_{body}$ |
| m03148[c] | xanthine | C00385 | $\alpha Syn_{body}$ |
| m03148[p] | xanthine | C00385 | $\alpha Syn_{body}$ |
| m02814[s] | pyridoxal-phosphate | C00018 | $\alpha Syn_{body}$ |
| m02813[s] | pyridoxal | C00250 | $\alpha Syn_{body}$ |

| | | | |
|---|---|---|---|
| m01450[m] | cholesterol | C00187 | $\alpha Syn_{body}$ |
| m02827[m] | reduced adrenal ferredoxin | C00662 | $\alpha Syn_{body}$ |
| m00592[m] | 20-hydroxycholesterol | C05500 | $\alpha Syn_{body}$ |
| m02663[m] | oxidized adrenal ferredoxin | C00667 | $\alpha Syn_{body}$ |
| m00579[m] | 20alpha,22beta dihydroxycholesterol | C05501 | $\alpha Syn_{body}$ |
