## Supplemental Table 3 for "Comparative Proteomic Analysis of Environmental and Genetic Models of Parkinson’s Disease Highlights the Role of Purine Metabolism"

**Table S3 |** Significantly enriched KEGG pathways (FDR < 0.05) of the genes in the differential *Drosophila* reactions by the PANGEA tool for paraquat treatment or α-synuclein expression.

| **Pathway ID** | **Pathway Term** | **Gene Set Size** | **Count Overlap Gene** | **Percentage Genes in Gene Set** | **Benjamini & Hochberg** | **Overlapping Gene IDs** | **PD Group** |
|---|---|---|---|---|---|---|---|
| path:map01100 | Metabolic pathways | 1153 | 199 | 51.554 | 2.92E-24 | FBgn0029648,FBgn0000479,FBgn0262738,FBgn028622 | $PQ_{head}$ |
| path:map00230 | Purine metabolism | 92 | 39 | 10.104 | 2.92E-24 | FBgn0000479,FBgn0259171,FBgn0030573,FBgn00033 | $PQ_{head}$ |
| path:map00970 | Aminoacyl-tRNA biosynthesis | 64 | 34 | 8.808 | 2.92E-24 | FBgn0030007,FBgn0027084,FBgn0027093,FBgn002708 | $PQ_{head}$ |
| path:map01232 | Nucleotide metabolism | 60 | 28 | 7.254 | 2.92E-24 | FBgn0030573,FBgn0052549,FBgn0024947,FBgn003148 | $PQ_{head}$ |
| path:map00240 | Pyrimidine metabolism | 38 | 24 | 6.218 | 2.92E-24 | FBgn0086450,FBgn0030573,FBgn0052549,FBgn003126 | $PQ_{head}$ |
| path:map01212 | Fatty acid metabolism | 53 | 22 | 5.699 | 1.46E-19 | FBgn0029648,FBgn0029975,FBgn0283427,FBgn002847 | $PQ_{head}$ |
| path:map00071 | Fatty acid degradation | 33 | 18 | 4.663 | 8.84E-19 | FBgn0028479,FBgn0040064,FBgn0032160,FBgn028672 | $PQ_{head}$ |
| path:map01240 | Biosynthesis of cofactors | 139 | 28 | 7.254 | 1.73E-15 | FBgn0014032,FBgn0030407,FBgn0030431,FBgn003046 | $PQ_{head}$ |
| path:map01200 | Carbon metabolism | 121 | 26 | 6.736 | 3.73E-15 | FBgn0286222,FBgn0027291,FBgn0001125,FBgn003206 | $PQ_{head}$ |
| path:map00620 | Pyruvate metabolism | 49 | 17 | 4.404 | 9.54E-14 | FBgn0286222,FBgn0033856,FBgn0034127,FBgn003435 | $PQ_{head}$ |
| md:M00089 | Triacylglycerol biosynthesis | 16 | 11 | 2.85 | 3.19E-13 | FBgn0030421,FBgn0263593,FBgn0016078,FBgn003662 | $PQ_{head}$ |
| path:map00564 | Glycerophospholipid metabolism | 63 | 18 | 4.663 | 6.02E-13 | FBgn0030421,FBgn0001128,FBgn0028848,FBgn003292 | $PQ_{head}$ |
| path:map00561 | Glycerolipid metabolism | 42 | 15 | 3.886 | 1.80E-12 | FBgn0030421,FBgn0033214,FBgn0033215,FBgn003321 | $PQ_{head}$ |
| path:map00280 | Valine, leucine and isoleucine degradation | 33 | 13 | 3.368 | 1.63E-11 | FBgn0030482,FBgn0021765,FBgn0031877,FBgn002847 | $PQ_{head}$ |
| md:M00938 | Pyrimidine deoxyribonucleotide biosynthesis, UDP => dTTP | 9 | 8 | 2.073 | 2.71E-11 | FBgn0030573,FBgn0011703,FBgn0250837,FBgn00117 | $PQ_{head}$ |
| path:map00270 | Cysteine and methionine metabolism | 36 | 13 | 3.368 | 5.32E-11 | FBgn0030482,FBgn0030558,FBgn0014455,FBgn003088 | $PQ_{head}$ |
| path:map00600 | Sphingolipid metabolism | 30 | 12 | 3.109 | 7.94E-11 | FBgn0030300,FBgn0002524,FBgn0016078,FBgn008653 | $PQ_{head}$ |
| path:map00900 | Terpenoid backbone biosynthesis | 27 | 11 | 2.85 | 4.43E-10 | FBgn0030683,FBgn0032811,FBgn0025373,FBgn00352 | $PQ_{head}$ |
| md:M00095 | C5 isoprenoid biosynthesis, mevalonate pathway | 8 | 7 | 1.813 | 7.00E-10 | FBgn0030683,FBgn0032811,FBgn0035203,FBgn003887 | $PQ_{head}$ |
| md:M00959 | Guanine ribonucleotide degradation, GMP => Urate | 13 | 8 | 2.073 | 2.51E-09 | FBgn0052549,FBgn0033538,FBgn0034225,FBgn003489 | $PQ_{head}$ |
| path:map00061 | Fatty acid biosynthesis | 13 | 8 | 2.073 | 2.51E-09 | FBgn0029648,FBgn0283427,FBgn0286723,FBgn003669 | $PQ_{head}$ |
| md:M00053 | Deoxyribonucleotide biosynthesis, ADP/GDP/CDP/UDP => dATP/ | 6 | 6 | 1.554 | 2.85E-09 | FBgn0030573,FBgn0011703,FBgn0011704,FBgn002899 | $PQ_{head}$ |
| path:map00983 | Drug metabolism - other enzymes | 96 | 17 | 4.404 | 6.06E-09 | FBgn0086450,FBgn0030573,FBgn0031489,FBgn003166 | $PQ_{head}$ |
| path:map00250 | Alanine, aspartate and glutamate metabolism | 29 | 10 | 2.591 | 1.76E-08 | FBgn0030653,FBgn0001125,FBgn0033543,FBgn000112 | $PQ_{head}$ |
| md:M00128 | Ubiquinone biosynthesis, eukaryotes, 4-hydroxybenzoate + polypre | 5 | 5 | 1.295 | 8.50E-08 | FBgn0030460,FBgn0031713,FBgn0032922,FBgn003757 | $PQ_{head}$ |
| md:M00169 | CAM (Crassulacean acid metabolism), light | 5 | 5 | 1.295 | 8.50E-08 | FBgn0034127,FBgn0029155,FBgn0002719,FBgn002915 | $PQ_{head}$ |
| md:M00172 | C4-dicarboxylic acid cycle, NADP - malic enzyme type | 5 | 5 | 1.295 | 8.50E-08 | FBgn0034127,FBgn0029155,FBgn0002719,FBgn002915 | $PQ_{head}$ |
| path:map00130 | Ubiquinone and other terpenoid-quinone biosynthesis | 9 | 6 | 1.554 | 1.69E-07 | FBgn0030460,FBgn0030558,FBgn0031713,FBgn003292 | $PQ_{head}$ |
| path:map00430 | Taurine and hypotaurine metabolism | 9 | 6 | 1.554 | 1.69E-07 | FBgn0030361,FBgn0030796,FBgn0030932,FBgn000451 | $PQ_{head}$ |
| path:map01230 | Biosynthesis of amino acids | 68 | 13 | 3.368 | 1.73E-07 | FBgn0030482,FBgn0027291,FBgn0031148,FBgn000112 | $PQ_{head}$ |
| path:map00650 | Butanoate metabolism | 15 | 7 | 1.813 | 2.92E-07 | FBgn0031877,FBgn0028479,FBgn0033879,FBgn00352 | $PQ_{head}$ |
| path:map01210 | 2-Oxocarboxylic acid metabolism | 22 | 8 | 2.073 | 3.29E-07 | FBgn0030482,FBgn0027291,FBgn0001125,FBgn000112 | $PQ_{head}$ |
| md:M00849 | C5 isoprenoid biosynthesis, mevalonate pathway, archaea | 6 | 5 | 1.295 | 4.08E-07 | FBgn0035203,FBgn0038876,FBgn0263782,FBgn001061 | $PQ_{head}$ |
| path:map00062 | Fatty acid elongation | 16 | 7 | 1.813 | 4.62E-07 | FBgn0029975,FBgn0028479,FBgn0040064,FBgn003387 | $PQ_{head}$ |
| path:map00020 | Citrate cycle (TCA cycle) | 43 | 10 | 2.591 | 8.19E-07 | FBgn0286222,FBgn0027291,FBgn0020236,FBgn003435 | $PQ_{head}$ |
| path:map00410 | beta-Alanine metabolism | 25 | 8 | 2.073 | 9.19E-07 | FBgn0086450,FBgn0028479,FBgn0033879,FBgn000451 | $PQ_{head}$ |
| path:map00010 | Glycolysis / Gluconeogenesis | 55 | 11 | 2.85 | 9.86E-07 | FBgn0033856,FBgn0034356,FBgn0003067,FBgn003676 | $PQ_{head}$ |

| | | | | | | | |
|---|---|---|---|---|---|---|---|
| md:M00099 | Sphingosine biosynthesis | 7 | 5 | 1.295 | 1.21E-06 | FBgn0002524,FBgn0086532,FBgn0039304,FBgn003977 | $PQ_{head}$ |
| md:M00087 | beta-Oxidation | 12 | 6 | 1.554 | 1.29E-06 | FBgn0028479,FBgn0040064,FBgn0033879,FBgn003443 | $PQ_{head}$ |
| path:map04146 | peroxisome | 96 | 14 | 3.627 | 1.34E-06 | FBgn0031860,FBgn0031877,FBgn0032061,FBgn002847 | $PQ_{head}$ |
| md:M00958 | Adenine ribonucleotide degradation, AMP => Urate | 19 | 7 | 1.813 | 1.54E-06 | FBgn0052549,FBgn0033538,FBgn0034225,FBgn003489 | $PQ_{head}$ |
| path:map00260 | Glycine, serine and threonine metabolism | 27 | 8 | 2.073 | 1.54E-06 | FBgn0031148,FBgn0031860,FBgn0032287,FBgn003620 | $PQ_{head}$ |
| md:M00086 | beta-Oxidation, acyl-CoA synthesis | 4 | 4 | 1.036 | 1.95E-06 | FBgn0286723,FBgn0036821,FBgn0027348,FBgn026312 | $PQ_{head}$ |
| path:map00740 | Riboflavin metabolism | 8 | 5 | 1.295 | 2.73E-06 | FBgn0030431,FBgn0032522,FBgn0038912,FBgn000003 | $PQ_{head}$ |
| path:map00380 | Tryptophan metabolism | 21 | 7 | 1.813 | 3.15E-06 | FBgn0032061,FBgn0028479,FBgn0033879,FBgn003520 | $PQ_{head}$ |
| md:M00532 | Photorespiration | 14 | 6 | 1.554 | 3.31E-06 | FBgn0032061,FBgn0032287,FBgn0036762,FBgn000026 | $PQ_{head}$ |
| path:map00770 | Pantothenate and CoA biosynthesis | 14 | 6 | 1.554 | 3.31E-06 | FBgn0086450,FBgn0030482,FBgn0023023,FBgn003751 | $PQ_{head}$ |
| path:map00630 | Glyoxylate and dicarboxylate metabolism | 33 | 8 | 2.073 | 7.29E-06 | FBgn0032061,FBgn0032287,FBgn0035203,FBgn003676 | $PQ_{head}$ |
| md:M00035 | Methionine degradation | 5 | 4 | 1.036 | 8.20E-06 | FBgn0014455,FBgn0031148,FBgn0035371,FBgn001501 | $PQ_{head}$ |
| md:M00085 | Fatty acid elongation in mitochondria | 5 | 4 | 1.036 | 8.20E-06 | FBgn0028479,FBgn0040064,FBgn0033879,FBgn002535 | $PQ_{head}$ |
| path:map00565 | Ether lipid metabolism | 25 | 7 | 1.813 | 1.04E-05 | FBgn0032923,FBgn0286511,FBgn0016078,FBgn003716 | $PQ_{head}$ |
| path:map00760 | Nicotinate and nicotinamide metabolism | 18 | 6 | 1.554 | 1.68E-05 | FBgn0052549,FBgn0033538,FBgn0034225,FBgn003489 | $PQ_{head}$ |
| md:M00036 | Leucine degradation, leucine => acetoacetate + acetyl-CoA | 11 | 5 | 1.295 | 1.70E-05 | FBgn0030482,FBgn0031877,FBgn0033761,FBgn003581 | $PQ_{head}$ |
| path:map00450 | Selenocompound metabolism | 11 | 5 | 1.295 | 1.70E-05 | FBgn0020653,FBgn0034401,FBgn0020389,FBgn003717 | $PQ_{head}$ |
| path:map00640 | Propanoate metabolism | 27 | 7 | 1.813 | 1.70E-05 | FBgn0028479,FBgn0033856,FBgn0033879,FBgn003676 | $PQ_{head}$ |
| md:M00052 | Pyrimidine ribonucleotide biosynthesis, UMP => UDP/UTP,CDP/C | 6 | 4 | 1.036 | 2.11E-05 | FBgn0030573,FBgn0028997,FBgn0039809,FBgn000015 | $PQ_{head}$ |
| md:M00088 | Ketone body biosynthesis, acetyl-CoA => acetoacetate/3-hydroxybu | 6 | 4 | 1.036 | 2.11E-05 | FBgn0031877,FBgn0035203,FBgn0262112,FBgn001061 | $PQ_{head}$ |
| path:map00480 | Glutathione metabolism | 79 | 11 | 2.85 | 2.62E-05 | FBgn0030361,FBgn0030411,FBgn0030796,FBgn003088 | $PQ_{head}$ |
| md:M00010 | Citrate cycle, first carbon oxidation, oxaloacetate => 2-oxoglutarate | 12 | 5 | 1.295 | 2.62E-05 | FBgn0027291,FBgn0035005,FBgn0038922,FBgn003935 | $PQ_{head}$ |
| md:M00050 | Guanine ribonucleotide biosynthesis, IMP => GDP,GTP | 7 | 4 | 1.036 | 4.58E-05 | FBgn0030573,FBgn0028997,FBgn0039809,FBgn000015 | $PQ_{head}$ |
| md:M00046 | Pyrimidine degradation, uracil => beta-alanine, thymine => 3-amin | 3 | 3 | 0.777 | 4.93E-05 | FBgn0086450,FBgn0023023,FBgn0037513 | $PQ_{head}$ |
| md:M00118 | Glutathione biosynthesis, glutamate => glutathione | 3 | 3 | 0.777 | 4.93E-05 | FBgn0030882,FBgn0040319,FBgn0046114 | $PQ_{head}$ |
| path:map00350 | Tyrosine metabolism | 22 | 6 | 1.554 | 5.09E-05 | FBgn0030558,FBgn0001125,FBgn0001124,FBgn001176 | $PQ_{head}$ |
| path:map00360 | Phenylalanine metabolism | 8 | 4 | 1.036 | 8.40E-05 | FBgn0030558,FBgn0001125,FBgn0001124,FBgn001054 | $PQ_{head}$ |
| md:M00009 | Citrate cycle (TCA cycle, Krebs cycle) | 35 | 7 | 1.813 | 9.02E-05 | FBgn0286222,FBgn0027291,FBgn0035005,FBgn003676 | $PQ_{head}$ |
| path:map00400 | Phenylalanine, tyrosine and tryptophan biosynthesis | 4 | 3 | 0.777 | 1.82E-04 | FBgn0030558,FBgn0001125,FBgn0001124 | $PQ_{head}$ |
| path:map00330 | Arginine and proline metabolism | 53 | 8 | 2.073 | 2.06E-04 | FBgn0001125,FBgn0031860,FBgn0001124,FBgn003640 | $PQ_{head}$ |
| md:M00049 | Adenine ribonucleotide biosynthesis, IMP => ADP,ATP | 10 | 4 | 1.036 | 2.27E-04 | FBgn0030573,FBgn0028997,FBgn0039809,FBgn000015 | $PQ_{head}$ |
| md:M00082 | Fatty acid biosynthesis, initiation | 5 | 3 | 0.777 | 4.19E-04 | FBgn0283427,FBgn0036691,FBgn0042627 | $PQ_{head}$ |
| md:M00094 | Ceramide biosynthesis | 5 | 3 | 0.777 | 4.19E-04 | FBgn0002524,FBgn0086532,FBgn0039304 | $PQ_{head}$ |
| path:map01040 | Biosynthesis of unsaturated fatty acids | 25 | 5 | 1.295 | 1.09E-03 | FBgn0029975,FBgn0035471,FBgn0260960,FBgn004304 | $PQ_{head}$ |
| path:map02010 | ABC transporters | 38 | 6 | 1.554 | 1.13E-03 | FBgn0031170,FBgn0028539,FBgn0034493,FBgn003674 | $PQ_{head}$ |
| path:map00590 | Arachidonic acid metabolism | 15 | 4 | 1.036 | 1.23E-03 | FBgn0030361,FBgn0030796,FBgn0030932,FBgn002583 | $PQ_{head}$ |
| path:map04213 | Longevity regulating pathway - multiple species | 55 | 7 | 1.813 | 1.48E-03 | FBgn0003301,FBgn0022710,FBgn0032061,FBgn002341 | $PQ_{head}$ |
| md:M00170 | C4-dicarboxylic acid cycle, phosphoenolpyruvate carboxykinase typ | 2 | 2 | 0.518 | 1.48E-03 | FBgn0001125,FBgn0001124 | $PQ_{head}$ |
| md:M00032 | Lysine degradation, lysine => saccharopine => acetoacetyl-CoA | 8 | 3 | 0.777 | 2.03E-03 | FBgn0028479,FBgn0033879,FBgn0036762 | $PQ_{head}$ |
| path:map00220 | Arginine biosynthesis | 18 | 4 | 1.036 | 2.45E-03 | FBgn0001125,FBgn0001124,FBgn0039071,FBgn000109 | $PQ_{head}$ |

| | | | | | | | |
|---|---|---|---|---|---|---|---|
| md:M00415 | Fatty acid elongation in endoplasmic reticulum | 9 | 3 | 0.777 | 2.87E-03 | FBgn0029975,FBgn0035471,FBgn0260960 | PQ$_{head}$ |
| path:map00920 | Sulfur metabolism | 9 | 3 | 0.777 | 2.87E-03 | FBgn0030465,FBgn0020389,FBgn0039698 | PQ$_{head}$ |
| md:M00015 | Proline biosynthesis, glutamate => proline | 3 | 2 | 0.518 | 3.92E-03 | FBgn0038516,FBgn0015781 | PQ$_{head}$ |
| md:M00100 | Sphingosine degradation | 3 | 2 | 0.518 | 3.92E-03 | FBgn0030300,FBgn0052484 | PQ$_{head}$ |
| md:M00911 | Riboflavin biosynthesis, fungi, GTP => riboflavin/FMN/FAD | 3 | 2 | 0.518 | 3.92E-03 | FBgn0030431,FBgn0032522 | PQ$_{head}$ |
| path:map00470 | D-Amino acid metabolism | 3 | 2 | 0.518 | 3.92E-03 | FBgn0031860,FBgn0033543 | PQ$_{head}$ |
| path:map00830 | Retinol metabolism | 34 | 5 | 1.295 | 3.92E-03 | FBgn0031630,FBgn0032910,FBgn0011768,FBgn000479 | PQ$_{head}$ |
| path:map00310 | Lysine degradation | 35 | 5 | 1.295 | 4.42E-03 | FBgn0028479,FBgn0033879,FBgn0035203,FBgn003676 | PQ$_{head}$ |
| md:M00130 | Inositol phosphate metabolism, PI=> PIP2 => Ins(1,4,5)P3 => Ins(1 | 11 | 3 | 0.777 | 4.97E-03 | FBgn0262738,FBgn0003416,FBgn0004611 | PQ$_{head}$ |
| md:M00083 | Fatty acid biosynthesis, elongation | 4 | 2 | 0.518 | 7.31E-03 | FBgn0283427,FBgn0042627 | PQ$_{head}$ |
| md:M00842 | Tetrahydrobiopterin biosynthesis, GTP => BH4 | 4 | 2 | 0.518 | 7.31E-03 | FBgn0014032,FBgn0003141 | PQ$_{head}$ |
| path:map00790 | Folate biosynthesis | 40 | 5 | 1.295 | 7.65E-03 | FBgn0014032,FBgn0030407,FBgn0003141,FBgn003596 | PQ$_{head}$ |
| path:map00562 | Inositol phosphate metabolism | 46 | 5 | 1.295 | 1.38E-02 | FBgn0262738,FBgn0003416,FBgn0004611,FBgn002588 | PQ$_{head}$ |
| md:M00873 | Fatty acid biosynthesis in mitochondria, animals | 6 | 2 | 0.518 | 1.70E-02 | FBgn0029648,FBgn0036691 | PQ$_{head}$ |
| md:M00171 | C4-dicarboxylic acid cycle, NAD - malic enzyme type | 7 | 2 | 0.518 | 2.32E-02 | FBgn0001125,FBgn0001124 | PQ$_{head}$ |
| path:map00340 | Histidine metabolism | 8 | 2 | 0.518 | 3.00E-02 | FBgn0051075,FBgn0010548 | PQ$_{head}$ |
| md:M00740 | Methylaspartate cycle | 9 | 2 | 0.518 | 3.75E-02 | FBgn0039071,FBgn0001098 | PQ$_{head}$ |
| path:map04070 | Phosphatidylinositol signaling system | 61 | 5 | 1.295 | 4.06E-02 | FBgn0262738,FBgn0030465,FBgn0003416,FBgn000461 | PQ$_{head}$ |
| path:map01100 | Metabolic pathways | 1153 | 72 | 40 | 3.15E-24 | FBgn0086450,FBgn0028342,FBgn0030222,FBgn003022 | PQ$_{body}$ |
| md:M00158 | F-type ATPase, eukaryotes | 18 | 18 | 10 | 3.15E-24 | FBgn0028342,FBgn0031941,FBgn0011211,FBgn003503 | PQ$_{body}$ |
| path:map00480 | Glutathione metabolism | 79 | 20 | 11.111 | 1.57E-19 | FBgn0030222,FBgn0030223,FBgn0030411,FBgn003386 | PQ$_{body}$ |
| path:map00071 | Fatty acid degradation | 33 | 15 | 8.333 | 3.32E-19 | FBgn0031813,FBgn0028479,FBgn0040064,FBgn003216 | PQ$_{body}$ |
| path:map01212 | Fatty acid metabolism | 53 | 17 | 9.444 | 9.42E-19 | FBgn0030731,FBgn0031813,FBgn0028479,FBgn004006 | PQ$_{body}$ |
| md:M00087 | beta-Oxidation | 12 | 8 | 4.444 | 3.24E-12 | FBgn0031813,FBgn0028479,FBgn0040064,FBgn003387 | PQ$_{body}$ |
| path:map00190 | Oxidative phosphorylation | 146 | 18 | 10 | 4.18E-12 | FBgn0028342,FBgn0031941,FBgn0011211,FBgn003503 | PQ$_{body}$ |
| path:map00410 | beta-Alanine metabolism | 25 | 7 | 3.889 | 1.63E-07 | FBgn0086450,FBgn0031813,FBgn0028479,FBgn003387 | PQ$_{body}$ |
| path:map00062 | Fatty acid elongation | 16 | 6 | 3.333 | 2.12E-07 | FBgn0028479,FBgn0040064,FBgn0033879,FBgn002535 | PQ$_{body}$ |
| md:M00085 | Fatty acid elongation in mitochondria | 5 | 4 | 2.222 | 8.32E-07 | FBgn0028479,FBgn0040064,FBgn0033879,FBgn002535 | PQ$_{body}$ |
| md:M00046 | Pyrimidine degradation, uracil => beta-alanine, thymine => 3-amino | 3 | 3 | 1.667 | 1.21E-05 | FBgn0086450,FBgn0023023,FBgn0037513 | PQ$_{body}$ |
| path:map00280 | Valine, leucine and isoleucine degradation | 33 | 6 | 3.333 | 1.83E-05 | FBgn0021765,FBgn0028479,FBgn0040064,FBgn003387 | PQ$_{body}$ |
| path:map04146 | peroxisome | 96 | 9 | 5 | 1.92E-05 | FBgn0030731,FBgn0031813,FBgn0028479,FBgn002757 | PQ$_{body}$ |
| md:M00086 | beta-Oxidation, acyl-CoA synthesis | 4 | 3 | 1.667 | 3.75E-05 | FBgn0036821,FBgn0027348,FBgn0263120 | PQ$_{body}$ |
| path:map01040 | Biosynthesis of unsaturated fatty acids | 25 | 5 | 2.778 | 6.09E-05 | FBgn0030731,FBgn0031813,FBgn0027572,FBgn003547 | PQ$_{body}$ |
| path:map00061 | Fatty acid biosynthesis | 13 | 4 | 2.222 | 6.86E-05 | FBgn0033246,FBgn0036821,FBgn0027348,FBgn026312 | PQ$_{body}$ |
| path:map00640 | Propanoate metabolism | 27 | 5 | 2.778 | 7.99E-05 | FBgn0031813,FBgn0028479,FBgn0033246,FBgn003387 | PQ$_{body}$ |
| path:map00770 | Pantothenate and CoA biosynthesis | 14 | 4 | 2.222 | 8.45E-05 | FBgn0086450,FBgn0023023,FBgn0037513,FBgn005209 | PQ$_{body}$ |
| path:map00330 | Arginine and proline metabolism | 53 | 6 | 3.333 | 1.94E-04 | FBgn0033860,FBgn0034132,FBgn0259795,FBgn003591 | PQ$_{body}$ |
| path:map02010 | ABC transporters | 38 | 5 | 2.778 | 3.76E-04 | FBgn0031170,FBgn0028539,FBgn0034493,FBgn003674 | PQ$_{body}$ |
| md:M00861 | beta-Oxidation, peroxisome, VLCFA | 9 | 3 | 1.667 | 5.01E-04 | FBgn0030731,FBgn0031813,FBgn0027572 | PQ$_{body}$ |
| md:M00013 | Malonate semialdehyde pathway, propanoyl-CoA => acetyl-CoA | 12 | 3 | 1.667 | 1.22E-03 | FBgn0031813,FBgn0033879,FBgn0027572 | PQ$_{body}$ |

| | | | | | | | |
|---|---|---|---|---|---|---|---|
| path:map00910 | Nitrogen metabolism | 18 | 3 | 1.667 | 4.08E-03 | FBgn0027844,FBgn0033542,FBgn0034560 | $PQ_{body}$ |
| md:M00113 | Jasmonic acid biosynthesis | 6 | 2 | 1.111 | 6.29E-03 | FBgn0031813,FBgn0027572 | $PQ_{body}$ |
| md:M00032 | Lysine degradation, lysine => saccharopine => acetoacetyl-CoA | 8 | 2 | 1.111 | 1.11E-02 | FBgn0028479,FBgn0033879 | $PQ_{body}$ |
| md:M00415 | Fatty acid elongation in endoplasmic reticulum | 9 | 2 | 1.111 | 1.31E-02 | FBgn0035471,FBgn0260960 | $PQ_{body}$ |
| path:map00920 | Sulfur metabolism | 9 | 2 | 1.111 | 1.31E-02 | FBgn0030465,FBgn0039698 | $PQ_{body}$ |
| path:map00051 | Fructose and mannose metabolism | 29 | 3 | 1.667 | 1.35E-02 | FBgn0033101,FBgn0035476,FBgn0086254 | $PQ_{body}$ |
| path:map04070 | Phosphatidylinositol signaling system | 61 | 4 | 2.222 | 1.71E-02 | FBgn0030465,FBgn0016984,FBgn0034789,FBgn003605 | $PQ_{body}$ |
| md:M00130 | Inositol phosphate metabolism, PI=> PIP2 => Ins(1,4,5)P3 => Ins(1 | 11 | 2 | 1.111 | 1.77E-02 | FBgn0016984,FBgn0034789 | $PQ_{body}$ |
| path:map00052 | Galactose metabolism | 35 | 3 | 1.667 | 2.07E-02 | FBgn0033101,FBgn0035476,FBgn0086254 | $PQ_{body}$ |
| md:M00959 | Guanine ribonucleotide degradation, GMP => Urate | 13 | 2 | 1.111 | 2.24E-02 | FBgn0034898,FBgn0035348 | $PQ_{body}$ |
| path:map00592 | alpha-Linolenic acid metabolism | 13 | 2 | 1.111 | 2.24E-02 | FBgn0031813,FBgn0027572 | $PQ_{body}$ |
| path:map00240 | Pyrimidine metabolism | 38 | 3 | 1.667 | 2.36E-02 | FBgn0086450,FBgn0023023,FBgn0037513 | $PQ_{body}$ |
| path:map00790 | Folate biosynthesis | 40 | 3 | 1.667 | 2.64E-02 | FBgn0033101,FBgn0035476,FBgn0086254 | $PQ_{body}$ |
| path:map00650 | Butanoate metabolism | 15 | 2 | 1.111 | 2.72E-02 | FBgn0028479,FBgn0033879 | $PQ_{body}$ |
| path:map00561 | Glycerolipid metabolism | 42 | 3 | 1.667 | 2.85E-02 | FBgn0033101,FBgn0035476,FBgn0086254 | $PQ_{body}$ |
| path:map00760 | Nicotinate and nicotinamide metabolism | 18 | 2 | 1.111 | 3.66E-02 | FBgn0034898,FBgn0035348 | $PQ_{body}$ |
| md:M00958 | Adenine ribonucleotide degradation, AMP => Urate | 19 | 2 | 1.111 | 3.95E-02 | FBgn0034898,FBgn0035348 | $PQ_{body}$ |
| path:map00040 | Pentose and glucuronate interconversions | 51 | 3 | 1.667 | 4.39E-02 | FBgn0033101,FBgn0035476,FBgn0086254 | $PQ_{body}$ |
| path:map00380 | Tryptophan metabolism | 21 | 2 | 1.111 | 4.54E-02 | FBgn0028479,FBgn0033879 | $PQ_{body}$ |
| path:map01100 | Metabolic pathways | 1153 | 194 | 51.187 | 5.53E-24 | FBgn0262738,FBgn0029969,FBgn0029975,FBgn001032 | $\alpha Syn_{head}$ |
| path:map00970 | Aminoacyl-tRNA biosynthesis | 64 | 34 | 8.971 | 5.53E-24 | FBgn0030007,FBgn0027084,FBgn0027093,FBgn002708 | $\alpha Syn_{head}$ |
| md:M00158 | F-type ATPase, eukaryotes | 18 | 18 | 4.749 | 5.53E-24 | FBgn0028342,FBgn0031941,FBgn0011211,FBgn003503 | $\alpha Syn_{head}$ |
| path:map01212 | Fatty acid metabolism | 53 | 25 | 6.596 | 5.88E-24 | FBgn0029969,FBgn0029975,FBgn0283427,FBgn002847 | $\alpha Syn_{head}$ |
| path:map00230 | Purine metabolism | 92 | 26 | 6.86 | 2.93E-18 | FBgn0003301,FBgn0022710,FBgn0024947,FBgn003166 | $\alpha Syn_{head}$ |
| path:map01240 | Biosynthesis of cofactors | 139 | 29 | 7.652 | 1.61E-16 | FBgn0014032,FBgn0003965,FBgn0030407,FBgn003104 | $\alpha Syn_{head}$ |
| path:map00071 | Fatty acid degradation | 33 | 16 | 4.222 | 1.15E-15 | FBgn0029969,FBgn0028479,FBgn0040064,FBgn003216 | $\alpha Syn_{head}$ |
| path:map00564 | Glycerophospholipid metabolism | 63 | 20 | 5.277 | 2.74E-15 | FBgn0003656,FBgn0030013,FBgn0052699,FBgn003042 | $\alpha Syn_{head}$ |
| path:map00561 | Glycerolipid metabolism | 42 | 15 | 3.958 | 2.26E-12 | FBgn0030421,FBgn0033101,FBgn0033214,FBgn003321 | $\alpha Syn_{head}$ |
| path:map00062 | Fatty acid elongation | 16 | 9 | 2.375 | 1.16E-09 | FBgn0029975,FBgn0028479,FBgn0040064,FBgn003239 | $\alpha Syn_{head}$ |
| md:M00095 | C5 isoprenoid biosynthesis, mevalonate pathway | 8 | 7 | 1.847 | 1.21E-09 | FBgn0029969,FBgn0030683,FBgn0032811,FBgn003520 | $\alpha Syn_{head}$ |
| path:map00600 | Sphingolipid metabolism | 30 | 11 | 2.902 | 2.42E-09 | FBgn0030300,FBgn0028916,FBgn0034997,FBgn003542 | $\alpha Syn_{head}$ |
| path:map00061 | Fatty acid biosynthesis | 13 | 8 | 2.111 | 3.99E-09 | FBgn0283427,FBgn0286723,FBgn0033246,FBgn003669 | $\alpha Syn_{head}$ |
| path:map00280 | Valine, leucine and isoleucine degradation | 33 | 11 | 2.902 | 6.82E-09 | FBgn0029969,FBgn0021765,FBgn0028479,FBgn004006 | $\alpha Syn_{head}$ |
| path:map00380 | Tryptophan metabolism | 21 | 9 | 2.375 | 1.77E-08 | FBgn0029969,FBgn0003965,FBgn0287695,FBgn002847 | $\alpha Syn_{head}$ |
| path:map00190 | Oxidative phosphorylation | 146 | 19 | 5.013 | 1.57E-07 | FBgn0028342,FBgn0031941,FBgn0011211,FBgn001668 | $\alpha Syn_{head}$ |
| path:map00650 | Butanoate metabolism | 15 | 7 | 1.847 | 5.34E-07 | FBgn0029969,FBgn0028479,FBgn0033879,FBgn003520 | $\alpha Syn_{head}$ |
| path:map00790 | Folate biosynthesis | 40 | 10 | 2.639 | 7.27E-07 | FBgn0014032,FBgn0030407,FBgn0003141,FBgn003310 | $\alpha Syn_{head}$ |
| md:M00849 | C5 isoprenoid biosynthesis, mevalonate pathway, archaea | 6 | 5 | 1.319 | 7.35E-07 | FBgn0029969,FBgn0035203,FBgn0263782,FBgn001061 | $\alpha Syn_{head}$ |
| path:map00565 | Ether lipid metabolism | 25 | 8 | 2.111 | 1.64E-06 | FBgn0030013,FBgn0052699,FBgn0032923,FBgn028651 | $\alpha Syn_{head}$ |
| md:M00087 | beta-Oxidation | 12 | 6 | 1.583 | 2.44E-06 | FBgn0028479,FBgn0040064,FBgn0033879,FBgn003443 | $\alpha Syn_{head}$ |

| | | | | | | | |
|---|---|---|---|---|---|---|---|
| path:map00860 | Porphyrin metabolism | 46 | 10 | 2.639 | 2.47E-06 | FBgn0032684,FBgn0027074,FBgn0027070,FBgn002707 | $\alpha Syn_{head}$ |
| path:map00260 | Glycine, serine and threonine metabolism | 27 | 8 | 2.111 | 2.78E-06 | FBgn0031148,FBgn0031860,FBgn0034276,FBgn000056 | $\alpha Syn_{head}$ |
| md:M00086 | beta-Oxidation, acyl-CoA synthesis | 4 | 4 | 1.055 | 3.69E-06 | FBgn0286723,FBgn0036821,FBgn0027348,FBgn026312 | $\alpha Syn_{head}$ |
| md:M00415 | Fatty acid elongation in endoplasmic reticulum | 9 | 5 | 1.319 | 1.10E-05 | FBgn0029975,FBgn0032394,FBgn0032524,FBgn003547 | $\alpha Syn_{head}$ |
| md:M00089 | Triacylglycerol biosynthesis | 16 | 6 | 1.583 | 1.49E-05 | FBgn0030421,FBgn0034971,FBgn0036622,FBgn003662 | $\alpha Syn_{head}$ |
| md:M00042 | Catecholamine biosynthesis, tyrosine => dopamine => noradrenalin | 5 | 4 | 1.055 | 1.49E-05 | FBgn0010329,FBgn0000422,FBgn0283437,FBgn026136 | $\alpha Syn_{head}$ |
| md:M00082 | Fatty acid biosynthesis, initiation | 5 | 4 | 1.055 | 1.49E-05 | FBgn0283427,FBgn0033246,FBgn0036691,FBgn004262 | $\alpha Syn_{head}$ |
| md:M00085 | Fatty acid elongation in mitochondria | 5 | 4 | 1.055 | 1.49E-05 | FBgn0028479,FBgn0040064,FBgn0033879,FBgn002535 | $\alpha Syn_{head}$ |
| path:map01040 | Biosynthesis of unsaturated fatty acids | 25 | 7 | 1.847 | 1.79E-05 | FBgn0029975,FBgn0032394,FBgn0032524,FBgn003547 | $\alpha Syn_{head}$ |
| path:map01232 | Nucleotide metabolism | 60 | 10 | 2.639 | 2.31E-05 | FBgn0024947,FBgn0031663,FBgn0032001,FBgn003200 | $\alpha Syn_{head}$ |
| path:map00640 | Propanoate metabolism | 27 | 7 | 1.847 | 2.86E-05 | FBgn0028479,FBgn0033246,FBgn0033879,FBgn003676 | $\alpha Syn_{head}$ |
| path:map00900 | Terpenoid backbone biosynthesis | 27 | 7 | 1.847 | 2.86E-05 | FBgn0029969,FBgn0030683,FBgn0032811,FBgn003520 | $\alpha Syn_{head}$ |
| path:map00040 | Pentose and glucuronate interconversions | 51 | 9 | 2.375 | 3.93E-05 | FBgn0032684,FBgn0027074,FBgn0027070,FBgn003310 | $\alpha Syn_{head}$ |
| path:map00350 | Tyrosine metabolism | 22 | 6 | 1.583 | 9.14E-05 | FBgn0010329,FBgn0030558,FBgn0000422,FBgn028343 | $\alpha Syn_{head}$ |
| path:map00601 | Glycosphingolipid biosynthesis - lacto and neolacto series | 3 | 3 | 0.792 | 9.14E-05 | FBgn0031491,FBgn0039378,FBgn0050037 | $\alpha Syn_{head}$ |
| md:M00129 | Ascorbate biosynthesis, animals, glucose-1P => ascorbate | 33 | 7 | 1.847 | 1.07E-04 | FBgn0032684,FBgn0027074,FBgn0027070,FBgn002707 | $\alpha Syn_{head}$ |
| path:map00562 | Inositol phosphate metabolism | 46 | 8 | 2.111 | 1.25E-04 | FBgn0262738,FBgn0003416,FBgn0004611,FBgn002588 | $\alpha Syn_{head}$ |
| path:map00830 | Retinol metabolism | 34 | 7 | 1.847 | 1.25E-04 | FBgn0032684,FBgn0027074,FBgn0027070,FBgn002707 | $\alpha Syn_{head}$ |
| md:M00032 | Lysine degradation, lysine => saccharopine => acetoacetyl-CoA | 8 | 4 | 1.055 | 0.00014216 | FBgn0028479,FBgn0033879,FBgn0036762,FBgn003789 | $\alpha Syn_{head}$ |
| path:map00053 | Ascorbate and aldarate metabolism | 36 | 7 | 1.847 | 0.000175449 | FBgn0032684,FBgn0027074,FBgn0027070,FBgn002707 | $\alpha Syn_{head}$ |
| path:map04146 | peroxisome | 96 | 11 | 2.902 | 0.000224168 | FBgn0031860,FBgn0028479,FBgn0032811,FBgn003322 | $\alpha Syn_{head}$ |
| md:M00014 | Glucuronate pathway (uronate pathway) | 38 | 7 | 1.847 | 0.000241371 | FBgn0032684,FBgn0027074,FBgn0027070,FBgn002707 | $\alpha Syn_{head}$ |
| md:M00049 | Adenine ribonucleotide biosynthesis, IMP => ADP,ATP | 10 | 4 | 1.055 | 0.000371292 | FBgn0033754,FBgn0283494,FBgn0022709,FBgn004209 | $\alpha Syn_{head}$ |
| path:map01250 | Biosynthesis of nucleotide sugars | 30 | 6 | 1.583 | 0.000483653 | FBgn0033377,FBgn0033969,FBgn0035147,FBgn003958 | $\alpha Syn_{head}$ |
| path:map00450 | Selenocompound metabolism | 11 | 4 | 1.055 | 0.000546164 | FBgn0000566,FBgn0034401,FBgn0037170,FBgn002708 | $\alpha Syn_{head}$ |
| md:M00120 | Coenzyme A biosynthesis, pantothenate => CoA | 5 | 3 | 0.792 | 0.000658146 | FBgn0031682,FBgn0011205,FBgn0050290 | $\alpha Syn_{head}$ |
| md:M00855 | Glycogen degradation, glycogen => glucose-6P | 5 | 3 | 0.792 | 0.000658146 | FBgn0033377,FBgn0033969,FBgn0003076 | $\alpha Syn_{head}$ |
| path:map00730 | Thiamine metabolism | 21 | 5 | 1.319 | 0.000689682 | FBgn0283494,FBgn0035619,FBgn0035620,FBgn002270 | $\alpha Syn_{head}$ |
| path:map04070 | Phosphatidylinositol signaling system | 61 | 8 | 2.111 | 0.000759127 | FBgn0262738,FBgn0003416,FBgn0004611,FBgn002874 | $\alpha Syn_{head}$ |
| path:map00983 | Drug metabolism - other enzymes | 96 | 10 | 2.639 | 0.000885291 | FBgn0031663,FBgn0032001,FBgn0032002,FBgn003268 | $\alpha Syn_{head}$ |
| path:map00592 | alpha-Linolenic acid metabolism | 13 | 4 | 1.055 | 0.000982634 | FBgn0030013,FBgn0036053,FBgn0036545,FBgn003965 | $\alpha Syn_{head}$ |
| path:map00052 | Galactose metabolism | 35 | 6 | 1.583 | 0.000982634 | FBgn0033101,FBgn0033377,FBgn0033969,FBgn003514 | $\alpha Syn_{head}$ |
| path:map00310 | Lysine degradation | 35 | 6 | 1.583 | 0.000982634 | FBgn0029969,FBgn0028479,FBgn0033879,FBgn003520 | $\alpha Syn_{head}$ |
| md:M00088 | Ketone body biosynthesis, acetyl-CoA => acetoacetate/3-hydroxybu | 6 | 3 | 0.792 | 0.001111357 | FBgn0029969,FBgn0035203,FBgn0010611 | $\alpha Syn_{head}$ |
| path:map00270 | Cysteine and methionine metabolism | 36 | 6 | 1.583 | 0.001111357 | FBgn0030558,FBgn0031148,FBgn0032726,FBgn003272 | $\alpha Syn_{head}$ |
| path:map00030 | Pentose phosphate pathway | 24 | 5 | 1.319 | 0.001158131 | FBgn0032424,FBgn0033377,FBgn0033969,FBgn003603 | $\alpha Syn_{head}$ |
| path:map01200 | Carbon metabolism | 121 | 11 | 2.902 | 0.001285672 | FBgn0029969,FBgn0033879,FBgn0035203,FBgn003603 | $\alpha Syn_{head}$ |
| md:M00126 | Tetrahydrofolate biosynthesis, GTP => THF | 15 | 4 | 1.055 | 0.001588925 | FBgn0035619,FBgn0035620,FBgn0004087,FBgn003884 | $\alpha Syn_{head}$ |
| path:map00590 | Arachidonic acid metabolism | 15 | 4 | 1.055 | 0.001588925 | FBgn0030013,FBgn0036053,FBgn0036545,FBgn003965 | $\alpha Syn_{head}$ |
| md:M00099 | Sphingosine biosynthesis | 7 | 3 | 0.792 | 0.001740371 | FBgn0039774,FBgn0040918,FBgn0045064 | $\alpha Syn_{head}$ |
| md:M00338 | Cysteine biosynthesis, homocysteine + serine => cysteine | 2 | 2 | 0.528 | 0.001875376 | FBgn0031148,FBgn0000566 | $\alpha Syn_{head}$ |

| | | | | | | | |
|---|---|---|---|---|---|---|---|
| md:M00373 | Ethylmalonyl pathway | 2 | 2 | 0.528 | 0.001875376 | FBgn0029969,FBgn0035203 | $\alpha Syn_{head}$ |
| md:M00374 | Dicarboxylate-hydroxybutyrate cycle | 2 | 2 | 0.528 | 0.001875376 | FBgn0029969,FBgn0035203 | $\alpha Syn_{head}$ |
| md:M00375 | Hydroxypropionate-hydroxybutylate cycle | 2 | 2 | 0.528 | 0.001875376 | FBgn0029969,FBgn0035203 | $\alpha Syn_{head}$ |
| md:M00549 | Nucleotide sugar biosynthesis, glucose => UDP-glucose | 8 | 3 | 0.792 | 0.002521945 | FBgn0033377,FBgn0033969,FBgn0003076 | $\alpha Syn_{head}$ |
| path:map00513 | Various types of N-glycan biosynthesis | 30 | 5 | 1.319 | 0.002890268 | FBgn0031149,FBgn0036446,FBgn0036485,FBgn003855 | $\alpha Syn_{head}$ |
| md:M00078 | Heparan sulfate degradation | 9 | 3 | 0.792 | 0.003598005 | FBgn0033836,FBgn0270927,FBgn0260475 | $\alpha Syn_{head}$ |
| path:map00520 | Amino sugar and nucleotide sugar metabolism | 48 | 6 | 1.583 | 0.004310455 | FBgn0033377,FBgn0033969,FBgn0035147,FBgn003958 | $\alpha Syn_{head}$ |
| path:map00630 | Glyoxylate and dicarboxylate metabolism | 33 | 5 | 1.319 | 0.004310455 | FBgn0029969,FBgn0287695,FBgn0035203,FBgn003676 | $\alpha Syn_{head}$ |
| md:M00027 | GABA (gamma-Aminobutyrate) shunt | 3 | 2 | 0.528 | 0.004852935 | FBgn0004516,FBgn0036927 | $\alpha Syn_{head}$ |
| md:M00038 | Tryptophan metabolism, tryptophan => kynurenine => 2-aminomuc | 3 | 2 | 0.528 | 0.004852935 | FBgn0003965,FBgn0287695 | $\alpha Syn_{head}$ |
| md:M00100 | Sphingosine degradation | 3 | 2 | 0.528 | 0.004852935 | FBgn0030300,FBgn0052484 | $\alpha Syn_{head}$ |
| md:M00957 | Lysine degradation, bacteria, L-lysine => glutarate => succinate/ace | 3 | 2 | 0.528 | 0.004852935 | FBgn0029969,FBgn0035203 | $\alpha Syn_{head}$ |
| md:M00009 | Citrate cycle (TCA cycle, Krebs cycle) | 35 | 5 | 1.319 | 0.00526429 | FBgn0010352,FBgn0036762,FBgn0037891,FBgn003935 | $\alpha Syn_{head}$ |
| md:M00130 | Inositol phosphate metabolism, PI=> PIP2 => Ins(1,4,5)P3 => Ins(1 | 11 | 3 | 0.792 | 0.006072924 | FBgn0262738,FBgn0003416,FBgn0004611 | $\alpha Syn_{head}$ |
| path:map00510 | N-Glycan biosynthesis | 37 | 5 | 1.319 | 0.006586919 | FBgn0031149,FBgn0032014,FBgn0036446,FBgn003855 | $\alpha Syn_{head}$ |
| md:M00011 | Citrate cycle, second carbon oxidation, 2-oxoglutarate => oxaloacet | 23 | 4 | 1.055 | 0.00667946 | FBgn0010352,FBgn0036762,FBgn0037891,FBgn003524 | $\alpha Syn_{head}$ |
| path:map00514 | Other types of O-glycan biosynthesis | 38 | 5 | 1.319 | 0.007140897 | FBgn0030930,FBgn0031765,FBgn0261403,FBgn000197 | $\alpha Syn_{head}$ |
| path:map02010 | ABC transporters | 38 | 5 | 1.319 | 0.007140897 | FBgn0031170,FBgn0028539,FBgn0034493,FBgn003674 | $\alpha Syn_{head}$ |
| md:M00083 | Fatty acid biosynthesis, elongation | 4 | 2 | 0.528 | 0.008490409 | FBgn0283427,FBgn0042627 | $\alpha Syn_{head}$ |
| md:M00842 | Tetrahydrobiopterin biosynthesis, GTP => BH4 | 4 | 2 | 0.528 | 0.008490409 | FBgn0014032,FBgn0003141 | $\alpha Syn_{head}$ |
| path:map00410 | beta-Alanine metabolism | 25 | 4 | 1.055 | 0.008490409 | FBgn0028479,FBgn0033879,FBgn0004516,FBgn003692 | $\alpha Syn_{head}$ |
| path:map00563 | Glycosylphosphatidylinositol (GPI)-anchor biosynthesis | 25 | 4 | 1.055 | 0.008490409 | FBgn0034649,FBgn0035464,FBgn0265174,FBgn026643 | $\alpha Syn_{head}$ |
| path:map00770 | Pantothenate and CoA biosynthesis | 14 | 3 | 0.792 | 0.01127602 | FBgn0031682,FBgn0011205,FBgn0050290 | $\alpha Syn_{head}$ |
| path:map00020 | Citrate cycle (TCA cycle) | 43 | 5 | 1.319 | 0.011408816 | FBgn0010352,FBgn0036762,FBgn0037891,FBgn003935 | $\alpha Syn_{head}$ |
| md:M00874 | Fatty acid biosynthesis in mitochondria, fungi | 5 | 2 | 0.528 | 0.012971938 | FBgn0033246,FBgn0036691 | $\alpha Syn_{head}$ |
| md:M00912 | NAD biosynthesis, tryptophan => quinolinate => NAD | 5 | 2 | 0.528 | 0.012971938 | FBgn0003965,FBgn0287695 | $\alpha Syn_{head}$ |
| path:map00603 | Glycosphingolipid biosynthesis - globo and isoglobo series | 5 | 2 | 0.528 | 0.012971938 | FBgn0031491,FBgn0039378 | $\alpha Syn_{head}$ |
| path:map00531 | Glycosaminoglycan degradation | 15 | 3 | 0.792 | 0.013047147 | FBgn0033836,FBgn0270927,FBgn0260475 | $\alpha Syn_{head}$ |
| path:map00250 | Alanine, aspartate and glutamate metabolism | 29 | 4 | 1.055 | 0.013514711 | FBgn0004516,FBgn0036927,FBgn0039580,FBgn028720 | $\alpha Syn_{head}$ |
| path:map00982 | Drug metabolism - cytochrome P450 | 67 | 6 | 1.583 | 0.01717036 | FBgn0032684,FBgn0027074,FBgn0027070,FBgn002707 | $\alpha Syn_{head}$ |
| md:M00065 | GPI-anchor biosynthesis, core oligosaccharide | 17 | 3 | 0.792 | 0.017563343 | FBgn0034649,FBgn0035464,FBgn0265174 | $\alpha Syn_{head}$ |
| path:map01230 | Biosynthesis of amino acids | 68 | 6 | 1.583 | 0.017563343 | FBgn0031148,FBgn0000566,FBgn0036030,FBgn003768 | $\alpha Syn_{head}$ |
| md:M00079 | Keratan sulfate degradation | 6 | 2 | 0.528 | 0.017563343 | FBgn0033836,FBgn0260475 | $\alpha Syn_{head}$ |
| path:map00440 | Phosphonate and phosphinate metabolism | 6 | 2 | 0.528 | 0.017563343 | FBgn0033844,FBgn0041342 | $\alpha Syn_{head}$ |
| path:map00620 | Pyruvate metabolism | 49 | 5 | 1.319 | 0.017563343 | FBgn0029969,FBgn0033246,FBgn0035203,FBgn003676 | $\alpha Syn_{head}$ |
| path:map00980 | Metabolism of xenobiotics by cytochrome P450 | 69 | 6 | 1.583 | 0.018512807 | FBgn0032684,FBgn0027074,FBgn0027070,FBgn002707 | $\alpha Syn_{head}$ |
| path:map00910 | Nitrogen metabolism | 18 | 3 | 0.792 | 0.020030712 | FBgn0027844,FBgn0033542,FBgn0034560 | $\alpha Syn_{head}$ |
| md:M00072 | N-glycosylation by oligosaccharyltransferase | 7 | 2 | 0.528 | 0.023386536 | FBgn0031149,FBgn0011336 | $\alpha Syn_{head}$ |
| path:map00010 | Glycolysis / Gluconeogenesis | 55 | 5 | 1.319 | 0.026724099 | FBgn0033377,FBgn0033969,FBgn0036762,FBgn001203 | $\alpha Syn_{head}$ |
| path:map04213 | Longevity regulating pathway - multiple species | 55 | 5 | 1.319 | 0.026724099 | FBgn0003301,FBgn0022710,FBgn0023416,FBgn000485 | $\alpha Syn_{head}$ |
| path:map00360 | Phenylalanine metabolism | 8 | 2 | 0.528 | 0.029735165 | FBgn0030558,FBgn0000422 | $\alpha Syn_{head}$ |
| path:map00240 | Pyrimidine metabolism | 38 | 4 | 1.055 | 0.030386578 | FBgn0024947,FBgn0032001,FBgn0032002,FBgn003790 | $\alpha Syn_{head}$ |

| | | | | | | | |
|---|---|---|---|---|---|---|---|
| path:map00511 | Other glycan degradation | 22 | 3 | 0.792 | 0.032911124 | FBgn0028916,FBgn0051148,FBgn0051414 | $\alpha Syn_{head}$ |
| path:map00430 | Taurine and hypotaurine metabolism | 9 | 2 | 0.528 | 0.036489446 | FBgn0034364,FBgn0004516 | $\alpha Syn_{head}$ |
| md:M00892 | UDP-N-acetyl-D-glucosamine biosynthesis, eukaryotes, glucose => | 10 | 2 | 0.528 | 0.042403767 | FBgn0039580,FBgn0287209 | $\alpha Syn_{head}$ |
| path:map01100 | Metabolic pathways | 1153 | 123 | 48.047 | 1.07E-23 | FBgn0003116,FBgn0030013,FBgn0014032,FBgn026154 | $\alpha Syn_{body}$ |
| path:map00480 | Glutathione metabolism | 79 | 25 | 9.766 | 6.45587E-23 | FBgn0030222,FBgn0030223,FBgn0030361,FBgn003079 | $\alpha Syn_{body}$ |
| path:map01240 | Biosynthesis of cofactors | 139 | 29 | 11.328 | 3.58491E-21 | FBgn0014032,FBgn0030407,FBgn0030431,FBgn003046 | $\alpha Syn_{body}$ |
| path:map00970 | Aminoacyl-tRNA biosynthesis | 64 | 20 | 7.813 | 2.22108E-18 | FBgn0027084,FBgn0027093,FBgn0031497,FBgn002709 | $\alpha Syn_{body}$ |
| path:map00561 | Glycerolipid metabolism | 42 | 15 | 5.859 | 7.96961E-15 | FBgn0261549,FBgn0033101,FBgn0085390,FBgn003367 | $\alpha Syn_{body}$ |
| path:map00564 | Glycerophospholipid metabolism | 63 | 17 | 6.641 | 1.48603E-14 | FBgn0030013,FBgn0261549,FBgn0052699,FBgn000112 | $\alpha Syn_{body}$ |
| path:map00790 | Folate biosynthesis | 40 | 13 | 5.078 | 2.30056E-12 | FBgn0014032,FBgn0030407,FBgn0003141,FBgn003310 | $\alpha Syn_{body}$ |
| path:map00900 | Terpenoid backbone biosynthesis | 27 | 11 | 4.297 | 8.5131E-12 | FBgn0030683,FBgn0032811,FBgn0025373,FBgn003520 | $\alpha Syn_{body}$ |
| md:M00095 | C5 isoprenoid biosynthesis, mevalonate pathway | 8 | 7 | 2.734 | 5.99601E-11 | FBgn0030683,FBgn0032811,FBgn0035203,FBgn003887 | $\alpha Syn_{body}$ |
| path:map00330 | Arginine and proline metabolism | 53 | 13 | 5.078 | 9.11121E-11 | FBgn0013307,FBgn0033860,FBgn0001124,FBgn003413 | $\alpha Syn_{body}$ |
| path:map01232 | Nucleotide metabolism | 60 | 13 | 5.078 | 4.53837E-10 | FBgn0024920,FBgn0031663,FBgn0011703,FBgn001170 | $\alpha Syn_{body}$ |
| path:map00590 | Arachidonic acid metabolism | 15 | 8 | 3.125 | 5.8631E-10 | FBgn0030013,FBgn0030361,FBgn0030796,FBgn003093 | $\alpha Syn_{body}$ |
| md:M00128 | Ubiquinone biosynthesis, eukaryotes, 4-hydroxybenzoate + polypre | 5 | 5 | 1.953 | 1.63763E-08 | FBgn0030460,FBgn0031713,FBgn0032922,FBgn003757 | $\alpha Syn_{body}$ |
| path:map00430 | Taurine and hypotaurine metabolism | 9 | 6 | 2.344 | 2.19209E-08 | FBgn0030361,FBgn0030796,FBgn0030932,FBgn000451 | $\alpha Syn_{body}$ |
| md:M00849 | C5 isoprenoid biosynthesis, mevalonate pathway, archaea | 6 | 5 | 1.953 | 8.3881E-08 | FBgn0035203,FBgn0038876,FBgn0263782,FBgn001061 | $\alpha Syn_{body}$ |
| path:map00051 | Fructose and mannose metabolism | 29 | 8 | 3.125 | 2.34812E-07 | FBgn0033101,FBgn0027552,FBgn0086254,FBgn003618 | $\alpha Syn_{body}$ |
| path:map00130 | Ubiquinone and other terpenoid-quinone biosynthesis | 9 | 5 | 1.953 | 1.48551E-06 | FBgn0030460,FBgn0031713,FBgn0032922,FBgn003757 | $\alpha Syn_{body}$ |
| path:map00250 | Alanine, aspartate and glutamate metabolism | 29 | 7 | 2.734 | 4.20668E-06 | FBgn0261625,FBgn0001124,FBgn0004516,FBgn003640 | $\alpha Syn_{body}$ |
| path:map00230 | Purine metabolism | 92 | 11 | 4.297 | 5.16331E-06 | FBgn0003116,FBgn0031663,FBgn0011703,FBgn001170 | $\alpha Syn_{body}$ |
| path:map00040 | Pentose and glucuronate interconversions | 51 | 8 | 3.125 | 1.96669E-05 | FBgn0033101,FBgn0027552,FBgn0086254,FBgn003618 | $\alpha Syn_{body}$ |
| md:M00911 | Riboflavin biosynthesis, fungi, GTP => riboflavin/FMN/FAD | 3 | 3 | 1.172 | 3.10077E-05 | FBgn0030431,FBgn0032522,FBgn0014930 | $\alpha Syn_{body}$ |
| path:map00760 | Nicotinate and nicotinamide metabolism | 18 | 5 | 1.953 | 6.75291E-05 | FBgn0033373,FBgn0033853,FBgn0053156,FBgn003489 | $\alpha Syn_{body}$ |
| path:map04070 | Phosphatidylinositol signaling system | 61 | 8 | 3.125 | 6.75291E-05 | FBgn0261549,FBgn0030465,FBgn0085390,FBgn002093 | $\alpha Syn_{body}$ |
| md:M00049 | Adenine ribonucleotide biosynthesis, IMP => ADP,ATP | 10 | 4 | 1.563 | 9.46095E-05 | FBgn0033754,FBgn0283494,FBgn0022709,FBgn004209 | $\alpha Syn_{body}$ |
| path:map00730 | Thiamine metabolism | 21 | 5 | 1.953 | 0.000136202 | FBgn0283494,FBgn0035619,FBgn0035620,FBgn002270 | $\alpha Syn_{body}$ |
| path:map00052 | Galactose metabolism | 35 | 6 | 2.344 | 0.000152824 | FBgn0033101,FBgn0027552,FBgn0086254,FBgn003618 | $\alpha Syn_{body}$ |
| path:map00240 | Pyrimidine metabolism | 38 | 6 | 2.344 | 0.000230504 | FBgn0024920,FBgn0011703,FBgn0011704,FBgn002202 | $\alpha Syn_{body}$ |
| path:map02010 | ABC transporters | 38 | 6 | 2.344 | 0.000230504 | FBgn0031170,FBgn0028539,FBgn0010241,FBgn003449 | $\alpha Syn_{body}$ |
| path:map00592 | alpha-Linolenic acid metabolism | 13 | 4 | 1.563 | 0.000255248 | FBgn0030013,FBgn0036053,FBgn0036545,FBgn003965 | $\alpha Syn_{body}$ |
| path:map00565 | Ether lipid metabolism | 25 | 5 | 1.953 | 0.000279098 | FBgn0030013,FBgn0052699,FBgn0036053,FBgn003654 | $\alpha Syn_{body}$ |
| path:map00770 | Pantothenate and CoA biosynthesis | 14 | 4 | 1.563 | 0.00032949 | FBgn0031682,FBgn0051075,FBgn0011205,FBgn005209 | $\alpha Syn_{body}$ |
| md:M00088 | Ketone body biosynthesis, acetyl-CoA => acetoacetate/3-hydroxybu | 6 | 3 | 1.172 | 0.000390624 | FBgn0031877,FBgn0035203,FBgn0010611 | $\alpha Syn_{body}$ |
| path:map00650 | Butanoate metabolism | 15 | 4 | 1.563 | 0.000416015 | FBgn0031877,FBgn0035203,FBgn0004516,FBgn001061 | $\alpha Syn_{body}$ |
| path:map00280 | Valine, leucine and isoleucine degradation | 33 | 5 | 1.953 | 0.000973089 | FBgn0031877,FBgn0033761,FBgn0035203,FBgn005107 | $\alpha Syn_{body}$ |
| path:map00740 | Riboflavin metabolism | 8 | 3 | 1.172 | 0.000973089 | FBgn0030431,FBgn0032522,FBgn0014930 | $\alpha Syn_{body}$ |
| md:M00938 | Pyrimidine deoxyribonucleotide biosynthesis, UDP => dTTP | 9 | 3 | 1.172 | 0.00139989 | FBgn0024920,FBgn0011703,FBgn0011704 | $\alpha Syn_{body}$ |
| md:M00015 | Proline biosynthesis, glutamate => proline | 3 | 2 | 0.781 | 0.002867796 | FBgn0038516,FBgn0015781 | $\alpha Syn_{body}$ |
| path:map00981 | Insect hormone biosynthesis | 27 | 4 | 1.563 | 0.003907612 | FBgn0024995,FBgn0010383,FBgn0003486,FBgn005107 | $\alpha Syn_{body}$ |
| md:M00842 | Tetrahydrobiopterin biosynthesis, GTP => BH4 | 4 | 2 | 0.781 | 0.00537558 | FBgn0014032,FBgn0003141 | $\alpha Syn_{body}$ |

| | | | | | | | |
|---|---|---|---|---|---|---|---|
| md:M00126 | Tetrahydrofolate biosynthesis, GTP => THF | 15 | 3 | 1.172 | 0.006290516 | FBgn0035619,FBgn0035620,FBgn0038845 | αSyn$_{body}$ |
| md:M00120 | Coenzyme A biosynthesis, pantothenate => CoA | 5 | 2 | 0.781 | 0.008419238 | FBgn0031682,FBgn0011205 | αSyn$_{body}$ |
| path:map00220 | Arginine biosynthesis | 18 | 3 | 1.172 | 0.010317076 | FBgn0261625,FBgn0001124,FBgn0001098 | αSyn$_{body}$ |
| md:M00053 | Deoxyribonucleotide biosynthesis, ADP/GDP/CDP/UDP => dATP/ | 6 | 2 | 0.781 | 0.01189612 | FBgn0011703,FBgn0011704 | αSyn$_{body}$ |
| path:map00350 | Tyrosine metabolism | 22 | 3 | 1.172 | 0.017610327 | FBgn0001124,FBgn0011768,FBgn0010548 | αSyn$_{body}$ |
| path:map00340 | Histidine metabolism | 8 | 2 | 0.781 | 0.020260884 | FBgn0051075,FBgn0010548 | αSyn$_{body}$ |
| path:map00360 | Phenylalanine metabolism | 8 | 2 | 0.781 | 0.020260884 | FBgn0001124,FBgn0010548 | αSyn$_{body}$ |
| path:map00410 | beta-Alanine metabolism | 25 | 3 | 1.172 | 0.023649641 | FBgn0004516,FBgn0051075,FBgn0010548 | αSyn$_{body}$ |
| md:M00131 | Inositol phosphate metabolism, Ins(1,3,4,5)P4 => Ins(1,3,4)P3 => m | 9 | 2 | 0.781 | 0.024160974 | FBgn0037063,FBgn0016672 | αSyn$_{body}$ |
| path:map00920 | Sulfur metabolism | 9 | 2 | 0.781 | 0.024160974 | FBgn0030465,FBgn0039698 | αSyn$_{body}$ |
| path:map00620 | Pyruvate metabolism | 49 | 4 | 1.563 | 0.026202121 | FBgn0035203,FBgn0037370,FBgn0011768,FBgn00510[illegible] | αSyn$_{body}$ |
| path:map00260 | Glycine, serine and threonine metabolism | 27 | 3 | 1.172 | 0.026984107 | FBgn0036208,FBgn0036997,FBgn0037370 | αSyn$_{body}$ |
| md:M00036 | Leucine degradation, leucine => acetoacetate + acetyl-CoA | 11 | 2 | 0.781 | 0.033313369 | FBgn0031877,FBgn0033761 | αSyn$_{body}$ |
| path:map00450 | Selenocompound metabolism | 11 | 2 | 0.781 | 0.033313369 | FBgn0034401,FBgn0027083 | αSyn$_{body}$ |
| path:map00600 | Sphingolipid metabolism | 30 | 3 | 1.172 | 0.03398311 | FBgn0034997,FBgn0035421,FBgn0037958 | αSyn$_{body}$ |
| path:map00071 | Fatty acid degradation | 33 | 3 | 1.172 | 0.040851344 | FBgn0035203,FBgn0011768,FBgn0051075 | αSyn$_{body}$ |
| md:M00959 | Guanine ribonucleotide degradation, GMP => Urate | 13 | 2 | 0.781 | 0.041431466 | FBgn0034898,FBgn0035348 | αSyn$_{body}$ |
