## Supplemental Table 4 for "Comparative Proteomic Analysis of Environmental and Genetic Models of Parkinson’s Disease Highlights the Role of Purine Metabolism"

**Table S4 |** Significantly enriched KEGG pathways (p-value < 0.05 and pathway impact > 0.10) of the metabolites in the differential *Drosophila* reactions by the MetaboAnalyst tool for paraquat treatment or α-

| **Pathway Term (*Homo sapiens*)** | **Total** | **Hits** | **Raw *p*-value** | **Impact** | **PD Group** |
|---|---|---|---|---|---|
| Biosynthesis of unsaturated fatty acids | 36 | 34 | 4.54E-26 | 0.99998 | $PQ_{head}$ |
| Pyrimidine metabolism | 39 | 25 | 6.90E-12 | 0.70247 | $PQ_{head}$ |
| One carbon pool by folate | 26 | 17 | 1.31E-08 | 0.66541 | $PQ_{head}$ |
| Pantothenate and CoA biosynthesis | 20 | 14 | 7.48E-08 | 0.61905 | $PQ_{head}$ |
| Purine metabolism | 70 | 28 | 5.73E-07 | 0.40806 | $PQ_{head}$ |
| Glyoxylate and dicarboxylate metabolism | 32 | 17 | 9.48E-07 | 0.65834 | $PQ_{head}$ |
| Glycine,serine and threonine metabolism | 33 | 17 | 1.68E-06 | 0.57724 | $PQ_{head}$ |
| Terpenoid backbone biosynthesis | 18 | 11 | 1.50E-05 | 0.87301 | $PQ_{head}$ |
| Alanine,aspartate and glutamate metabolism | 28 | 14 | 2.29E-05 | 0.77405 | $PQ_{head}$ |
| Glycerophospholipid metabolism | 36 | 15 | 1.66E-04 | 0.61908 | $PQ_{head}$ |
| Pyruvate metabolism | 23 | 11 | 3.01E-04 | 0.48516 | $PQ_{head}$ |
| Arginine biosynthesis | 14 | 8 | 4.59E-04 | 0.60106 | $PQ_{head}$ |
| Nitrogen metabolism | 6 | 5 | 5.12E-04 | 1 | $PQ_{head}$ |
| Lipoic acid metabolism | 28 | 12 | 5.63E-04 | 0.32273 | $PQ_{head}$ |
| Glutathione metabolism | 28 | 12 | 5.63E-04 | 0.51516 | $PQ_{head}$ |
| Butanoate metabolism | 15 | 8 | 8.50E-04 | 0.42857 | $PQ_{head}$ |
| beta-Alanine metabolism | 21 | 9 | 2.84E-03 | 0.61567 | $PQ_{head}$ |
| Cysteine and methionine metabolism | 33 | 12 | 3.10E-03 | 0.54321 | $PQ_{head}$ |
| Sulfur metabolism | 8 | 5 | 3.62E-03 | 0.78724 | $PQ_{head}$ |
| Taurine and hypotaurine metabolism | 8 | 5 | 3.62E-03 | 1 | $PQ_{head}$ |
| Glycerolipid metabolism | 16 | 7 | 7.43E-03 | 0.27071 | $PQ_{head}$ |
| Citrate cycle (TCA cycle) | 20 | 8 | 8.04E-03 | 0.42457 | $PQ_{head}$ |
| Phenylalanine,tyrosine and tryptophan biosynthesis | 4 | 3 | 1.40E-02 | 1 | $PQ_{head}$ |
| Riboflavin metabolism | 4 | 3 | 1.40E-02 | 1 | $PQ_{head}$ |
| Retinol metabolism | 17 | 6 | 4.02E-02 | 0.72165 | $PQ_{head}$ |
| Glycolysis or Gluconeogenesis | 26 | 8 | 4.21E-02 | 0.28432 | $PQ_{head}$ |
| Arginine and proline metabolism | 36 | 10 | 4.76E-02 | 0.42325 | $PQ_{head}$ |
| Glutathione metabolism | 28 | 9 | 5.66E-06 | 0.44798 | $PQ_{body}$ |
| Pantothenate and CoA biosynthesis | 20 | 7 | 3.63E-05 | 0.4966 | $PQ_{body}$ |
| Glyoxylate and dicarboxylate metabolism | 32 | 8 | 1.47E-04 | 0.17833 | $PQ_{body}$ |
| Biosynthesis of unsaturated fatty acids | 36 | 8 | 3.56E-04 | 0.54545 | $PQ_{body}$ |
| Nicotinate and nicotinamide metabolism | 15 | 5 | 6.79E-04 | 0.42544 | $PQ_{body}$ |
| Purine metabolism | 70 | 11 | 6.88E-04 | 0.15225 | $PQ_{body}$ |
| Glycine,serine and threonine metabolism | 33 | 7 | 1.15E-03 | 0.36486 | $PQ_{body}$ |
| One carbon pool by folate | 26 | 6 | 1.66E-03 | 0.1954 | $PQ_{body}$ |
| Nitrogen metabolism | 6 | 3 | 2.44E-03 | 1 | $PQ_{body}$ |
| Inositol phosphate metabolism | 30 | 6 | 3.63E-03 | 0.33864 | $PQ_{body}$ |

| | | | | | |
|---|---|---|---|---|---|
| Pyruvate metabolism | 23 | 5 | 5.46E-03 | 0.26414 | $PQ_{body}$ |
| Propanoate metabolism | 22 | 4 | 2.45E-02 | 0.31539 | $PQ_{body}$ |
| Biosynthesis of unsaturated fatty acids | 36 | 34 | 1.91E-26 | 0.99998 | $\alpha Syn_{head}$ |
| One carbon pool by folate | 26 | 21 | 1.52E-13 | 0.94524 | $\alpha Syn_{head}$ |
| Glycerophospholipid metabolism | 36 | 18 | 9.94E-07 | 0.69008 | $\alpha Syn_{head}$ |
| Glycine,serine and threonine metabolism | 33 | 16 | 6.93E-06 | 0.75636 | $\alpha Syn_{head}$ |
| Purine metabolism | 70 | 24 | 5.54E-05 | 0.39704 | $\alpha Syn_{head}$ |
| Glyoxylate and dicarboxylate metabolism | 32 | 14 | 1.12E-04 | 0.47167 | $\alpha Syn_{head}$ |
| Glycerolipid metabolism | 16 | 9 | 1.91E-04 | 0.71775 | $\alpha Syn_{head}$ |
| Inositol phosphate metabolism | 30 | 13 | 2.24E-04 | 0.33216 | $\alpha Syn_{head}$ |
| Nitrogen metabolism | 6 | 5 | 4.55E-04 | 1 | $\alpha Syn_{head}$ |
| Butanoate metabolism | 15 | 8 | 7.19E-04 | 0.34921 | $\alpha Syn_{head}$ |
| Pyrimidine metabolism | 39 | 14 | 1.28E-03 | 0.39172 | $\alpha Syn_{head}$ |
| Pantothenate and CoA biosynthesis | 20 | 9 | 1.58E-03 | 0.66327 | $\alpha Syn_{head}$ |
| Sphingolipid metabolism | 32 | 12 | 1.85E-03 | 0.74253 | $\alpha Syn_{head}$ |
| Alanine,aspartate and glutamate metabolism | 28 | 11 | 1.85E-03 | 0.72838 | $\alpha Syn_{head}$ |
| Cysteine and methionine metabolism | 33 | 12 | 2.51E-03 | 0.76449 | $\alpha Syn_{head}$ |
| Arginine biosynthesis | 14 | 6 | 1.32E-02 | 0.3617 | $\alpha Syn_{head}$ |
| Propanoate metabolism | 22 | 8 | 1.33E-02 | 0.35642 | $\alpha Syn_{head}$ |
| Terpenoid backbone biosynthesis | 18 | 7 | 1.37E-02 | 0.68095 | $\alpha Syn_{head}$ |
| Glutathione metabolism | 28 | 9 | 2.07E-02 | 0.44646 | $\alpha Syn_{head}$ |
| Sulfur metabolism | 8 | 4 | 2.37E-02 | 0.53192 | $\alpha Syn_{head}$ |
| Folate biosynthesis | 27 | 8 | 4.59E-02 | 0.40348 | $\alpha Syn_{head}$ |
| One carbon pool by folate | 26 | 16 | 4.66E-09 | 0.58624 | $\alpha Syn_{body}$ |
| Terpenoid backbone biosynthesis | 18 | 11 | 1.59E-06 | 0.87301 | $\alpha Syn_{body}$ |
| Pantothenate and CoA biosynthesis | 20 | 11 | 6.62E-06 | 0.55102 | $\alpha Syn_{body}$ |
| Glyoxylate and dicarboxylate metabolism | 32 | 14 | 1.12E-05 | 0.48667 | $\alpha Syn_{body}$ |
| Pyrimidine metabolism | 39 | 15 | 3.52E-05 | 0.45249 | $\alpha Syn_{body}$ |
| Glycerophospholipid metabolism | 36 | 14 | 5.63E-05 | 0.58098 | $\alpha Syn_{body}$ |
| Glutathione metabolism | 28 | 12 | 6.36E-05 | 0.49056 | $\alpha Syn_{body}$ |
| Arginine biosynthesis | 14 | 8 | 9.29E-05 | 0.42553 | $\alpha Syn_{body}$ |
| Nitrogen metabolism | 6 | 5 | 1.74E-04 | 1 | $\alpha Syn_{body}$ |
| Alanine,aspartate and glutamate metabolism | 28 | 11 | 3.31E-04 | 0.77165 | $\alpha Syn_{body}$ |
| Glycine,serine and threonine metabolism | 33 | 12 | 4.12E-04 | 0.58209 | $\alpha Syn_{body}$ |
| Butanoate metabolism | 15 | 7 | 1.29E-03 | 0.42857 | $\alpha Syn_{body}$ |
| Taurine and hypotaurine metabolism | 8 | 5 | 1.30E-03 | 0.82857 | $\alpha Syn_{body}$ |
| Pyruvate metabolism | 23 | 8 | 5.41E-03 | 0.29379 | $\alpha Syn_{body}$ |
| Phenylalanine,tyrosine and tryptophan biosynthesis | 4 | 3 | 7.41E-03 | 1 | $\alpha Syn_{body}$ |
| Riboflavin metabolism | 4 | 3 | 7.41E-03 | 1 | $\alpha Syn_{body}$ |
| Glycerolipid metabolism | 16 | 6 | 1.06E-02 | 0.38878 | $\alpha Syn_{body}$ |
| Purine metabolism | 70 | 16 | 1.19E-02 | 0.16387 | $\alpha Syn_{body}$ |

| | | | | | |
|---|---|---|---|---|---|
| Folate biosynthesis | 27 | 8 | 1.56E-02 | 0.31237 | $\alpha Syn_{body}$ |
| Arginine and proline metabolism | 36 | 9 | 3.18E-02 | 0.5872 | $\alpha Syn_{body}$ |
| Porphyrin metabolism | 31 | 8 | 3.53E-02 | 0.18789 | $\alpha Syn_{body}$ |
